# Comparative Analysis of RNA Secondary Structure Accuracy on Predicted RNA 3D Models

**DOI:** 10.1101/2022.10.16.512453

**Authors:** Mandar Kulkarni, Jayaraman Thangappan, Indrajit Deb, Sangwook Wu

## Abstract

RNA structure is conformationally dynamic, and accurate all-atom tertiary (3D) structure modeling of RNA remains challenging with the prevailing tools. Secondary structure (2D) information is the standard prerequisite for most RNA 3D modeling. Despite several 2D and 3D structure prediction tools proposed in recent years, one of the challenges is to choose the best combination for accurate RNA 3D structure prediction. Here, we benchmarked seven small RNA PDB structures (40 to 90 nucleotides) with different topologies to understand the effects of different 2D structure predictions on the accuracy of 3D modeling. The current study explores the blind challenge of 2D to 3D conversions and highlights the performances of *de novo* RNA 3D modeling from their predicted 2D structure constraints. Our results show that conformational sampling-based methods such as SimRNA and IsRNA1 depend less on 2D accuracy, whereas motif-based methods account for 2D evidence. Our observations illustrate the disparities in available 3D and 2D prediction methods and may further offer insights into developing topology-specific or family-specific RNA structure prediction pipelines.

## INTRODUCTION

Ribonucleic acids (RNAs) are versatile molecules that leverage the transfer of genetic information, regulation of gene expression, catalytic function, and many other significant roles in cellular metabolism. Unlike protein, RNA is sequentially encoded with only four canonical nucleotides, A, U, C, and G, and some modified nucleotides. To perform its function, RNA must fold into specific secondary (2D) and tertiary (3D) structures that are inherently related to each other. The importance of RNA structures and the challenges associated with experimental determination have briskly promoted computational RNA modeling in recent times. At the 2D level, RNA canonical base pairing covers A-U, G-C, and G-U pairs.

Furthermore, noncanonical base pair inclusion extends the 2D structure repertoire with the formation of pseudoknots^1^. It has been hypothesized that the folding of small RNAs is hierarchical and sequential, i.e., the sequence undergoes canonical 2D structure formation at first, followed by stable 3D structure formation^2,3^. Therefore, predicting accurate RNA 2D structures is the foremost logical step in modeling RNA 3D structures. Multiple approaches have been proposed for RNA 2D structure prediction based on thermodynamic nearest-neighbor parameters, statistical learning, knowledge, and homology. Many machine learning methods are also being utilized to solve RNA 2D structure prediction problems.

On the other hand, RNA tertiary interactions are based on a negatively charged phosphate backbone, sugar moiety, and aromatic heteroatomic bases. Such interactions involve Watson-Crick base pairing, non-Watson-Crick pairing, stacking, base-phosphate interactions, sugar-phosphate interactions, and interactions with microenvironment ions or cosolutes. Structural rearrangements due to these diverse interactions generate multiple kinetic traps, slowing the folding of large RNAs^2,3^. Modeling these interactions, which lead to a rugged free energy landscape with conformationally alternative yet energetically similar folds, confounds definite RNA 3D modeling. RNA 3D modeling tools are classified into conformational sampling-based and motif-based methods. Despite the availability of several 2D and 3D tools, it is often difficult to obtain predicted models that fit native structure homogeneity. It is currently considered possible that the structural prediction precision and correlation between 2D and 3D structure modeling were not established properly. Moreover, no unified 2D or 3D structure prediction methods are suitable for predicting all RNA types.

In this study, in an attempt to assess the performance of available 2D and 3D modeling together, we carried out two stages of evaluations as follows. (A) We performed 2D and 3D modeling prediction of a given RNA sequence using multiple 2D and 3D tools, respectively. (B) The correlation of the predicted models with the native 2D and 3D structures was extracted from experimentally solved structures. The RNA 2D prediction tools used in this study included ViennaRNA 2.0 RNAfold, RNAstructure, NUPACK, LinearFold, UNAFold/MFold, RNAfold_MC-Fold, CONTRAfold, MXfold2, SPOTRNA, UFold, ContextFold, RNAPKplex, HotKnots, pKiss, IPknot, DotKnot, CentroidFold, CentroidHomfold-LAST and R2DT. Similarly, the 3D modeling tools considered here were IsRNA1, SimRNA, and RNAComposer. RNA-Puzzles (https://www.rnapuzzles.org/) has highlighted the important aspects of RNA 3D structure predictions by evaluating the accuracies and limitations of numerous tools. We chose small RNA PDB structures ranging between 40 and 90 nucleotides from RNA Puzzle number 3 (PDB 3OWZ), number 11 (PDB 5LYU), number 21 (PDB 5NWQ), number 25 (PDB 6P2H), and number 29 (PDB 6TB7). Additionally, we included the recent SARS-COV-2 frameshift pseudoknot (PDB 7LYJ) and the classical fluoride riboswitch (PDB 4ENC). Various 2D predictions followed by RNA 3D modeling were performed for all these RNA candidates. Consistently, with the help of the RNA-Puzzles toolkit^4^, the quality and accuracy of the predicted 3D structures were evaluated in terms of clashes between atoms, RMSD, and interaction fidelity network matrix.

The main objective of our work is to display the effect of 2D structure accuracy on the 3D structure modeling of RNAs (<100 nucleotides). Our results show the RNA 2D structure prediction accuracies for each RNA candidate used in this study, followed by correlations between the 2D and 3D modeling predictions. We found some discrepancies and advantages between motif-based and sampling-based 3D modeling and discussed them in detail. Finally, we demonstrated the convergence issues for sampling-based methods. Our analyses correlating the accuracies of the RNA 2D and 3D structural predictions suggest that considering structural topologies and RNA families in the pipeline of RNA structure prediction would lead to improved accuracies.

## MATERIALS AND METHODS

### RNA structures used in this study

Three RNA structures (PDB 5LYU, 6TB7, 7LYJ) were used, in the order of relatively simple (stem–loop) to complex folds (such as pseudoknots and long-range interactions). Then, four RNA structures (PDB 5NWQ, 4ENC, 6P2H, 3OWZ) with relatively complex folds were chosen. The RNA structures were grouped based on their topological complexities to reveal the prediction limitations of the existing tools. The RNA structures examined are listed in **Table 1**.

**Table 1.** List of RNAs used for 2D and 3D structure prediction.

| PDB ID | No of bases | 3D topology | Riboswitch | Pseudoknot | RNA puzzle ID |
| --- | --- | --- | --- | --- | --- |
| 5LYU | 58 | Stem-loop | No | No | 11 |
| 6TB7 | 52 | Three-way junction | Yes | No | 29 |
| 7LYJ | 66 | Pseudoknot | No | Yes | NA |
| 5NWQ | 41 | H-type pseudoknot | Yes | Yes | 21 |
| 4ENC | 52 | Pseudoknot | Yes | Yes | NA |
| 6P2H | 69 | Three-way junction | Yes | No | 25 |
| 3OWZ<sup>a</sup> | 88 | Three-way junction | Yes | No | 3 |

<sup>a</sup>Chain A as reference for 3D structure analysis

Briefly, PDB 5LYU^5^ is a simple stem–loop structure with one internal loop, one bulge, and the tetraloop UUCG. PDB 6TB7^6^ is an NAD+ riboswitch with a tandem bulged stem–loop structure. PDB 7LYJ^7^ is a programmed −1 ribosomal frameshifting pseudoknot of SARS-COV-2, which contains GAAA tetraloop and three planar triplets. PDBs 5NWQ^8^ and 4ENC^9^ are pseudoknot structures with guanidine III riboswitch and fluoride riboswitch, respectively. PDB 5NWQ contains a left-handed triple-helix core. PDB 6P2H^10^ corresponds to a 2’-deoxyguanosine riboswitch with a signature three-way junction. PDB 3OWZ^11^ is a glycine riboswitch. We considered chain A of PDB 3OWZ with the terminal nucleotide changed for this study. RNA-Puzzle 3OWZ (RPZ3) differs from the actual crystallized structure by two terminal nucleotides (GG on the 5’ side and UC on the 3’-side). Additionally, the loop region “CAUAU” in RNA-Puzzle 3 is replaced by “GAAAC” in the crystallized structure. Thus, we considered 88 nucleotides in the crystal structure sequence.

PDB 3OWZ_A:

5’-GGCUCUGGAGAGAACCGUUUAAUCGGUCGCCGAAGGAGCAAGCUCUGCGGAAACGCAGAGUGAAACUCUCAGGCAAAAGGACAGAGUC-3’

RNA-Puzzle 3 sequence:

5’-CUCUGGAGAGAACCGUUUAAUCGGUCGCCGAAGGAGCAAGCUCUGCGCAUAUGCAGAGUGAAACUCUCAGGCAAAAGGACAGAG-3’

A total of 27 secondary (2D) structure prediction tools were utilized (**Table 2)**. The tools were broadly grouped into four major classes: thermodynamics-based, machine learning-based, pseudoknot-based, and homology-based tools. Despite being classified into different categories, some packages have overlapping features, such as RNAPKplex, which takes advantage of thermodynamic constraints to predict pseudoknots^12,13^. All the command-line usages for the local runs are mentioned in the **Supplementary Information**. Most of the tools were used with the default parameters. In a few cases, nondefault parameters were applied. For example, Das and coworkers observed^14^ RNA 2D prediction results that better agreed with the chemical probing data when a high temperature of 60° C was considered. We therefore used 60° C for the RNAfold_MFE (RNAfold_MFE_60) predictions in addition to the standard MFE prediction.

**Table 2.**
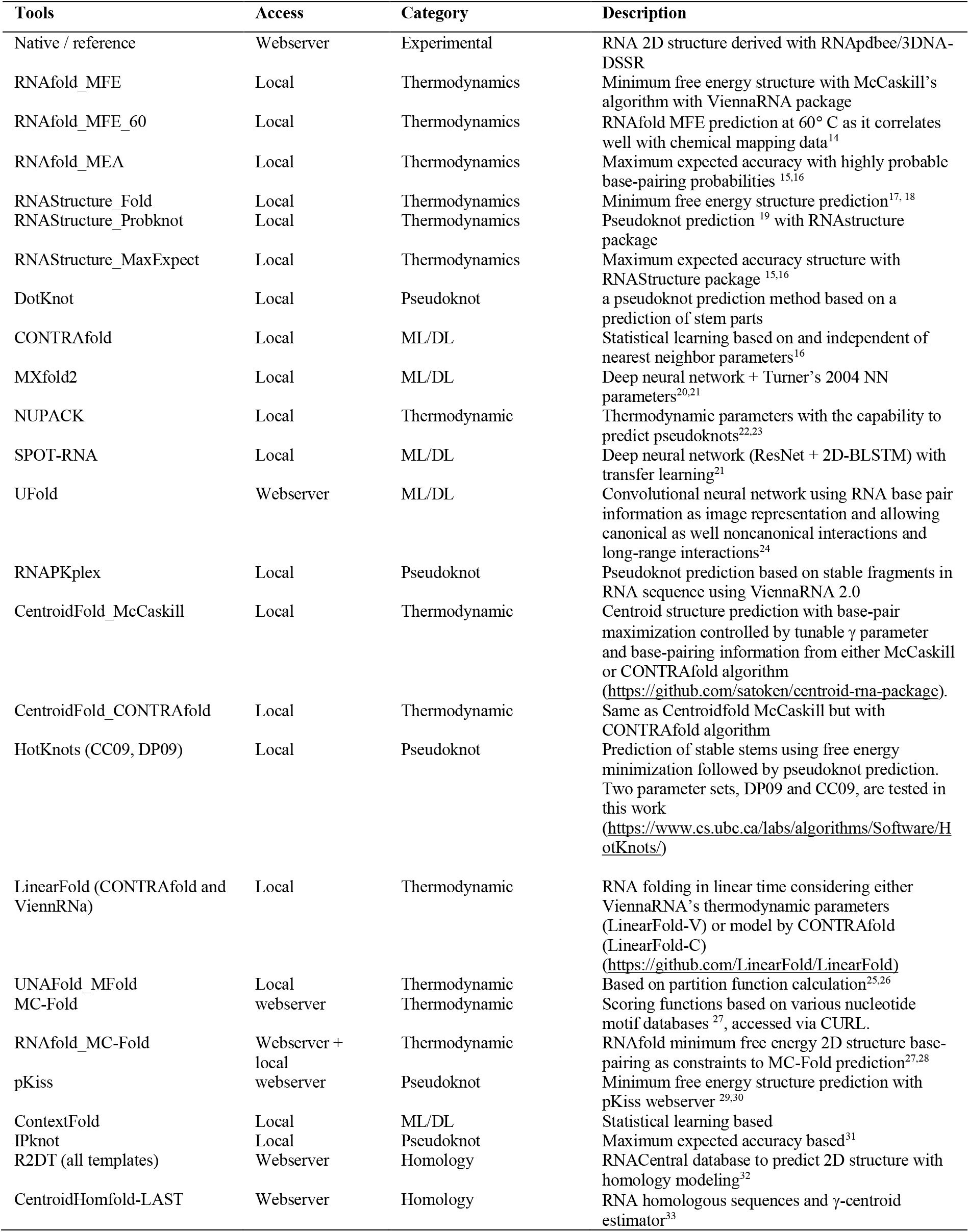
List of RNA 2D structure prediction tools.

### 3D tools

The three software packages used for RNA 3D structure prediction were (1) IsRNA1, (2) RNAComposer, and (3) SimRNA. Detailed descriptions of the usage of each of these tools are summarized below, and corresponding scripts are provided in the **Supplementary Information**.

### 2D Software Packages

### IsRNA1

IsRNA1^34^ combines a coarse-grained force field and replica-exchange simulations to model RNA 3D structures. In IsRNA1, coarse-grained energy functions are inspired by the probability distributions of bonds, angles, torsions, and pairwise distances from experimental PDB structures. The automated 3D structure prediction and clustering protocol proposed by IsRNA1 developers was used in this study without manual intervention. The details of this automated protocol are provided in the **Supplementary Information**. An initial RNA sequence and corresponding 2D structure were provided to predict the 3D structures.

### RNAComposer

The RNAComposer webserver was used in this study. RNAComposer uses known 3D structural motifs that can be accessed through RNA FRABASE 2.0 to build the 3D model based on the user-defined RNA sequence and 2D structural information. RNA FRABASE 2.0 is a web-accessible database^35^ of 2753 (accessed on 31 August 2022) PDB-deposited RNA structures and provides a search engine for the 3D fragments within those PDB-derived structures. RNAComposer can build large 3D structures with up to 500 nucleotides within a short time given an RNA sequence and its predicted 2D structure. We provided the RNA sequence and the predicted 2D structural information to the RNAComposer webserver to build the 3D models. We observed that the modeled 3D structure is sensitive to the input format if pseudoknot or long-range interactions are involved.

Our results show that the RNAComposer webserver is a unique and fast RNA structure building with accuracies equivalent to or sometimes better than sampling-based methods.

### SimRNA

SimRNA utilizes coarse-grained modeling and replica-exchange Monte Carlo (REMC) simulations to predict the RNA 3D structure by conformational sampling. The energy function of SimRNA is based on the base and backbone parameter distribution, including the RNA pseudotorsion η-θ distribution from experimental structures. SimRNA utilizes RNA 2D structural information in the form of distance restraints. We used the standard protocol of REMC simulations for the main results.

However, to understand convergence issues with SimRNA, we performed three sets of repeated REMC simulations with the 2D structures derived from the crystal structures using the RNApdbee _3DNA-DSSR approach, which we otherwise called native 2D structures. Each set consisted of 8 repetitions of the REMC simulations but with ~30% of the standard steps, i.e., 4800000 with the same replica-exchange frequency of 16000. Each REMC simulation consisted of 10 trajectories. Thus, the structures were collected from 80 trajectories (10 replicas x 8 repetitions) for one set. The same clustering protocol was followed to obtain the final centroid structure from the first majorly populated cluster. In the Results section, we have presented all the results from the REMC simulations with the default simulation setup and from all three repeated sets.

### Evaluation of RNA 2D structure prediction tools

We evaluated 28 RNA 2D structure prediction tools, including the RNApdbee_3DNA-DSSR approach, to extract the native 2D structures from the crystal structures. To evaluate the accuracy of the predictions, the positive predictive value (PPV, precision), sensitivity (recall), and F1-score values were calculated as suggested in reference^1^ by comparing the base pairs observed in the predicted structures to those in experimental structures. The definitions of true positive (TP), false positive (FP), and false negative (FN) are summarized in **Table 3**.

**Table 3.** Definition of true positive (TP), false positive (FP), and false-negative (FN)

| Classification | Base pairs in experimental structure | Base pairs in the predicted structure |
| --- | --- | --- |
| TP | Yes | Yes |
| FP | No | Yes |
| FN | Yes | No |

Briefly, the dot-bracket notation was converted to connectivity table (CT) file format using the *dot2ct* program. The PPV and sensitivity values were predicted using the scorer program of RNAStructure software. As shown in Equations (1)–(3), F1-score values were derived using PPV and sensitivity values. In the present work, the scorer program enforced exact base pairing for the PPV and sensitivity calculations. This criterion allows differentiation between the native and near-native 2D structures to better understand the influence of the 2D structural accuracy in translating to accurate 3D structures.

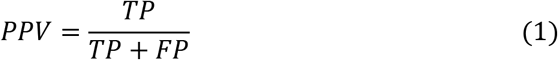

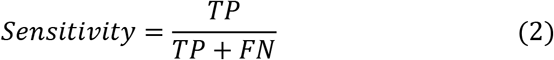

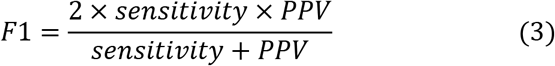

### Assessing RNA 3D structure quality

The predicted 3D models were normalized by the *RNA_normalizer* tool from the RNA-Puzzles toolkit. The normalization of the PDB file altered the modified nucleotides in the PDB to the nearest possible unmodified nucleotides. The residue numbers and chain termination information were preserved, and hydrogens were removed. Although root mean square deviation (RMSD) measurements are widely used, RMSD values may not be the ideal measure to estimate the accuracy of predicted RNA 3D structures compared to the crystal (native) structure.

For example, two RNA structures can have similar all-atom RMSD values with different base orientations. In such cases, the RMSD alone could be ambiguous. Magnus et al.^4^ proposed an RNA-Puzzle Toolkit: a collection of RNA structure assessment tools for prediction benchmarking. Major and coworkers^36^ proposed RNA structure descriptors, deformation index (DI), and interaction network fidelity (INF) parameters. In addition to RMSD, the following metrics (**Table 4**) were considered to evaluate the RNA 3D structure quality and accuracy compared to the crystal (native) structure.

**Table 4.**
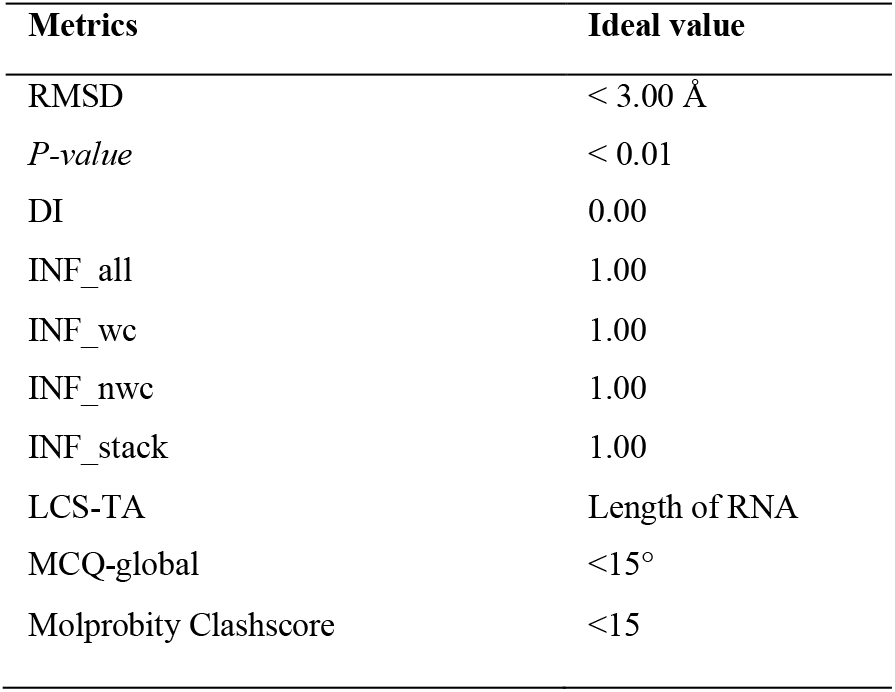
RNA 3D structure assessment metrics.

#### a. Interaction Network Fidelity (INF)

Interaction network fidelity is an important metric that compares the Watson-Crick (WC) and non-Watson-Crick (nWC) base pairs and stacked bases in the modeled structure to those in the native structure (**Table 5**).

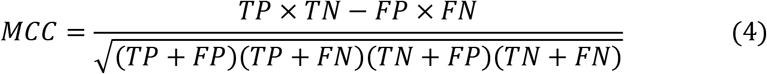

**Table 5.** Definitions of true positive (TP), true negative (TN), false positive (FP), and false-negative (FN).

| Classification | PARAM<sup>a</sup> in Reference PDB | PARAM<sup>a</sup> in modeled PDB |
| --- | --- | --- |
| TP | Y | Y |
| TN | N | N |
| FP | N | Y |
| FN | Y | N |

<sup>a</sup>PARAM = Watson-Crick base pairs, non-Watson-Crick base pairs, base stacking.

Thus, the INF serves as a metric for assessing local 3D structure quality. The MC-annotate program calculated INF and defined it as the Mathews correlation coefficient (MCC) (Equation 4) between the reference structure and the predicted structure.

#### b. Deformation Index (DI)

The deformation index (DI) is the ratio of RMSD to INF_all (Equation 5) and is used to compare global and local structural accuracy. If the RMSD values are similar, DI values help to understand the local structural quality of the 3D models of the sequence of interest.

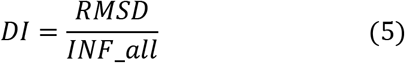

#### c. *P*-value

**The** *P*-value^37^ for the predicted RMSD is independent of RNA size, and we chose a cutoff value of less than 0.01, similar to previous studies, to decide on a successful prediction. The *P*-value is based on the range of the ideal RMSD values for a given length of an RNA. It is an important measure for understanding the significance of high RMSD values. The ideal RMSD value for an RNA of length N is given as

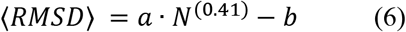

where a = 0.51 and b=19.80 if 2D structural constraints are applied for 3D structure prediction. If *m* is the RMSD of the predicted 3D model with respect to the reference structure, the *P*-value is calculated as

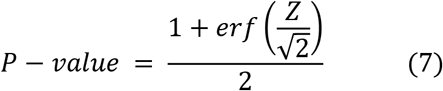

where 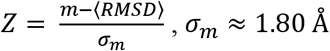

#### d. Clashscore

The clashscore indicates the number of serious steric clashes in an RNA structure model per 1000 atoms. We used the MolProbity software package to determine the clashscores, and a probe radius of 0.40 Å was used for the calculation.

#### e. Mean Circular Quantities (MCQ)

The mean of circular quantities (MCQ)^38^ is a metric to compare RNA 3D structures in the torsion angle space. The residue-wise comparison in the torsion angle space highlights the dissimilarity in the local structure. An MCQ score of less than 15° suggests a higher similarity with the native or reference structure.

#### f. Longest Continuous Segments in Torsion Angle space (LCS-TA)

LCS-TA^39^ identifies the longest continuous segments that display local similarity in the torsion angle space.

### Molecular Dynamics (MD) simulations

For PDB 4ENC, the SimRNA simulations with the native RNA 2D structural restraints predicted a high-RMSD (19.65 Å) structure. We performed 2 μs long simulations using the SimRNA-predicted 3D model of PDB 4ENC as a starting structure to understand the efficiency of long MD simulations in improving RNA structure. Many recent studies have pointed out that the present RNA force fields exhibit an imbalance in nonbonded interactions^40^ and torsional parameters^41^. Additionally, conventional RNA simulations do not assure reliable and converged conformational sampling^40,42^. Thus, MD simulations with current RNA force fields and conformational sampling limitations do not guarantee a substantial improvement of 3D predictions that are structurally distant from the native state. However, with the present 2 μs simulations, we have attempted to estimate the effect of conventional MD simulations with different water models on the RMSD values and alterations of RNA structures that are distant from the native state, specific to the case study of the fluoride riboswitch (PDB 4ENC).

Three different sets of simulations were performed with three water models, TIP3P, OPC, and SPC/E, and the AMBER OL3^43–46^ RNA force field. Na^+^ ions were added for charge neutralization, and NaCl salt was used to obtain a 0.15 M concentration. Each water model-specific Joung-Cheatham ion parameter^47^ was used for Na^+^ and Cl^-^ ions. The RNA molecule was placed in a dodecahedron box, maintaining a minimum distance of 10 Å from the box edges. The hydrogen mass repartitioning scheme^48^ was applied with a 4 fs timestep. First, the system was minimized to remove any clashes. Then, the whole system was slowly heated to 300 K within 100 ps. The simulation was continued for another 100 ps with restraints on the heavy atoms of RNA to allow the equilibration of water and ions. At this equilibration stage, the Berendsen thermostat and barostat were used to maintain temperature (300 K) and pressure (1 bar), respectively. Next, a 200 ps NPT equilibration was performed without restraints before the 2 μs production run simulations. The neighbor list cutoff was set to 10 Å. The PME method was applied for electrostatics with a cutoff of 10 Å. The van der Waals interactions were also truncated at 10 Å. Long-range corrections were applied to pressure and energy. All bonds were constrained with the LINCS algorithm. RNA and solvent ions were coupled separately to a V-rescale thermostat (300 K, coupling constant 0.10 ps) and a Parrinello-Rahman barostat to maintain pressure (1 bar, coupling constant 1 ps). The coordinates were saved every 10 ps.

## RESULTS

### Accuracy of RNA 2D structure predictions

First, the accuracy of the predictions of RNA 2D structures using various tools (**Supplementary Tables S2-S8, Figures S1-S7**) was evaluated to establish the correlation between 2D and 3D predictions. As described in the Methods section, the metrics for evaluating the accuracy of the RNA 2D structure predictions, such as PPV, sensitivity, and F1-score, were calculated for the seven RNA candidates (**Table 1**). The distributions of the PPV, sensitivity and F1-score from all 2D prediction tools (**Table 2)** are plotted in **Figure 1**.

**Figure 1.**
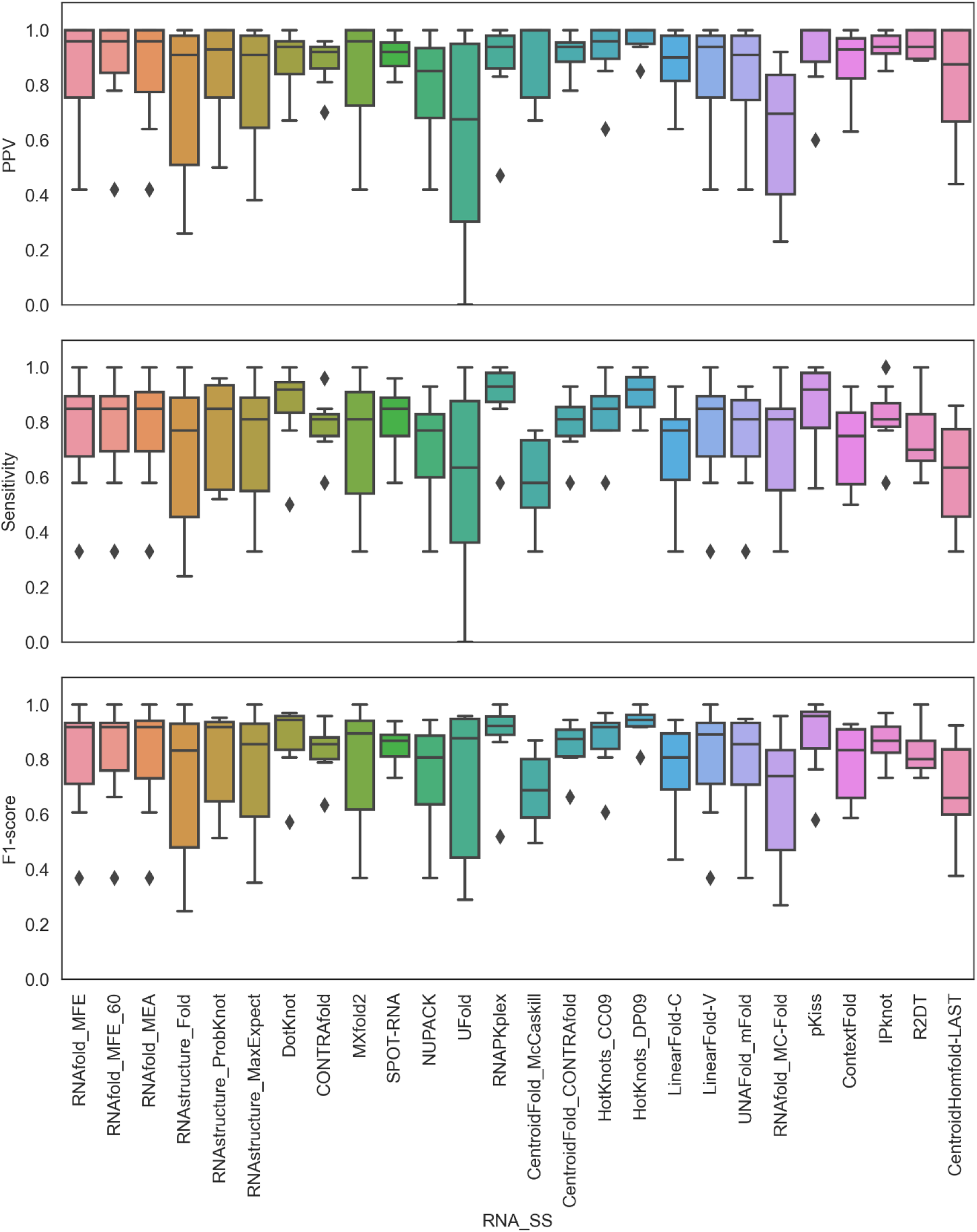
Distributions of the PPV, sensitivity, and F1-score values for each of the seven RNA candidates under study. The predicted 2D structures were compared to their corresponding native structures. These metrics range from 0 to 1, and a higher value indicates better prediction. Outliers are shown as diamond shape points and labeled with the PDB IDs.

In general, similar trends were observed for the medians of the PPV, sensitivity, and F1-score distributions (**Figure 1**). The distributions were found to be negatively skewed for most of the tools, indicating a central tendency to predict accurate 2D structures for most of the RNAs under study. Notably, for R2DT and CentroidHomefold-LAST, the distributions were found to be positively skewed, showing their limitations in predictions, at least for the RNAs under study. Considering all 27 prediction tools and 7 RNAs, we observed a high correlation (0.80) between the PPV and sensitivity metrics, with correlation coefficients of 0.92 and 0.97, respectively, with the F1 scores (**Supplementary Figures S8 and S12**). Ideally, F1-scores, a harmonic mean of PPV and sensitivity, should capture a substantial variation between them. However, it would be interesting to check the extent of variation between PPV and sensitivity that would significantly impact the F1-score. For instance, Ufold’s median F1-score was ~0.90, while the median PPV and sensitivity were ~0.60. Similar results were observed for RNAStructure Probknot. In both cases, for most RNAs, the base pairings were incorrectly predicted (quantified by PPV), and the native base pairs were missed (quantified by sensitivity) in the prediction. These observations can be attributed to the broad interquartile ranges for the PPV and sensitivity calculated for the seven RNAs with simple to complex folds.

In contrast, the median F1-score for RNAstructure_Fold was greater than 0.80, and the median PPV and median sensitivity were greater than 0.90 and 0.70, respectively. RNAstructure_MaxExpect showed similar trends. Interestingly, predictions by DotKnot, CONTRAfold, RNAPKplex, CentroidFold_CONTRAfold, HotKnots_CC09, HotKnots_DP09, IPknot, and SPOT-RNA had a median F1-score value of ~0.90 with narrow interquartile ranges. RNAfold_MFE, RNAfold_MFE_60, RNAfold_MEA, and pKiss predictions displayed a high median F1-score, while the interquartile range illustrated that the prediction accuracy may vary depending on the complexity of the RNA structure.

In general, our results suggest that the predictive accuracy of DotKnot, CONTRAfold, RNAPKplex, CentroidFold_CONTRAfold, HotKnots_CC09, HotKnots_DP09, IPknot, and SPOT-RNA are similar for the range of structurally diverse RNA sequences studied in this work.

### Relationship between predicted RNA 2D and 3D structural accuracies

First, the RNA 2D structure prediction accuracy metric F1-scores were clustered as follows: (a) high (F1-score >= 0.95), (b) medium (0.80 < F1-score < 0.95), and (c) low (F1-score =< 0.80) to understand the effect of prediction accuracy on the 3D modeling results. Subsequently, those F1-score clusters were compared against three important 3D metrics (INF_stack, INF_wc, RMSD), considering the results of all three 3D tools for the seven RNAs used in this study (**Figure 2a**). A gradual increase in the median RMSD values was observed for the 2D tools of the high, medium, and low F1-score groups. Similar results were observed for INF_all and INF_stack, where the groups of 2D tools with higher F1-scores predicted 3D structures with higher median INF_stack and INF_wc values compared to those with lower F1-scores. Notably, considering all 27 2D structure prediction tools and three 3D modeling tools applied to seven RNAs, we calculated correlation coefficients of −0.46, 0.92, and 0.53 for RMSD, INF_wc, and INF_stack values against the F1-scores (**Supplementary Figure S12**). These observations, in general, suggest that 2D structures predicted with higher local base pairing accuracy translate to 3D models with higher accuracy for the stem parts of RNA, irrespective of the 3D modeling approaches under study.

**Figure 2.**
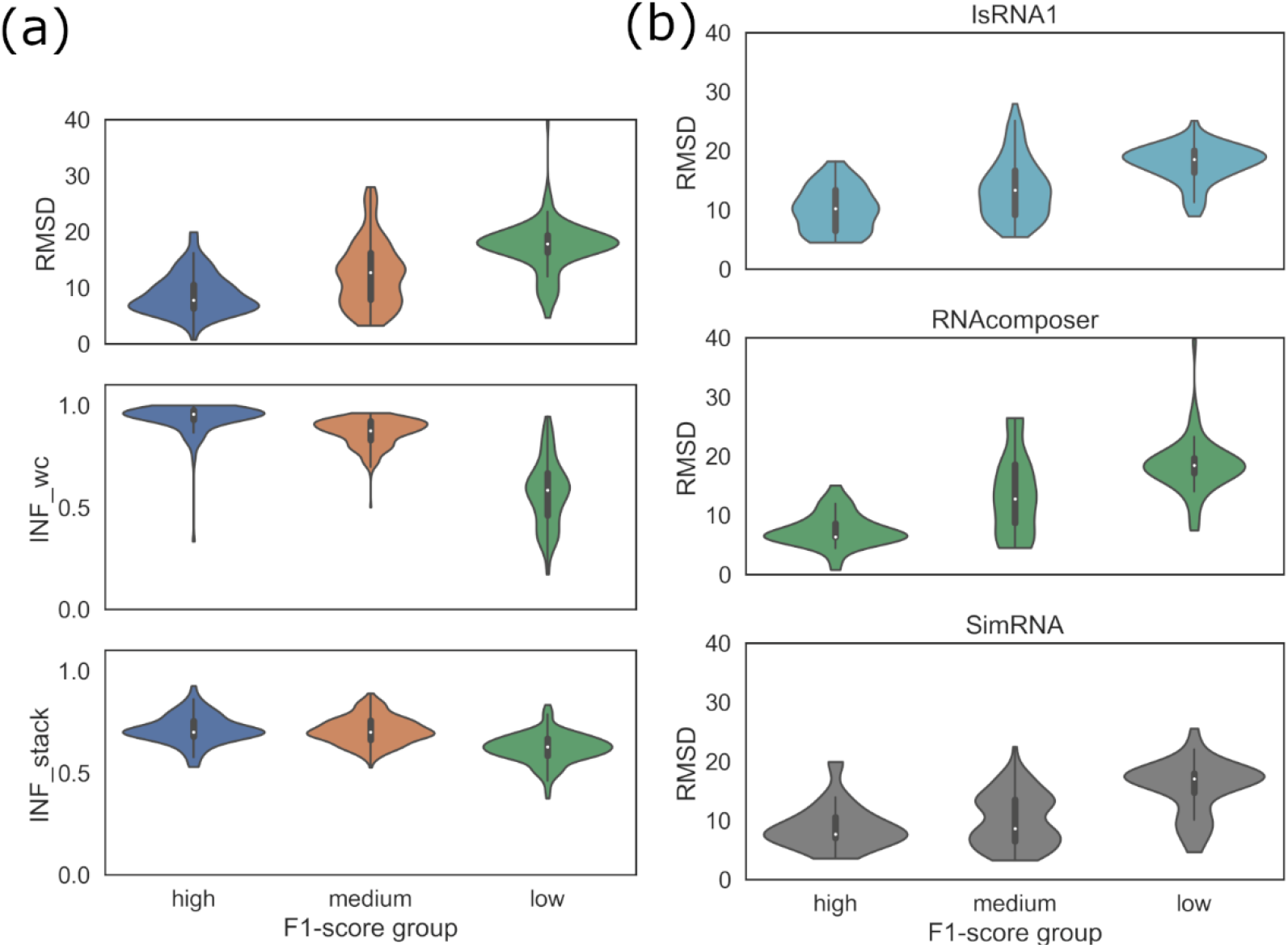
(a) Distributions of the RMSD, INF_wc, INF_stack values, (b) method-specific RMSD values for high, medium, and low F1-score groups.

However, 2D structures with higher local base pairing accuracy may not ensure the accurate prediction of base stacking in 3D models. In the case of the deformation index (DI), a correlation coefficient of −0.65 was calculated against the F1-scores (**Supplementary Figure S12**). This observation and the weak correlations of the F1-scores with the clashscore metric and the mean circular quantity (MCQ) and longest continuous segments in torsion angle space (LCS-TA) metrics indicate that the accurate translation of 2D structures to 3D models may not be guided by the accurate 2D structural inputs (base pairing) for the prediction of 3D models; rather, 2D to 3D translation largely depends on the 3D modeling approach. To examine this further, we compared the F1-score clusters against the distribution of RMSDs for three different 3D tools, IsRNA1, RNAComposer, and SimRNA (**Figure 2b**). At the individual package level, we also observed a gradual increase in the median RMSD values for the 2D tools of the high, medium, and low F1-score groups.

High F1-score 2D structural inputs led to 3D structures with a median RMSD of less than 10 Å with all three methods, IsRNA1, RNAComposer, and SimRNA. At the same time, a few models from IsRNA1 and SimRNA also tended to have relatively higher RMSD values, even with a high F1 score. Interestingly, for SimRNA, the medium F1-score displayed a bimodal RMSD distribution arising from low and high RMSD values. **Figure 2b** reveals that the RMSD vs. F1-score groups among 3D tools reflected that lower F1-score of each tool correlated with higher RMSDs. Moreover, a few predicted 3D models with IsRNA1 and SimRNA also tended to have higher RMSD values (less accurate models) for a high F1-score (accurate 2D structure). Additionally, the observed low F1-score of each tool correlated with higher RMSDs. For example, IsRNA1-, RNAComposer-, and SimRNA-modeled structures with high F1-score 2D structural inputs showed a median RMSD of less than 10 Å.

Furthermore, to correlate, the 2D and 3D modeling accuracies, the RMSD, deformation index, interaction network fidelity parameters, MCQ, and LCS-TA of each candidate were investigated with the F1-score. Specifically, two clusters were chosen for the F1-score: (a) F1-score >= 0.95 (**left, Figure 3**), and (b) F1-score < 0.95 (**right, Figure 3**). Therefore, RNAs were also grouped into *Group I* (5LYU, 6TB7, and 7LYJ) and *Group II* (5NWQ, 4ENC, 6P2H, 3OWZ_A) to demonstrate their structural complexities.

**Figure 3.**
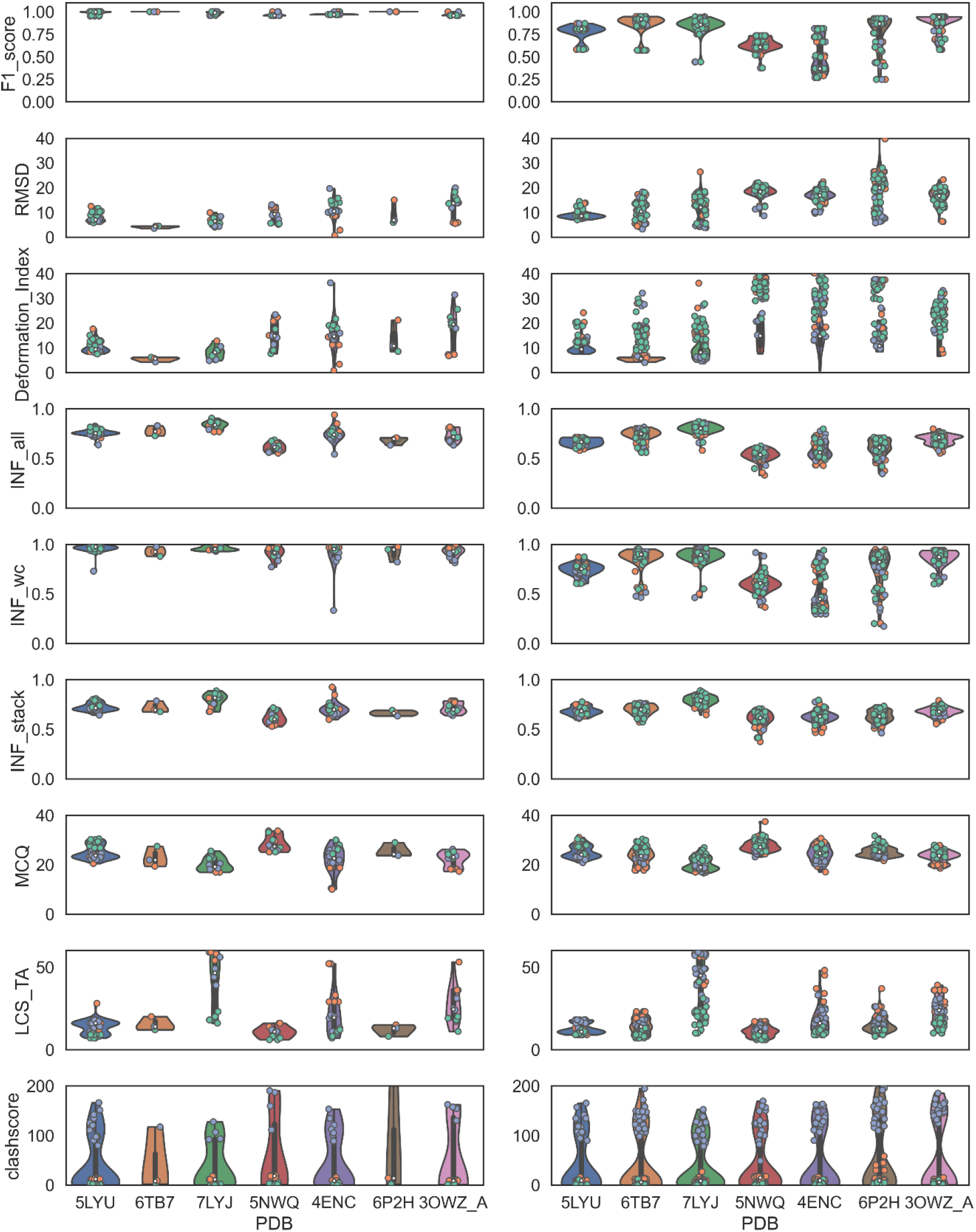
RMSD, deformation index, INF parameters, MCQ, and LCS-TA values for F1-scores greater than or equal to 0.95 (left) and less than 0.95 (right). The color codes are IsRNA1: green, RNAComposer: orange, and SimRNA: violet. Median values are indicated by white dots.

Although 5LYU from *Group I* with high and low F1-scores displayed similar median RMSD values, high F1-scores had median RMSDs lower than 10 Å for 6TB7 and 7LYJ with some outliers. Interestingly, INF_all and INF_wc values for 5LYU showed strong differences based on F1-scores. Meanwhile, INF_all and INF_wc values of 6TB7 and 7LYJ showed similar median values regardless of F1-scores. This observation explained the better INF_all and INF_wc values for simpler 3D topologies such as 6TB7 and 7LYJ. MCQ and LCS_TA were good indicators of 3D structural accuracy in RNA torsions. Overall, for *Group I*, the MCQ values were similar, with overlapping value distributions from5LYU to 7LYJ for low and high F1-scores. It should be noted that none of the MCQ values dropped below the ideal value of 15°. The striking difference of 5LYU became apparent with the LCS-TA values obtained a high F1-score and with a low F1-score, signifying more accurate torsion angles and similar RNA folds.

For the rest of the *Group I and II* RNAs, the LCS-TA values were largely condensed within a distinct range. For Group II RNAs, 5NWQ and 4ENC demonstrated a clear partition between RMSD, INF_all, and INF_wc parameter value distributions in the low and high F1-score categories. Overall, all such observations supported the importance of the correct base-pairing information from the 2D structure, which could manifest as correct Watson-Crick base pairing, ultimately affecting the interaction network fidelity value and RMSD. These results confirmed that accurate Watson-Crick base pairing at the 2D structure level significantly affected complex RNA folds. Stacking patterns (INF_stack) and torsional parameters such as MCQ and LCS-TA did not affect the 2D structures.

In **Figure 4**, we present scatter plots of various accuracy parameters, such as F1-score, RMSD, INF_all, INF_wc, MCQ, and LCS-TA, to understand the relationships between them. Furthermore, package-specific data are colored according to the 3D tools: IsRNA1, RNAComposer, and SimRNA data are shown in *green, blue, and orange*, respectively (**Figure 4)**. The F1-score (*2D structure accuracy parameter*) and INF_wc (*accuracy parameter of predicted 3D model*) demonstrated a linear relationship with a correlation coefficient of 0.92 (**Supplementary Figure S12**). Similarly, the distributions of INF_wc and F1-score against RMSD were nearly identical. The RMSD of structures can vary up to ~ 12 Å with high INF_wc values. The RMSD values increased sharply to 25 Å when the INF_wc values decreased below 0.50.

**Figure 4.**
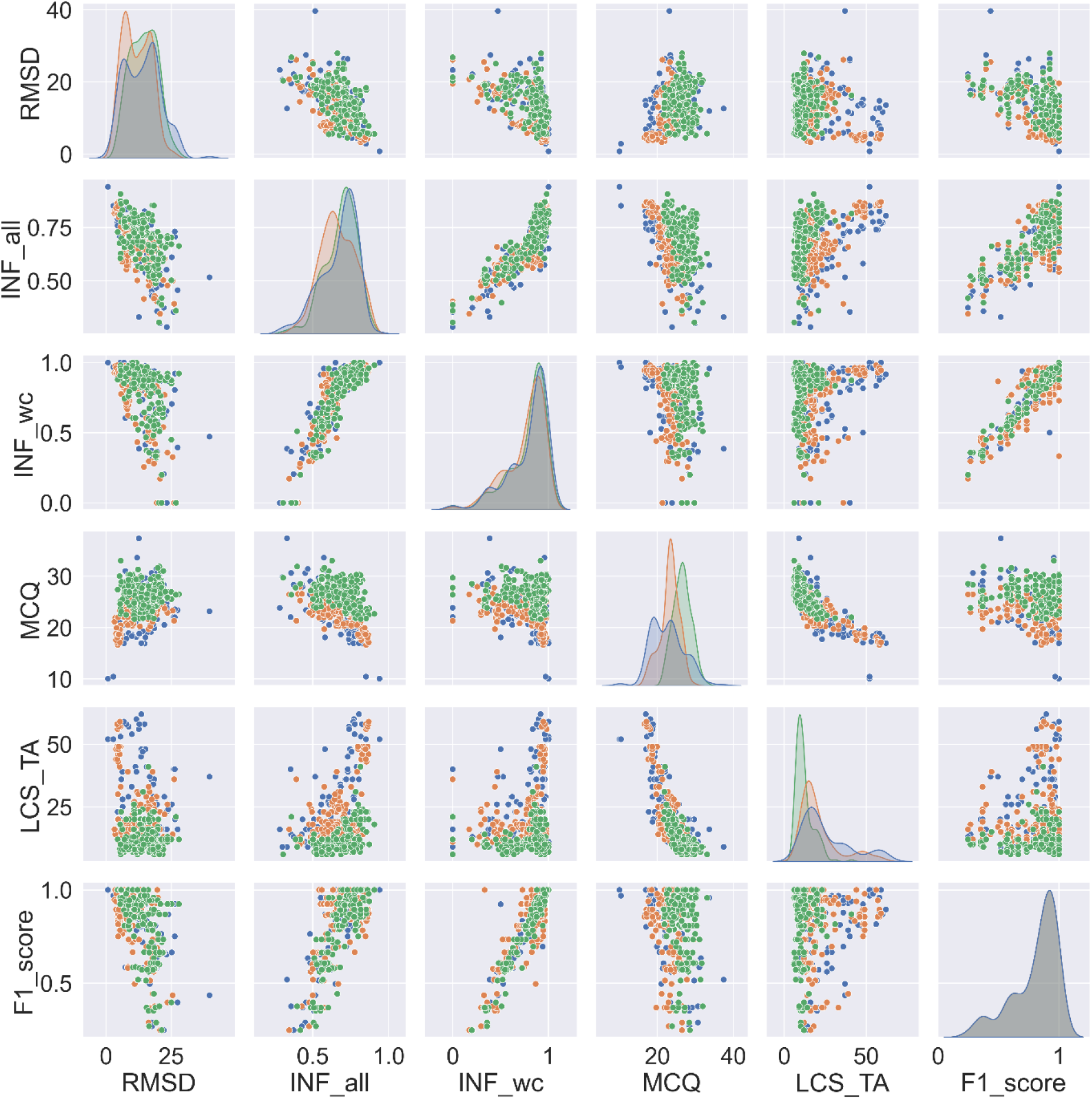
Pair plot distribution of F1-score values and RNA 3D accuracy parameters RMSD, INF_all, INF_wc, MCQ, and LCS-TA for RNAComposer (blue), SimRNA (orange), and IsRNA1 (green) methods.

INF_all and F1-score also showed correlation (cc = 0.74), as Watson-Crick interactions contribute to INF_all. For the IsRNA1 simulations (*green*), the MCQ values were higher (20-30); nonetheless, similar F1-score or INF_wc or INF_all values were observed. Compared with IsRNA1, RNAComposer and SimRNA produced moderately low MCQ values, representing better predictions in torsional space. RMSD vs. LCS-TA and MCQ vs. LCS-TA distribution plots further showed a native torsion correlation for SimRNA (*orange*), and RNAComposer (*blue*) predicted 3D models with RMSDs less than 10 Å.

To reflect the RMSD dependencies of all the 3D models on 2D structure prediction accuracy, the RMSDs for each 2D tool were plotted (**Figure 5**). Since simple and complex folds were included in our dataset, we believed that the median RMSD values would be a good indicator for understanding general trends.

**Figure 5.**
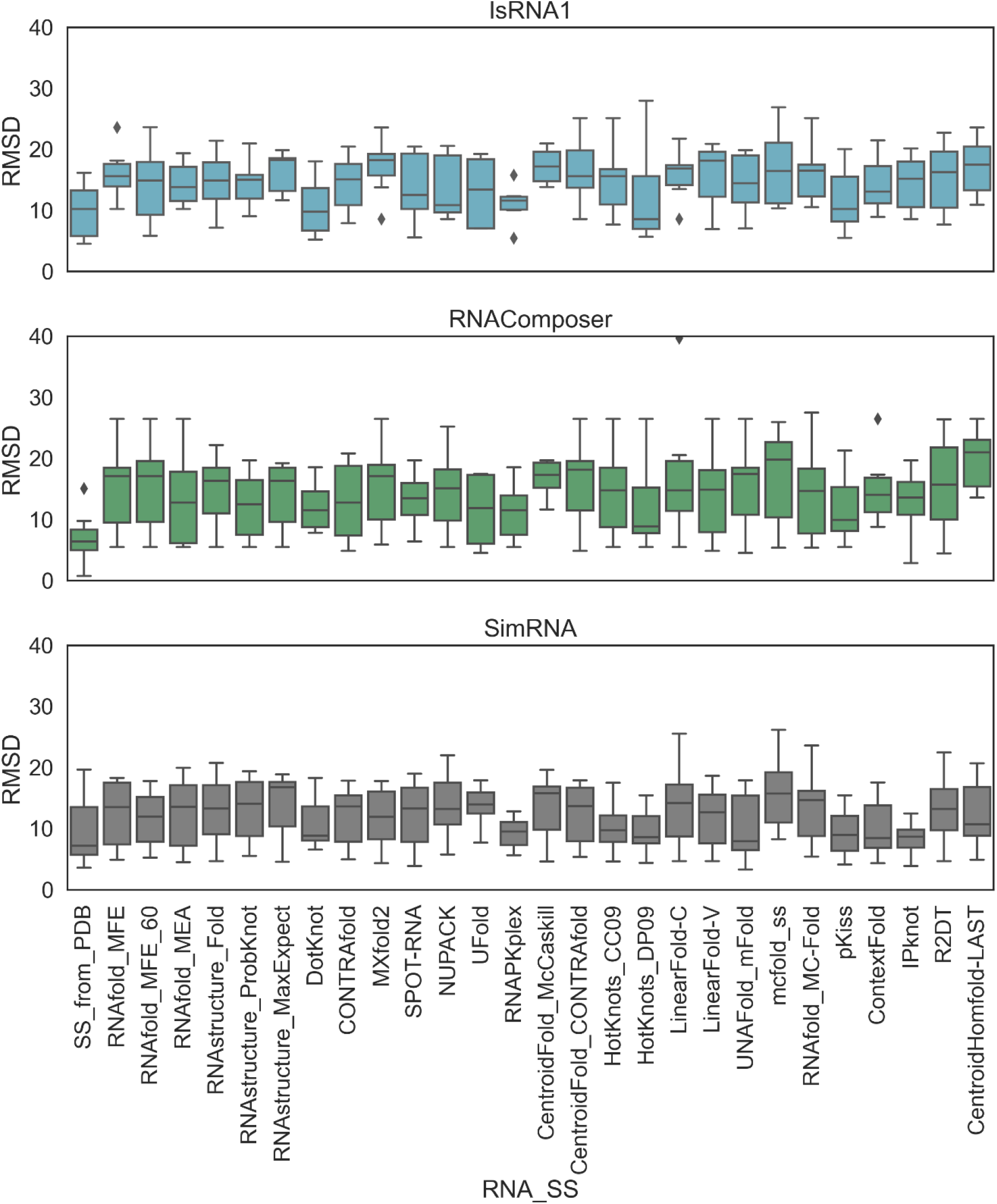
RMSD values of all seven PDBs determined with (a) IsRNA1 simulations, (b) the RNAComposer web server, and (c) SimRNA simulations with 2D structural constraints based on various methods (x-axis).

When only the native (SS_from_PDB) 2D structure was given, RNAComposer, which is widely considered a motif-based method, consistently produced lower RMSDs (< 10 Å) for both simple and complex folds. In contrast, the sampling-based methods, SimRNA and IsRNA1, also produced lower median RMSDs for other 2D structure methods. In particular, SimRNA often produced lower median RMSDs (< 10 Å), as several RMSD values of approximately 5 Å were observed. RNAPKplex 2D structural inputs produced lower RMSDs for both the IsRNA1 and SimRNA models. However, we also observed that sampling-based methods produced higher RMSDs even when the native 2D structural inputs were provided. We found that RNA 3D modeling with sampling-based methods could generate higher-RMSD structures due to simulation sampling issues. Thus, native 2D structural information did not always guarantee better 3D structures for sampling-based methods. These methods depend on simulation sampling, complex folds in RNAs, and the underlying effect of force fields. The results of the 3D structures with native 2D structural inputs are shown below in **Figure 6**. In addition, we observed that even when the 2D structure input differed from the native or reference 2D structure, different algorithm-based 2D predictions could impact 3D structure modeling.

**Figure 6.**
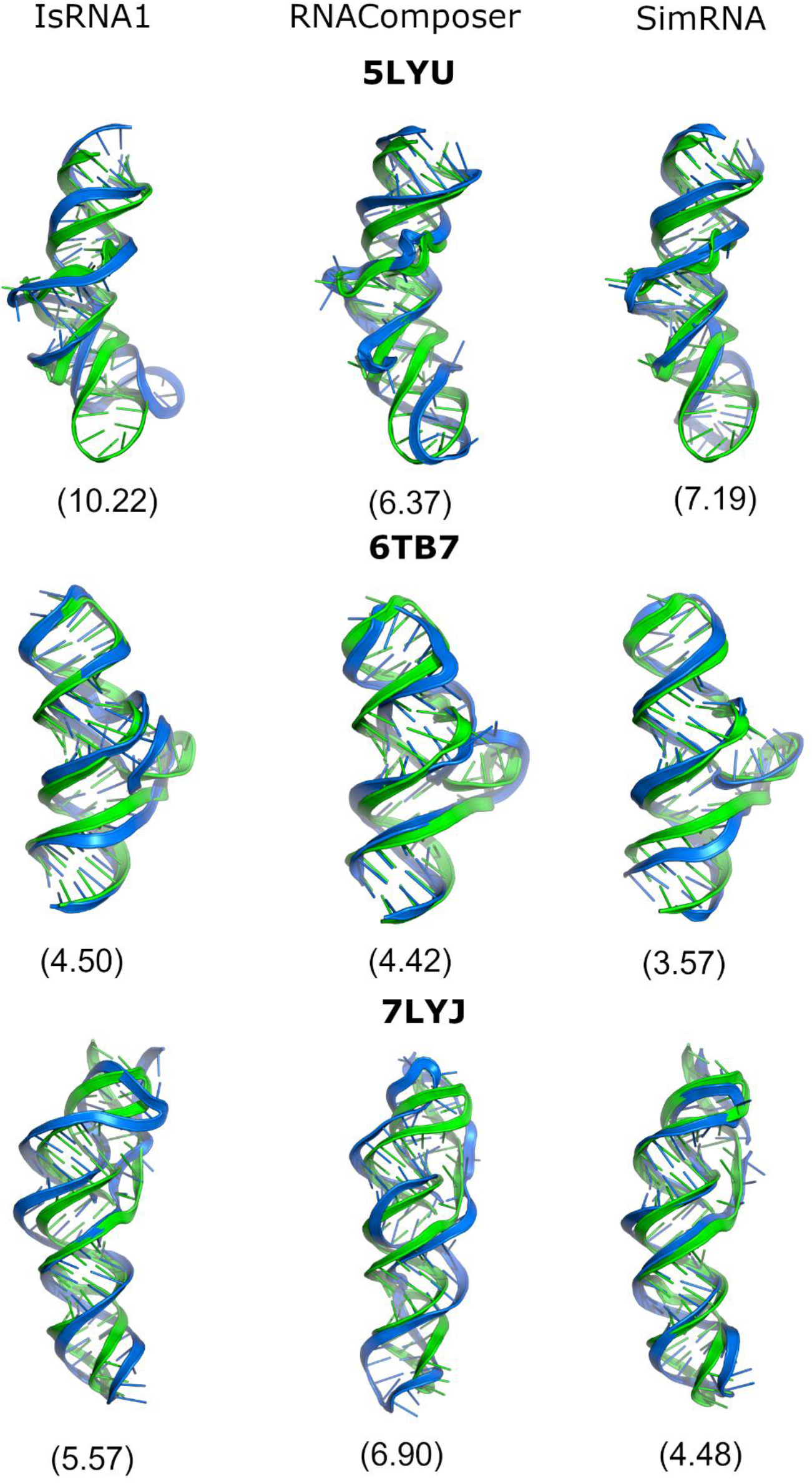
Predicted 3D structures (blue) aligned with the native 3D structure (green) for IsRNA1 (left), RNAComposer (middle), and SimRNA (right) with native 2D structural constraints for Group I PDBs (5LYU, 6TB7, 7LYJ). The RMSDs are mentioned in parentheses.

### Accuracy of predicted 3D structures with native 2D structural input

We systematically tested the influence of native 2D structural input alone on the accuracy of the 3D modeling outcomes of Group I and II RNAs (**Figures 6–7, Supplementary Figures S13-S96**). Reasonably, Group I RNA models displayed lower RMSDs with all three methods than Group II RNAs. The predicted models of IsRNA1 and RNAComposer for Group I RNAs demonstrated low clashscores, whereas SimRNA-predicted models had very high prediction clashscores (>100). We have provided the results case by case as follows.

**Figure 7.**
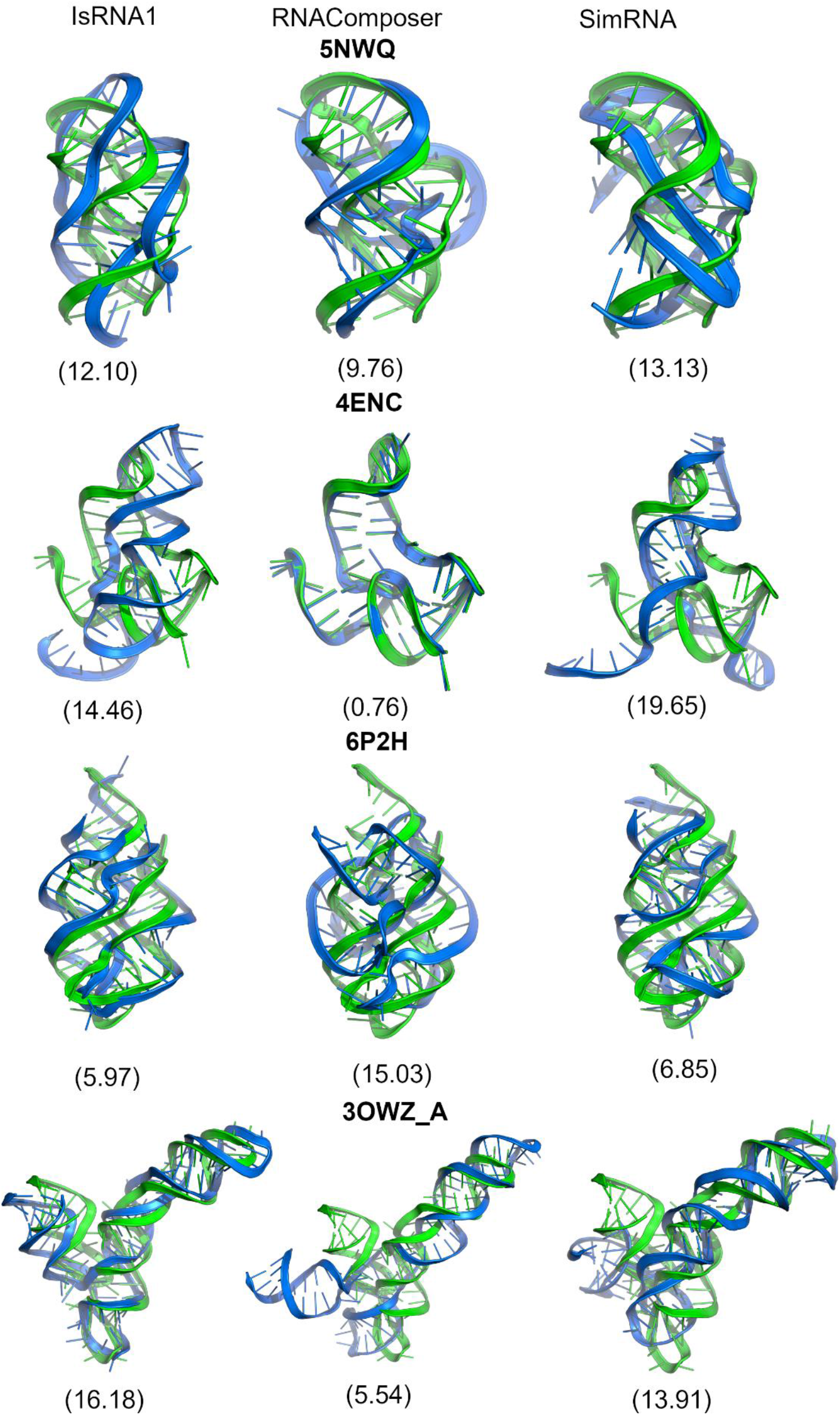
Predicted 3D structures (blue) aligned with native 3D structure (green) for IsRNA1 (left), RNAComposer (middle), and SimRNA (right) with native 2D structural constraints for group II PDBs (5NWQ, 4ENC, 6P2H, 3OWZ (chain A)). The RMSDs are given in parentheses.

For 5LYU, the IsRNA1 RMSD was higher than those of RNAComposer and SimRNA due to the differences in the loop regions. We observed a more bent structure and heterogeneity in the loop, possibly arising from the deviation in torsion values relative to the native structure. Thus, the highest MCQ (29.23) and lowest LCS-TA (10) values were observed. In the case of 6TB7 with SimRNA, the lowest RMSD of 3.57 Å and the highest INF_wc (0.98) value were obtained, albeit with intermediate LCS-TA and MCQ values. The IsRNA1 and RNAComposer models had lower RMSDs. For 7LYJ, the SimRNA prediction produced the lowest RMSD. IsRNA1 and RNAComposer produced similar RMSDs, but significant differences in the LCS-TA (IsRNA1: 18, SimRNA: 56) and clashscore values (IsRNA1: 0.47, SimRNA: 106.40) were observed. This illustrated that the torsional values could differ among similar RMSD structures.

We then checked the effect of native 2D structures on the 3D modeling of complex *Group II* RNA folds such as 5NWQ, 4ENC, 6P2H, and 3OWZ_A, as these were relatively challenging structures for 3D modeling. For 5NWQ, the IsRNA1 and SimRNA RMSD values were similar, but the SimRNA Model *p-value* (0.03) indicated a lack of confidence. RNAComposer achieved the lowest RMSD for 4ENC (0.76 Å), which reflected that their RNA fragments were typically selected from the PDB 4ENC as part of FRABASE 2.0. SimRNA and IsRNA1 modeling both exhibited higher RMSDs, and SimRNA had a higher deformation index and *p-value* for 4ENC.

Due to the differences in their terminal ends, the models showed overall high RMSDs. To verify this, we performed a 2 μs long molecular dynamics simulation with the SimRNA prediction as a starting structure; however, the overall heavy-atom RMSD remained approximately 16 Å, as described in detail in the next result section. For 6P2H, lower RMSDs were observed for the sampling-based methods with lower INF_stack values (~ 0.60), suggesting their base stacking orientations are inaccurate to the native topology. However, a different topology was observed for RNAComposer with an RMSD of 15.03 Å and a *p-value* less than 0.01. For 3OWZ (chain A), the RNAComposer RMSD values were lower than those of the other tools. Despite the terminal end differences, their MCQ (17.27) and LCS-TA (53.00) values and Watson-Crick base-pairing (INF_wc = 1.00) were consistent. IsRNA1 and SimRNA for 3OWZ_A structures had RMSD *p* values below a significance level of 0.01 and lower LCS-TA values (IsRNA1: 19.00 and SimRNA: 24.00).

Overall, none of the methods could generate lower RMSD models for 5NWQ owing to its pseudoknot formation. The lowest RMSD of 4ENC with RNAComposer only proved the fragmentation assembly. Except for RNA 6P2H, the IsRNA1 and SimRNA models produced higher RMSD structures than RNAComposer. Again, the complex folding of 3OWZ_A was apparent with sampling-based methods.

### Effect of long MD simulation

As mentioned above, the 4ENC model from SimRNA showed a higher RMSD than the native 3D structure. We sought to confirm the misaligned loop regions and extended single strand in the terminal region through MD simulation. We performed a 2 μs long simulation with different water models (TIP3P, SPC/E, and OPC), and no variation in RMSD values was found, as presented below in **Figure 8**.

**Figure 8.**
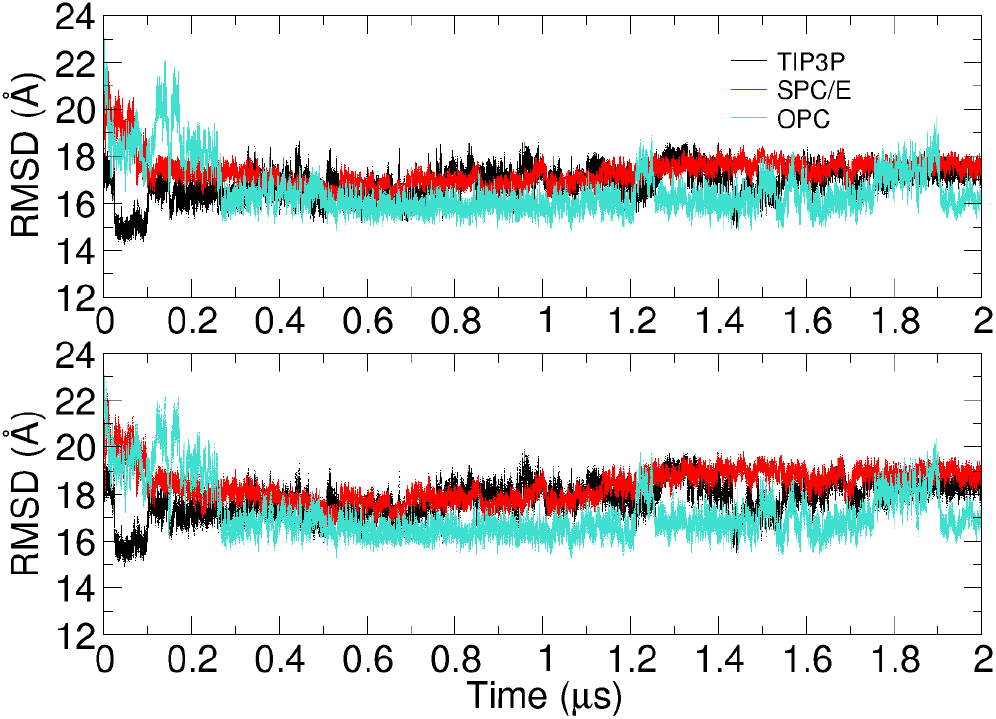
Heavy atom RMSD (top) and backbone RMSD (bottom) to the native 4ENC structure with AMBER OL3 and different water models (TIP3P (black), SPC/E (red), and OPC (green)).

## DISCUSSION

The heatmaps showing RNA 3D structure accuracy parameters are provided in the **Supplementary Information** (**Supplementary Figures S17-S24, S29-S36, S41-S48, S53-S60, S65-S72, S77-S84, S89-S96**), and the general discussion is provided below. RNA 2D structures provide base-pairing information that can act as distance restraints for sampling-based methods, such as IsRNA1 and SimRNA, for efficient conformational sampling. However, conformational sampling of RNA molecules becomes challenging with increasing complexity of RNA folds and increasing sequence length even with correct 2D structural information as input. PDB 3OWZ in our work (RNA-Puzzle 3 with 88 nucleotides) shows that even with the correct RNA 2D structural restraints, sampling-based methods could not fold to the correct topology and produced high RMSDs compared to the native crystal structure.

### IsRNA1

#### PDB 3OWZ_A

For 3OWZ_A, none of the 2D structures exactly matched the native base-paired 2D structure, as shown in **Table S8** and **Figure S7** of the **Supplementary Information**. RNAfold_MEA, CONTRAfold, and MXfold2 predicted close to native 2D structures with F1-scores of 0.959 for each of them, and with these three 2D structural constraints, higher RMSDs (13.60 Å, 15.05 Å, and 18.22 Å, respectively) were obtained (**Supplementary Figure S89**). The combination of the standard IsRNA1 protocol with 2D structural constraints from NUPACK and RNAPKplex provided the lowest RMSDs of all trials, 10.69 Å and 12.17 Å, respectively (**Supplementary Figure S89**). These 2D structures differed from the native 2D structure at the active site bulge and the four-nucleotide bulge defined by G8-A11 (**Supplementary Figure S7**). With these less accurate 2D structural constraints, IsRNA1 provided lower RMSDs for the 3OWZ_A riboswitch, suggesting its lower dependency on the 2D structural accuracy.

#### PDB_4ENC

In the case of 4ENC, among all methods, the 3D structure based on the MC-Fold-derived 2D structure provided the lowest RMSD structure, albeit 12.53 Å relative to the native all-atom structure (**Supplementary Figure S65**). The MC-Fold-derived 2D structure has noncanonical base pairs, G29-A32, U23-U38, and U7-U48, along with incorrect pseudoknot prediction (**Supplementary Table S6 and Figure S5**). In the native structure, a pseudoknot is formed by pairing G2-G3-C4-G5 of the 5’-overhang segment and residues C14-G15-C16-C17 of the internal loop. However, in MC-Fold, the pseudoknot is formed by bases G1-G2-G3-C4-G5 and C14-G15-C16-C17-C18. Notably, this completely different pattern of base pairing and pseudoknot led to a low-RMSD structure. The MC-Fold 2D structure maintained all native base pairing and predicted extra noncanonical base pairs (**Supplementary Table S6 and Figure S5**). We hypothesize that these extra base pairs restricted the conformational sampling extensively in IsRNA1, reducing the overall RMSD.

The RNA 2D structures predicted by IPknot, pKiss, DotKnot, RNAPKplex, and HotKnots (CC09 and DP09 parameters) are very similar to the native RNA 2D structure (**Supplementary Table S6 and Figure S5**). However, the RMSDs for these predictions range from 13.35 Å to 15.74 Å, whereas the native 2D structure-derived structure possesses an RMSD of 14.46 Å (**Supplementary Figure S65**). For IsRNA1 simulations, the highest RMSD (21.43 Å) was obtained with ContextFold RNA 2D structural constraints (**Supplementary Figure S65**). For ContextFold, the native pseudoknot base pairs were wrongly predicted as a stem part, whereas the G24-C37 stem–loop was correctly predicted (**Supplementary Table S6 and Figure S5**). Interestingly, the CentroidHomfold-LAST RNA 2D structure differs significantly from the native RNA 2D structure but still produces 3D structures with similar RMSDs of 14.61 Å and 14.46 Å, respectively. The INF_wc values for the modeled 3D structures with CentroidHomfold-LAST and native 2D structural constraints are 0.64 and 0.97, respectively (**Supplementary Figure S68**). This observation suggests that these relatively high-RMSD structures have different structural arrangements. Thus, the RMSD values from CentroidHomfold-LAST constraints and their comparison to that from native base pairing indicate that RMSD may not be a good measure for comparing RNA 3D structural quality.

#### PDB 5LYU

PDB 5LYU is a simple stem–loop structure, and many RNA 2D tools predicted the correct RNA 2D structure for this case. Thus, in this case, 2D to 3D translation could help us understand the reproducibility of prediction for simple stem–loop structures with all three 3D modeling approaches used here.

The RMSDs of the 3D structures corresponding to the native 2D structures from RNAfold_MFE, RNAfold_MEA, RNAstructure_Fold, RNAstructure_MaxExpect, MXfold2, LinearFold-V, RNAPKplex, R2DT, and pKiss were 10.22 Å, 10.25 Å, 10.22 Å, 9.00 Å, 11.66 Å, 8.56 Å, 10.22 Å, 10.22 Å, 10.25 Å, and 10.23 Å, respectively (**Supplementary Figure S17**). The lowest IsRNA1 RMSD (5.85 Å) was obtained with RNAfold_MFE_60 2D structural input, which was perfectly base-paired compared to the native 2D structure (**Supplementary Figure S17**). Thus, the 3D RMSD with the RNAfold_MFE_60 2D constraints could be considered a rare event of low-RMSD prediction, as with the same input, IsRNA1 mostly predicted structures with similar RMSDs (approximately 10.00 Å). The high INF_wc values for these 3D structures (0.93-1.00) indicated that stems might be formed properly (**Supplementary Figure S20**); however, the different arrangements of nucleotides in unpaired regions led to RMSDs from 8 Å to 10 Å.

Apart from the above RNA 2D structure prediction tools, DotKnot, NUPACK, CentroidFold_CONTRAfold, HotKnots_CC09, HotKnots_DP09, LinearFold-C, UNAFold_mFold, and IPknot produced the same 2D structure (**Supplementary Table S2 and Figure S1**). This 2D structure has significant stem and bulge structures (U10-G20, C38-G50 residues), leading to two large interior loops. The PPV, sensitivity, and F1-scores for these RNA 2D structures compared to the native 2D structure were 0.85. 0.77, 0.81, respectively. However, these 2D structural constraints in IsRNA1 led to consistent RMSD values of ~ 8.50 Å (except the RMSD of 7.05 Å for DotKnot).

#### PDB 5NWQ

PDB 5NWQ is a pseudoknot structure and one of the interesting cases for RNA 2D to 3D structure prediction. None of the 2D tools could predict the correct 2D structure for this RNA (**Supplementary Table S5 and Figure S4**). The 2D structural constraints provided by the native RNA 2D structure, RNAPKplex, pKiss, DotKnot, and HotKnots_DP09 provided structures with RMSDs of 5.19 Å – 12.31 Å (**Supplementary Figure S53**). All other methods produced structures with higher RMSDs and *p-value* higher than 0.01 (**Supplementary Figure S50**).

For this pseudoknot RNA, IsRNA1 with native base-paired 2D structural constraints exhibited an RMSD of 12.08 Å. Surprisingly, the pseudoknot structures predicted by DotKnot (5.19 Å), HotKnots_DP09 (5.65 Å), and pKiss (5.47 Å), which differed only by the C20-G29 canonical base pair compared to the native reference 2D structure, led to the lowest-RMSD structures. The active site region was correctly positioned by DotKnot, HotKnots_DP09, and pKiss compared to the native structure. The highest RMSDs were produced with the 2D structural constraints by CentroidFold_CONTRAfold (21.80 Å), LinearFold-C (21.73 Å), and R2DT (22.00 Å) (**Supplementary Figure S53**). The R2DT-predicted 2D structure has a large flexible loop region, A19-G34, whereas this region is divided into bulge and apical loop due to the base pairing of C21-G29 and C22-G28 in the CentroidFold_CONTRAfold and LinearFold-C predicted 2D structures (**Supplementary Table S5 and Figure S4**).

For the nonpseudoknot-specific RNA 2D structure prediction tools, the lowest RMSD of 16.59 Å was given by RNAfold_MFE_60, which matched CentroidFold_CONTRAfold, LinearFold-C, and R2DT. This might indicate a convergence issue with IsRNA1 simulation involving complex folds. SPOT-RNA, CentroidFold_McCaskill, RNAstructure_Probknot, IPknot, and R2DT predicted RNA 2D structures with a 5’-overhang strand, stem, and large loop (A19-G34) (**Supplementary Table S5 and Figure S4**). With these RNA 2D structural constraints, IsRNA1 provided RMSDs higher than 20.00 Å (**Supplementary Figure S53**). Thus, with IsRNA1, the predictions in the absence of pseudoknot 2D structural constraints led to large-RMSD structures incompatible with the expected RMSD distribution.

#### PDB 6P2H

The native RNA 2D structural constraints led to the lowest-RMSD (5.97 Å) structure. The RNA 2D structure prediction tools IPknot, RNAPKplex, RNAstructure Probknot, SS_from_PDB, ContextFold, DotKnot, RNAStructure MaxExpect, SpotRNA, and UNAFold_mFold led to structures with RMSD *p-value* lower than the significance value of 0.01 (**Supplementary Figure S74**). The DotKnot, RNAPKplex, and IPknot 2D structure-based ISRNA1 simulations produced RNA structures with RMSD values less than 10.00 Å, and SPOTRNA led to an RMSD of 11.27 Å (**Supplementary Figure S77**).

The PDB 6P2H sequence with IsRNA1 modeling demonstrates the importance of pseudoknot base-pair constraints in 3D structure prediction. The native 2D structure and CONTRAfold-predicted 2D structure are the same except for the absence of a pseudoknot base pair in the CONTRAfold prediction (**Supplementary Table S7 and Figure S6**). With the lack of information on pseudoknot base pairs but with canonical base pairs, the CONTRAfold-predicted 2D structural constraints led to a 3D structure with a large RMSD of 20.44 Å (**Supplementary Figure S77**). Similarly, if we compare the RMSDs of the 3D structures predicted with the 2D structural restraints from HotKnots_DP09 (27.96 Å, the largest RMSD in the dataset) and RNAPKplex (10.01 Å), which differ only by the pseudoknot base-pairing information, the results differ significantly, with RMSDs of 27.96 Å and 10.10 Å, respectively.

Although the UNAFold_mFold-derived 2D structure is an exact match with that of HotKnots_DP09, the RMSD of the IsRNA1-modeled 3D structure is 14.04 Å (*p-value* < 0.01) (**Supplementary Figure S74 and Figure S77**). The observed differences in the modeled 3D structure using the same 2D structural input can be attributed to the reproducibility issue with IsRNA1 when the additional 2D structural constraints of pseudoknots are not provided. The RMSDs of the 3D structures derived from HotKnots_DP09, RNAfold_MFE, RNAfold_MEA, HotKnots_CC09, UNAFold_mFold, RNAfold_MFE_60, MXfold2, CentroidFold_CONTRAfold, and ContextFold are all above 18.00 Å, which is surely above significance level of 0.01, thus indicating that the predictions did not come from the expected distribution.

#### PDB 6TB7

For this RNA sequence, pseudoknot-specific RNA 2D structure prediction tools, namely, RNAstructure_ProbKnot, DotKnot, RNAPKplex, HotKnots_CC09, HotKnots_DP09, pKiss, and IPknot, predicted incorrect pseudoknot base pairing, incorrect bulge, and apical loop prediction (**Supplementary Table S3 and Figure S2**). The native 2D structural input generated the lowest-RMSD (4.50 Å) 3D structure **(Supplementary Figure S29**). Most RNA 2D tools led to lower RMSDs for 3D structures, even if the predicted 2D structures differed significantly from the native 2D structure. This was counterintuitive with respect to the results displayed in the case of PDB 6P2H. Only DotKnot and CentroidFold_McCaskill led to RMSDs exceeding the significance threshold of 0.01 (**Supplementary Figure S26**). Even in the absence of pseudoknot base-pairing 2D structural constraints and inaccurate predictions on the apical loop and bulge, structures with accepted RMSD values were predicted (**Supplementary Figure S26**).

#### PDB 7LYJ

PDB 7LYJ is the SARS-COV-2 frameshifting pseudoknot and was an interesting test system for this study. The lowest-RMSD 3D structures were obtained with the native 2D structure, which was predicted by the SPOT-RNA 2D tool (**Supplementary Figure S41**). The 2D structures differ in the depiction of apical loop G40-C45. SPOT-RNA predicted the noncanonical base pair G41-A44, thus changing the G41-A42-A43-A44 tetraloop to the two-residue loop A42-A43 (**Supplementary Table S4 and Figure S3**). Thus, IsRNA1 seemed tolerant to small changes in the 2D structural constraints at the base-pairing level. The largest RMSD of 26.85 Å was obtained for the 2D structural constraints predicted by the MC-Fold 2D tool (**Supplementary Figure S41**).

Similar to PDB 6P2H, in the case of PDB 7LYJ, IsRNA1 also provided 3D structures with larger RMSDs for the 2D structural inputs without pseudoknot base pairing information compared to those with pseudoknot base pairing information. For example, IPknot predicted accurate 2D topology but missed the pseudoknot base pairing information, leading to a 3D structure with a large RMSD of 18.38 Å (**Supplementary Figure S41**). Notably, the 2D predictions of HotKnots_CC09, LinearFold-C, and ContextFold also matched the native 2D structure except for the prediction of the pseudoknot base pairing information, whereas when translated to 3D, they possessed RMSDs of 16.27 Å, 16.25 Å, and 10.93 Å, respectively. These observations indicated the possible convergence issues of IsRNA1.

#### RNAComposer

In general, RNAComposer provided the lowest RMSD 3D structures with RNA 2D structural inputs close to the native 2D structures. RNAComposer, a fragment building-based software, has no reproducibility issues. This means that with the same 2D structure, the same 3D structure can be obtained every time. A minor change in the 2D base pairing constraints may have a remarkable effect on the RNAComposer prediction of the 3D structure, as illustrated below.

#### PDB 3OWZ_A

For 3OWZ_A, among all three methods, RNAComposer created the lowest-RMSD structures with the native 2D structure and the 2D structures predicted by RNAfold_MEA, CONTRAfold, MXfold2, RNAstructure_ProbKnot, SpotRNA, and RNAfold_MEA (**Supplementary Table S8, Figure S7, and Figure S89**). As the 2D structures deviated increasingly near the three stem junctions (loop and bulge regions), the RMSDs of the modeled 3D structures increased (**Supplementary Table S8, Figure S7, and Figure S89**).

#### PDB 4ENC

In this case, the native RNA 2D structure translated to the lowest RMSD (0.76 Å) 3D structure (**Supplementary Figure S65**). Notably, PDB 4ENC is included in the underlying fragment database of FRABASE 2.0; thus, RNAComposer picked up the correct fragments, leading to an accurate 3D structure when sufficiently accurate 2D structural constraints were provided as input. In the native 2D structure, a pseudoknot is formed by the pairing of G2-G3-C4-G5 of the 5’-overhang segment with residues C14-G15-C16-C17 of the internal loop (**Supplementary Table S6 and Figure S5**). However, in IPknot, pseudoknot base pairing started from G1-G2-G3-G4. This shift by one base pair in pseudoknot led to a 3D structure with an RMSD of 2.86 Å (**Supplementary Figure S65**). This showed the sensitivity of 3D structure building by RNAComposer to the input 2D structural constraints. The largest RMSD was given by the 3D structure modeled with the CentroidHomfold-LAST 2D structural constraints, which are topologically distinct from the native 2D structure.

The IPknot and pKiss 2D structural notations are the same in the VARNA diagrams (**Supplementary Figure S5**). However, we observed an RMSD difference of 7.0 Å for the 3D structures predicted using the 2D structural constraints for these two 2D tools. Notably, the dot-bracket notations for the pseudoknot base pairing representation (either “[” and “]” or and “}” or “<” and “>”) as inputs to RNAComposer were different because we used the dot-bracket notations as they were output from the corresponding 2D tools. We found that RNAComposer is sensitive to 2D structural input representations and can predict different 3D structures if the representations of the same 2D structural input are changed. We assume that RNAComposer’s motif searching algorithm varies based on the 2D structural input representations.

#### PDB 5LYU

For this simple stem–loop structure, the 2D structure was correctly predicted by almost all of the conventional thermodynamics-based 2D tools and by a few pseudoknot-specific methods, for instance, RNAfold_MFE, RNAfold_MFE_60, RNAfold_MEA, RNAstructure_Fold, RNAstructure_MaxExpect, MXfold2, RNAPKplex, LinearFold-V, R2DT, and pKiss. (**Supplementary Table S2 and Figure S1**). These predicted 2D structural constraints and the native ones led to similar 3D structures with the same RMSD value of 6.4 Å (**Supplementary Figure S17**). The highest RMSDs were obtained for the 2D structural inputs predicted by CentroidHomfold-LAST and SPOT-RNA, 14.02 Å and 13.84 Å, respectively, with F1-scores of 0.581 and 0.808, respectively (**Supplementary Figure S17**). Thus, even though the F1-score was 0.808 for SPOT-RNA, the RNA 2D structural topology was very different from the native 2D structure, leading to high RMSD. This illustrated that a high F1-score does not always lead to low RMSD structures.

#### PDB 5NWQ

This structure is a pseudoknot, and the modeled 3D structures were very sensitive to secondary structural inputs. The 3D structures predicted with the native secondary structure, obtained from DotKnot, HotKnots_DP09, and pKiss, had RMSDs with a significance level below 0.01 (**Supplementary Figure S50**). The rest of the 2D tools led to higher RMSDs with large significance values (**Supplementary Figure S50**). Notably, the RNAPKplex-predicted 2D structural input-derived 3D structural RMSD (12.62 Å) was almost the same as that for pKiss (12.34 Å) (**Supplementary Figure S53**); however, the significance value was higher than that for pKiss, indicating that RMSDs produced by DotKnot, HotKnots_DP09, and pKiss were borderline higher. The predictions of DotKnot (11.94 Å), HotKnots_DP09 (11.94 Å), and pKiss (12.34 Å) differed from the native secondary structure by one base pair, C20-G29, and the pKiss secondary structure dot-bracket representation was different, demonstrating the effect of representation on the RNAComposer prediction again (**Supplementary Table S5, Figure S4, and Figure S53**).

#### PDB 6P2H

This structure provided an example of obtaining high RMSD 3D structures from RNAComposer even with correct secondary structural input (**Supplementary Table S7, Figure S6, and Figure S77**). Relatively low RMSDs, 11.54 Å, and 11.54 Å, respectively, were produced by DotKnot and RNAPKplex compared to the RMSD of 15.03 Å for the native secondary structural constraint-driven 3D structure (**Supplementary Figure S77**). The lack of pseudoknot base pairing information in the 2D structural input predicted by HotKnots_DP09 can explain the higher RMSD 3D structures for this 2D tool compared to that for the 2D tools DotKnot and RNAPKplex.

The 3D structures predicted with secondary structural constraints obtained from IPknot, RNAPKplex, RNAstructure_ProbKnot, native secondary structure, CentroidFold_McCaskill, DotKnot, RNAStructure MaxExpect, and SPOT-RNA possessed RMSDs between 10 Å and 20 Å, with the lowest being 11.54 Å (*p*-value < 0.01) (**Supplementary Figure S74**). The crystal structure of PDB 6P2H is a recently (2019) solved 2’-deoxyguanosine riboswitch with a three-way junction at the binding site of the ligand. We assume that as the underlying motif database of RNAComposer, RNA FRABASE 2.0, is not updated yet with the recently solved structural data, this may lead to high RMSD 3D structures even with accurate 2D structural input.

#### PDB 6TB7

RNAComposer provided low-RMSD 3D structures for PDB 6TB7, the lowest being 4.42 Å for the native 2D structural input (**Supplementary Figure S29**). PDB 6TB7 is the crystal structure of the NAD+ riboswitch with a flexible bulge and an internal loop (**Supplementary Table S3, Figure S2**). Although this is not a pseudoknot structure, it has signature canonical interactions between the bulge and internal loop residues, C14-G36 and C13-G38, respectively. UNAFold_MFold and UFold predicted 2D structures similar to the native 2D structure, albeit with the C14-G36 and C13-G38 base pairs missing, and these 2D structural inputs led to 3D structures with an RMSD value of 4.48 Å, which is similar to that obtained with the native 2D structural input. As this base pairing information contributes negligibly to the RMSD value, it can be assumed that these base pairs have low weight in the 3D structure prediction of PDB 6TB7. The 2D tools CONTRAfold, CentroidFold_CONTRAfold, and LinearFold-V displayed similar 2D structures to UFold, with a difference in a bulge near position A35, shifting the internal bulge at G23-G24-G25 to U22-G23-G24 (**Supplementary Table S3 and Figure S2**). However, this small change in the 2D structural rearrangement did not impact the 3D structural RMSD (4.84 Å) compared to the prediction by the 2D tool UFold and the native 2D structural input (Supplementary Information, Figure S22). DotKnot, MXfold2, and CentroidFold-McCaskill produced predictions that are topologically similar and very different from the native 2D structure and thus led to 3D structures with a high RMSD value of 17.28 Å and a *p-value* > 0.01 (**Supplementary Figure S26 and Figure S29**).

#### PDB 7LYJ

This SARS-COV-2 frameshift RNA with a pseudoknot displayed RMSDs between 10-15 Å for most of the 2D tool-derived structural constraints, with MC-Fold-, E2Efold-, and R2DT-derived 3D RMSDs of 25.96, 22.66, and 26.38, respectively, with the *p*-value leading tor ejection (**Supplementary Figure S38 and Figure S41**). The native 2D structure and the 2D structural constraints predicted by HotKnots_DP09 led to the lowest RMSD of 6.90 Å (**Supplementary Figure S41**). The DotKnot 2D structure has the 5’-C1 residue hanging and an additional G28-U58 base pair, and the G28-C1 base pair is missing compared to the native 2D structure; these differences led to a 3D structure with an RMSD of 7.82 Å (**Supplementary Table S4, Figure S3, and Figure S41**). The ContextFold- and MXfold2-derived 2D topologies are the same as the native topologies but lack pseudoknot base pairing information, which led to an increase of ~7 Å in the 3D RMSDs. Thus, RNAComposer is quite sensitive to 2D pseudoknot input information, as demonstrated by the PDBs 5NWQ, 6P2H, and 7LYJ for IsRNA1 and 6P2H for RNAComposer.

### SimRNA

#### PDB 3OWZ_A

For SimRNA, the lowest RMSD 3D structure of PDB 3OWZ_A was modeled by the 2D structural constraints predicted by RNAstructure_Fold, which differ from the native 2D structure in the region from G8 to A13 (**Supplementary Table S8, Figure S7, and Figure S89**). The native 2D structure has a four-nucleotide bulge (G8 to A11), whereas RNAstructure_Fold predicted a bulge comprising six residues from G8 to A13 and a missing G12-C28 base pair (**Supplementary Table S8 and Figure S7**). The 2D structures predicted by RNAfold_MEA, CONTRAfold, and MXfold2 match the native 2D structure (**Supplementary Table S8 and Figure S7**). However, SimRNA predictions with these three 2D structural inputs led to significantly different RMSDs ranging from 11.92 Å to 19.92 Å (**Supplementary Figure S89**). The largest RMSD obtained was 20.7 Å for the 2D structural constraints predicted by CentroidHomfold-LAST with a significantly lower F1-score of 0.654 than that for RNAfold_MEA (0.959). However, the 2D structure predicted by RNAfold_MEA translated to a 3D structure with an RMSD of 19.92 Å. Thus, it was quite surprising that with the more accurate native 2D structural constraints, the obtained RMSD was similar. This can be attributed to insufficient conformational sampling.

#### PDB 4ENC

With the native 2D structural constraints, SimRNA modeled the 3D structure with an RMSD of 19.65 Å for PDB 4ENC (**Supplementary Figure S65**). The calculated INF_wc for this 3D model was as low as 0.33, suggesting severe deviation from Watson-Crick base pairing compared to the crystal structure (**Supplementary Figure S68**). This observation indicated that conformations were insufficiently sampled during the simulations. The 2D structures predicted by IPknot, pKiss, DotKnot, RNAPKplex, HotKnots_CC09, and HotKnots_DP09 translated to 3D structures with consistent RMSDs of ~10.00 Å (**Supplementary Figure S65**). Notably, for CentroidHomfold-LAST and ContextFold, even though the topology of the predicted 2D structures is significantly different from the native 2D structure and pseudoknot base pairing information is missing in them, SimRNA modeled the 3D structures with an RMSD of ~10.0 Å (**Supplementary Table S6, Figure S5, and Figure S65**). CentroidFold_McCaskill predicted a large unpaired region and only partially predicted the stem–loop structure from G24 to C37 (**Supplementary Table S6 and Figure S5**). Intriguingly, in this case, even with minimal 2D structural constraints, SimRNA modeled the 3D structure with an RMSD value of ~11.0 Å (**Supplementary Figure S65**). SimRNA uses 2D structural information as distance restraints; however, the unpaired regions fold under the influence of the underlying SimRNA force field (which does not fully account for the molecular context). The CentroidHomfold-LAST-derived 3D structure demonstrated the efficiency of the SimRNA force field in folding RNAs.

#### PDB 5LYU

For PDB 5LYU, the native 2D structural constraints led to a 3D structure with an RMSD of 7.19 Å (**Supplementary Figure S17**). The 2D predictions by RNAfold_MFE, RNAfold_MFE_60, RNAfold_MEA, RNAstructure_Fold, RNAstructure_MaxExpect, LinearFold-V, RNAPKplex, R2DT, and pKiss were the same (**Supplementary Table S2 and Figure S1**). For these abovementioned 2D tools, SimRNA simulations led to similar RMSDs (6.47 Å – 7.66 Å) (**Supplementary Figure S17**). The MXfold2 2D structure matches the native 2D structure but still led to a 3D structure with an RMSD of 8.52 Å (**Supplementary Figure S17**). This observation indicated the reproducibility capability of SimRNA for simple stem–loop structures, but the same could not always be expected for complex fold structures such as PDBs 3OWZ and 4ENC. Notably, for PDB 5LYU, all three methods, IsRNA1, RNAComposer, and SimRNA, modeled 3D structures with RMSDs less than 10.0 Å.

#### PDB 5NWQ

The lowest-RMSD (7.17 Å) structure was produced with the 2D structural constraints obtained from the 2D tool HotKnots_DP09 (**Supplementary Figure S53**). The 2D structure predicted by HotKnots_DP09 deviates from the native 2D structure by the absence of a large loop comprising residues from C20 to G29 (**Supplementary Table S5 and Figure S4**). The 2D structure predicted by IPknot lacks pseudoknot base pairing information (**Supplementary Table S5 and Figure S4**); however, it produced a low-RMSD structure (8.65 Å) (**Supplementary Figure S53**). This observation demonstrates the capability of SimRNA to predict pseudoknot structures. However, obtaining diverse RMSD values for the 3D structures with the same 2D structural constraints predicted with IPknot, SPOT-RNA, CentroidFold_McCaskill, RNAstructure_ProbKnot, and R2DT pointed toward the insufficient conformational sampling issue in SimRNA simulations. The remaining 2D structures have distinct topologies compared to the native and HotKnots_DP09 predicted 2D structures and therefore led to RMSDs higher than 15.0 Å (**Supplementary Table S5, Figure S4, and Figure S53**).

#### PDB 6P2H

SimRNA simulations generally generated low-RMSD structures (~6-8 Å) compared to IsRNA1 and RNAComposer for PDB 6P2H (**Supplementary Figure S77**). Here, the lowest RMSD (5.79 Å) structure was produced by the 2D structural constraints predicted from IPknot. Interestingly, the 2D topology predicted by IPknot is distinctly different from that of the native 2D structure in terms of the expanded three-way junction (**Supplementary Table S7 and Figure S6**). The 2D structures predicted by DotKnot and the native one are different but still led to very similar 3D structures with RMSDs of 6.85 Å and 6.53 Å, respectively (**Supplementary Table S7, Figure S6, and Figure S77**). RNAstructure_Fold, RNAstructure_ProbKnot, RNAstructure_MaxExpect, and NUPACK predicted significantly different base pairs compared to the native 2D structure; this led to large RMSDs with lower s for these 2D tools.

#### PDB 6TB7

Overall, SimRNA produced higher RMSDs for this structure than the other methods (**Supplementary Figure S29**). However, with native and UNAFold_mFold-predicted 2D structural constraints, the 3D structures with the lowest RMSDs, 3.57 Å, and 3.28 Å, respectively, were generated by SimRNA compared to those generated by IsRNA1 and RNAComposer (**Supplementary Figure S29**). Although UFold and UNAFold_mFold predicted the same 2D structure, it translated to the 3D structure with an RMSD of 13.38 Å (**Supplementary Table S3, Figure S2, and Figure S29)**. The RNAfold, RNAStructure, and HotKnots variants, NUPACK, pKiss, and RNAPKplex, all predicted similar 2D structures, and those 2D structural constraints, except for those from RNAPKplex, generated 3D structures with a range of RMSDs ~13-14 Å relative to the native 3D structure. The 2D structural constraints predicted by RNAPKplex translated to a 3D structure with an RMSD of 9.53 Å. These observations indicate that although SimRNA simulations produce similar 3D structures with similar 2D structural constraints, the possibility of producing outliers cannot be ignored. As expected, the distinct topology predicted by MXfold2, DotKnot, and CentroidFold_McCaskill led to higher RMSDs (*p-value*>0.01) (**Supplementary Figure S26 and Figure S29**).

#### PDB 7LYJ

In general, for most of the 2D structural constraints, except for those from UFold and RNAfold_MC-Fold, SimRNA-generated 3D structures were close to the crystal structure (**Figure S41)**. Even if pseudoknot base pairing information was not predicted, SimRNA efficiently folded to 3D structures with low RMSD values for the 2D structural constraints predicted by IPknot, ContextFold, and many other 2D tools (**Supplementary Table S4, Figure S3, and Figure S41)**. These observations illustrated the remarkable ability of SimRNA simulations to predict nearly accurate 3D structures in the absence of pseudoknot base pairing information as 2D structural constraints. These results are consistent with the observations in Reference 49.

#### SimRNA simulation convergence

We found that with the same RNA 2D structure input and the default single REMC simulation run, SimRNA-predicted 3D models displayed a wide range of RMSD values with respect to the reference crystal structure. Thus, assuming convergence as one of the reasons, we followed the protocol of repeated SimRNA REMC simulations as suggested by the SimRNA developers to mitigate the convergence issues.

We found that at least for 4 RNAs, namely, 3OWZ_A, 4ENC, 6TB7, and 5NWQ, the accuracy of 3D structure prediction may suffer from the issue of insufficient conformational sampling in SimRNA simulations with the default setup. For instance, with the native 2D structural constraints, the RMSD of the SimRNA-derived 3D structure was as high as 19.70 Å for PDB 4ENC. Similarly, 2D structural constraints with a high F1-score (0.96) for PDB 3OWZ_A translated to a high-RMSD (19.65 Å) 3D structure.

To check whether these observations arose due to insufficient conformational sampling, we performed SimRNA replica exchange Monte Carlo (REMC) simulations repeatedly as described in the methods section. In short, the same REMC simulation was performed in three sets (set 1, set 2, and set 3) and then analyzed individually, and RMSDs were compared with each other and to those obtained from the default simulations to ensure whether the results were reproducible. Notably, for set 1, set 2, and set 3, only the native 2D structural constraints were used. The native 2D structural constraints with repeated simulations could help shed light on the variations in RMSD values in terms of RNA topology complexity.

The RMSD results are summarized in **Figure 9** (**Supplementary Table S1**). Except for PDB 3OWZ_A and 6P2H, the predicted RMSDs did not vary much between sets and default simulations. Set 1, Set 2, and Set 3 demonstrated varying RMSDs for 3OWZ_A and 6P2H, thus indicating the possibility of different outcomes even if native 2D structural information is provided. We attribute this variation to the complex topologies of these RNAs.

**Figure 9.**
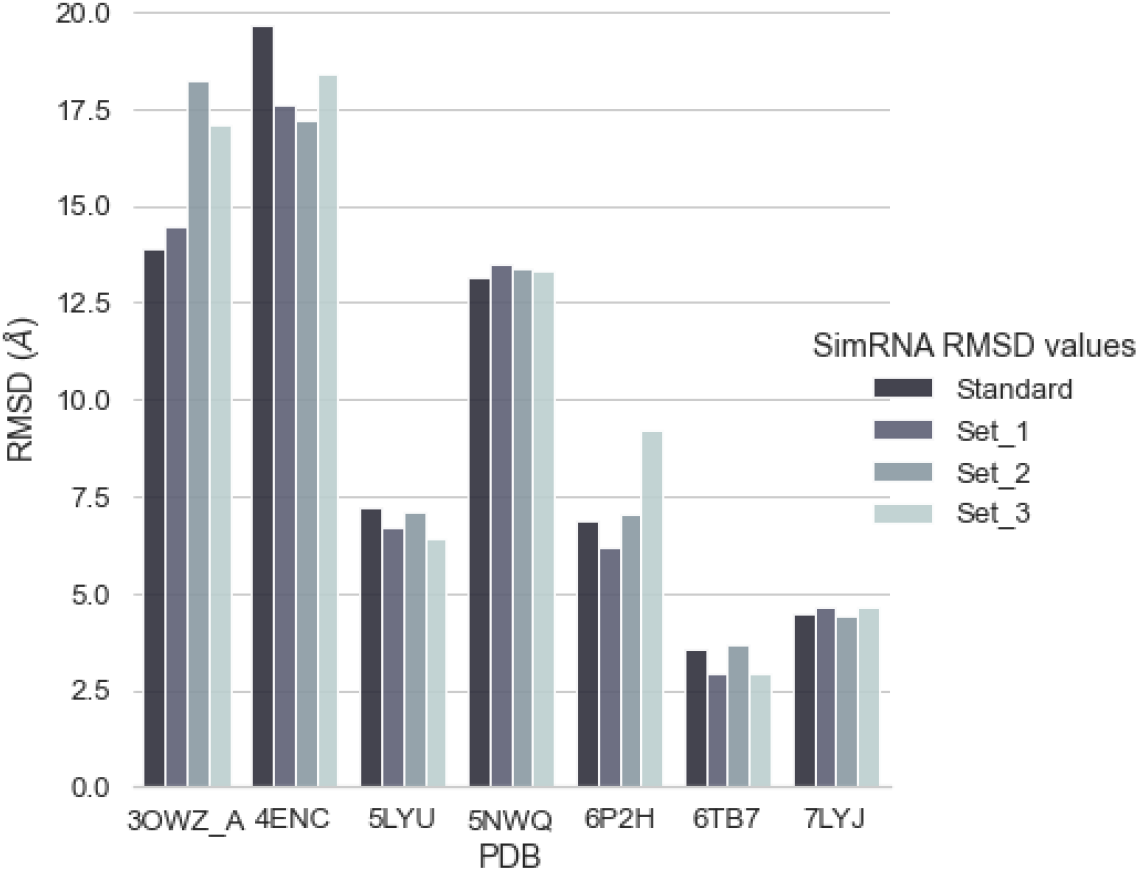
RMSD values of repeated SimRNA simulations with native 2D structural constraints.

## CONCLUSIONS

In this work, we assessed the transferability of 2D structural constraints to 3D models for seven RNAs with different complex topologies. The 2D structural constraints predicted by 27 different tools were utilized as inputs in widely known software packages for RNA 3D structure prediction, IsRNA1, RNAComposer, and SimRNA. Extensive comparative analyses were performed to correlate the accuracy of the 2D structure predictions and the 3D model quality.

We found that for the predicted 2D structures with high F1-scores, the motif-based tool RNAComposer efficiently modeled the 3D structures with a median RMSD of ~5.00 Å. Accordingly, the 3D tools can be ordered as RNAComposer > SimRNA > IsRNA1 based on their efficiency of translating 2D structural constraints into 3D models. We observed that the accuracy of the 2D structure prediction, the complexity of the structural folds, and the availability of 3D structural motifs in the RNA FRABASE 2.0 database played a significant role in the 2D to 3D translation accuracy of RNAComposer. For the group of 2D tools with medium F1-scores, SimRNA modeled the 3D structures more efficiently than RNAComposer and IsRNA1. Our results suggested that the sampling efficiency and the underlying force field of SimRNA played a major role in compensating for the lack of accuracy in the 2D structural inputs. Irrespective of motif-based or sampling-based approaches, the modeled 3D structures deviated greatly from the crystal structures for the 2D structures predicted with F1-score =< 0.80.

However, further extensions of the current study could confirm the performance and accuracy of structural prediction and establish the correlation between 2D and 3D modeling of RNA. This will include the use of state-of-the-art methods in both 2D (recent ML/DL methods such as CNNFold, CROSS, DMfold, and RPRes) and 3D (Rosetta FARFAR2 and 3dRNA). Further improvements can be expected by considering a larger dataset, including alternative conformations for each RNA sequence, and incorporating auxiliary information such as chemically modified free energy nearest neighbor parameters as folding constraints.

## SUPPLEMENTARY DATA

Supplementary information is available Online.

