## Supplementary Data for "Comparative Analysis of RNA Secondary Structure Accuracy on Predicted RNA 3D Models"

### Commands used for secondary structure methods:

```
## RNAfold MFE
RNAfold -p -d2 --noLP < seq.fa > rnafold.out

RNAfold -p -d2 --noLP -T 60 < seq.fa > rnafold_60C.out ## RNAfold MFE
structure at 60


## RNA Structure Fold sequence
Fold seq.fa Fold.ct --loop 30 --maximum 20 --percent 10 --temperature
310.15 --window 3

## create partition function
partition seq.fa partition.pfs --temperature 310.15

## RNAStructure Maximum expected secondary structure
MaxExpect partition.pfs MaxExpect.ct --gamma 1 --percent 10 --structures
20 --window 3

### Pseudoknot probable structure
ProbKnot partition.pfs ProbKnot.ct --iterations 1 --minimum 3

## converting .ct files to dot files
ct2dot Fold.ct 1 Fold.dot -f side
ct2dot ProbKnot.ct 1 ProbKnot.dot  -f side
ct2dot MaxExpect.ct 1 MaxExpect.dot -f si

## Dotknot
mkdir Dotknot

## needed to activate conda in Bash
eval "$(conda shell.bash hook)"

##
conda activate py2

cd /home/mandar/softwares/dotknot/DotKnot_1.3.2/src

## -k == kissing loop
## -l = loop
## -g == global structure

## writting FASTA file
cat << EOF > seq.fa
>seq
$sequence
EOF

python dotknot.py seq.fa -k -l -g > seq.out
```

**#Contrafold**

```
mkdir contrafold ; cd contrafold

cp ../seq.fa .

/home/mandar/softwares/CONTRAfold/contrafold/src/contrafold predict seq.fa
> contrafold.out
```

**# MXFOLD2**

```
mkdir mxfold2 ; cd mxfold2

eval "$(conda shell.bash hook)"

conda activate mxfold2_local

cp ../seq.fa .

mxfold2 predict seq.fa  > mxfold2.out
```

**#NUPACK**

```
mkdir NUPACK; cd NUPACK

## writing FASTA file
cat << EOF > seq.in
$sequence
EOF

/home/mandar/softwares/NUPACK/nupack3.0.6/bin/mfe -batch seq -material
rna1995 -dangles 1 -T 37.0 -sodium 1.0 -magnesium 0.0 nupac_out

### output is seq.mfe
```

#### Centroidfold

```
#!/bin/bash
### Note: -g is gamma factor, gamma tunes sensitivity and PPV
### The original publication " Prediction of RNA secondary structure using
generalized centroid estimators" suggests
### if RNA length < 80 --> highest MCC value is given by
###   gamma = 4 for CONTRAfold model
###   gamma = 1 for McCaskill model

echo ""
echo ""
echo "Centroidfold McCaskill model"
~/softwares/centroidfold_0_0_16/centroid-rna-package-0.0.16-linux-
x86_64/centroid_fold -g 1 -e "McCaskill" seq.fa > seq_c_m.out
cat seq_c_m.out

echo ""
echo ""

echo "Centroidfold CONTRAFOLD model"
~/softwares/centroidfold_0_0_16/centroid-rna-package-0.0.16-linux-
x86_64/centroid_fold -g 4 -e "CONTRAfold" seq.fa > seq_c_contra.out
cat seq_c_contra.out
echo ""
echo ""
```

#### Linearfold-V and Linearfold-C

```
#!/bin/bash

## Linearfold-V
echo ""
echo ""
echo "Linearfold-c results!!"
tail -n1 seq.fa | ~/softwares/linearfold/LinearFold-
master/bin/linearfold_c | tail -n1

## Linearfold-C
echo ""
echo ""
echo "Linearfold-v results!!"
tail -n1 seq.fa | ~/softwares/linearfold/LinearFold-
master/bin/linearfold_v | tail -n1
```

#### RNAPKplex

```
#!/bin/bash

RNAPKplex  --noLP < seq.fa > seq.out
```

#### Contrafold

```
~/softwares/CONTRAfold/contrafold/src/contrafold predict seq.fa
```

**#Hotknot**

```
cp -r ~/softwares/hotknots/HotKnots_v2.0/bin/ .
cd bin/

echo ""
echo "CC09 parameters !!!"

./HotKnots -s `tail -n1 ../seq.fa` -m CC -p params/parameters_CC09.txt  |
grep "S0:"

echo ""
echo "DP09 parameters !!!"

./HotKnots -s `tail -n1 ../seq.fa` -m DP -p params/parameters_DP09.txt |
grep "S0:"

echo ""
echo ""
```

**### ipknot**

rna_benchmark_seq.fa has all the sequences.

```
./ipknot rna_benchmark_seq.fa
```

**Webservers:**

Ufold
Mcfold
Mcfold + RNAfold – RNAFold MFE as input to Mcfold to constrain folding
pkiss
R2DT
CentroidHomfold

### RNA 3D Structure Prediction Tools and Input file Sample scripts

The steps involved in this automated protocol are described below. An initial RNA sequence and corresponding 2D structure were provided to predict the 3D structures. The IsRNA1 scripts predicted three models were predicted using Vfold3D/VfoldLA approach. Then, for each model, replica exchange simulations (10 replicas for a temperature window from 200 K to 425 K) were performed for $5 \times 10^7$ steps (50 ns) and 2000 structures per replica were recorded. The last 1000 coarse-grained structures from each of the three replica exchange simulations were combined for clustering. The clustering was performed on a total of 3000 coarse-grained structures with an RMSD cutoff of 10% of the sequence length and centroid structures corresponding to each cluster were extracted. The predicted structures were back-mapped to all-atom models. These cluster's centroid structures were sorted with the filtering protocol of IsRNA1 using standard molecular dynamics (MD) simulations. Five repeats of 0.5 ns of MD simulations were executed for each modeled structure to generate 1000 structures. These structures were then clustered with a 5 Å RMSD cutoff. The average IsRNA1 model energy of this structural ensemble was used as one of the criteria to rank structures along with RMSD relative to modeled cluster's centroid structures. As suggested in Ref. (1), this protocol promotes the selection of structures with stable, low RMSD (possibly close to native) and with high INF_wc values as top-ranked structures.

#### IsRNA1:

```
work_dir=${PWD}

#work_dir=/nfs/mkulkarni/IsRNA1/PDB_7LYJ

jobname="PDB_7LYJ"

## writing seq2D.txt

cat <<EOF > seq2D.txt

CGGUGUAAGUGCAGCCCGUCUUACACCGUGCGGCACAGCGGAAACGCUGAUGUCGUAUACAGGGCU

(((((((((...[[[[[..)))))))))((((((((((((....))))).)))))))....]]]]]

EOF
```

```
### modifying conf-TF.dat if needed

cat << EOF > conf-TF.dat
Nstep 50000000     # Number of simulation steps.
Nstep_Ter 30000000 #Not used as no tertiary structure folding
Nstru 2000         #Number of recorded structures from each replica
PS 0.5             #Last 50% recorded structures for conformational
ensemble
PE 0.1             #Top 10% lowest-energy structures are clustered to
predict the 3D structures
RMSD_Cut -1        #RMSD cutoff 10% of sequence length
Nout 20            #Number of predicted 3D structures
EOF


###excluded pdb-list
cat << EOF > excluded_pdblist.txt
PDB ID of case study
EOF


######################### CHANGES END
!!!!!##########################################################

seq2D_file=seq2D.txt
excluded_pdblist_file="excluded_pdblist.txt"
config_file="conf-TF.dat"


/home/sysguru/IsRNA_Vfold3DLA_standalone/bin/IsRNA-Tertiary_Folding.out
$Input_Data $seq2D_file $work_dir $jobname $config_file 0
$excluded_pdblist_file 0
```

#### Clustering in IsRNA1:

```
## absolute paths are given here
### Do not change Input_Data
Input_Data=/home/sysguru/IsRNA_Vfold3DLA_standalone/Data/

########################## CHANGE HERE !!!!!!!!!!!!!!!!!!!!!!
###################################

work_dir=${PWD}
#work_dir=/nfs/mkulkarni/IsRNA1/PDB_7LYJ
jobname="PDB_3OWZ_A"

## writing seq2D.txt
cat <<EOF > seq.txt
GGCUCUGGAGAGAACCGUUUAAUCGGUCGCCGAAGGAGCAAGCUCUGCGGAAACGCAGAGUGAAACUCUCAGGC
AAAAGGACAGAGUC
EOF
```

```
### modifying conf-TF.dat if needed

cat << EOF > conf-TF.dat
Nstep 500000     #Number of steps for MD simulations, 1 ns = 1000000 steps
Nstru 1000       #Number of recorded structures from each trajectory
Nduplicate 5     #Number of duplicate simulations for each model
RMSD_Cut 5.0     #RMSD cutoff to collect close decoy structures
Energy_Cut 0.975 #Energy threshold to filter good models
EOF


######################### CHANGES END
!!!!!##############################################################

seq_file=seq.txt
config_file="conf-TF.dat"

mkdir modeled_pdbs
mv "$jobname"_IsRNA-*.pdb modeled_pdbs/


numfile=`ls modeled_pdbs | wc -l`

rm -rf pdblist.txt
for i in $( seq 1 $numfile ); do echo "modeled_pdbs/"$jobname"_IsRNA-
$i.pdb" >> pdblist.txt ; done

pdblist="pdblist.txt"
energy_file="energy.txt"

/home/sysguru/IsRNA_Vfold3DLA_standalone/bin/IsRNA-Filtration_Protocol.out
$Input_Data $seq_file $work_dir $pdblist $energy_file $config_file 0
```

#### SimRNA

The sampled conformations are generally clustered further to obtain an ensemble of RNA or minimum energy structures. For all PDBs listed in Table 1, the default SimRNA simulation protocol is followed. 160000000 steps of REMC simulations were performed with 10 replicas distributed over a SimRNA specific temperature range of 0.90 to 1.35. Please note that these temperature values are specific to SimRNA model energies, and it lacks relation with actual physical temperature values. A total of 1600 structures were collected for each replica, and all replicas were catenated. Clustering was performed with a 5 Å cutoff. The centroid structure of the first cluster is reported here. Although the thumb rule of 10% of RNA length was suggested by both IsRNA1 and SimRNA developers, we followed 5 Å cutoff to have a consistent workflow.

```
set name=4ENC
### Give sequence here
cat << EOF > sequence.dat
GGGCGAUGAGGCCCGCCCAAACUGCCCUGAAAAGGGCUGAUGGCCUCUACUG
EOF
################ Write config file ###########
cat  <<EOF > config.dat
## These are default parameters
NUMBER_OF_ITERATIONS 16000000
TRA_WRITE_IN_EVERY_N_ITERATIONS 16000
INIT_TEMP 1.35
FINAL_TEMP 0.90
SECOND_STRC_RESTRAINTS_WEIGHT 1.0
BONDS_WEIGHT 1.0
ANGLES_WEIGHT 1.0
TORS_ANGLES_WEIGHT 0.0
ETA_THETA_WEIGHT 0.40
EOF

## if RNA SS does not have pseudoknot, put RNA SS on first line only
cat <<EOF  > rna_ss.dat
(((((........))))).....(((((....)))))...............
.......((((((............................)))))).....
EOF
################################# MAKE CHANGES ENDS
####################################################

## linking data folder in WORKING DIR
ln -s /md_data/rosetta/SimRNA_64bitIntel_Linux/data/ data

# Execute SimRNA
${SimRNA} -s sequence.dat -c config.dat -S rna_ss.dat -E 10 -o
OUT_seq_${name}_simRNA >& mytestE.log
```

#### Clustering in SimRNA:

```
SIMRNA_PATH=/data/mkulkarni/softwares/SimRNA/SimRNA_64bitIntel_Linux
### cleaning up exisitng data
rm -rf *.trafl *.pdb *.ss_detected clustered_trafl_files
### catenating all trajectory files
cat ../OUT_seq_4ENC_simRNA_??.trafl > catenated_ALL.trafl
## Syntax: clustering "catenated_trafl_traj" "energy_cutoff_percentage
(0.01 means 1%)" "RMSD cutoff (angstrom) thumb rule: 10% of sequence
length"
${SIMRNA_PATH}/clustering catenated_ALL.trafl 0.01 5.0 >& mytest_clust.log
### printing number of structures per cluster
```

```
grep  "contains " mytest_clust.log | grep -w  "cluster" >
num_str_per_cluster.dat
## checking "data" folder in working directory
 if [ ! -d "data" ]
then
    echo "Directory "data" DOES NOT exist !! COPYING!!\n"
    ln -s ${SIMRNA_PATH}/data/ data
else
        echo "Directory "data" exists !! \n"
 fi
 ## copy PDB file from previous folder
cp ../OUT_seq_4ENC_simRNA_01-000001.pdb .
 ### Convert coarse-grained models to all-atom
${SIMRNA_PATH}/SimRNA_trafl2pdbs OUT_seq_4ENC_simRNA_01-000001.pdb
catenated_ALL_thrs*_clust01.trafl 1  AA

 ### move all cluster trafl to one folder
mkdir clustered_trafl_files
mv catenated_ALL_thrs*.00A_clust??.trafl clustered_trafl_files
```

#### RNAComposer

RNAComposer divides the user-defined RNA 2D structure into structural elements, namely stem, loop, single strands, and n-way junctions capped with canonical base pairs. These 2D structure elements are screened against a prebuilt FRABASE dictionary to get 3D structure elements. For a missing 3D structural element in FRABASE corresponding to a certain sequence, alternate 3D structures in single strands and stems are modeled using the NAB utility of AmberTools while the loops are built in the torsion angle space using the CYANA structure calculation program. These loop structures are initiated using A-RNA torsion angles extracted from NDB and distance restraints were applied for the closure of the sugar rings and loop closing residues.

### Table S1. RMSD values of 4ENC Standard and Repeated SimRNA simulations

| PDB | Standard | Mean value from three sets [a] |
|---|---|---|
| 5LYU | 7.19 | 6.74 (0.36) |
| 6TB7 | 3.57 | 3.17 (0.41) |
| 7LYJ | 4.48 | 4.56 (0.14) |
| 5NWQ | 13.13 | 13.40 (0.06) |
| 4ENC | 19.65 | 17.73 (0.61) |
| 6P2H | 6.85 | 7.47 (1.56) |
| 3OWZ_A | 13.91 | 16.59 (1.95) |

[a] Standard deviation value in parentheses.

#### Table S2. Predicted 2D structures of PDB 5LYU sequence in dot-bracket notation.

| | |
|---|---|
| **PDB 5LYU** | **GGGGAUCUGUCACCCCAUUGAUCGCCUUCGGGCUGAUCUGGCUGGCUAGGCGGGUCCC** |
| **RNApdbee_3DNA-DSSR** | **.((((((((((.(((((..(((((((....))).)))))))..))...))))))))))** |
| RNAfold_MFE | .((((((((((.(((((..(((((((....))).)))))))..))...)))))))))) |
| RNAfold_MFE_60 | .((((((((((.(((((..(((((((....))).)))))))..))...)))))))))) |
| RNAfold_MEA | .((((((((((.(((((..(((((((....))).)))))))..))...)))))))))) |
| RNAstructure_Fold | .((((((((((.(((((..(((((((....))).)))))))..))...)))))))))) |
| RNAstructure_ProbKnot | .((((((((((...(((..(((((((....))).))))))).......)))))))))) |
| RNAstructure_MaxExpect | .((((((((((.(((((..(((((((....))).)))))))..))...)))))))))) |
| DotKnot | .((((((((((...(((..(((((((....))).))))....)))...)))))))))) |
| CONTRAfold | .((((((((((...((((.(((((((....))).)))).)..)))...)))))))))) |
| MXfold2 | .((((((((((.(((((..(((((((....))).)))))))..))...)))))))))) |
| SPOT-RNA | <((((((((((....>(..((((((((..)))).))))....).....)))))))))) |
| NUPACK | .((((((((((...(((..(((((((....))).))))....)))...)))))))))) |
| UFold | .((((((((((((((((..((((((((..)))).)))))))..)).).)))))))))) |
| RNAPKplex | .((((((((((.(((((..(((((((....))).)))))))..))...)))))))))) |
| CentroidFold_McCaskill | .((((((((((........(((((((....))).))))..........)))))))))) |
| CentroidFold_CONTRAfold | .((((((((((...(((..(((((((....))).))))....)))...)))))))))) |
| HotKnots_CC09 | .((((((((((...(((..(((((((....))).))))....)))...)))))))))) |
| HotKnots_DP09 | .((((((((((...(((..(((((((....))).))))....)))...)))))))))) |
| LinearFold-C | .((((((((((...(((..(((((((....))).))))....)))...)))))))))) |
| LinearFold-V | .((((((((((.(((((..(((((((....))).)))))))..))...)))))))))) |
| UNAFold_mFold | .((((((((((...(((..(((((((....))).))))....)))...)))))))))) |
| MC-Fold | .((((((((((((((((((((((((((...)))))))))))..))))))))))))))) |
| RNAfold_MC-Fold | .((((((((((((((((..((((((((..)))).)))))))..))..))))))))))) |

```
E2Efold                ....((....((.....).(...).............).......(......)))...
pKiss                  .((((((((((.(((((..(((((((....))).)))))))..))...))))))))))
ContextFold            ((((.((((((...(((..(((.(((....)))..)))....)))...))))))))))
IPknot                 .((((((((((...(((..(((((((....))).))))....)))...))))))))))
R2DT                   .((((((((((.(((((..(((((((....))).)))))))..))...))))))))))
CentroidHomfold-LAST   ......(((((........((((...........))))..........))))).....
```

**Table S3. Predicted 2D structures of PDB 6TB7 sequence in dot-bracket notation.**

| | |
|---|---|
| **PDB 6TB7** | **GGCUUCAACAACCCCGUAGGUUGGGCCGAAAGGCAGCGAAUCUACUGGAGCC** |
| **RNApdbee_3DNA-DSSR** | **(((((((.....[[.((((((((.(((....))).])]))))))))))))))** |
| RNAfold_MFE | (((((((........(((((((..(((....)))....)))))))))))))) |
| RNAfold_MFE_60 | (((((((........(((((((..(((....)))....)))))))))))))) |
| RNAfold_MEA | (((((((........(((((((..(((....)))....)))))))))))))) |
| RNAstructure_Fold | (((((((........(((((((..(((....)))....)))))))))))))) |
| RNAstructure_ProbKnot | (((((((........(((((((..(((....)))....)))))))))))))) |
| RNAstructure_MaxExpect | (((((((........(((((((..(((....)))....)))))))))))))) |
| DotKnot | (((((((.(((((.....))))).(((....)))...........))))))) |
| CONTRAfold | (((((((........(((((((.((((....)))..).)))))))))))))) |
| MXfold2 | (((((((.(((((.....))))).(((....)))...........))))))) |
| SPOT-RNA | (((((((.<......(((((((>.((((..))))....)))))))))))))) |
| NUPACK | (((((((........(((((((..(((....)))....)))))))))))))) |
| UFold | (((((((........((((((((.(((....)))..).)))))))))))))) |
| RNAPKplex | (((((((........(((((((..(((....)))....)))))))))))))) |
| CentroidFold_McCaskill | (((((((.(((((.....))))).(((....)))...........))))))) |
| CentroidFold_CONTRAfold | (((((((........(((((((.((((....)))..).)))))))))))))) |
| HotKnots_CC09 | (((((((........(((((((..(((....)))....)))))))))))))) |
| HotKnots_DP09 | (((((((........(((((((..(((....)))....)))))))))))))) |
| LinearFold-C | (((((((........(((((((..(((....)))....)))))))))))))) |
| LinearFold-V | (((((((........(((((((.((((....)))..).)))))))))))))) |
| UNAFold_mFold | (((((((........((((((((.(((....)))..).)))))))))))))) |

| | |
|---|---|
| MC-Fold | (((((((........(((((((((((((..))))..)))))))))))))))) |
| RNAfold_MC-Fold | (((((((........(((((((((((((..))))..)))))))))))))))) |
| E2Efold | ..(..(.....(.................(.)...)......)...)..... |
| pKiss | (((((((........(((((((..(((....)))....)))))))))))))) |
| ContextFold | (((((..........(((((((.((((....)))..).)))))))..))))) |
| IPknot | (((((((.[[.....((((((]].(((....))).....))))))))))))) |
| R2DT | (((((((........((((((((((((....)))))..)))))))))))))) |
| CentroidHomfold-LAST | No output |

#### Table S4. Predicted 2D structures of PDB 7LYJ sequence in dot-bracket notation.

| | |
|---|---|
| **PDB 7LYJ** | **CGGUGUAAGUGCAGCCCGUCUUACACCGUGCGGCACAGCGGAAACGCUGAUGUCGUAUACAGGGCU** |
| **RNApdbee_3DNA-DSSR** | **(((((((((...[[[[[..)))))))))((((((((((((....))))).)))))))....]]]]]** |
| RNAfold_MFE | .((((((((.((.....))))))))))(((((((((((((....))))).))))))))........ |
| RNAfold_MFE_60 | (((((((((.((.....)))))))))))((((((((((((....))))).))))))))......... |
| RNAfold_MEA | (((((((((.((.....)))))))))))((((((((((((....))))).))))))))......... |
| RNAstructure_Fold | (((((((((.((.....))))))))))).(((((((((((....))))).))))))).......... |
| RNAstructure_ProbKnot | (((((((((.((<<<<.)))))))))))((((((((((((....))))).))))))).....>>>> |
| RNAstructure_MaxExpect | (((((((((.((.....)))))))))))((((((((((((....))))).))))))))......... |
| DotKnot | .((((((((...[[[[[..))))))))(((((((((((((....))))).))))))))...]]]]] |
| CONTRAfold | (((((((((.((.....)))))))))))((((((((((((....))))).))))))))......... |
| MXfold2 | (((((((((..........)))))))))((((((((((((....))))).))))))))......... |
| SPOT-RNA | (((((((((...<<<<<..))))))))).(((((.((((((..))))))..))))).....>>>>> |
| NUPACK | (((((((((.((.....)))))))))))((((((((((((....))))).))))))))......... |
| UFold | (((((((((.((((((()))))))))))((((((((((((....))))).)))))))....))))) |
| RNAPKplex | .((((((((.(([[[[[))))))))))(((((((((((((....))))).))))))))...]]]]] |

```
Centroid-
Fold_McCaskill        .((((((((..........)))))))).((((((((((((....))))).)))))))).........
CentroidFold_CONTRA-
fold                  (((((((((..(.....).)))))))))((((((((((((....))))).))))))).........
HotKnots_CC09         (((((((((..........))))))))).(((((((((((....))))).))))))..........
HotKnots_DP09         (((((((((...[[[[[..)))))))))((((((((((((....))))).)))))))....]]]]]
LinearFold-C          (((((((((..........))))))))).(((((((((((....))))).))))))..........
LinearFold-V          .((((((((.((.....))))))))))(((((((((((((....))))).))))))))........
UNAFold_mFold         (((((((((.((.....)))))))))))((((((((((((....))))).))))))).........
MC-Fold               (((((((((.((((((.(((((((((((.......).)))))))))))))))))))))))))....
RNAfold_MC-Fold       .((((((((.((.(..)))))))))))((((((((((((((..)))))).))))))))........
E2Efold               (<.(.(....<......(...(.>..<(..........)....)).(.(.{>.)).}...).))>.
pkiss webserver       [[[[[[[[[...{{{{{..]]]]]]]]].(((((((((((....))))).)))))).....}}}}}
ContextFold           (((((((((..........)))))))))((((((((((((....))))).))))))).........
IPknot                (((((((((..........)))))))))((((((((((((....))))).))))))).........
R2DT                  ((((((((..((.....)).))))))))..((.(((((((....))))).)).))...........
CentroidHomfold-LAST  (((((((((..........)))))))))((((((((((((....))))).))))))).........
```

**Table S5. Predicted 2D structures of PDB 5NWQ sequence in dot-bracket notation.**

| | |
|---|---|
| **PDB 5NWQ** | **CCGGACGAGGUGCGCCGUACCCGGUCAGGACAAGACGGCGC** |
| **RNApdbee_3DNA-DSSR** | **[[[[.......(((((((.(]]]]....).....)))))))** |
| RNAfold_MFE | ((......)).(((((((..((.....)).....))))))) |
| RNAfold_MFE_60 | ...........(((((((..((.....)).....))))))) |
| RNAfold_MEA | ((......)).(((((((..((.....)).....))))))) |
| RNAstructure_Fold | ((......)).(((((((..((.....)).....))))))) |
| RNAstructure_ProbKnot | ...........(((((((................))))))) |
| RNAstructure_MaxExpect | ((......)).(((((((..((.....)).....))))))) |
| DotKnot | ((((.......[[[[[[[..))))..........]]]]]]] |
| CONTRAfold | .(......)..(((((((..((.....)).....))))))) |
| MXfold2 | ...........(((((((..((.....)).....))))))) |
| SPOT-RNA | ...........(((((((................))))))) |
| NUPACK | ((......)).(((((((..((.....)).....))))))) |
| UFold | No output |
| RNAPKplex | ((.(((..((([[[[[[[)))..))).)).....]]]]]]] |
| CentroidFold_McCaskill | ...........(((((((................))))))) |
| CentroidFold_CONTRAfold | ...........(((((((..((.....)).....))))))) |
| HotKnots_CC09 | ((......)).(((((((..((.....)).....))))))) |
| HotKnots_DP09 | ((((.......[[[[[[[..))))..........]]]]]]] |
| LinearFold-C | ...........(((((((..((.....)).....))))))) |

| | |
|---|---|
| LinearFold-V | ((......)).(((((((..((.....)).....))))))) |
| UNAFold_mFold | ((......)).(((((((..((.....)).....))))))) |
| MC-Fold | ..((((((((((.(((((((...)))))))))))))))).) |
| RNAfold_MC-Fold | No output |
| E2Efold | ((.(.((.....).........)............).).). |
| pKiss | [[[[.......{{{{{{{..]]]]..........}}}}}}} |
| ContextFold | ...........((((((...((.....))......)))))) |
| IPknot | ...........(((((((................))))))) |
| R2DT | ...........(((((((................))))))) |
| CentroidHomfold-LAST | ...........((((((....))(((.......))).)))) |

**Table S6. Predicted 2D structures of PDB 4ENC sequence in dot-bracket notation.**

| | |
|---|---|
| **PDB 4ENC** | **GGGCGAUGAGGCCCGCCCAAACUGCCCUGAAAAGGGCUGAUGGCCUCUACUG** |
| **RNApdbee_3DNA-DSSR** | **.[[[[..((((((]]]]......(((((....)))))....)))))).....** |
| RNAfold_MFE | ((((......))))(((......(((((....)))))....)))........ |
| RNAfold_MFE_60 | ((((......))))(((......(((((....)))))....)))........ |
| RNAfold_MEA | ((((......))))(((......(((((....)))))....)))........ |
| RNAstructure_Fold | ((((......))))(((.(....(((((....)))))...))))........ |
| RNAstructure_ProbKnot | ((((...<<<))))(((.(....(((((....)))))...))))>>>..... |
| RNAstructure_MaxExpect | ((((......))))(((.(....(((((....)))))...))))........ |

| | |
|---|---|
| DotKnot | (((((..[[[[[[))))).....(((((....)))))....]]]]]]..... |
| CONTRAfold | .......((((((........(.(((((....))))).)..))))))..... |
| MXfold2 | ((((......))))(((......(((((....)))))....)))........ |
| SPOT-RNA | .......((((((..........((((((..))))))....))))))..... |
| NUPACK | ((((......))))..(((....(((((....)))))...)))......... |
| UFold | ((((((.(((((()))))).....((((((..))))))).(.))))))).... |
| RNAPKplex | (((((..[[[[[[))))).....(((((....)))))....]]]]]]..... |
| CentroidFold_McCaskill | .......................(((((....))))).............. |
| CentroidFold_CONTRAfold | .......((((((........(.(((((....))))).)..))))))..... |
| HotKnots_CC09 | (((((..[[[[[[))))).....(((((....)))))....]]]]]]..... |
| HotKnots_DP09 | (((((..[[[[[[))))).....(((((....)))))....]]]]]]..... |
| LinearFold-C | (((((........))))).....(((((....))))).............. |
| LinearFold-V | ((((......))))(((......(((((....)))))....)))........ |
| UNAFold_mFold | ((((......))))..(((....(((((....)))))...)))......... |
| MC-Fold | (((((.[[[[[[[))))).((((((((((..))))))))))]]]]]]].... |
| RNAfold_MC-Fold | ((((.((((())))(((..((((((((((..))))))))))))))))))... |
| E2Efold | .((.(.((...(.))................(...)(.....)..).).)). |
| pKiss | [[[[[..{{{{{{]]]]].....(((((....)))))....}}}}}}..... |
| ContextFold | (((((........))))).....(((((....))))).............. |
| IPknot | [[[[[..((((((]]]]].....(((((....)))))....))))))..... |
| R2DT | .......((((((..........((((......))))....))))))..... |
| CentroidHomfold-LAST | (((((........)))))...(.(((((....))))).).....(......) |

**Table S7. Predicted 2D structures of PDB 6P2H sequence in dot-bracket notation.**

| | |
|---|---|
| PDB 6P2H | GGGUGUAAUCUCCAAAAUAUGGUUGGGGAGCCUCCACCAGUGAACCGUAAAAUCGCUGUCACCACCCAG |
| RNApdbee_3DNA-DSSR | ((((((..(((((((.....[[)))))))......).((((((.]]......))))))....))))).. |
| RNAfold_MFE | (((((...(((((((.......)))))))........((((((.........))))))....))))).. |
| RNAfold_MFE_60 | (((((...(((((((.......)))))))........((((((.........))))))....))))).. |
| RNAfold_MEA | (((((...(((((((.......)))))))........((((((.........))))))....))))).. |
| RNAstructure_Fold | (((((..............((((.(((.....)))))))((((.(.((......)).)))))))))).. |
| RNAstructure_ProbKnot | (((((......(((.....))).((((.....)))).((((((.........))))))....))))).. |
| RNAstructure_MaxExpect | (((((......((.......)).((((.....)))).((((((.........))))))....))))).. |
| DotKnot | (((((...((((((...[[[[[[))))))........((((((]]]]]]...))))))....))))).. |
| CONTRAfold | (((((...(((((((.......))))))).....)..((((((.........)))))).....)))).. |
| MXfold2 | (((((...(((((((.......)))))))........((((((.........))))))....))))).. |
| SPOT-RNA | (((((...(((((((......<)))))))).......>((((((.........))))))....))))).. |
| NUPACK | (((.((..(((((((.......))))))))).)))..((((((.........))))))........... |
| UFold | (((((((((((((((.(((.(()))))))).....).((((((())()))))))))))))..())))).) |
| RNAPKplex | (((((...((((((...[[[[[[))))))........((((((]]]]]]...))))))....))))).. |
| CentroidFold_McCaskill | (((((...................((....)).....(((((...........)))))....))))).. |
| CentroidFold_CONTRAfold | (((((...(((((((.......)))))))........((((((.........))))))....))))).. |
| HotKnots_CC09 | (((((...(((((((.......)))))))........((((((.........))))))....))))).. |
| HotKnots_DP09 | (((((...(((((((.......)))))))........((((((.........))))))....))))).. |
| LinearFold-C | .((((...(((((((.......))))))).....)))).............................. |
| LinearFold-V | (((((...(((((((.......)))))))........((((((.........))))))....))))).. |
| UNAFold_mFold | (((((...(((((((.......)))))))........((((((.........))))))....))))).. |
| MC-Fold | (((((((((((((((((.[[[[)))))))))))))))))(((((((((....))))).))))...]]]] |

| | |
|---|---|
| RNAfold_MC-Fold | (((((((((((((((((...))))))))))..)))).((((((.((((((..))))))))))))))).. |
| E2Efold | .(..((...(.(.........)...)(..(.......)..)........(..(.....).)..)).).. |
| pKiss | ........[[[[[[[..{{{{{]]]]]]]........<<<<<<.}}}}}...>>>>>>........... |
| ContextFold | (((((...(((((((.......)))))))........((((((.........))))))....))))).. |
| ipknot | .((.....(((((((....[[[)))))))........((((((.]]].....))))))......))... |
| R2DT | (((((...(((((((.......))))))).........((((((.......)))))).....))))).. |
| CentroidHomfold-LAST | (((((...(((((((.......)))))))........((((((.........))))))....))))).. |

#### Table S8. Predicted 2D structures of PDB 3OWZ_A sequence in dot-bracket notation.

| | |
|---|---|
| **3OWZ_A considered in standardize PDB** | **CUCUGGAGAGAACCGUUUAAUCGGUCGCCGAAGGAGCAAGCUCUGCGCAUAUGCAGAGUG-<br>AAACUCUCAGGCAAAAGGACAGAG** |
| **PDB 3OWZ_A** | **GGCUCUGGAGAGAACCGUUUAAUCGGUCGCCGAAGGAGCAAGCUCUGCGGAAACGCAGAGUG-<br>AAACUCUCAGGCAAAAGGACAGAGUC** |
| **RNApdbee_3DNA-DSSR** | **(((((((....(.((((......)))))(((...((((...((((((((....))))))))....))).).))).......<br>)))))))** |
| RNAfold_MFE | (((((((......((((......)))).(((...((((...((((((((....))))))))....))))..))).......<br>))))))) |
| RNAfold_MFE_60 | (((((((......((((......)))).(((...((((...((((((((....))))))))....))))..))).......<br>))))))) |
| RNAfold_MEA | (((((((....(.((((......)))))(((...((((...((((((((....))))))))....))))..))).......<br>))))))) |
| RNAstructure_Fold | (((((((......((((......)))).(((...((((...((((((((....))))))))....))))..))).......<br>))))))) |
| RNAstructure_ProbKnot | (((((((....(.(((((...).)))))((((..((((...((((((((....))))))))....))))).))).......<br>))))))) |

| | |
|---|---|
| RNAstructure_MaxExpect | (((((((......((((......)))).(((...((((...((((((((....))))))))....))))..))).......))))))) |
| DotKnot | (((((((......((((......)))).(((...((((...((((((((....))))))))....))))..))).......))))))) |
| CONTRAfold | (((((((....(.((((......)))))(((...((((...((((((((....))))))))....))))..))).......))))))) |
| MXfold2 | (((((((....(.((((......)))))(((...((((...((((((((....))))))))....))))..))).......))))))) |
| SPOT-RNA | ((((((((...(.((((......)))))((((..(((((.((((((((((..))))))))))..)))))).)))......))))))) |
| NUPACK | (((((((......((((......)))).(((...((((...((((((((....))))))))....))))..))).......))))))) |
| UFold | (((((((......((((......)))).......((((...((((((((....))))))))....))))............))))))) |
| RNAPKplex | (((((((......((((......)))).(((...((((...((((((((....))))))))....))))..))).......))))))) |
| CentroidFold_McCaskill | ((((((((((((.((((......)))).(((....).))..((((((((....))))))))....)))))...(....)..))))))) |
| CentroidFold_CONTRAfold | (((((((......((((......)))).(((...((((...((((((((....))))))))....))))..))).......))))))) |
| HotKnots_CC09 | (((((((......((((......)))).(((...((((...((((((((....))))))))....))))..))).......))))))) |
| HotKnots_DP09 | (((((((......((((......)))).(((...((((...((((((((....))))))))....))))..))).......))))))) |
| LinearFold-C | (((((((......((((......)))).(((...((((...((((((((....))))))))....))))..))).......))))))) |
| LinearFold-V | (((((((......((((......)))).(((...((((...((((((((....))))))))....))))..))).......))))))) |

| | |
|---|---|
| UNAFold_mFold | (((((((......((((......)))).(((...((((...((((((((....))))))))....))))..)))....... ))))))) |
| MC-Fold | No output |
| Mcfold + RNA-fold | ((((((((((.(.((((((..)))))))((((((((((((((((((((((..))))))))))))))))))))))....))) ))))))) |
| E2Efold | ((((((...(.(...........)...).(...........).....(.........)........(.........)..)) )))..). |
| pKiss | .[[[[[.......((((......)))).{{{...]]]]]..((((((((....))))))))...<<<<<..}}}....... .>>>>>. |
| ContextFold | (((((((...(..((((......)))).(((...((((...((((((((....))))))))....))))..)))...)... ))))))) |
| IPknot | (((((((......((((......)))).(((...((((...((((((((....))))))))....))))..)))....... ))))))) |
| R2DT | (((((((......((((......)))).(((((..(((...((................))....))))).)))....... ))))))) |
| CentroidHom-fold-LAST | .(((((((((((.((((......))))((((....).))..((((((((....)))))))))...)))))...(....).. ))))))). |

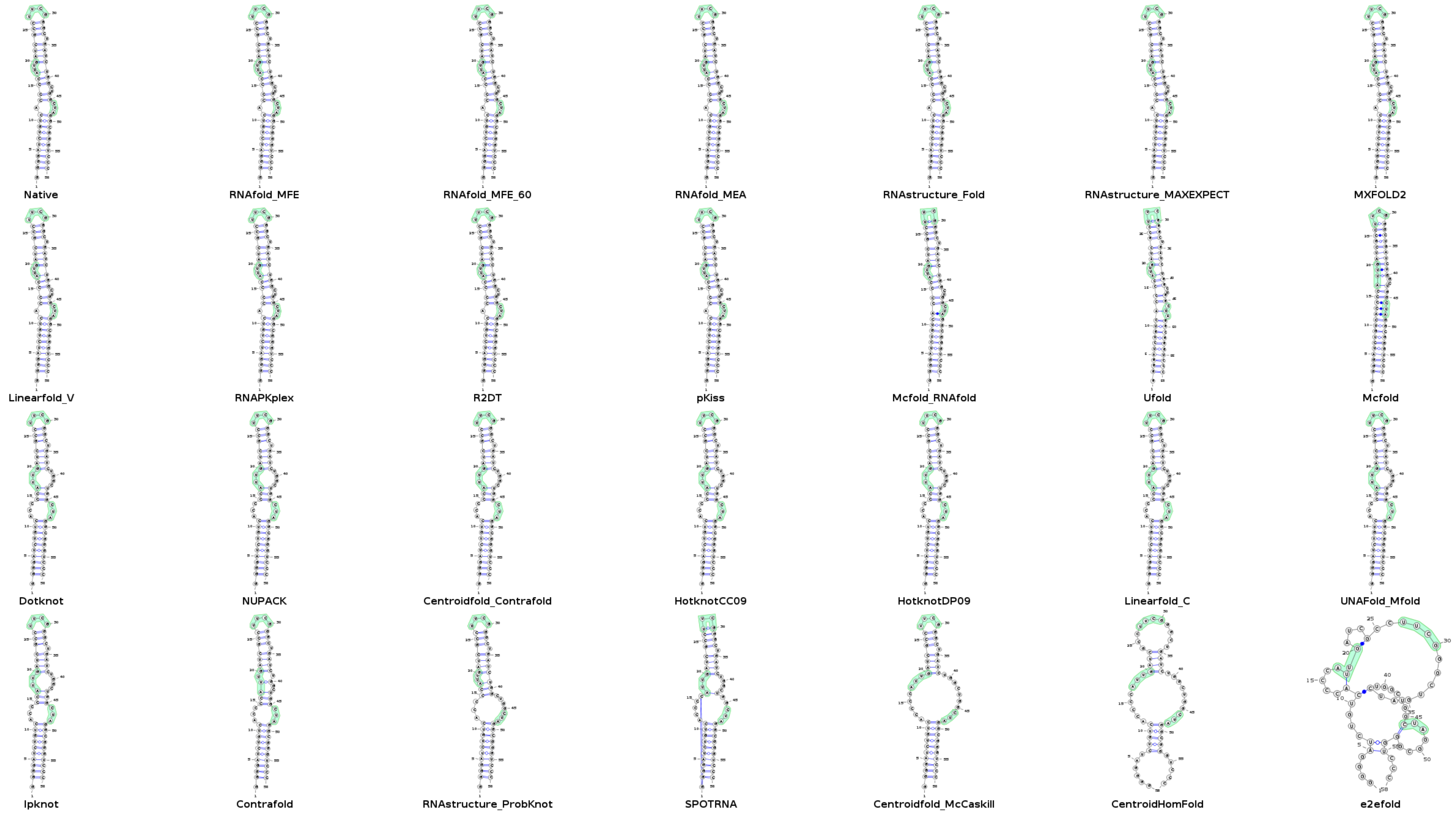


**Figure S1. P**redicted secondary structures of PDB 5LYU plotted with VARNA.

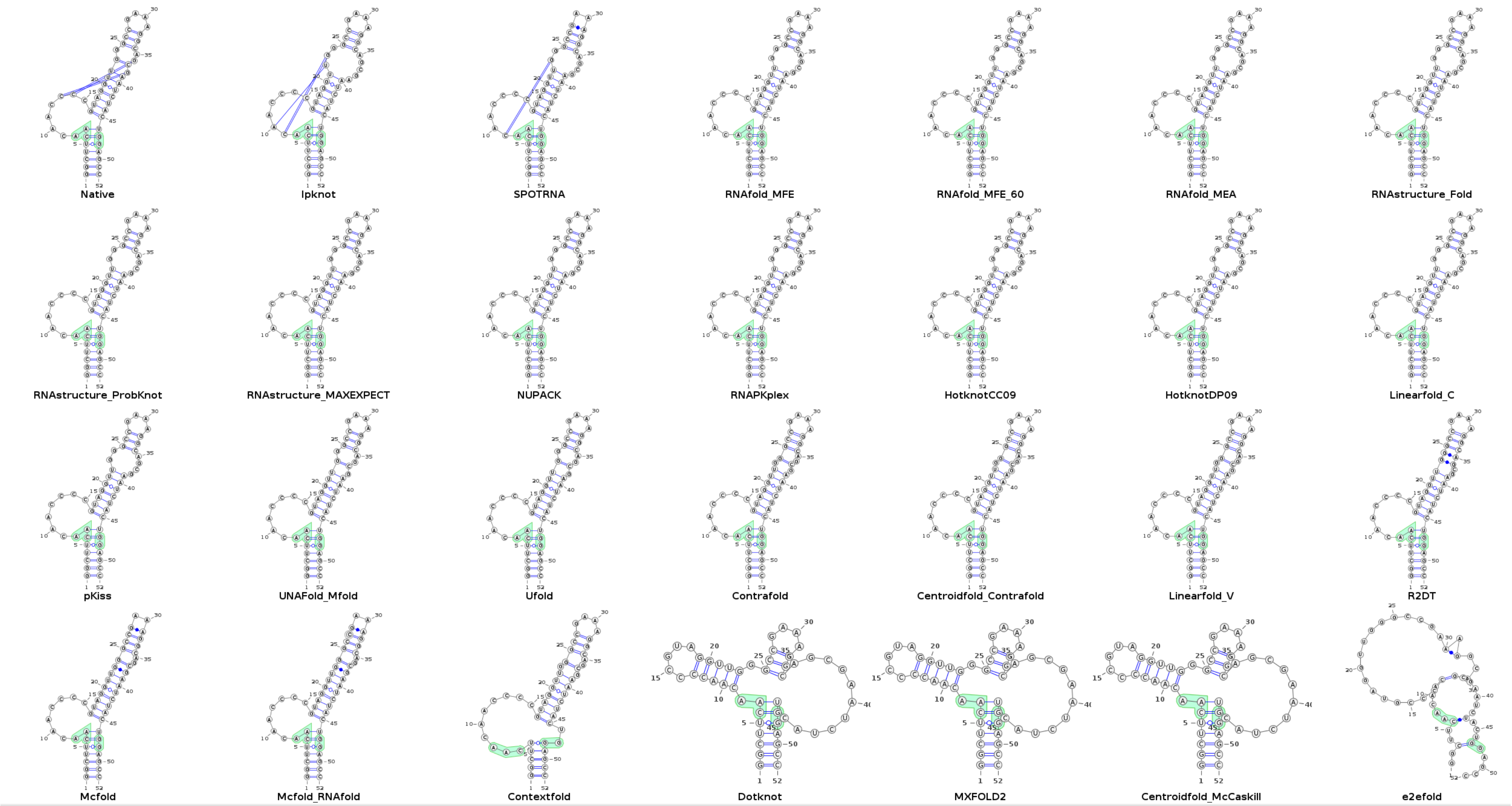


**Figure S2.** Predicted secondary structures of PDB 6TB7 plotted with VARNA.

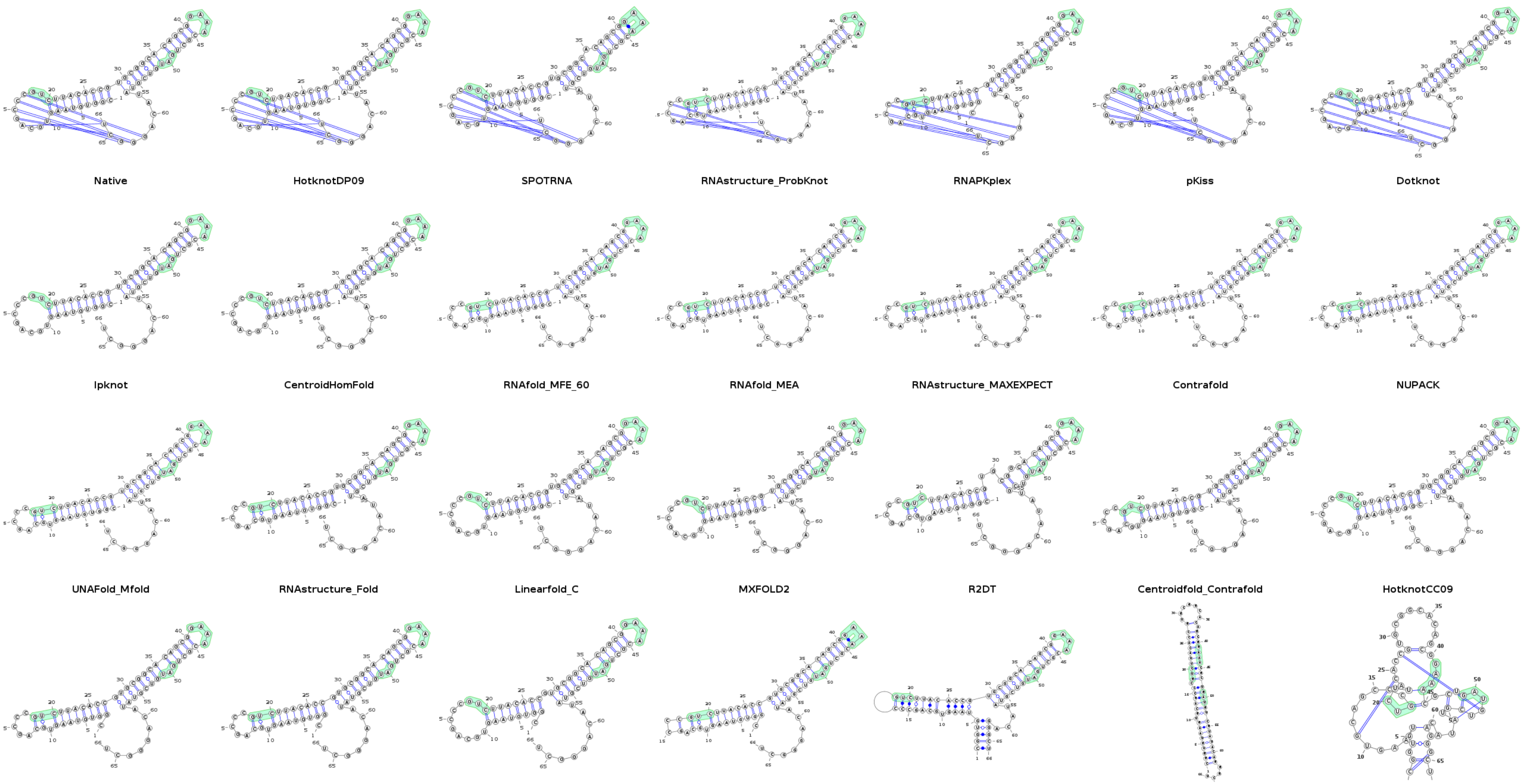


**Figure S3.** Predicted secondary structures of PDB 7LYJ plotted with VARNA.

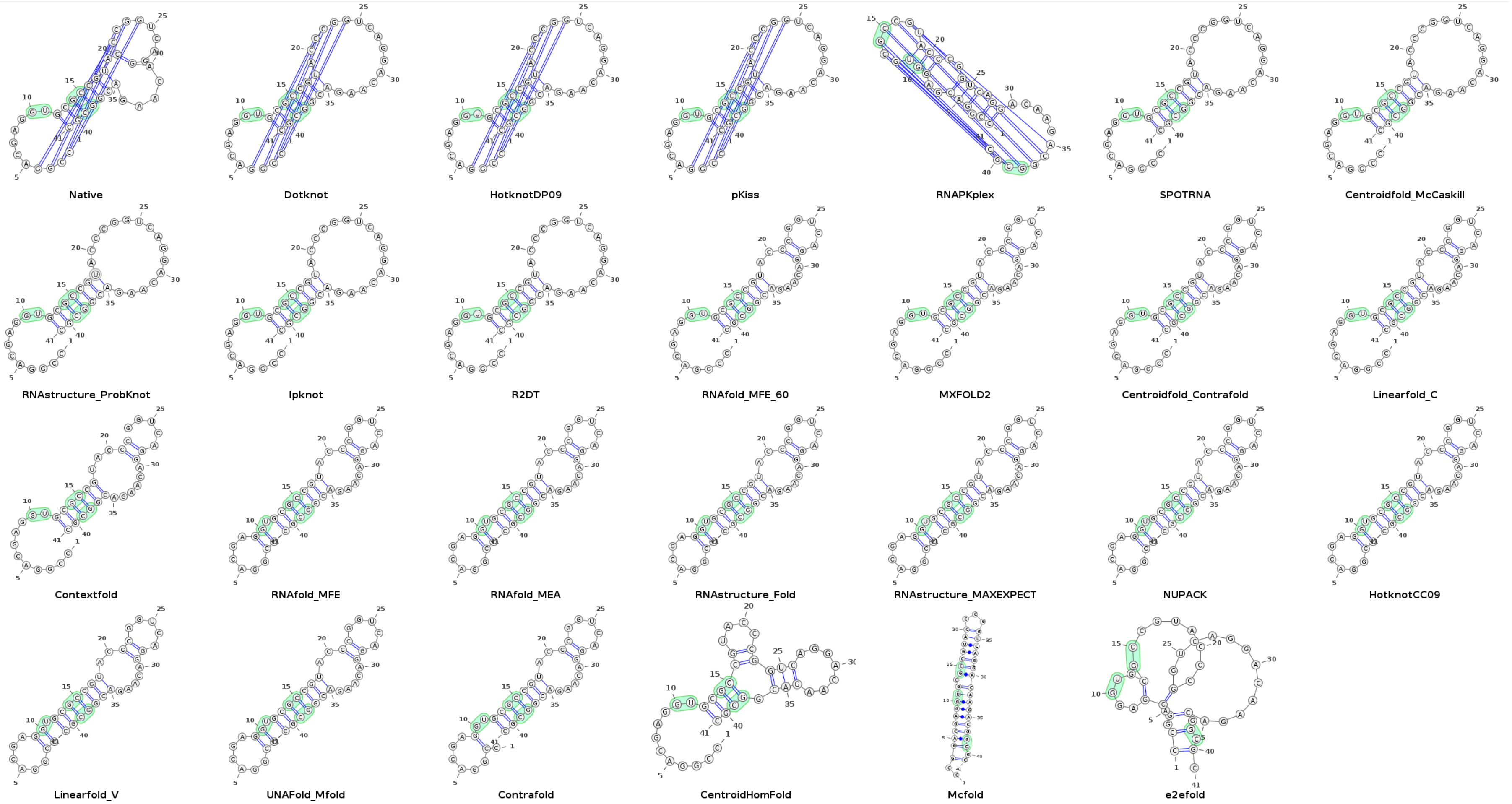


**Figure S4.** Predicted secondary structures of PDB 5NWQ plotted with VARNA.

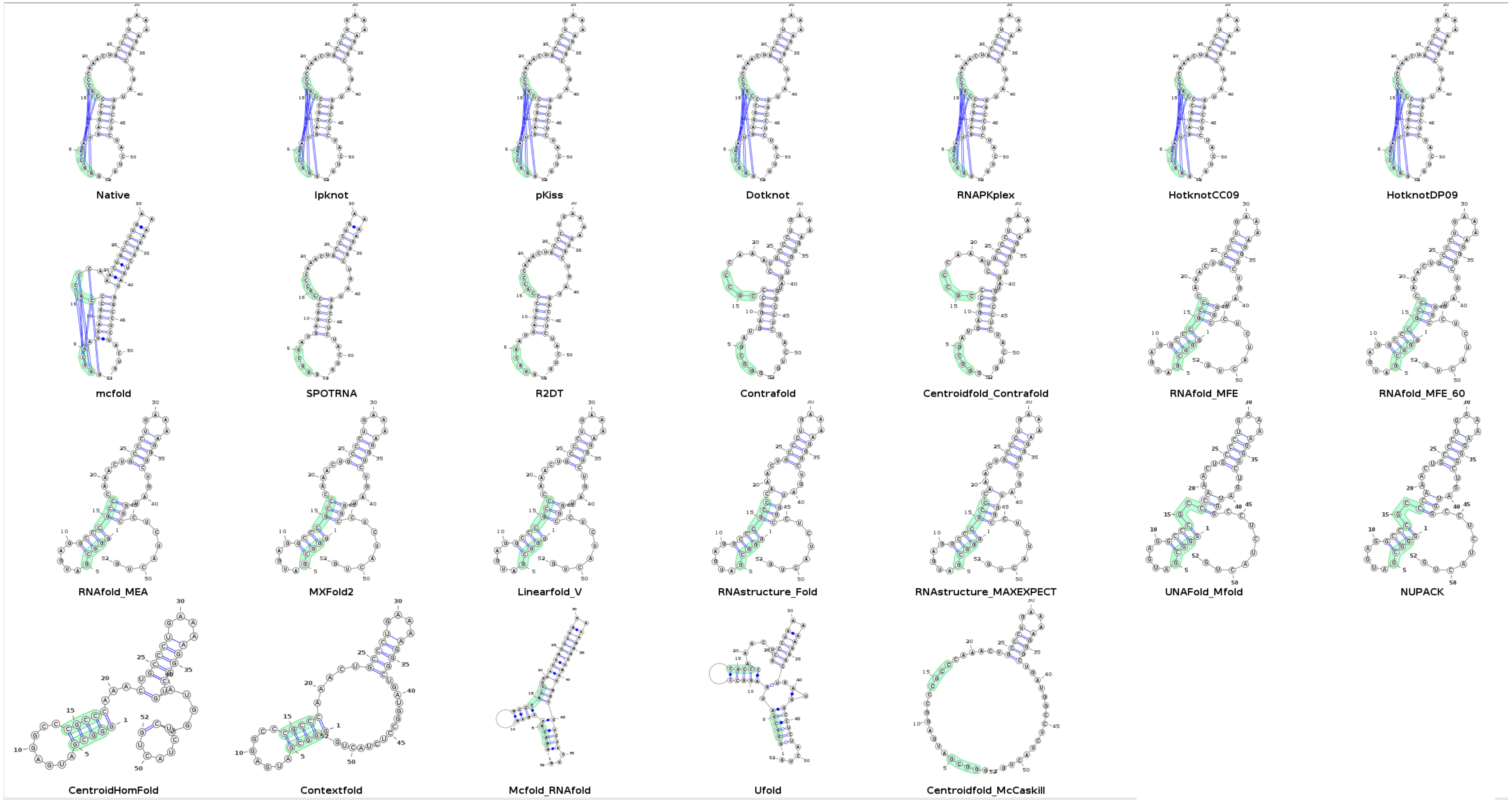


**Figure S5. P**redicted secondary structures of PDB 4ENC plotted with VARNA.

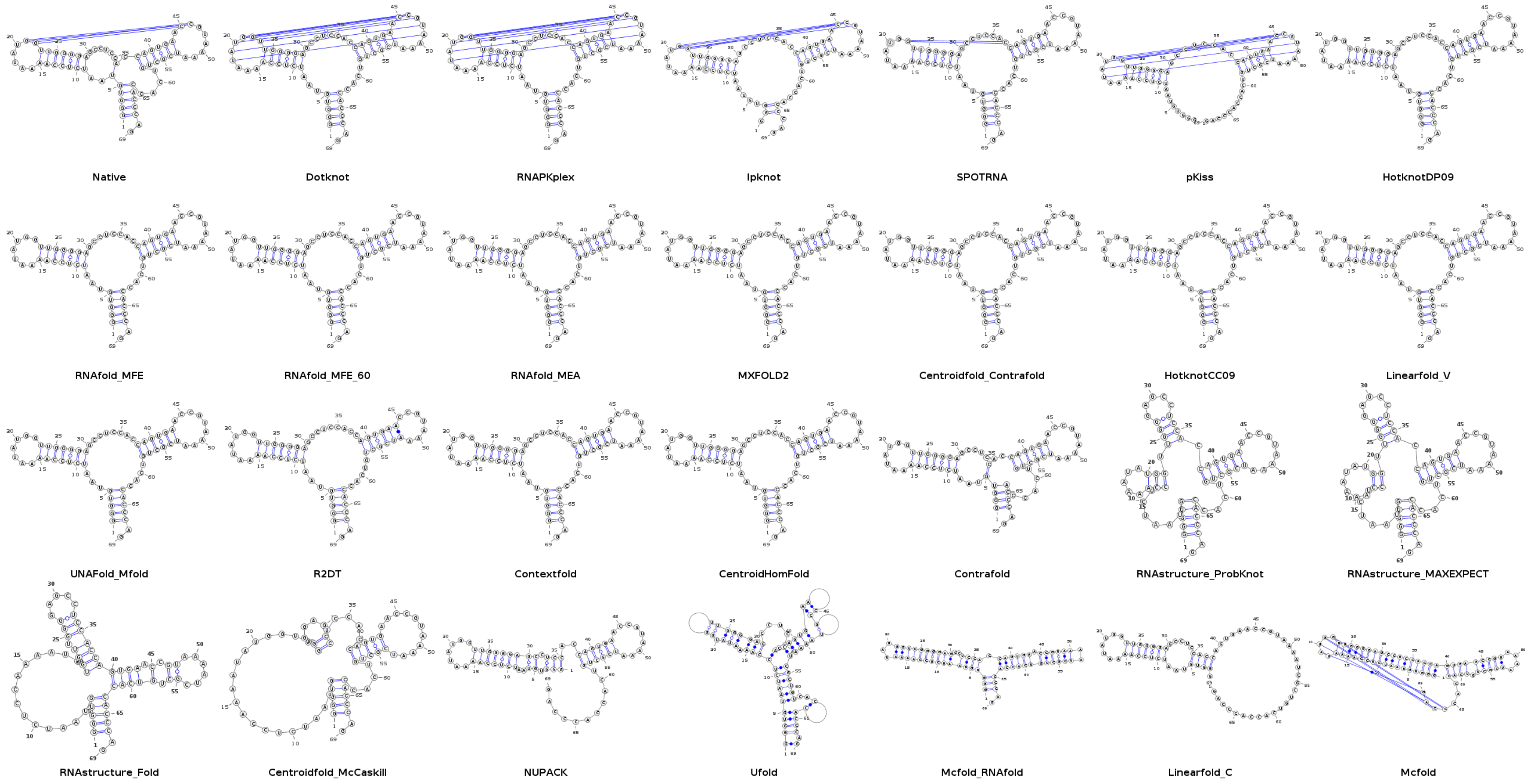


**Figure S6. P**redicted secondary structures of PDB 6P2H plotted with VARNA.

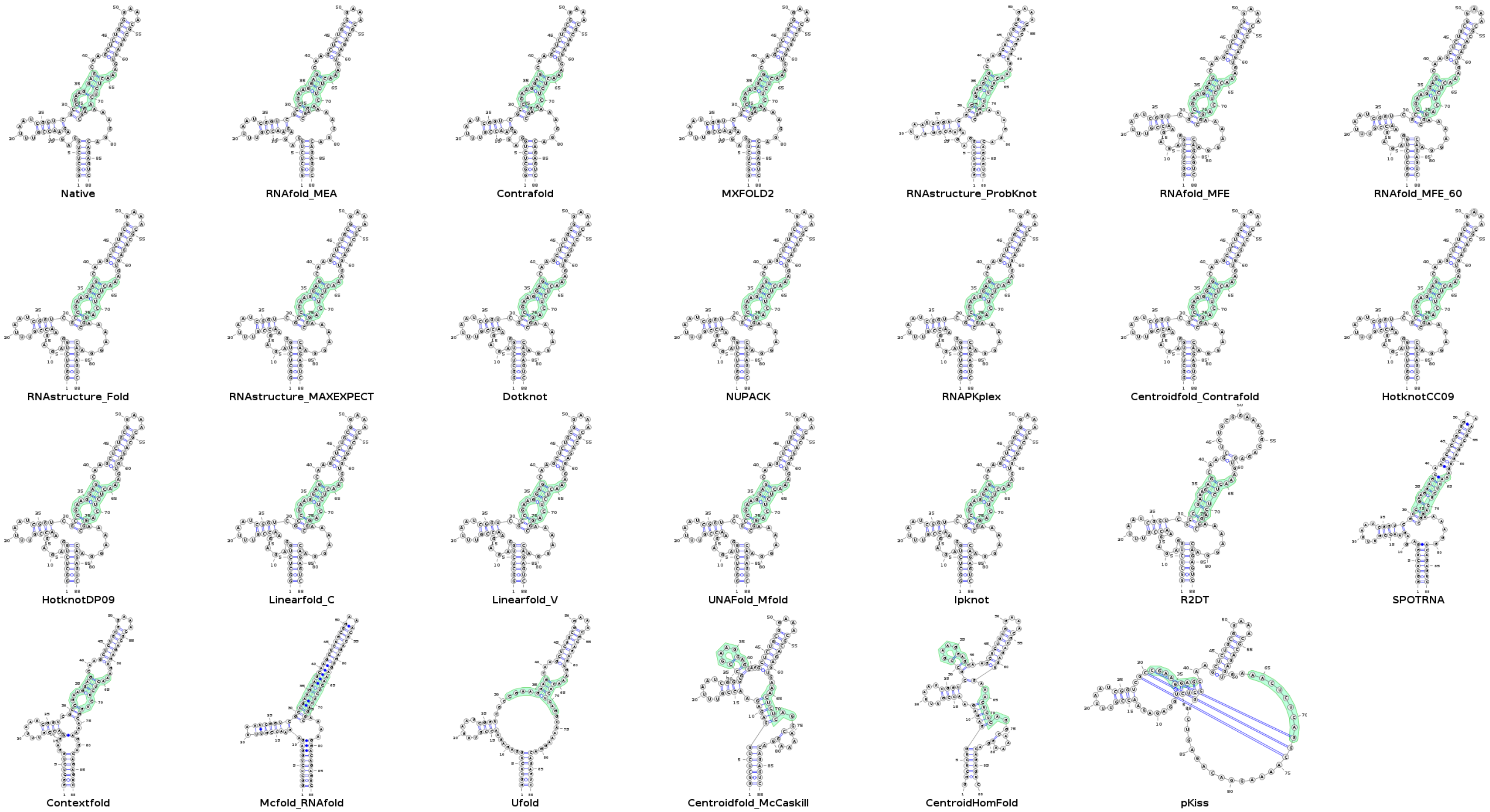


**Figure S7.** Predicted secondary structures of PDB 3OWZ_A plotted with VARNA.

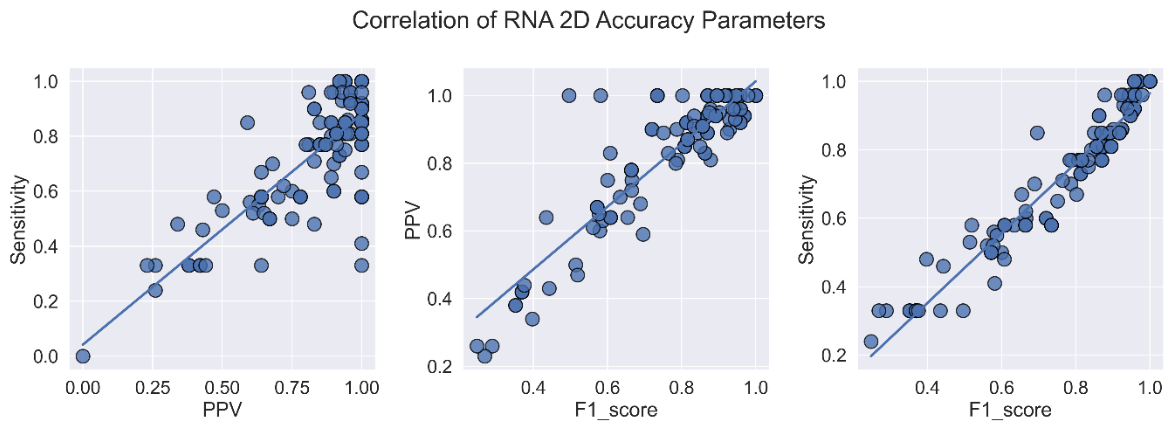


**Figure S8.** Correlation plot between PPV, sensitivity, and F1-score for all predicted secondary structures of seven sequences (correlation coefficients in **Figure S12**)

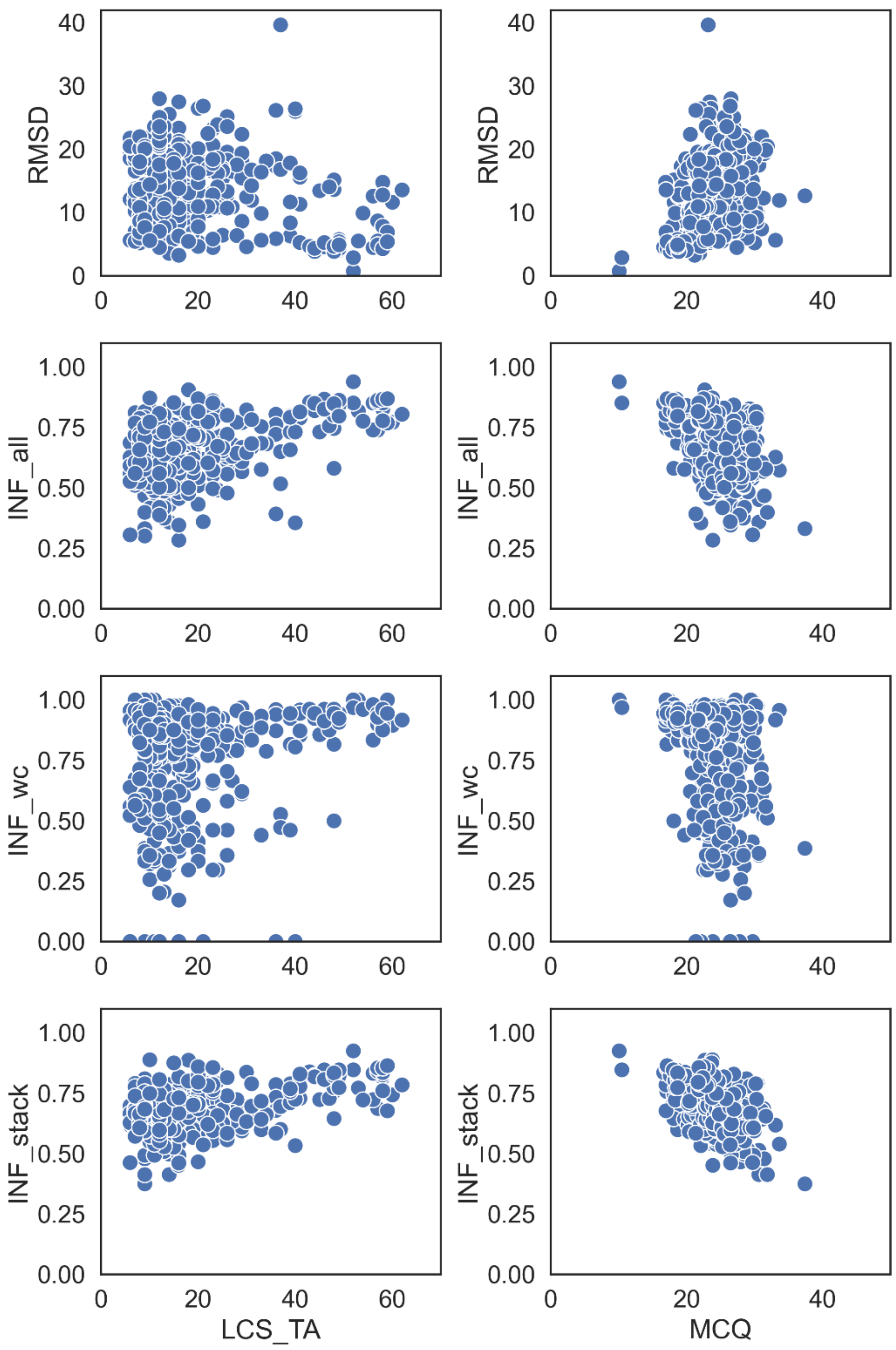


**Figure S9.** Comparison of RNA 3D structure accuracy parameters RMSD, INF_all, INF_wc, INF_stack to the torsional accuracy parameters LCS_TA and MCQ.

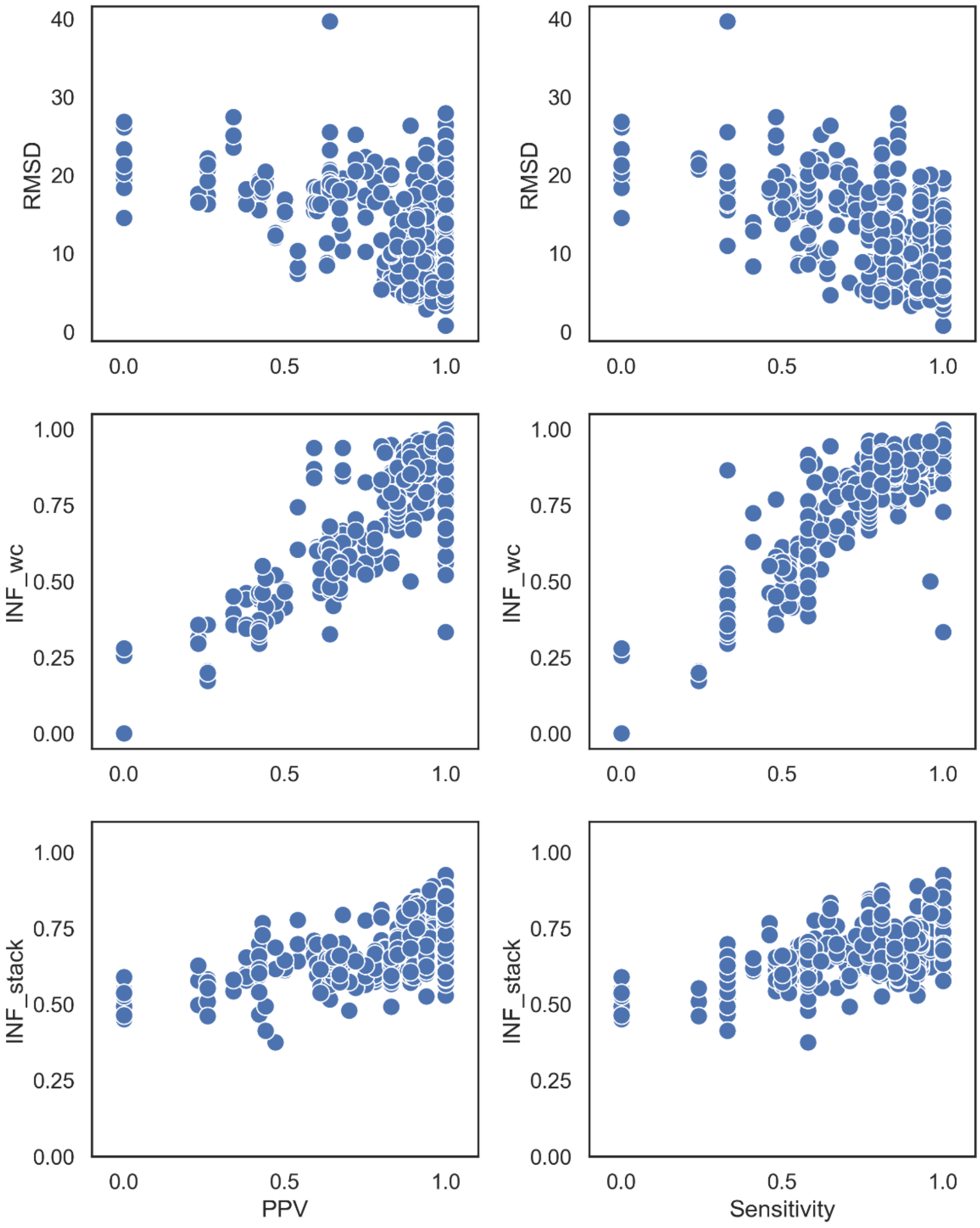


**Figure S10.** Comparison of RNA secondary structure accuracy parameters PPV and Sensitivity with 3D structure quality parameters RMSD, INF_wc, INF_stack.

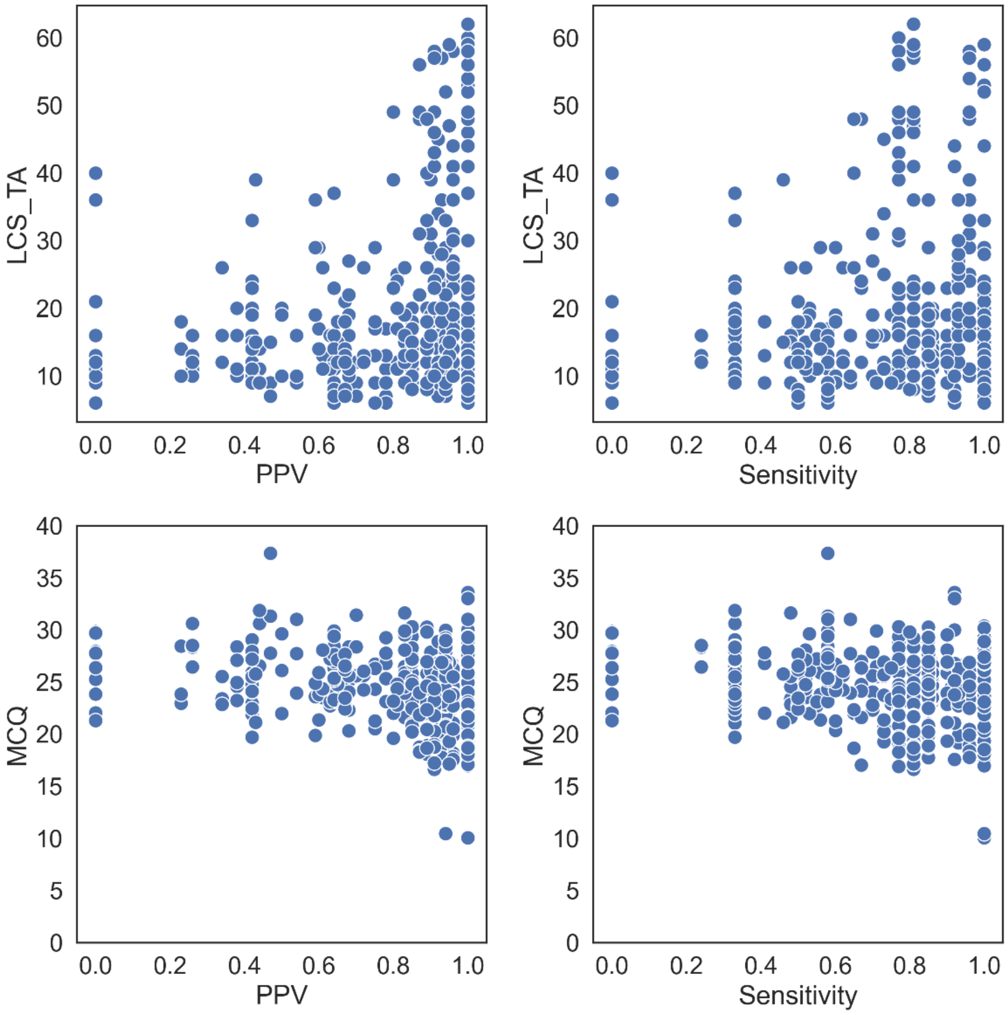


**Figure S11.** Comparison of RNA secondary structure accuracy parameters PPV and Sensitivity with 3D structure quality parameters LCS_TA and MCQ.

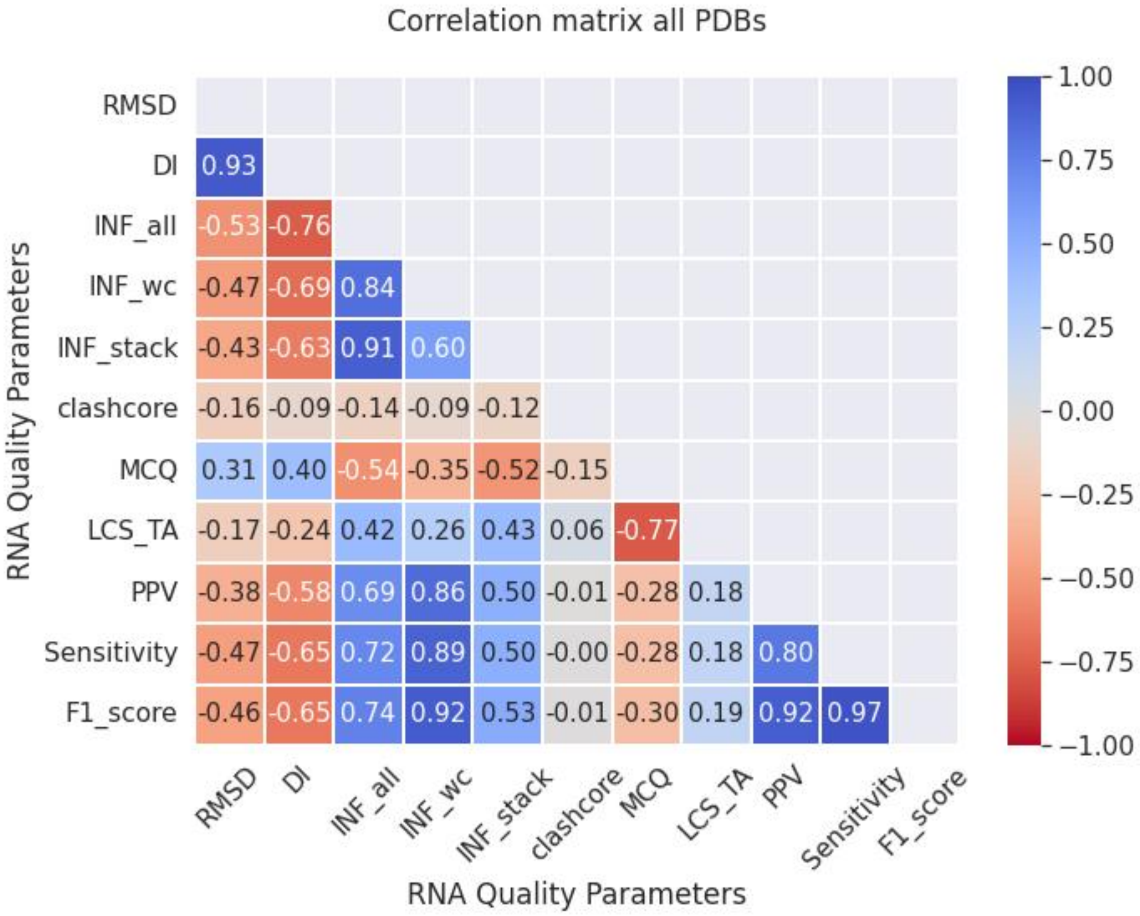


**Figure S12.** Correlation matrix for secondary structure accuracy parameters (PPV, Sensitivity, F1-score) and 3D structure parameters (RMSD, Deformation Index (DI), INF_all, INF_wc, INF_stack, clashscore, MCQ, LCS_TA )

### PDB 5LYU

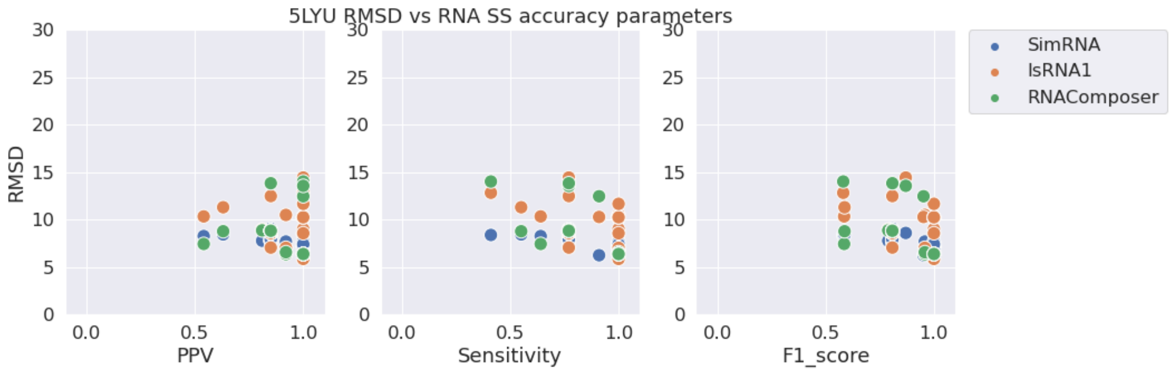


**Figure S13.** RMSD vs. PPV (left), Sensitivity (middle), and F1-score (right) for stem-loop structure PDB 5LYU.

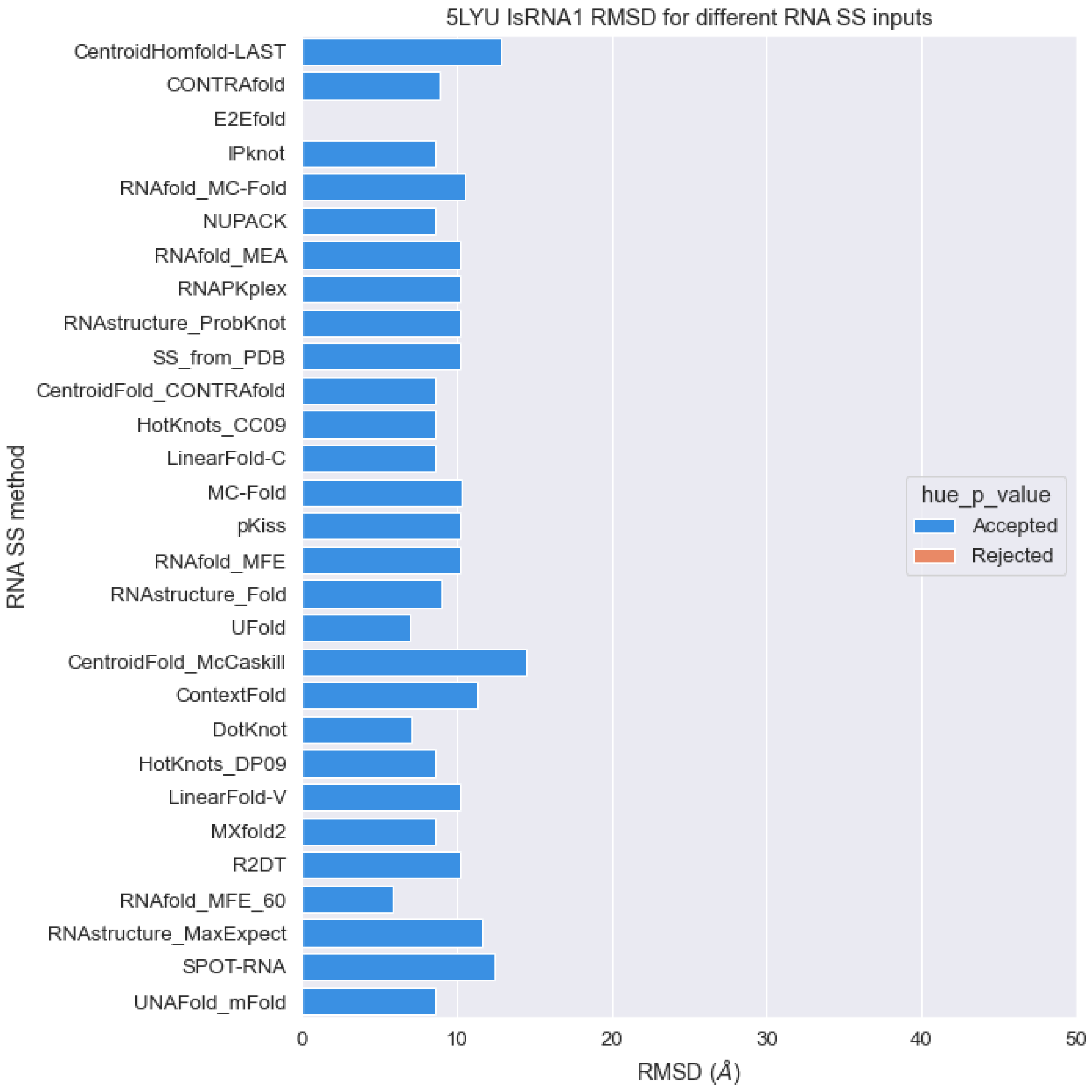


**Figure S14.** RMSD values of predicted structures for PDB 5LYU sequence with different RNA secondary structures as constraints with IsRNA1 package (orange: p-value > 0.01, blue: p-value < 0.01).

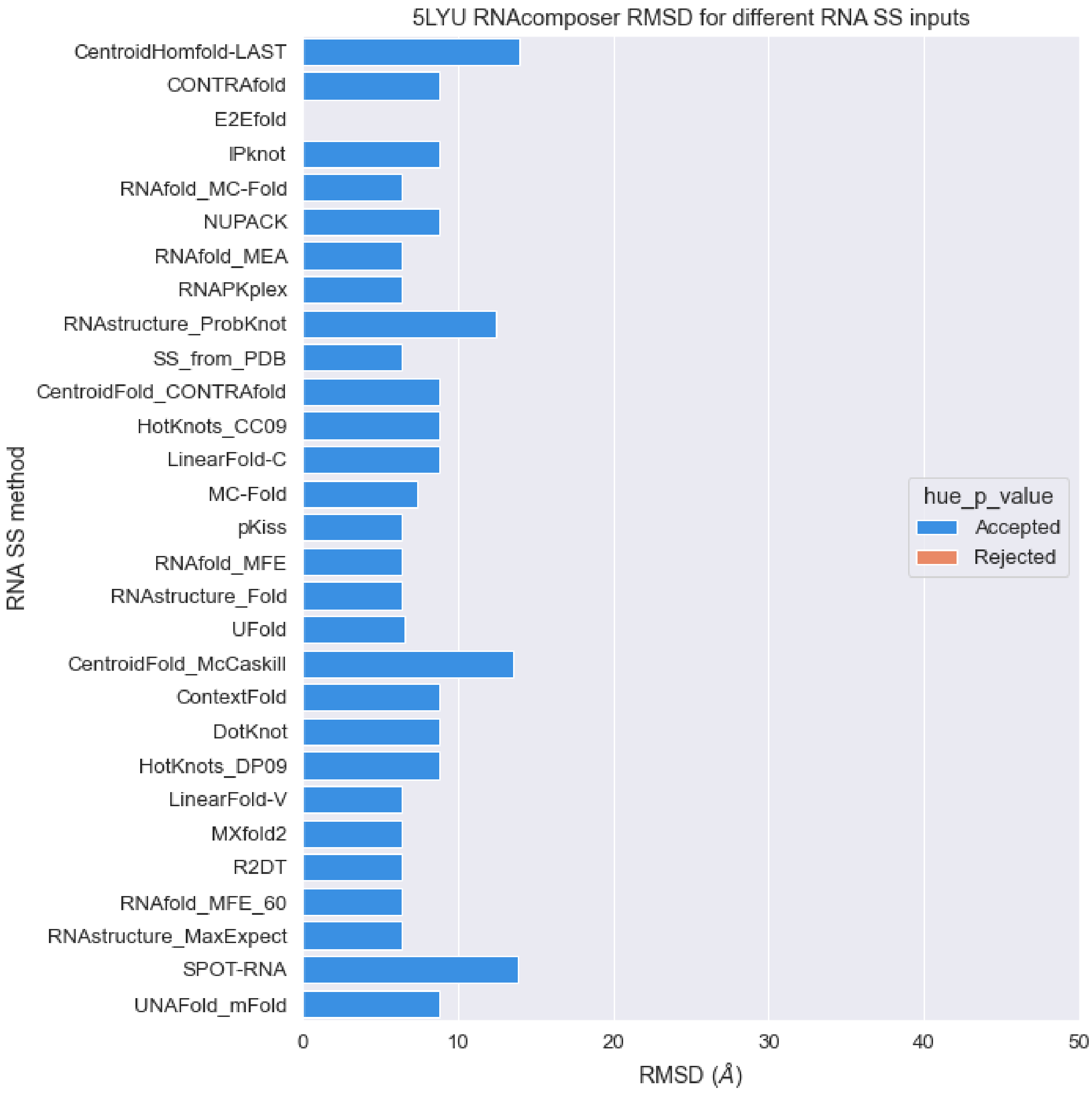


**Figure S15.** RMSD values of predicted structures for PDB 5LYU sequence with different RNA secondary structures as constraints with RNAComposer package (orange: p-value > 0.01, blue: p-value < 0.01).

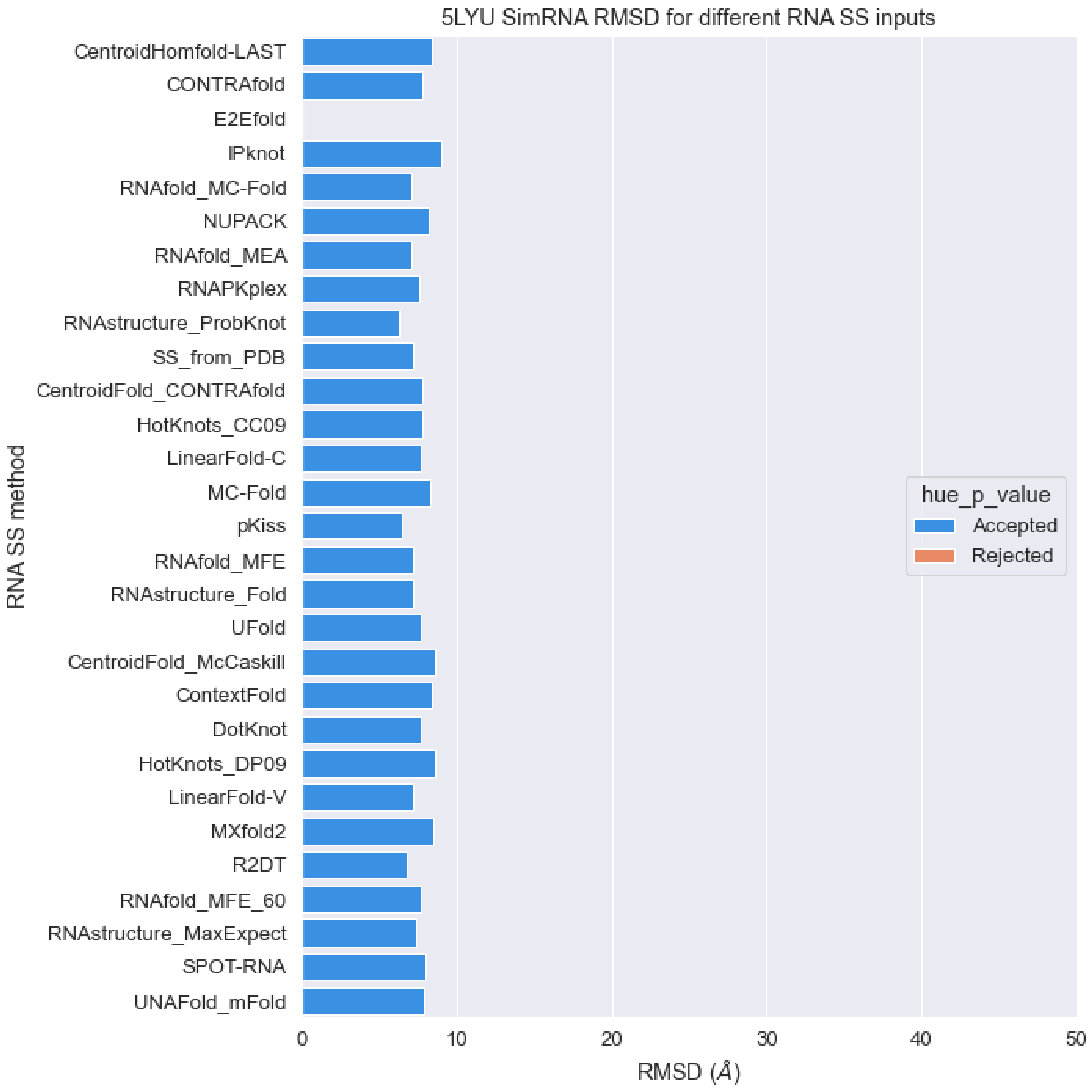


**Figure S16.** RMSD values of predicted structures for PDB 5LYU sequence with different RNA secondary structures as constraints with SimRNA package (orange: p-value > 0.01, blue: p-value < 0.01).

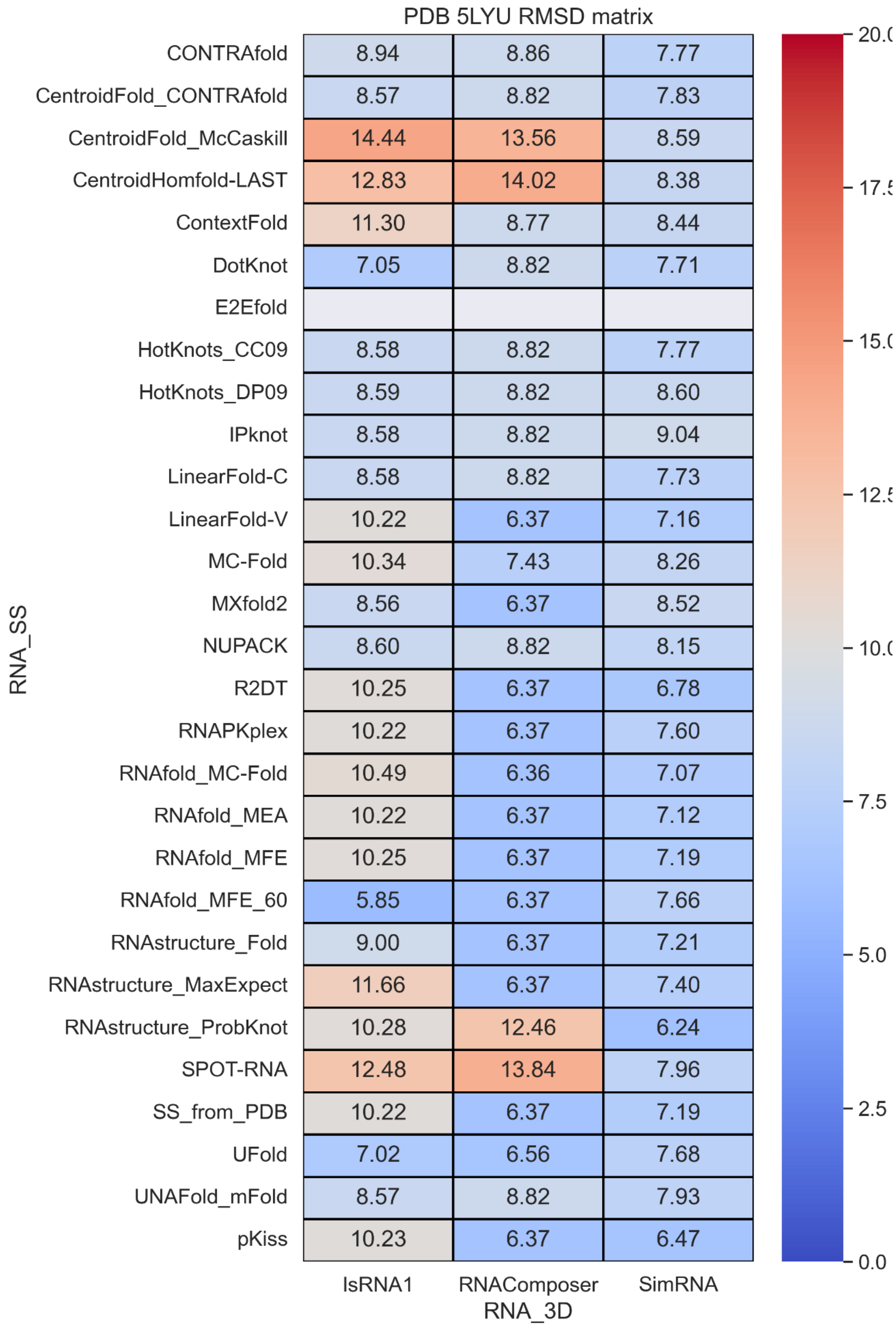


**Figure S17.** RMSD values with different RNA secondary structure methods and RNA 3D methods.

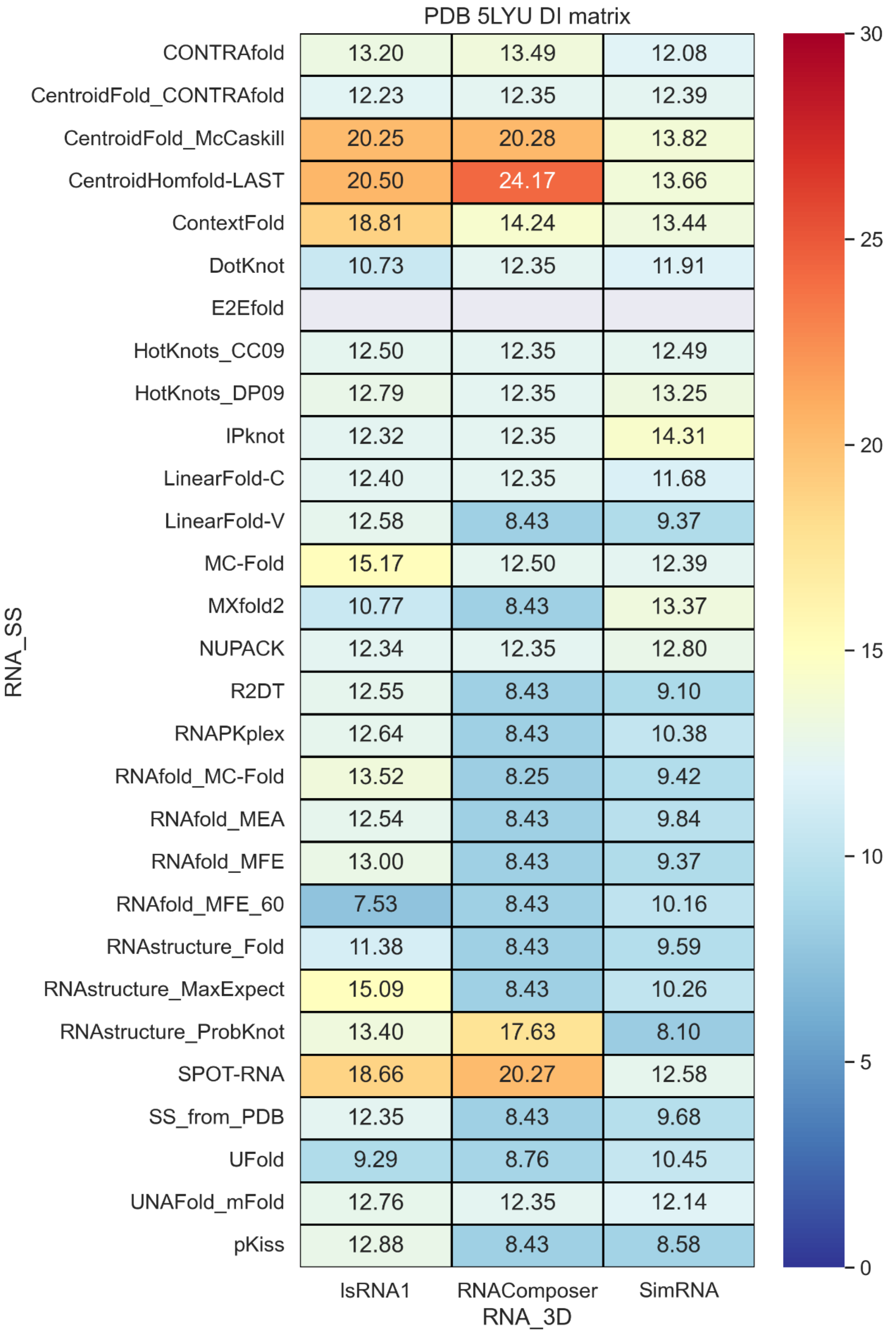


**Figure S18.** Deformation Index values with different RNA secondary structure methods and RNA 3D methods.

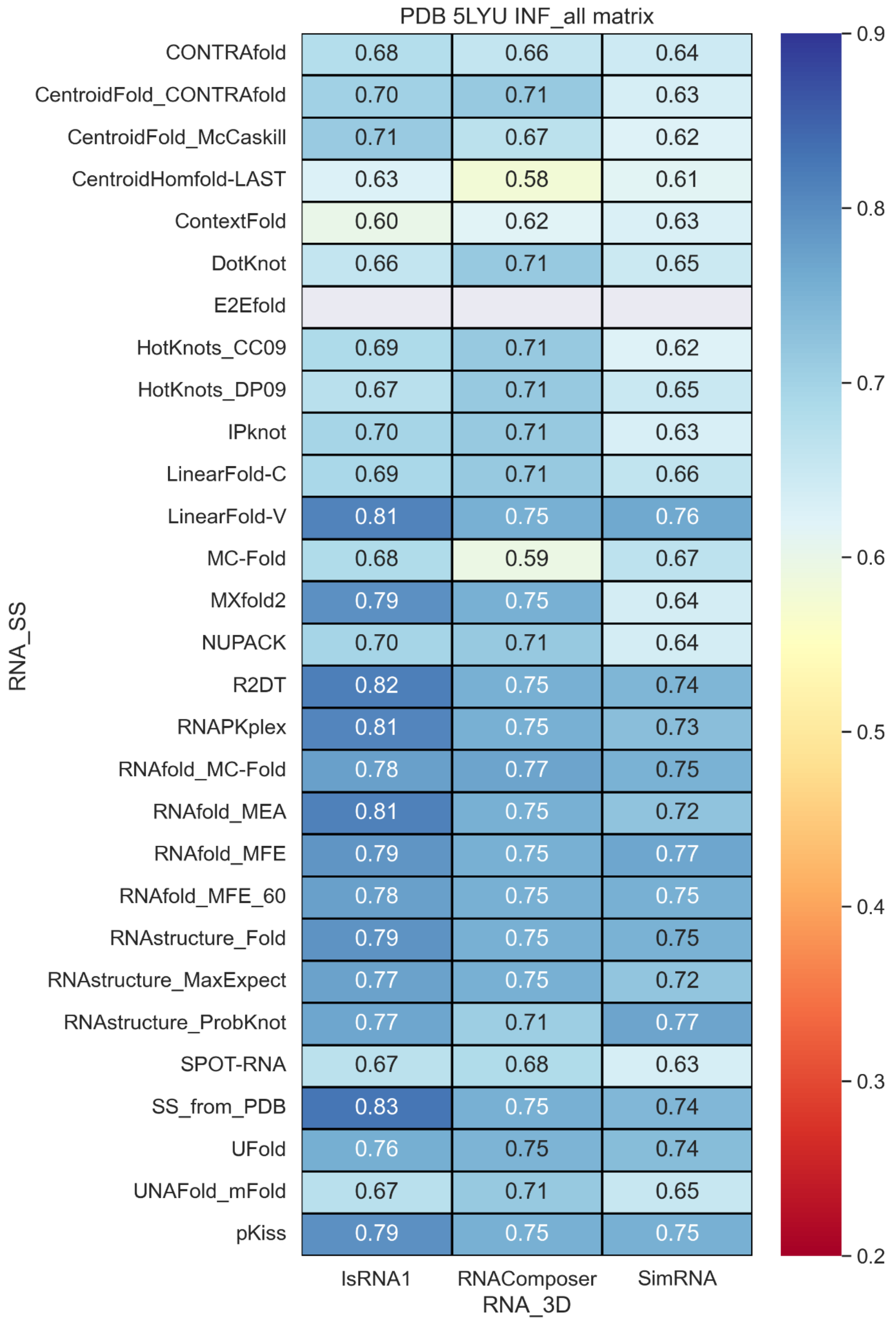


**Figure S19.** INF_all values with different RNA secondary structure methods and RNA 3D methods.

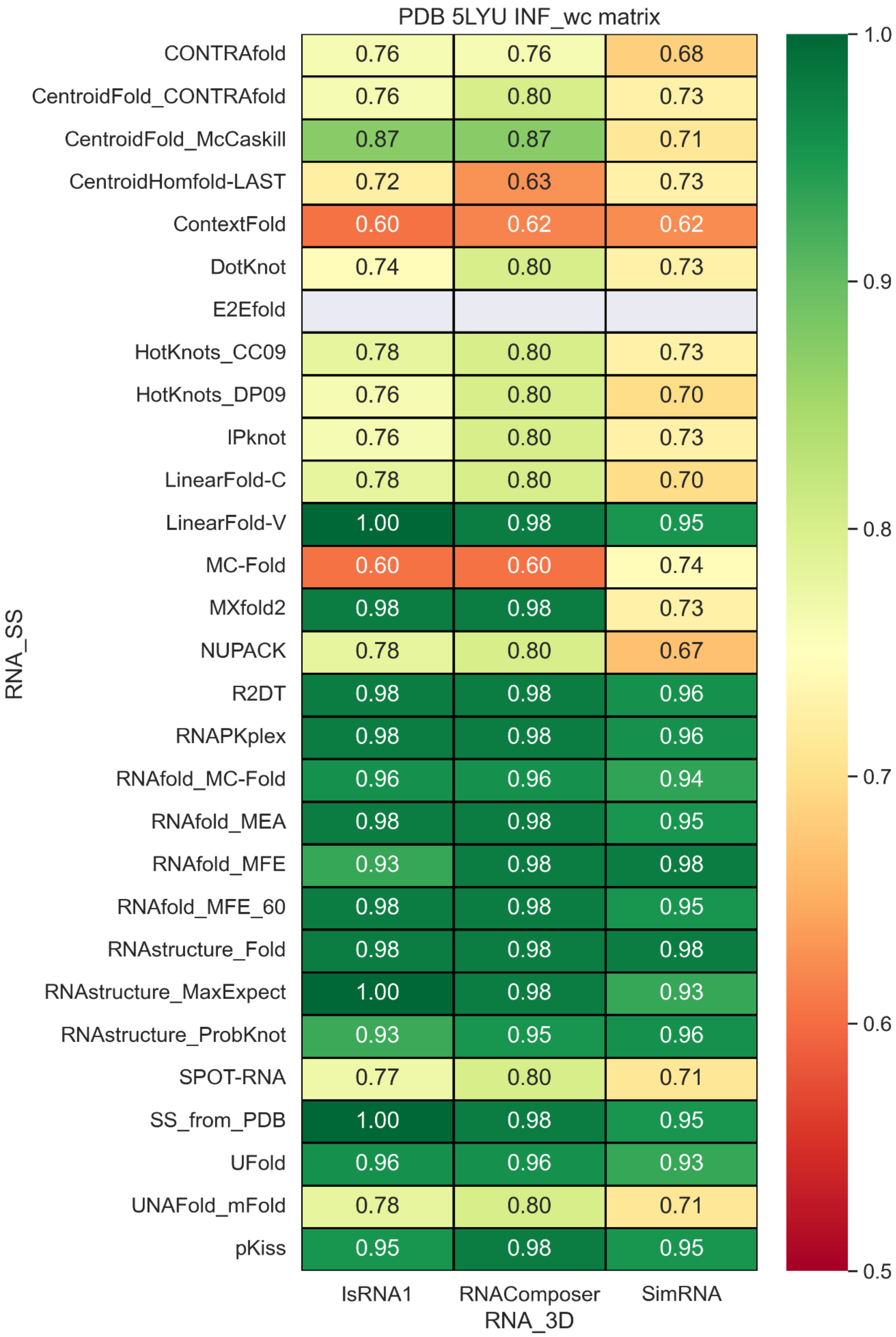


**Figure S20.** INF_wc values with different RNA secondary structure methods and RNA 3D methods.

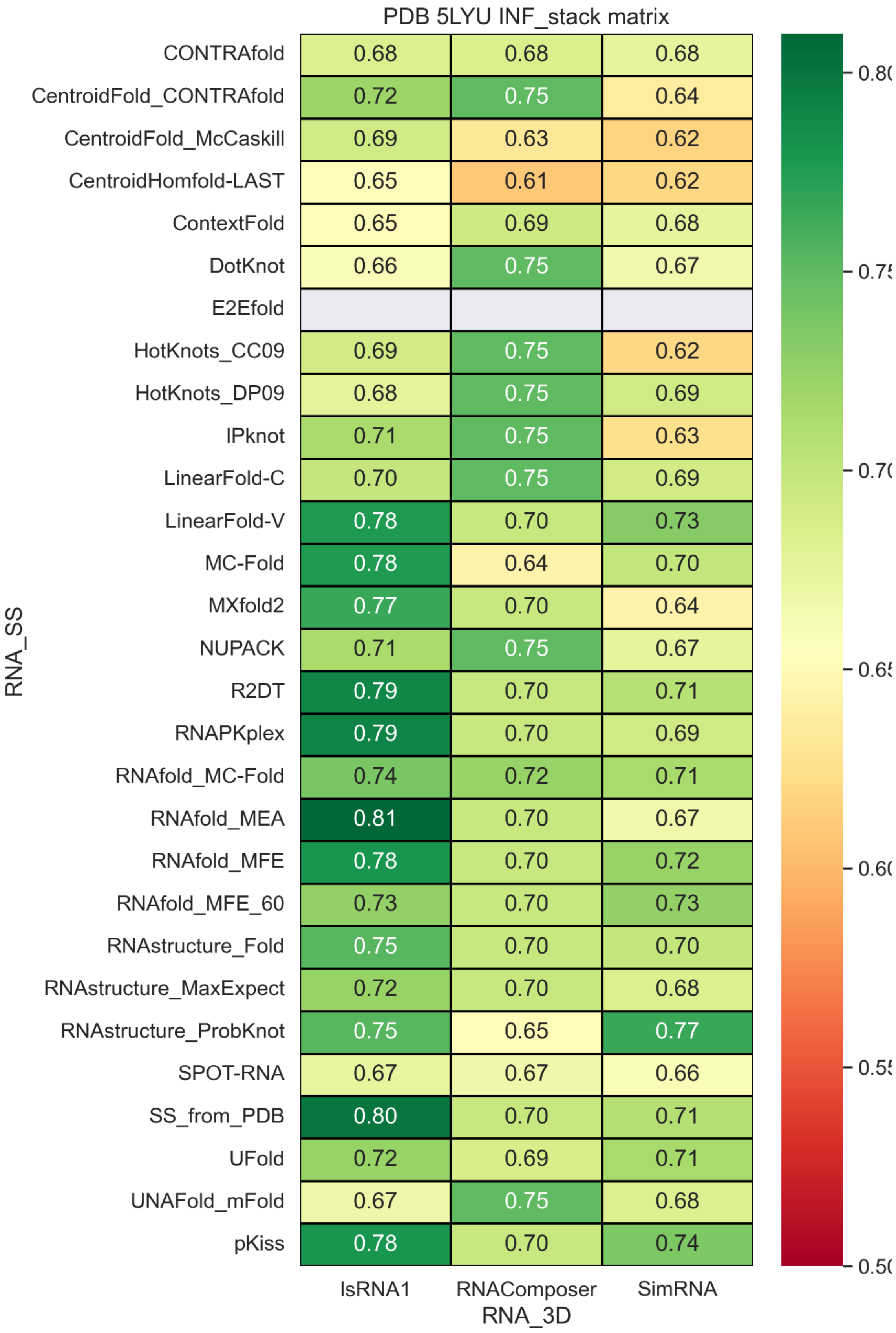


**Figure S21.** INF_stack values with different RNA secondary structure methods and RNA 3D methods.

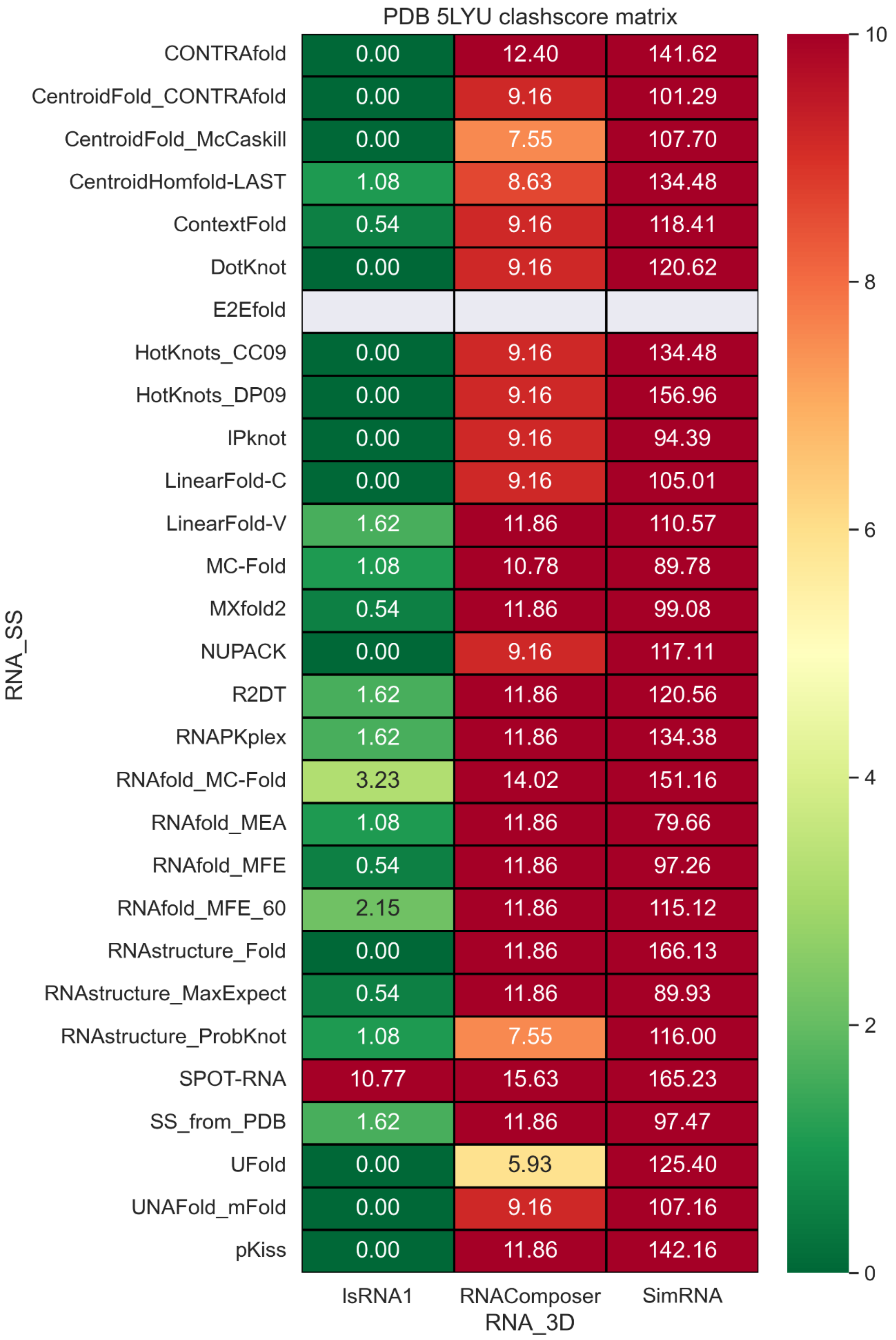


**Figure S22.** Clashscore values with different RNA secondary structure methods and RNA 3D methods.

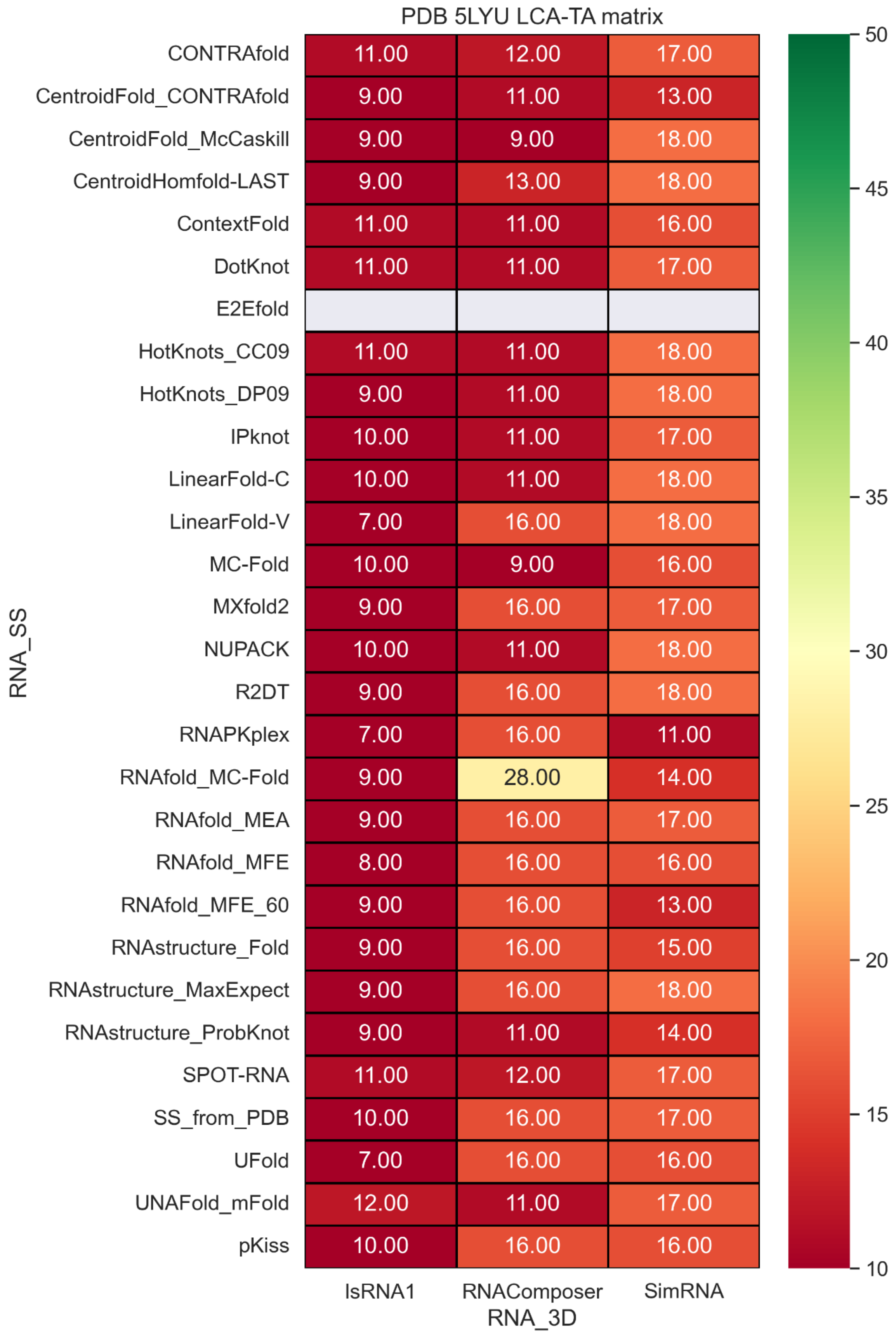


**Figure S23.** LCA-TA values with different RNA secondary structure methods and RNA 3D methods.

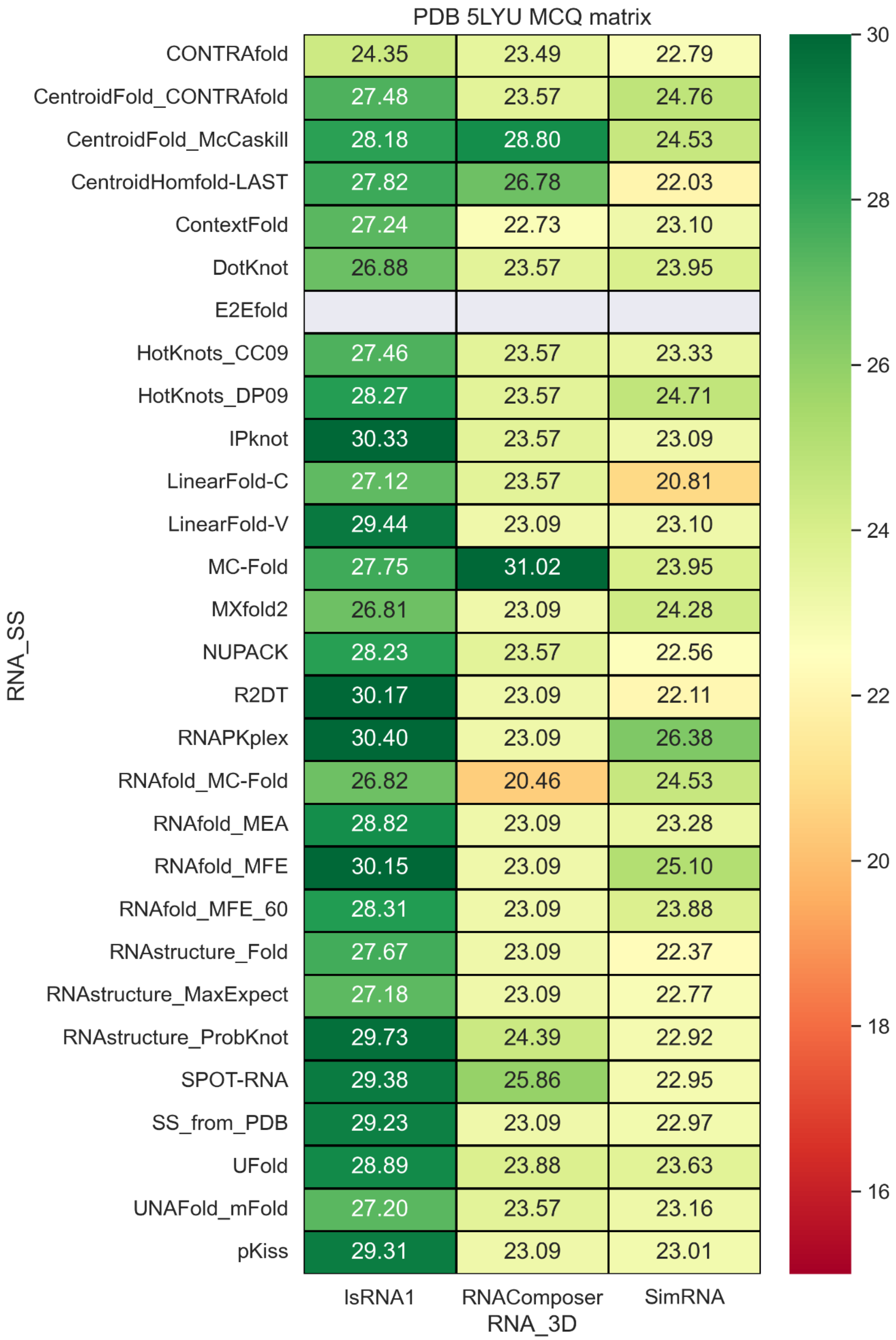


**Figure S24.** MCQ values with different RNA secondary structure methods and RNA 3D methods.

### PDB 6TB7

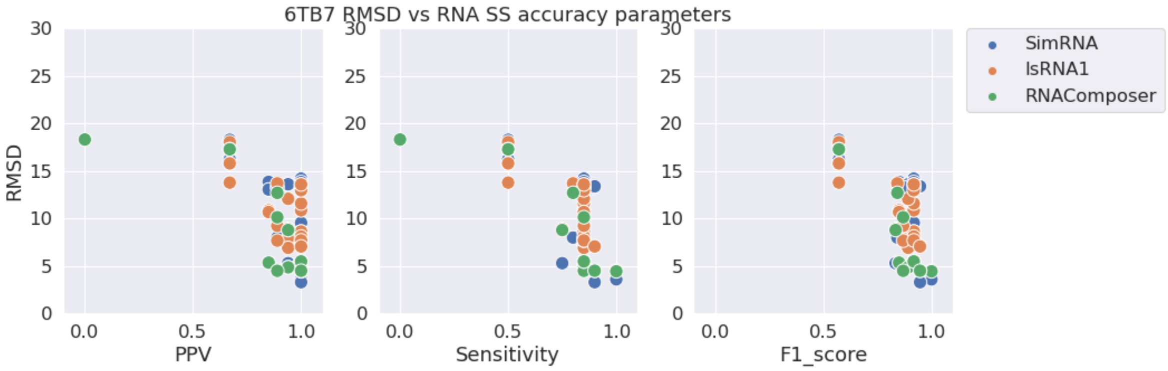


**Figure S25.** RMSD vs. PPV (left), Sensitivity (middle), and F1-score (right) for stem-loop structure PDB 6TB7.

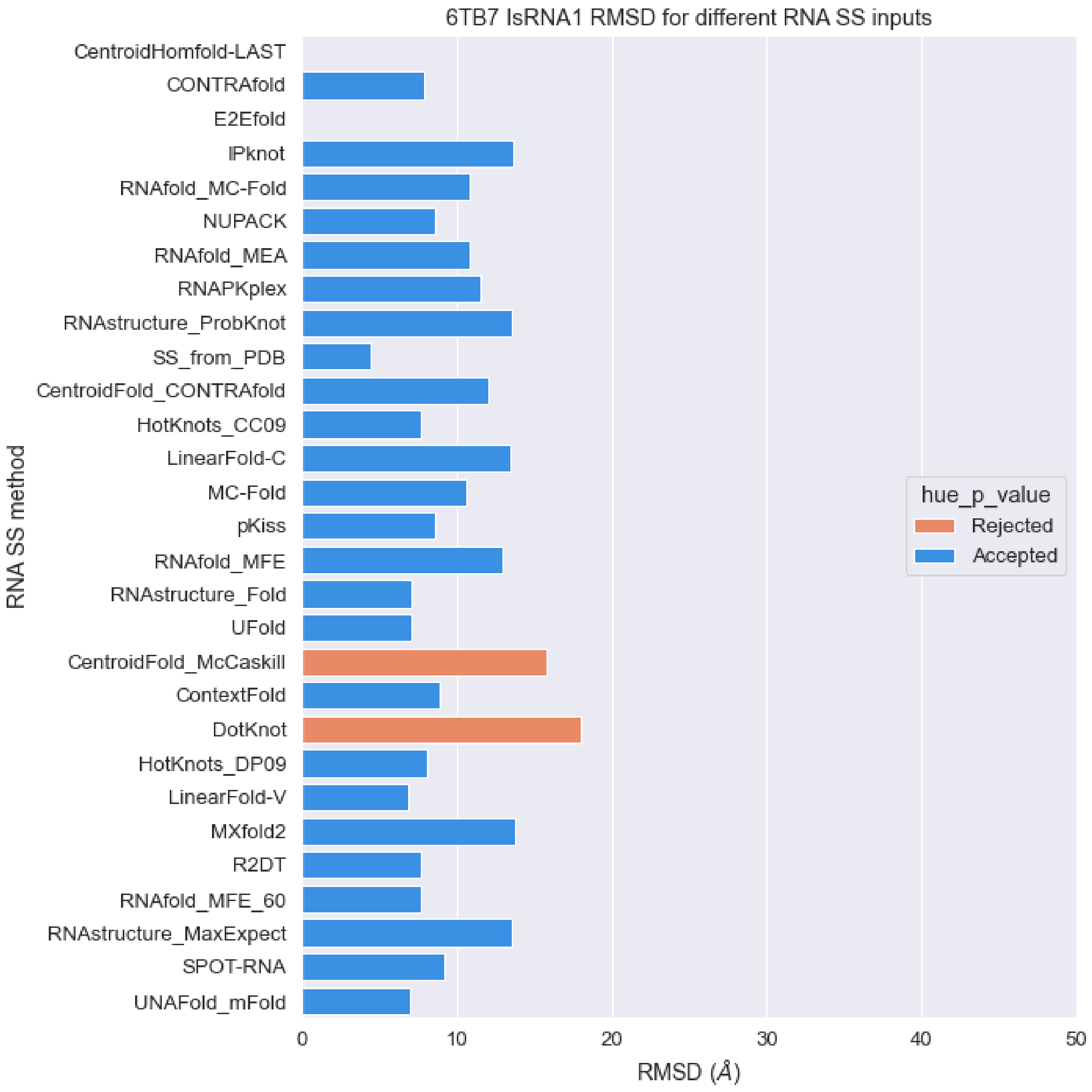


**Figure S26.** RMSD values of predicted structures for PDB 6TB7 sequence with different RNA secondary structures as constraints with IsRNA1 package (orange: p-value > 0.01, blue: p-value < 0.01).

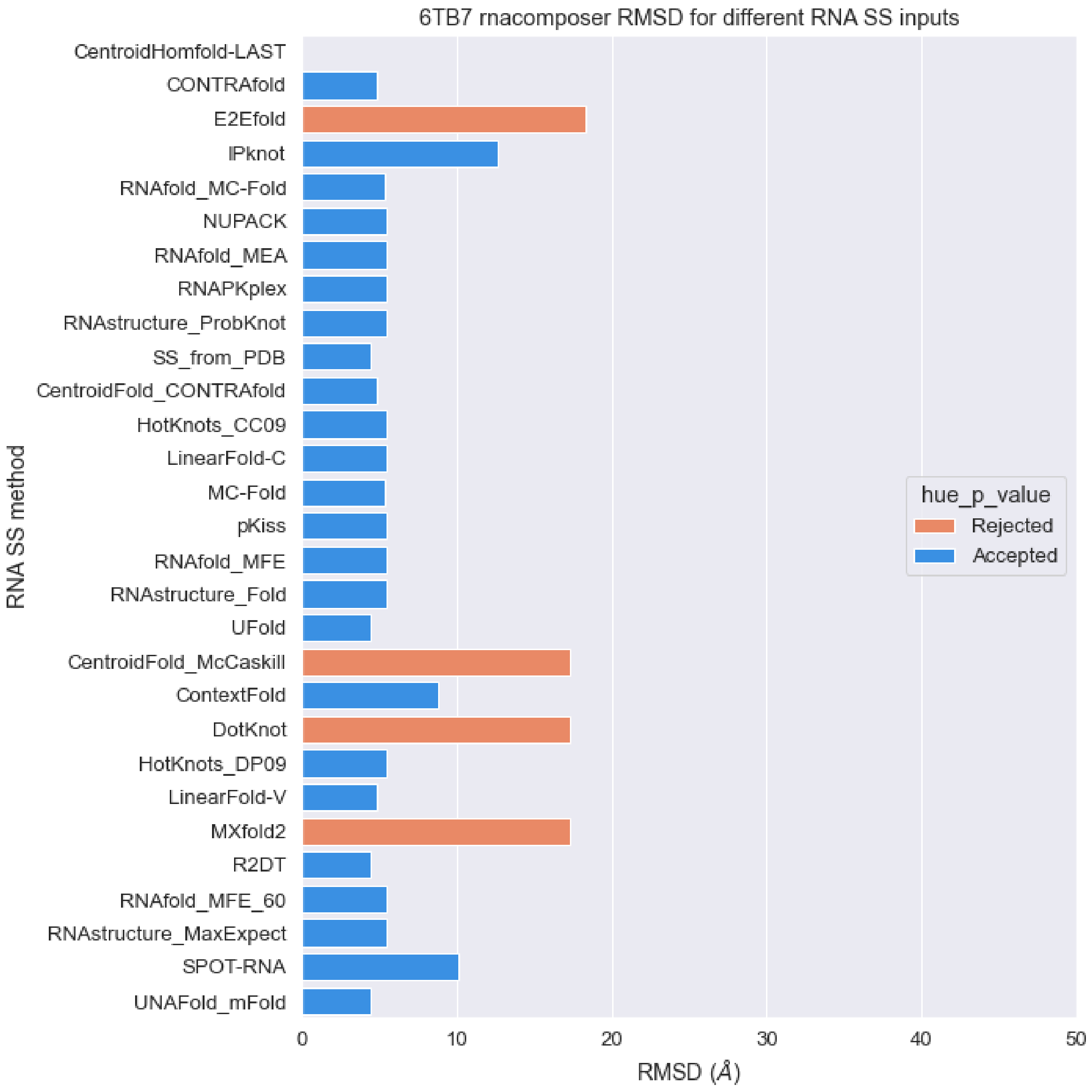


**Figure S27.** RMSD values of predicted structures for PDB 6TB7 sequence with different RNA secondary structures as constraints with RNAcomposer package (orange: p-value > 0.01, blue: p-value < 0.01).

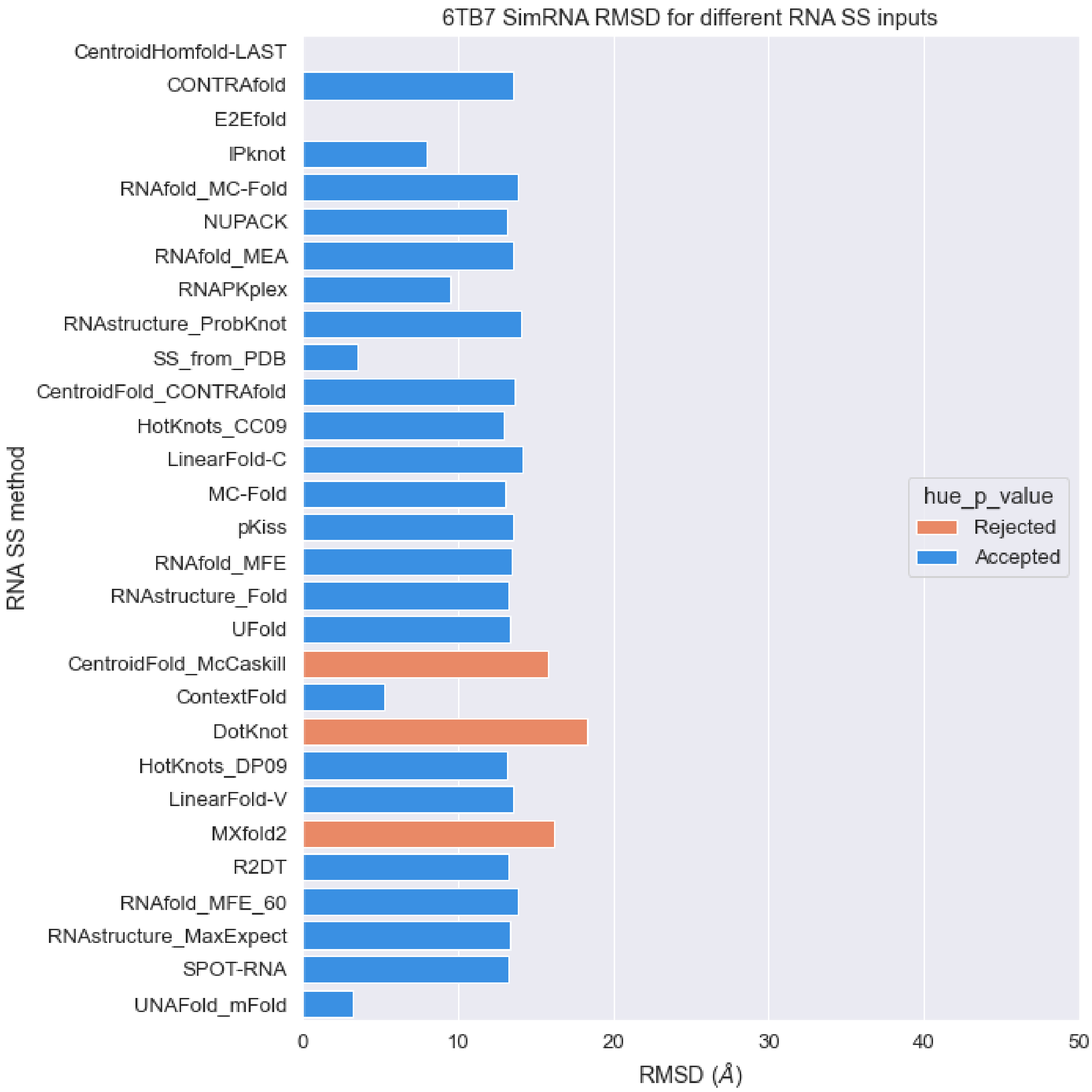


**Figure S28.** RMSD values of predicted structures for PDB 6TB7 sequence with different RNA secondary structures as constraints with SimRNA package (orange: p-value > 0.01, blue: p-value < 0.01).

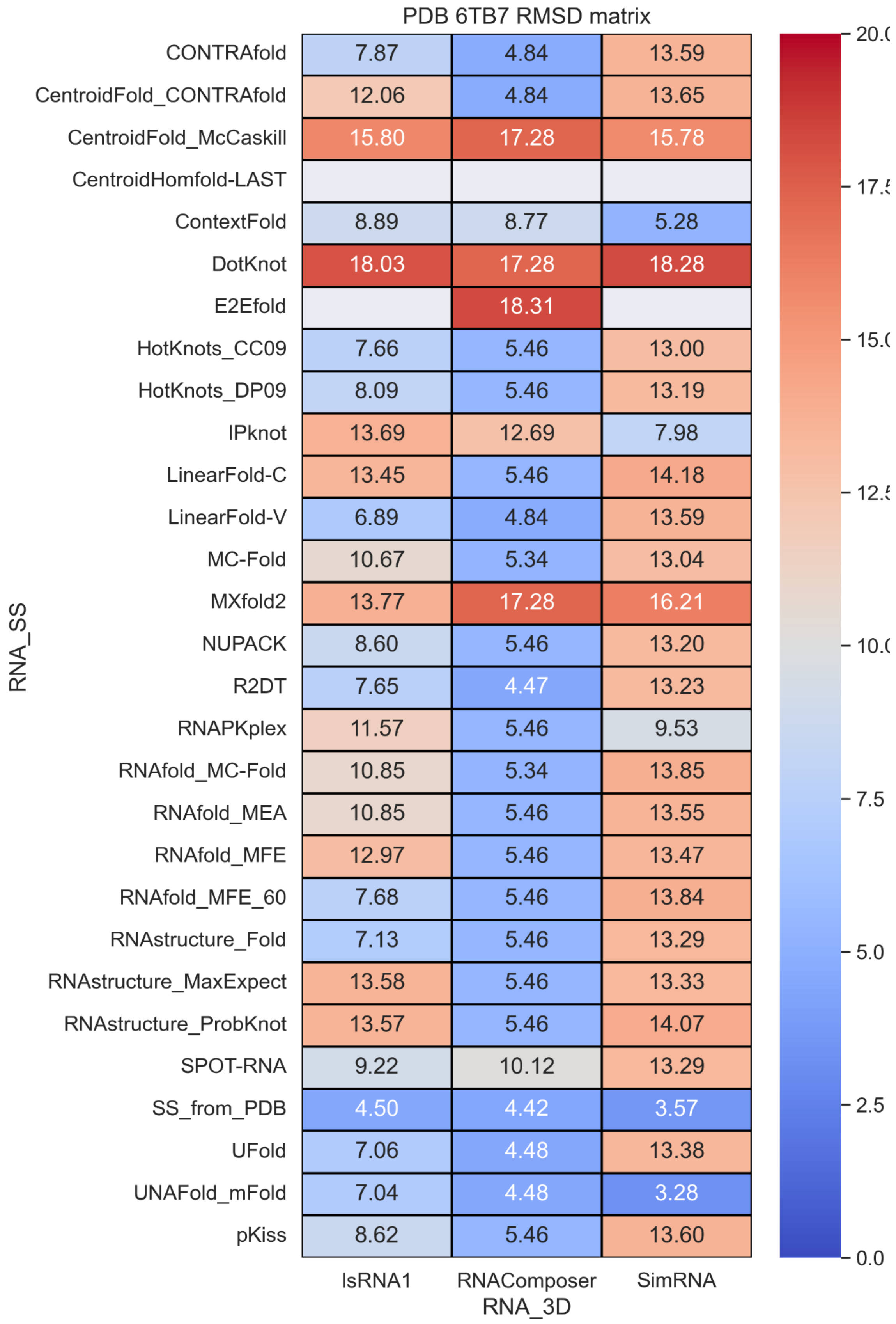


**Figure S29.** RMSD values with different RNA secondary structure methods and RNA 3D methods.

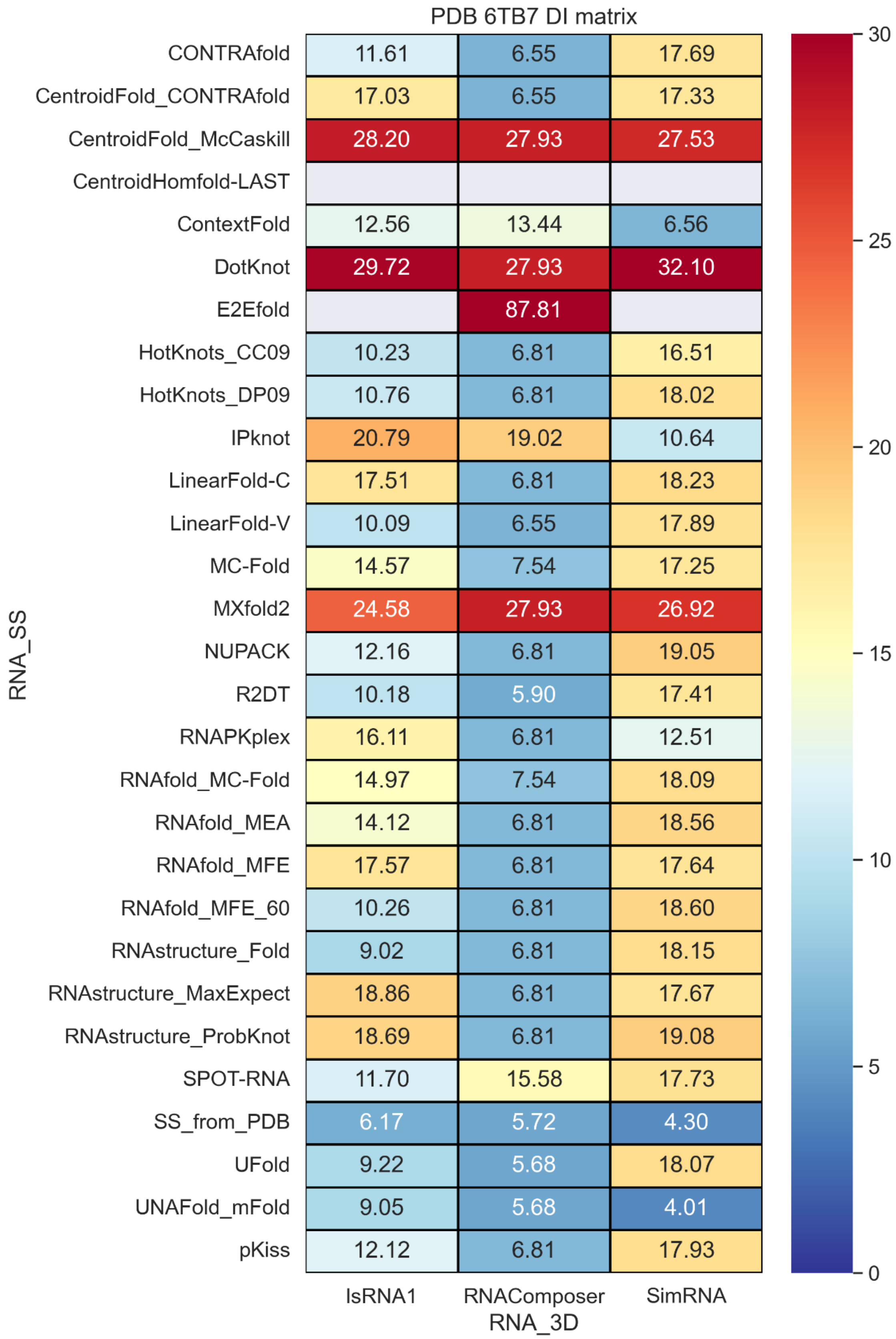


**Figure S30. Deformation Index** values with different RNA secondary structure methods and RNA 3D methods.

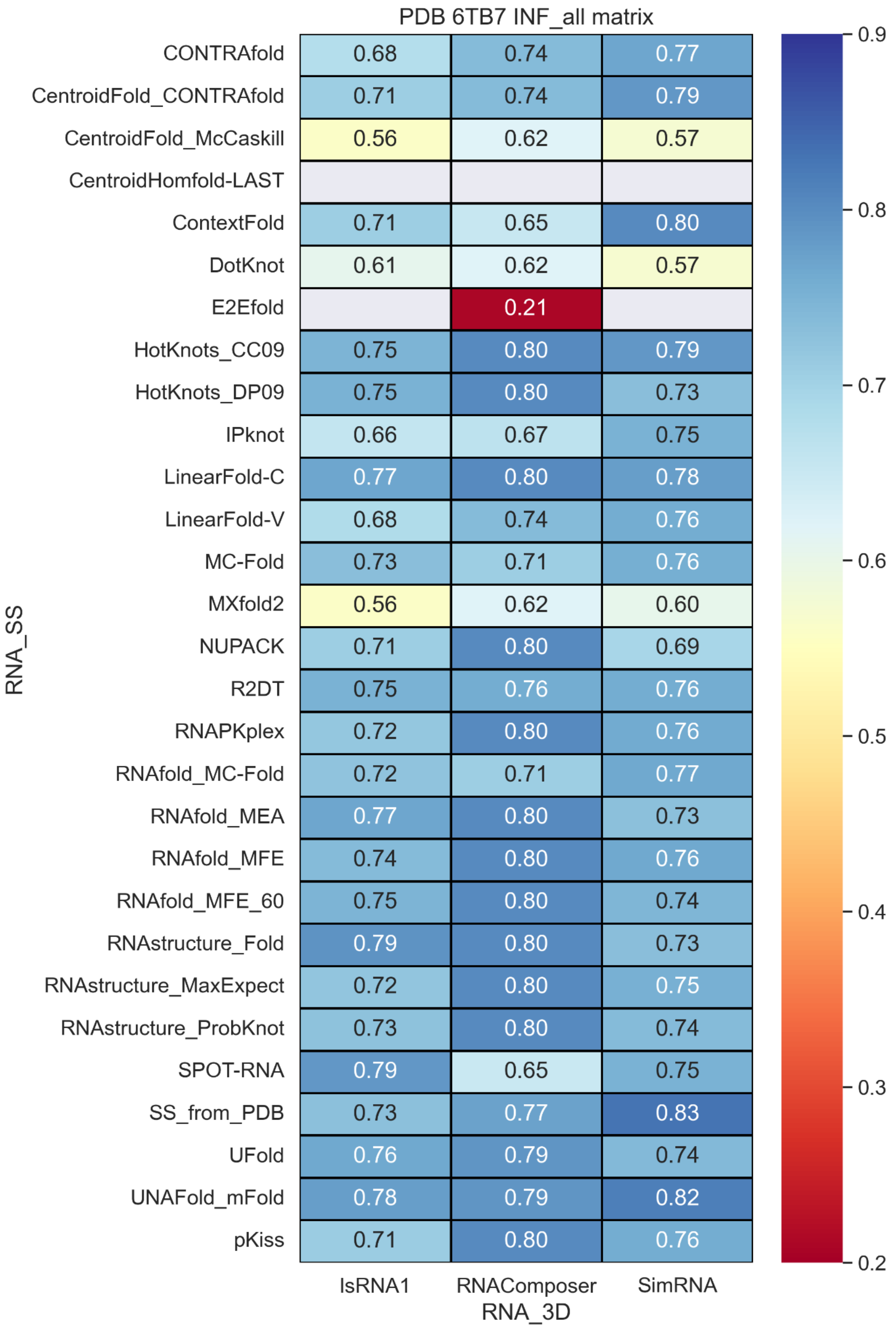


**Figure S31.** INF_all values with different RNA secondary structure methods and RNA 3D methods.

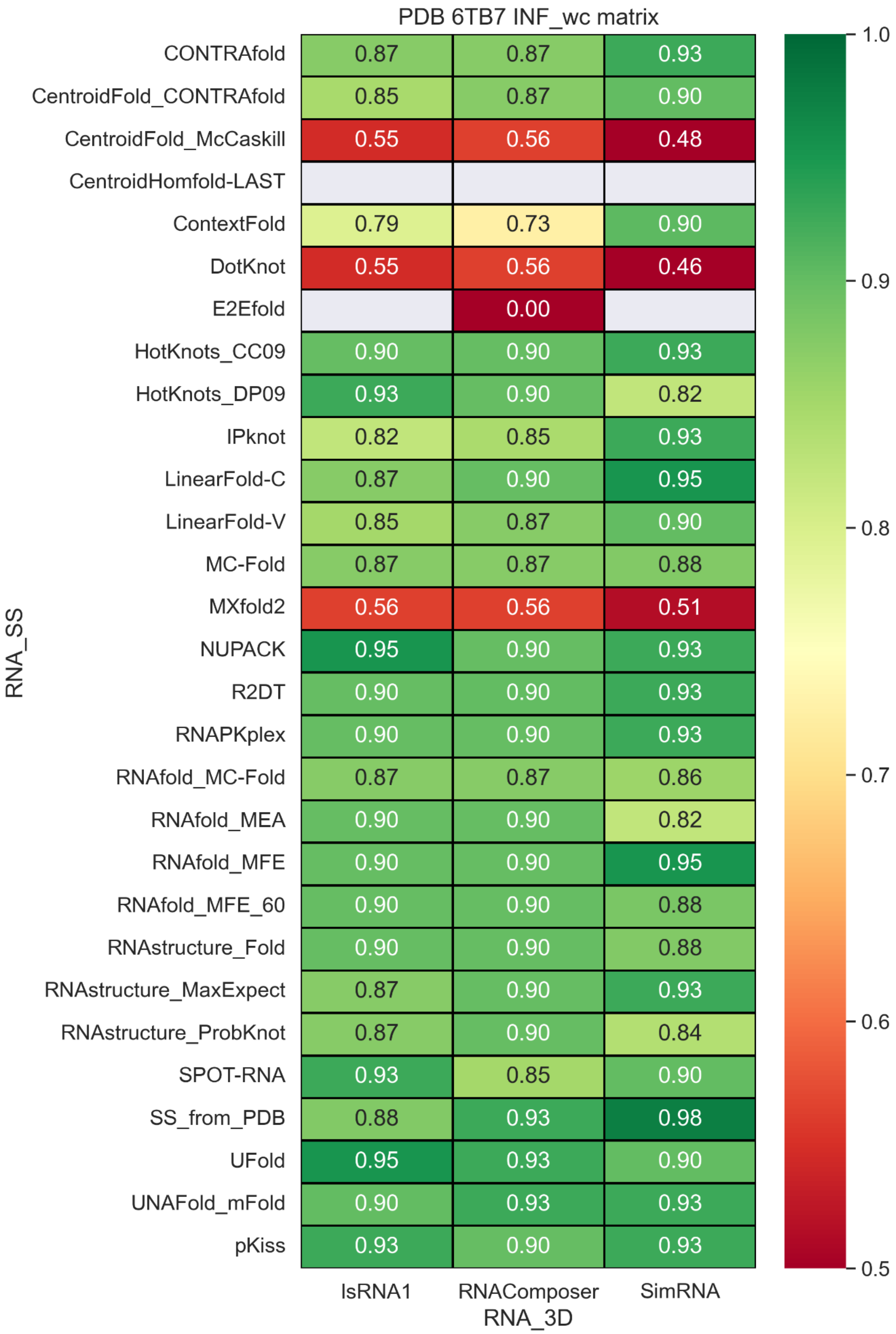


**Figure S32.** INF_wc values with different RNA secondary structure methods and RNA 3D methods.

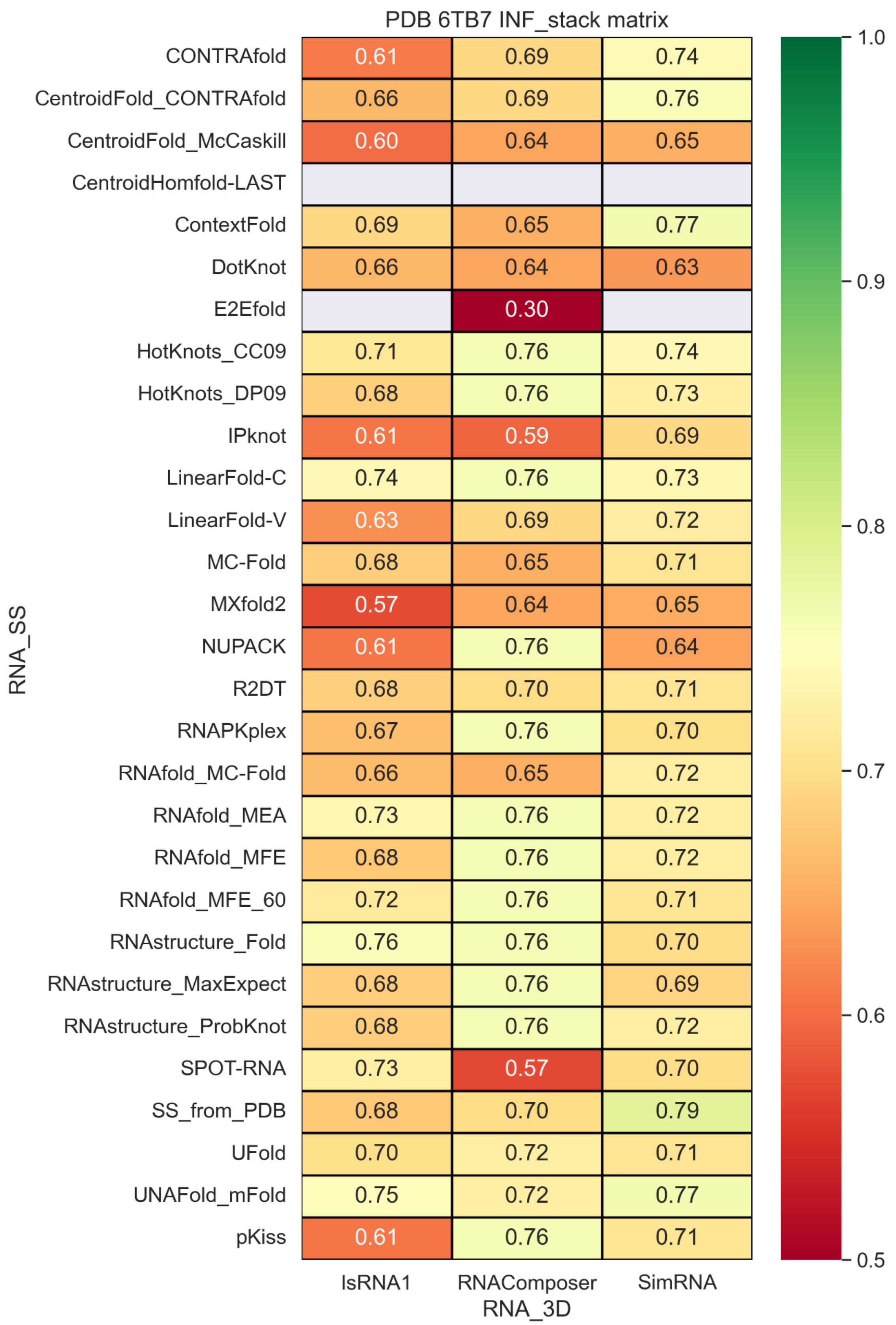


**Figure S33. INF_stack** values with different RNA secondary structure methods and RNA 3D methods.

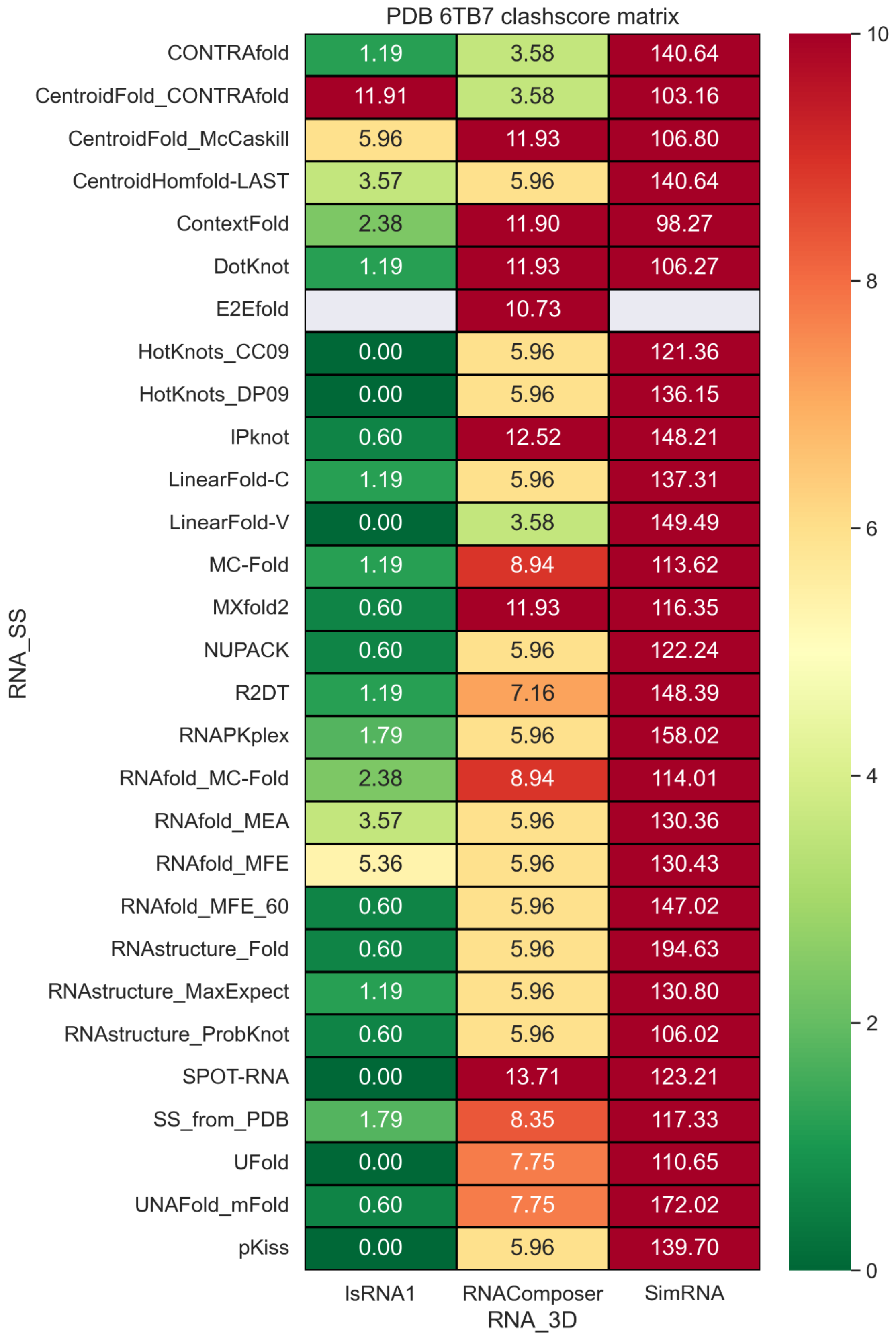


**Figure S34.** clashscore values with different RNA secondary structure methods and RNA 3D methods.

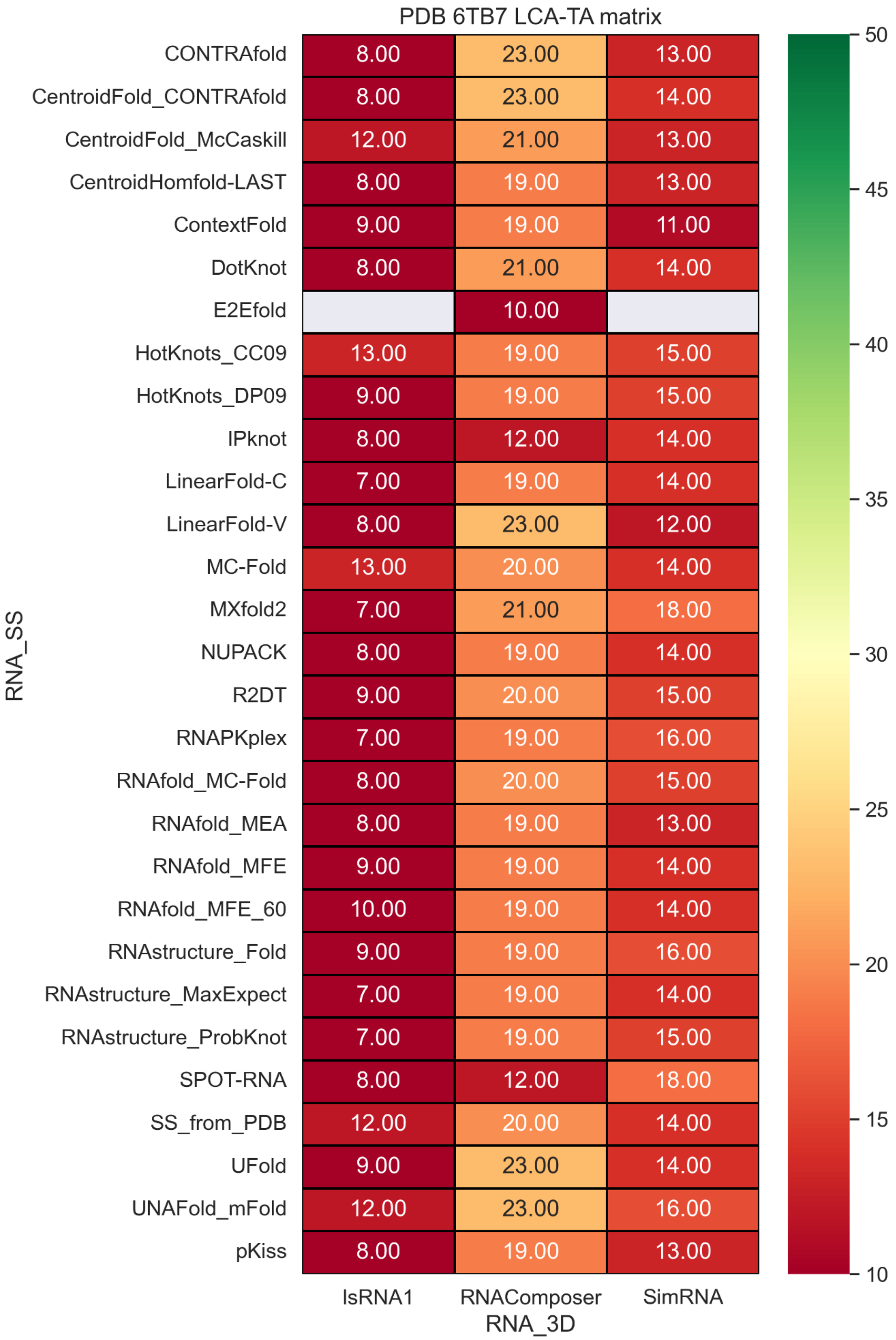


**Figure S35.** LCS-TA values with different RNA secondary structure methods and RNA 3D methods.

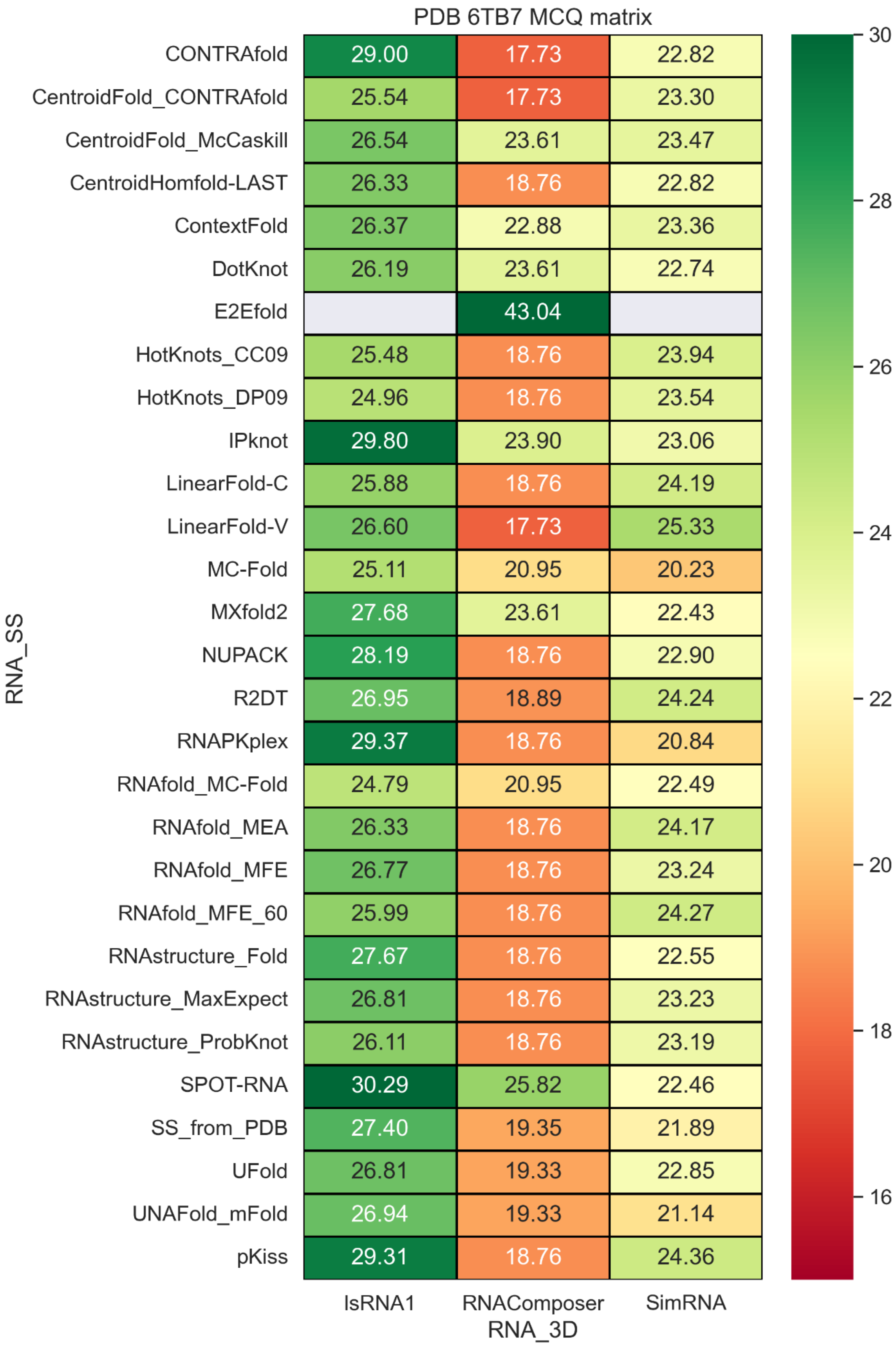


| RNA_SS | IsRNA1 | RNAComposer | SimRNA |
|---|---|---|---|
| CONTRAfold | 29.00 | 17.73 | 22.82 |
| CentroidFold_CONTRAfold | 25.54 | 17.73 | 23.30 |
| CentroidFold_McCaskill | 26.54 | 23.61 | 23.47 |
| CentroidHomfold-LAST | 26.33 | 18.76 | 22.82 |
| ContextFold | 26.37 | 22.88 | 23.36 |
| DotKnot | 26.19 | 23.61 | 22.74 |
| E2Efold | | 43.04 | |
| HotKnots_CC09 | 25.48 | 18.76 | 23.94 |
| HotKnots_DP09 | 24.96 | 18.76 | 23.54 |
| IPknot | 29.80 | 23.90 | 23.06 |
| LinearFold-C | 25.88 | 18.76 | 24.19 |
| LinearFold-V | 26.60 | 17.73 | 25.33 |
| MC-Fold | 25.11 | 20.95 | 20.23 |
| MXfold2 | 27.68 | 23.61 | 22.43 |
| NUPACK | 28.19 | 18.76 | 22.90 |
| R2DT | 26.95 | 18.89 | 24.24 |
| RNAPKplex | 29.37 | 18.76 | 20.84 |
| RNAfold_MC-Fold | 24.79 | 20.95 | 22.49 |
| RNAfold_MEA | 26.33 | 18.76 | 24.17 |
| RNAfold_MFE | 26.77 | 18.76 | 23.24 |
| RNAfold_MFE_60 | 25.99 | 18.76 | 24.27 |
| RNAstructure_Fold | 27.67 | 18.76 | 22.55 |
| RNAstructure_MaxExpect | 26.81 | 18.76 | 23.23 |
| RNAstructure_ProbKnot | 26.11 | 18.76 | 23.19 |
| SPOT-RNA | 30.29 | 25.82 | 22.46 |
| SS_from_PDB | 27.40 | 19.35 | 21.89 |
| UFold | 26.81 | 19.33 | 22.85 |
| UNAFold_mFold | 26.94 | 19.33 | 21.14 |
| pKiss | 29.31 | 18.76 | 24.36 |

**Figure S36.** MCQ values with different RNA secondary structure methods and RNA 3D methods.

### PDB 7LYJ

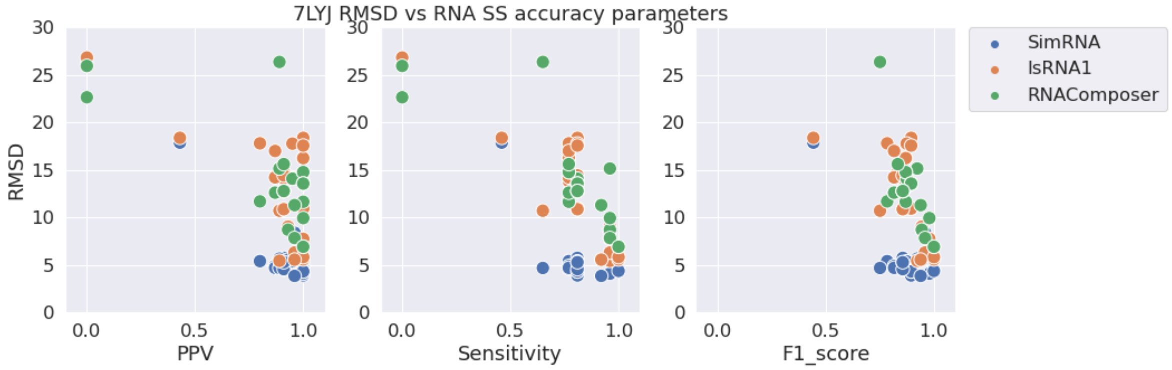


**Figure S37.** RMSD vs. PPV (left), Sensitivity (middle), and F1-score (right) for stem-loop structure PDB 7LYJ.

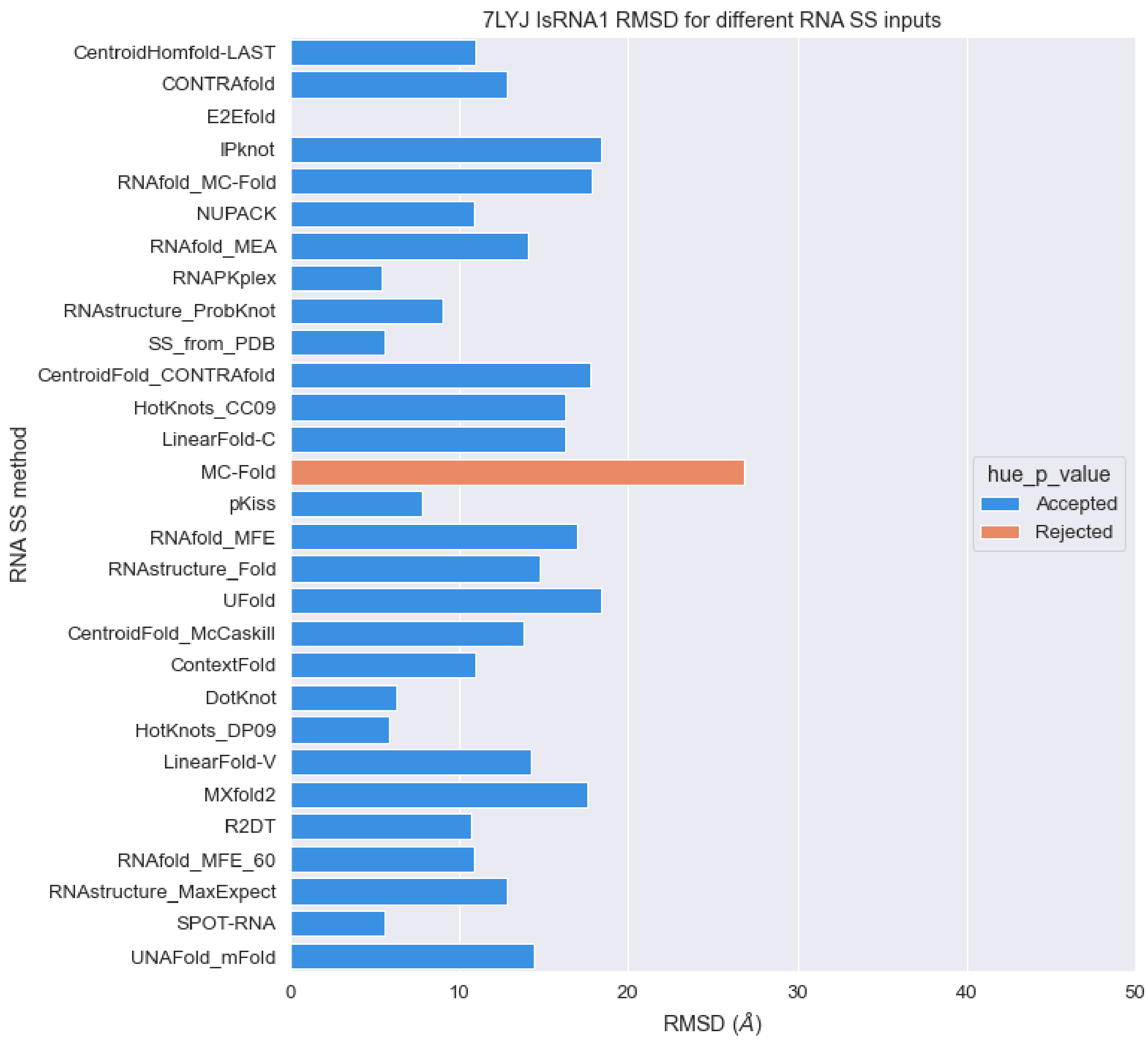


**Figure S38.** RMSD values of predicted structures for PDB 7LYJ sequence with different RNA secondary structures as constraints with IsRNA1 package (orange: p-value > 0.01, blue: p-value < 0.01).

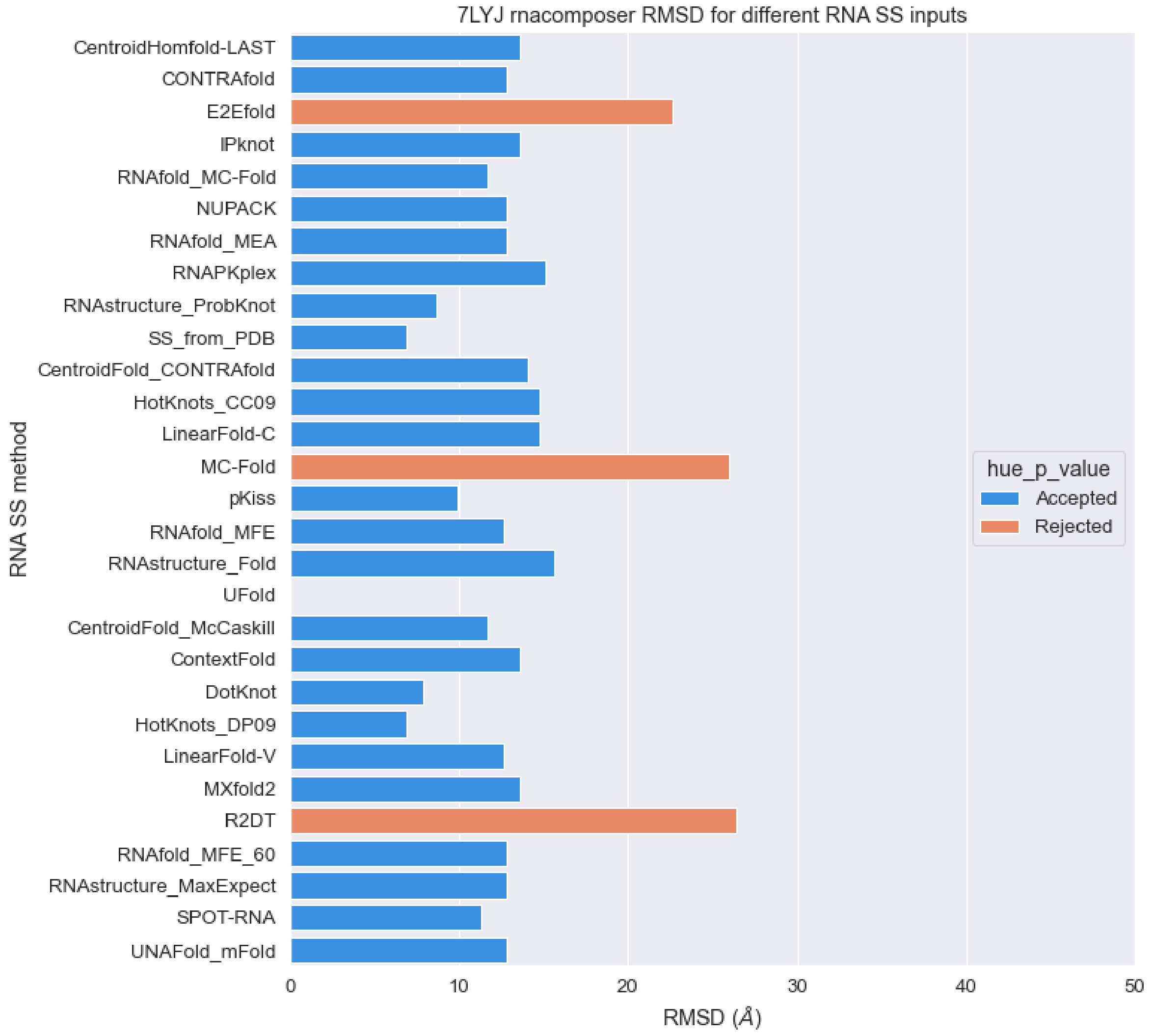


**Figure S39.** RMSD values of predicted structures for PDB 7LYJ sequence with different RNA secondary structures as constraints with RNAcomposer package (orange: p-value > 0.01, blue: p-value < 0.01).

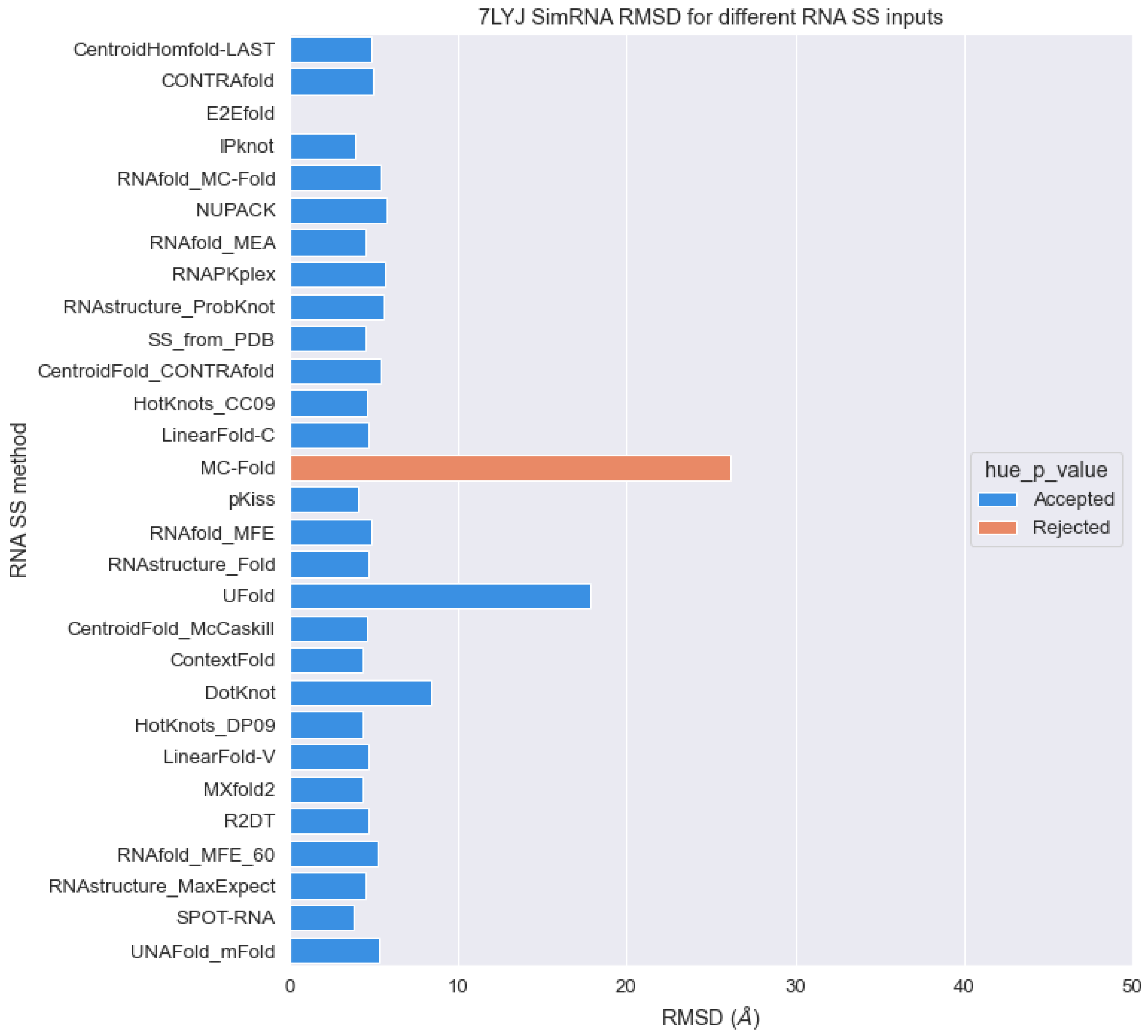


**Figure S40.** RMSD values of predicted structures for PDB 7LYJ sequence with different RNA secondary structures as constraints with SimRNA package (orange: p-value > 0.01, blue: p-value < 0.01).

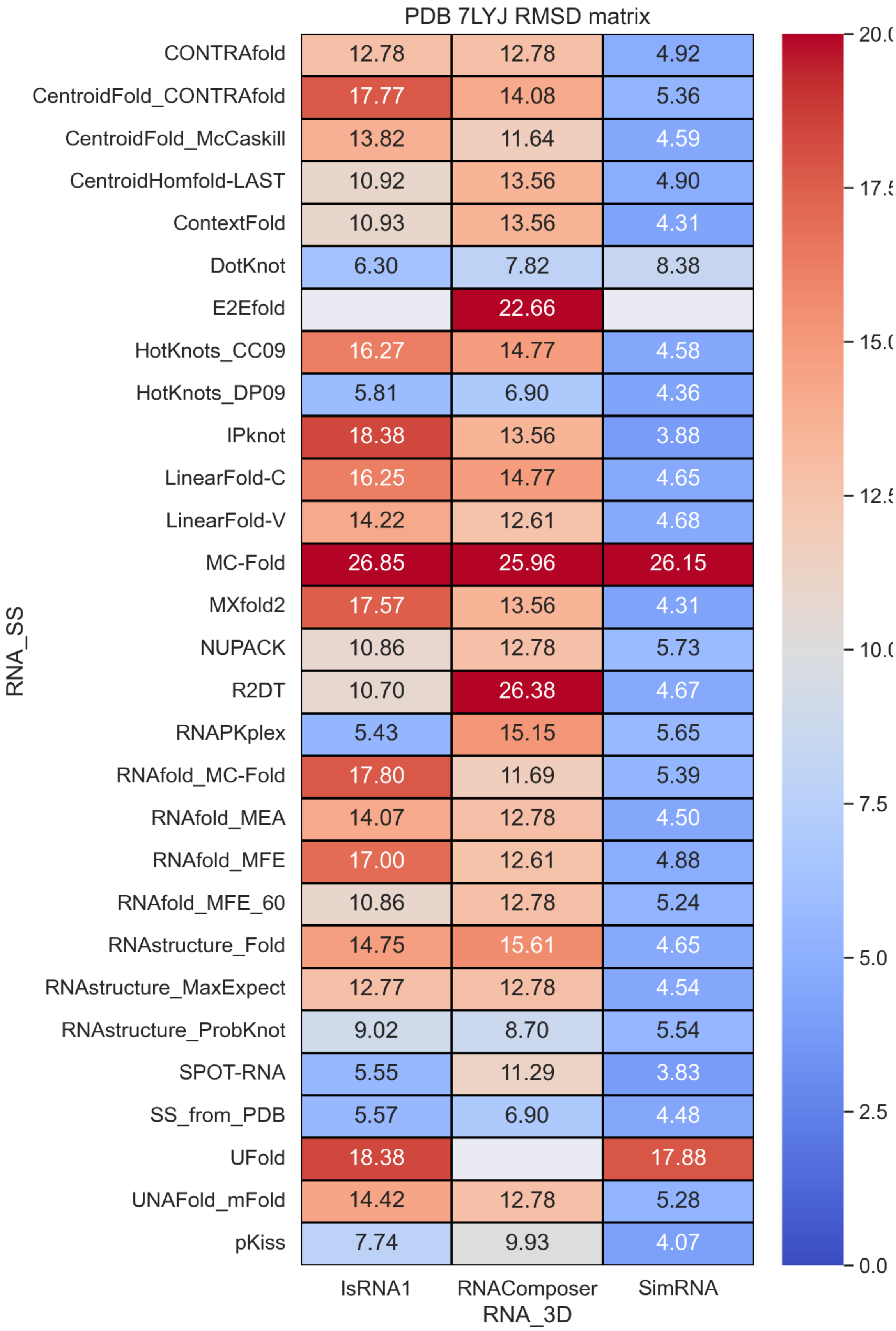


**Figure S41.** RMSD values with different RNA secondary structure methods and RNA 3D methods.

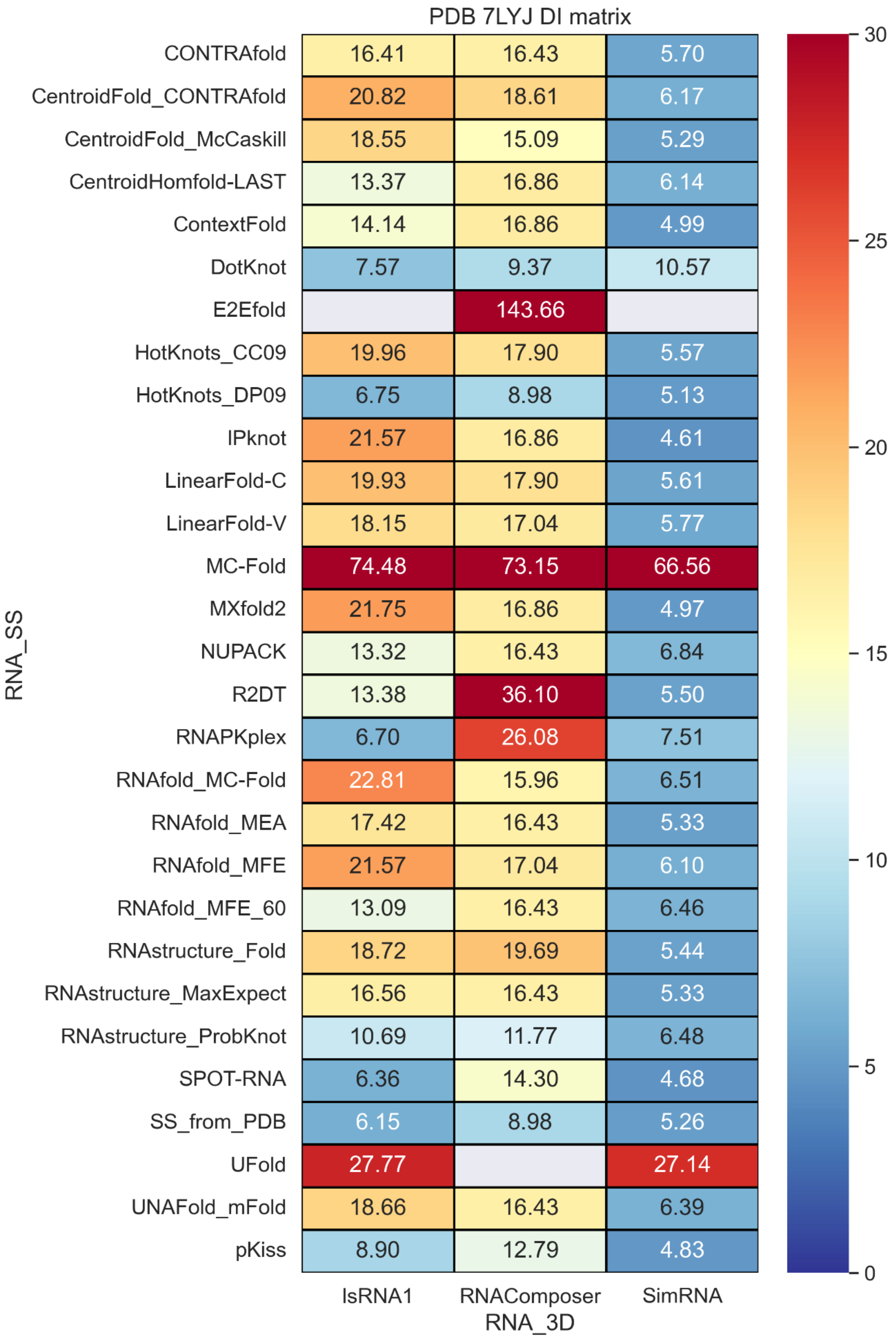


**Figure S42.** Deformation Index values with different RNA secondary structure methods and RNA 3D methods.

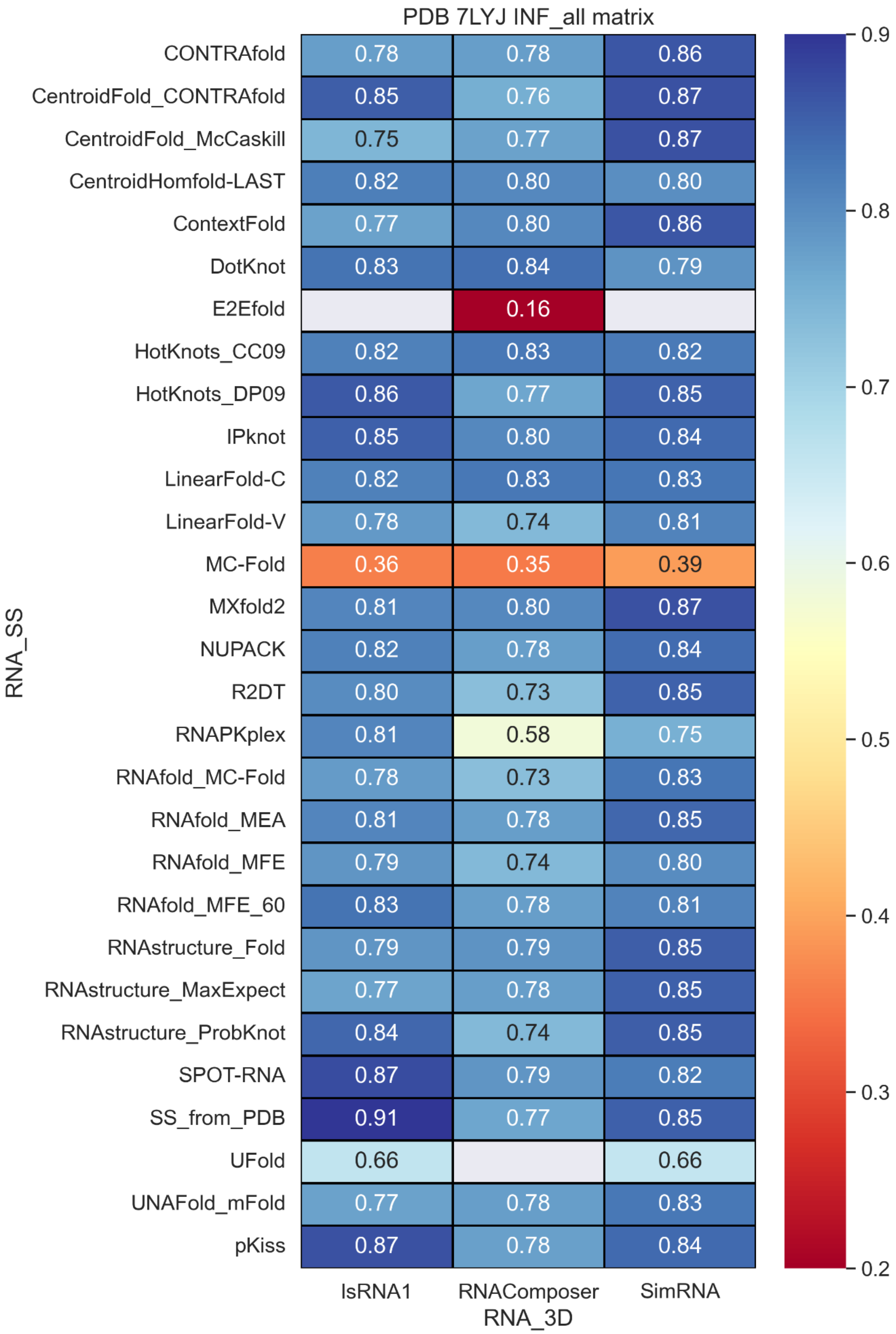


**Figure S43.** INF_all values with different RNA secondary structure methods and RNA 3D methods.

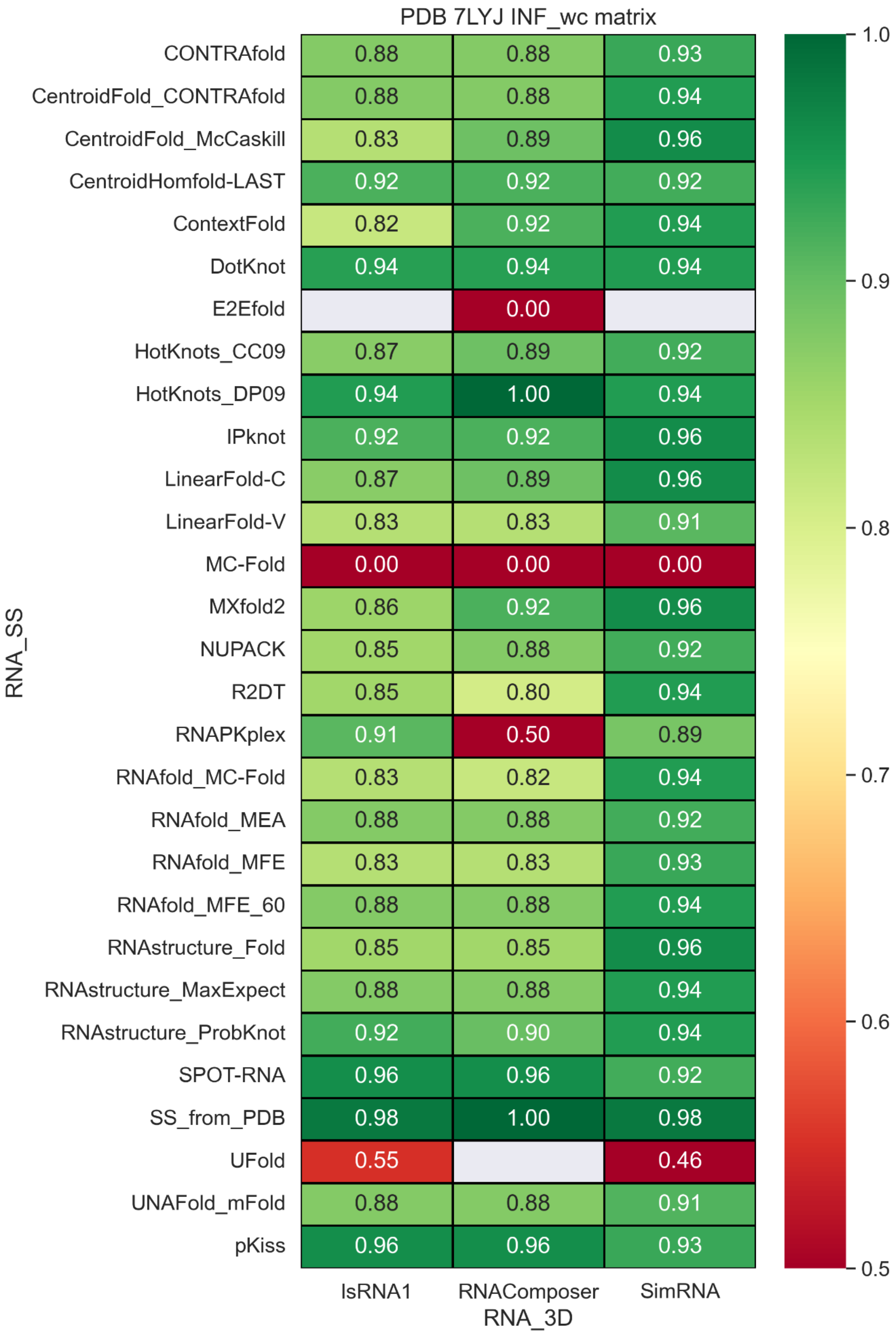


**Figure S44.** INF_wc values with different RNA secondary structure methods and RNA 3D methods.

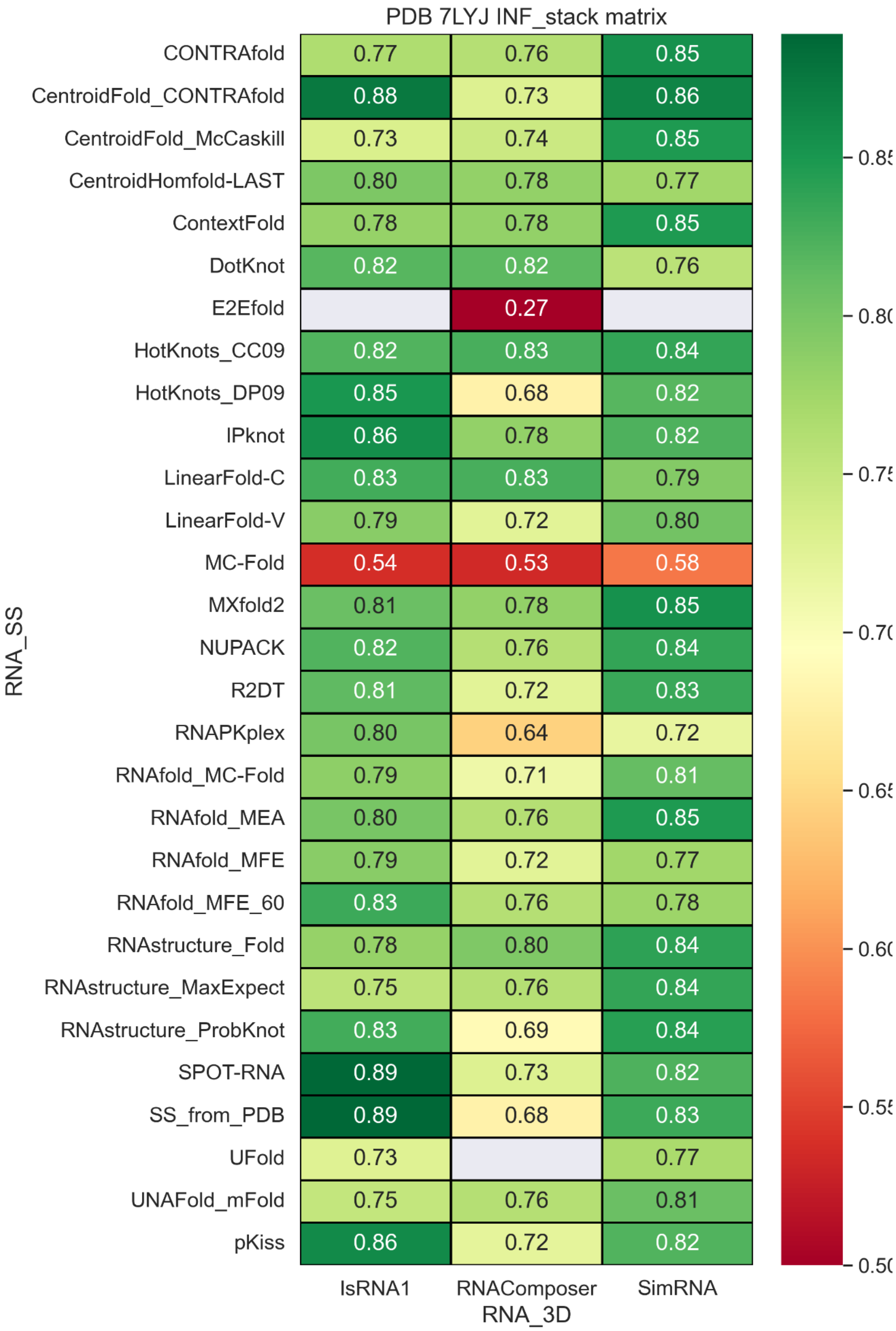


**Figure S45.** INF_stack values with different RNA secondary structure methods and RNA 3D methods.

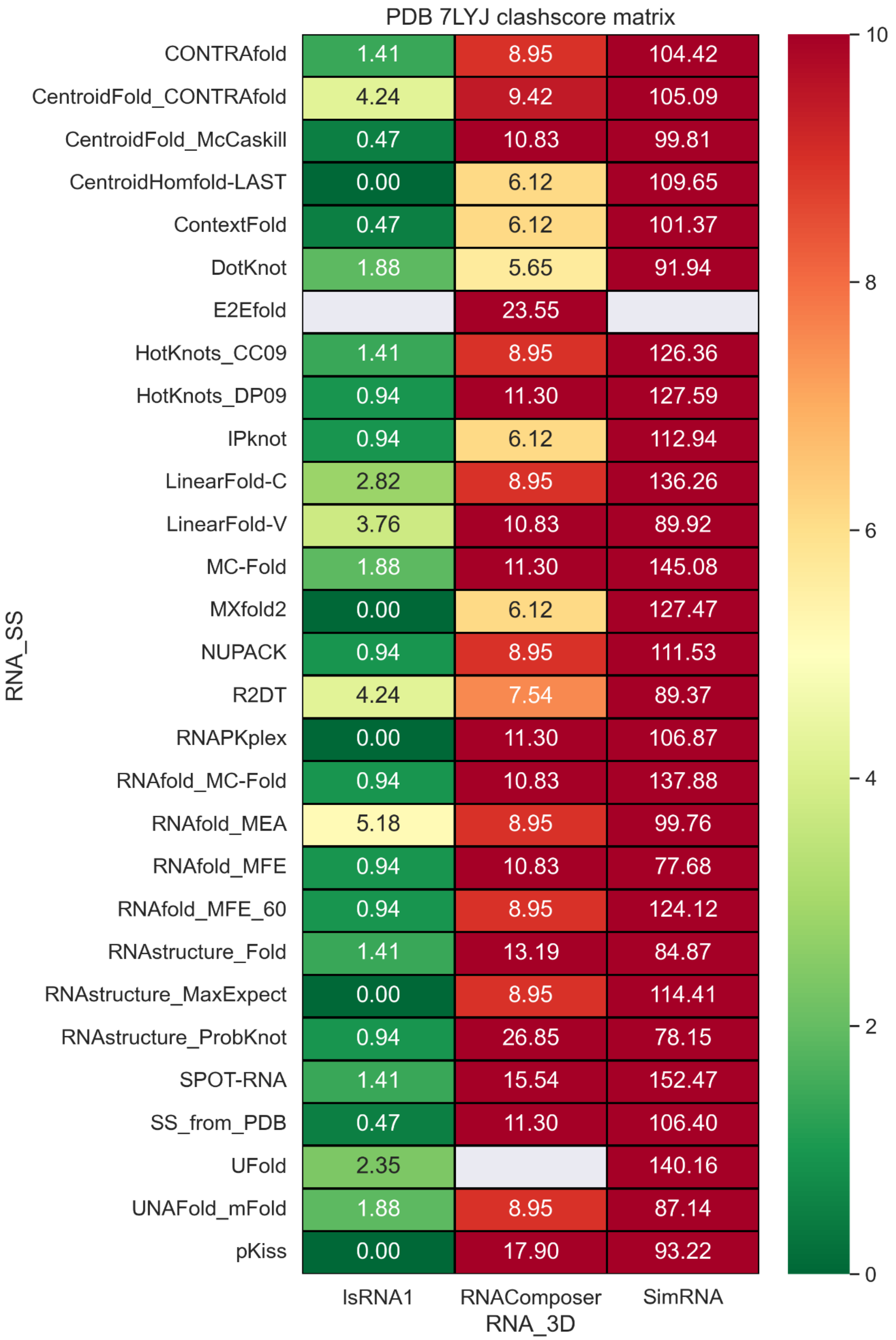


**Figure S46.** Classcore values with different RNA secondary structure methods and RNA 3D methods.

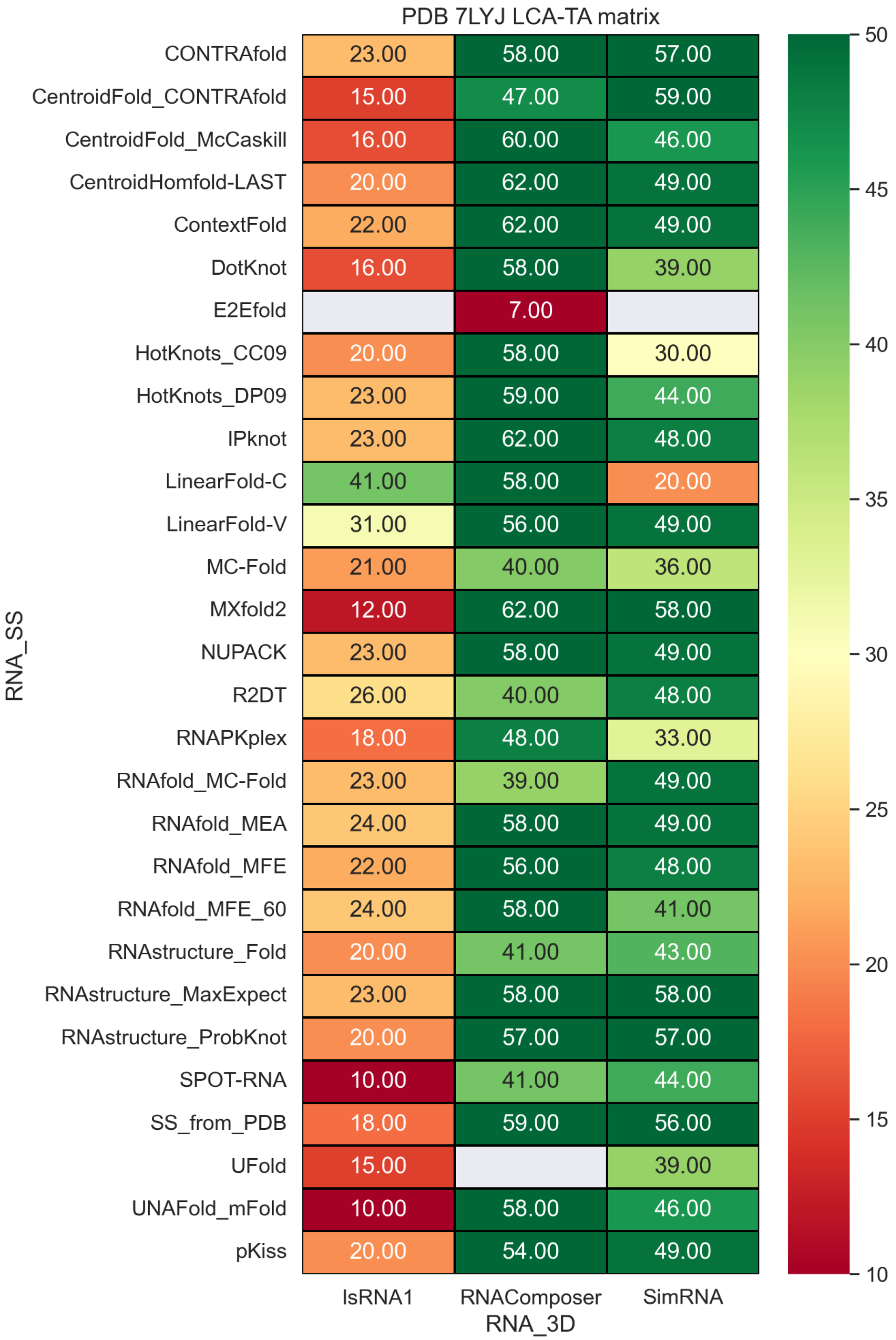


**Figure S47.** LCS-TA values with different RNA secondary structure methods and RNA 3D methods.

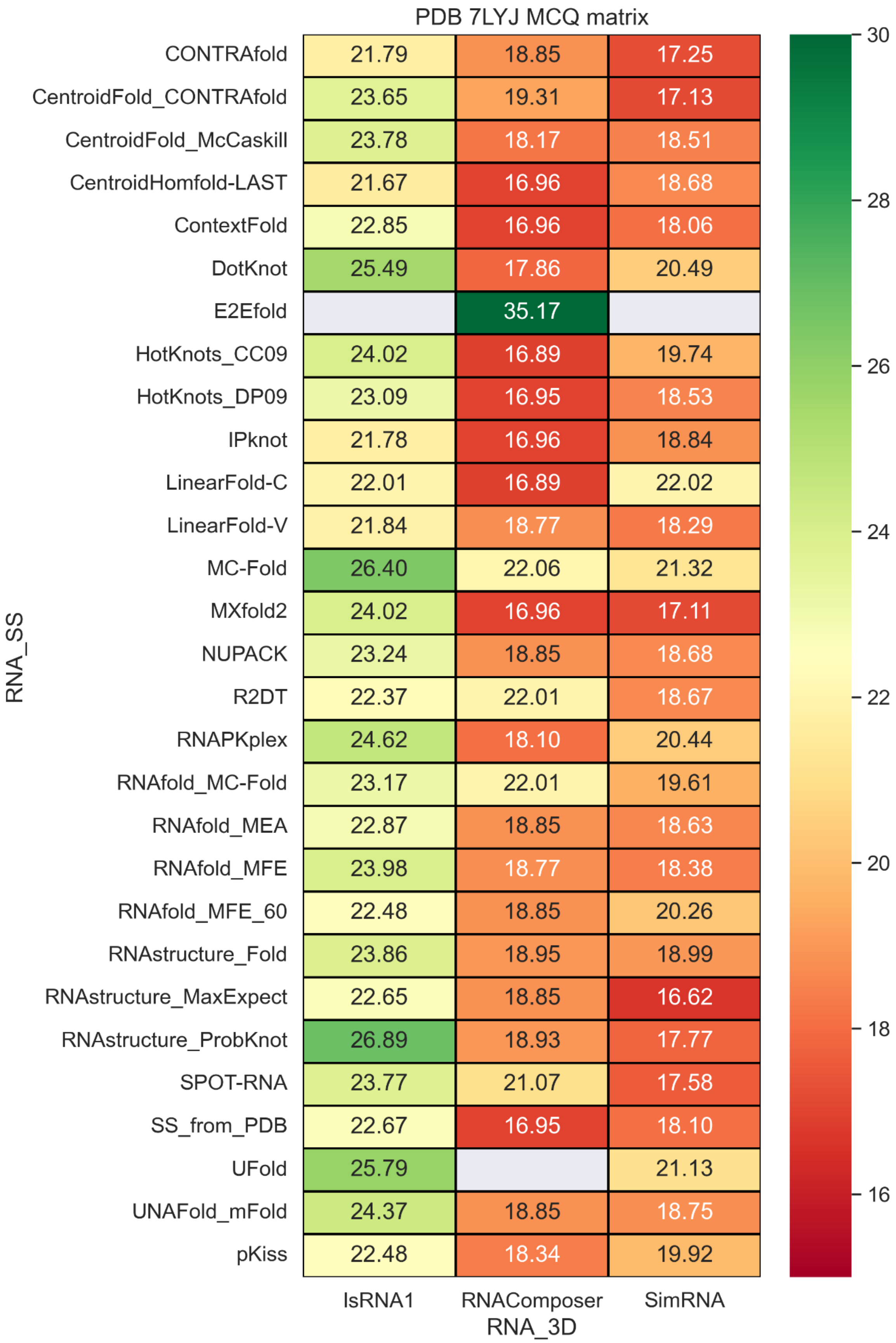


**Figure S48.** MCQ values with different RNA secondary structure methods and RNA 3D methods.

### PDB 5NWQ

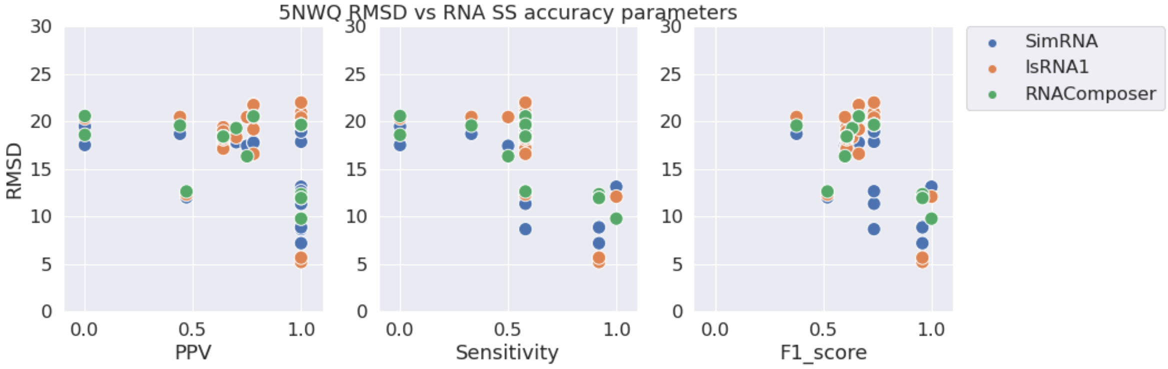


**Figure S49.** RMSD vs. PPV (left), Sensitivity (middle), and F1-score (right) for stem-loop structure PDB 5NWQ.

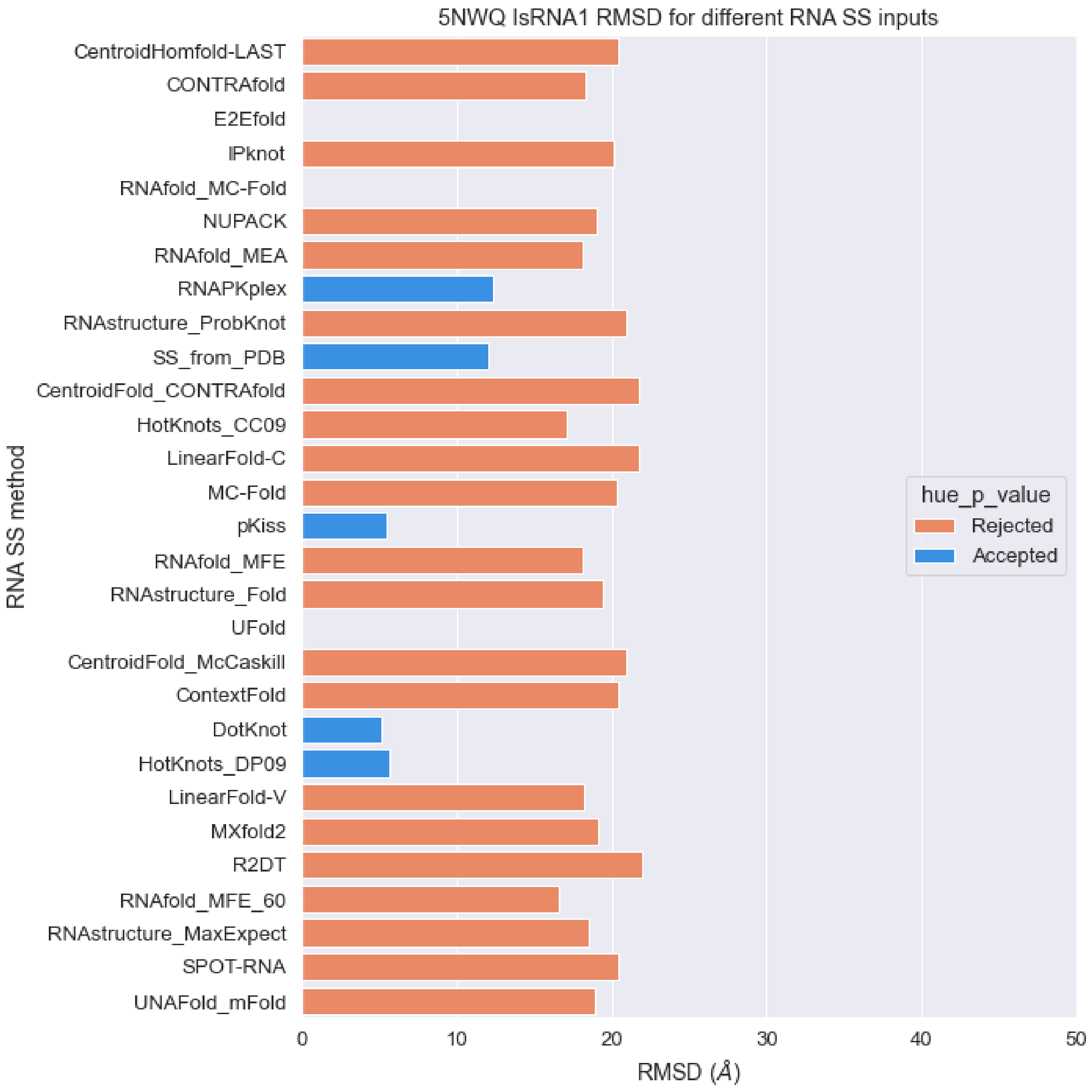


**Figure S50.** RMSD values of predicted structures for PDB 5NWQ sequence with different RNA secondary structures as constraints with IsRNA1 package (orange: p-value > 0.01, blue: p-value < 0.01).

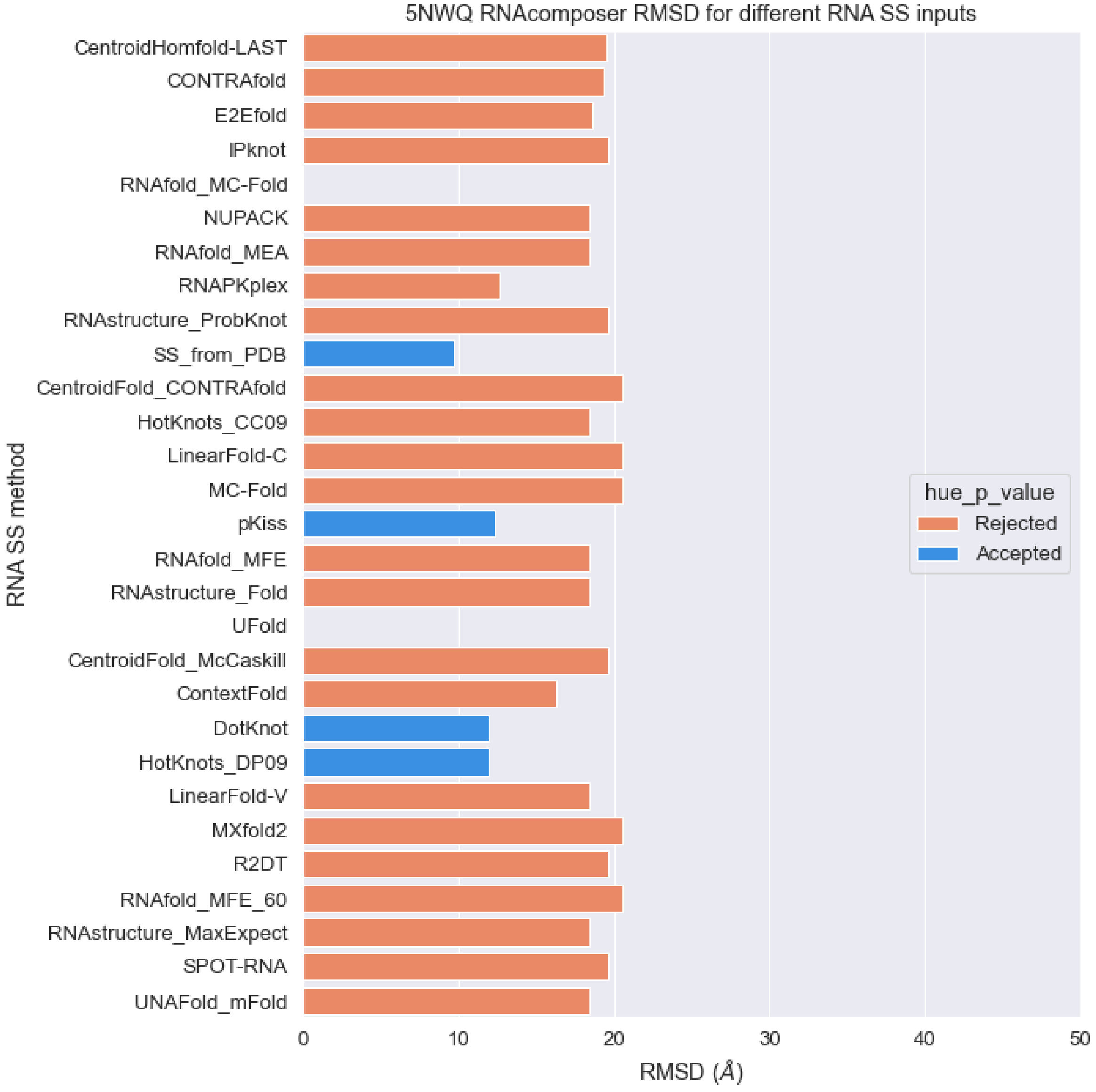


**Figure S51.** RMSD values of predicted structures for PDB 5NWQ sequence with different RNA secondary structures as constraints with RNAcomposer package (orange: p-value > 0.01, blue: p-value < 0.01).

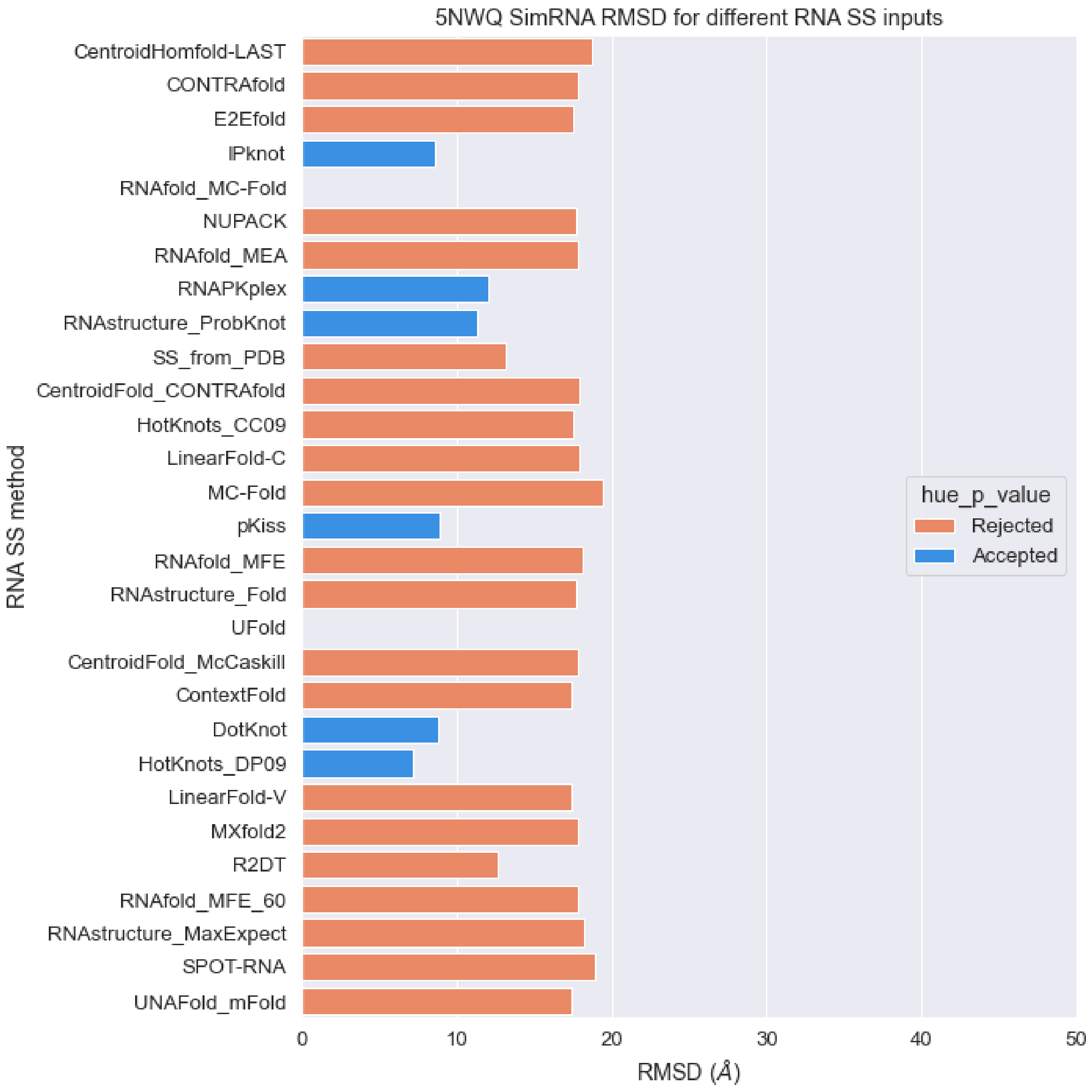


**Figure S52.** RMSD values of predicted structures for PDB 5NWQ sequence with different RNA secondary structures as constraints with SimRNA package (orange: p-value > 0.01, blue: p-value < 0.01).

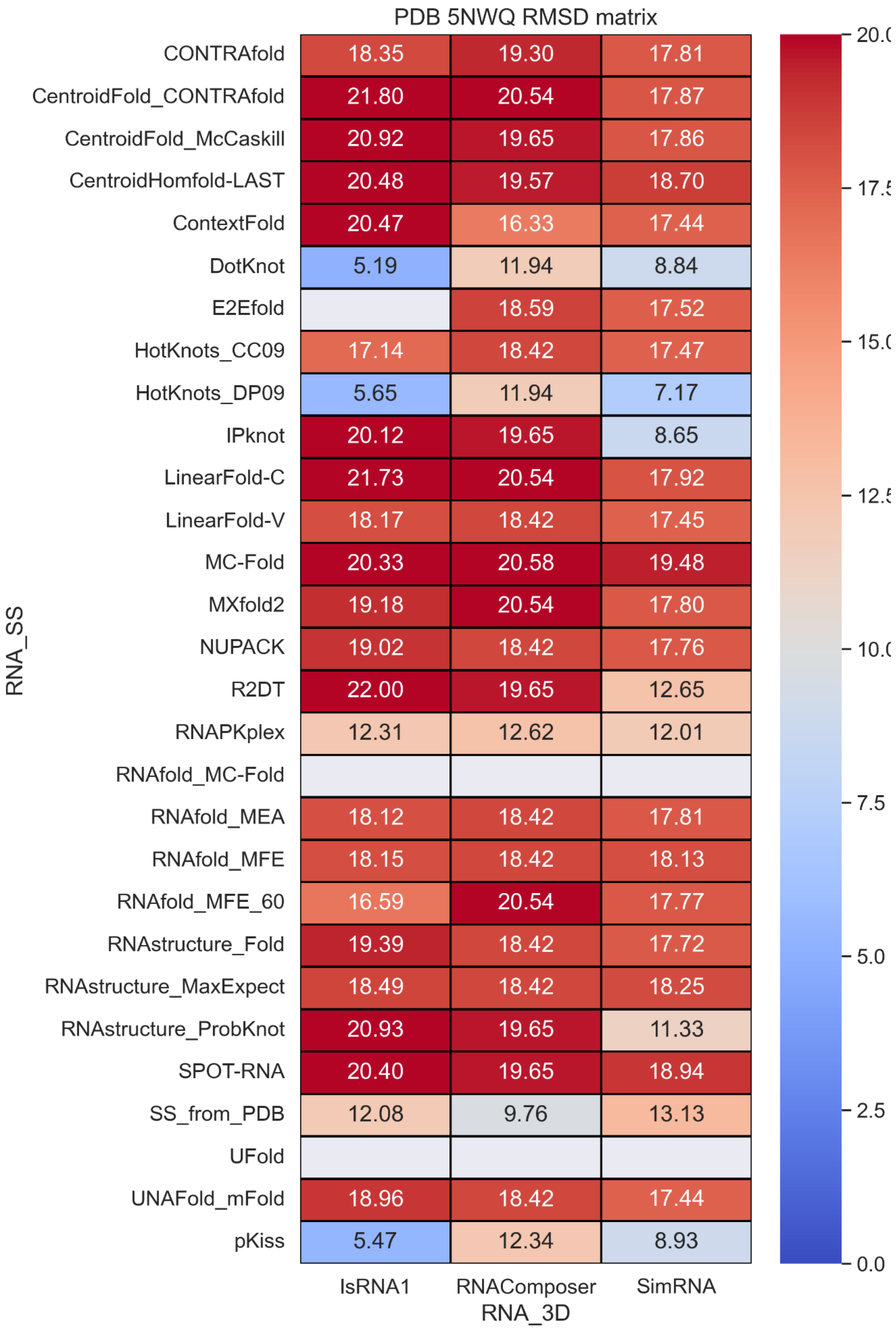


**Figure S53.** RMSD values with different RNA secondary structure methods and RNA 3D methods.

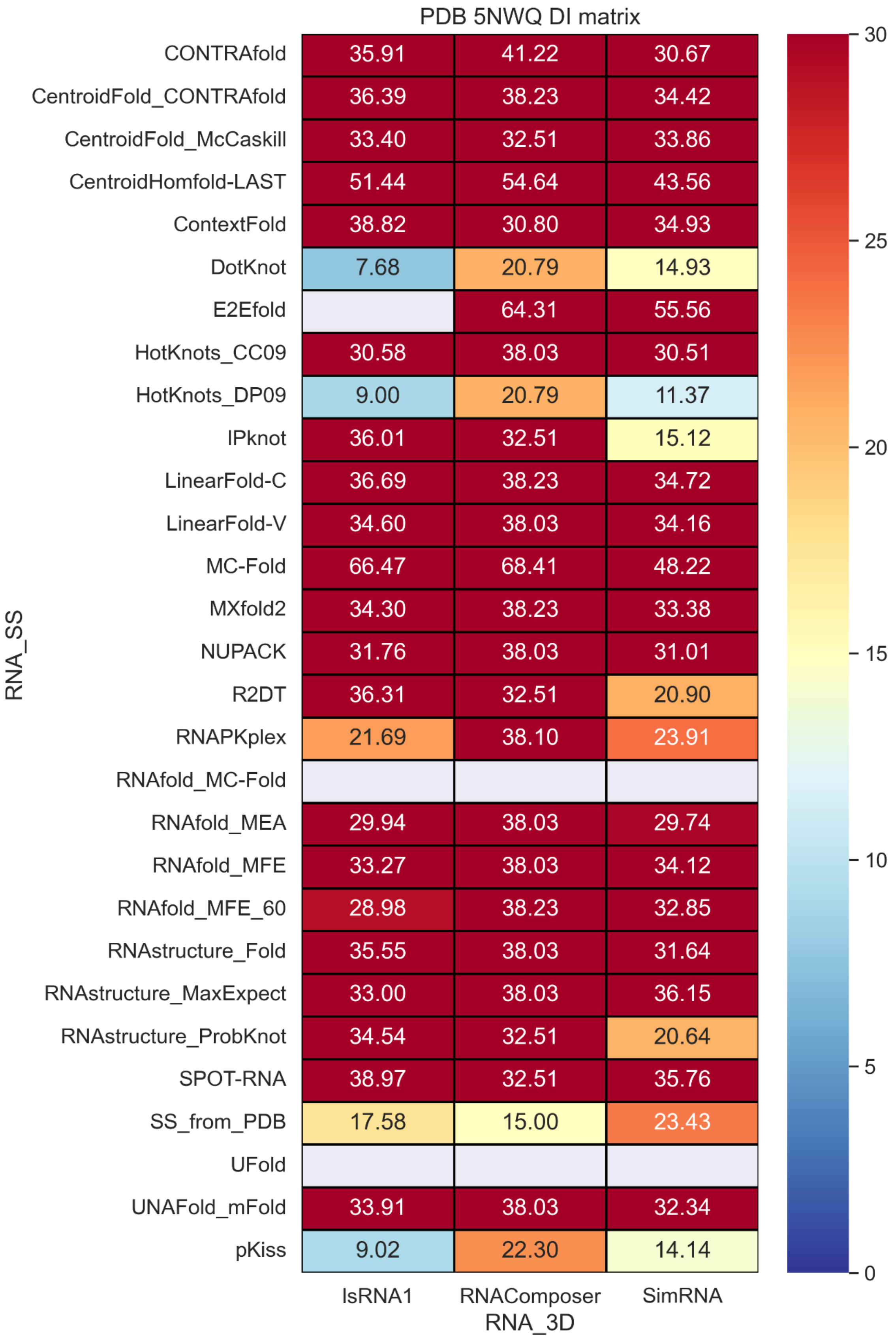


**Figure S54.** Deformation index values with different RNA secondary structure methods and RNA 3D methods.

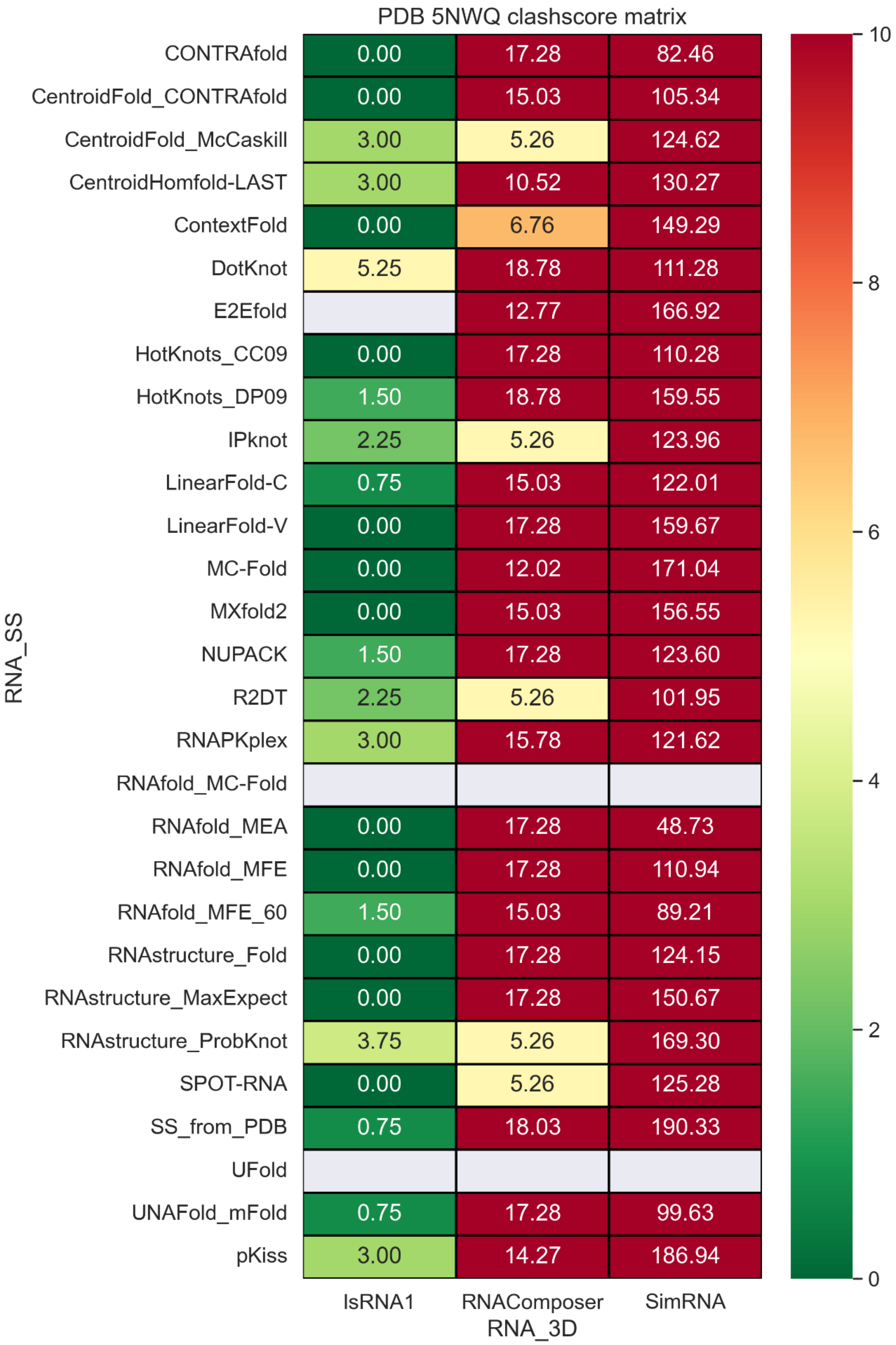


**Figure S55.** Clashscore values with different RNA secondary structure methods and RNA 3D methods.

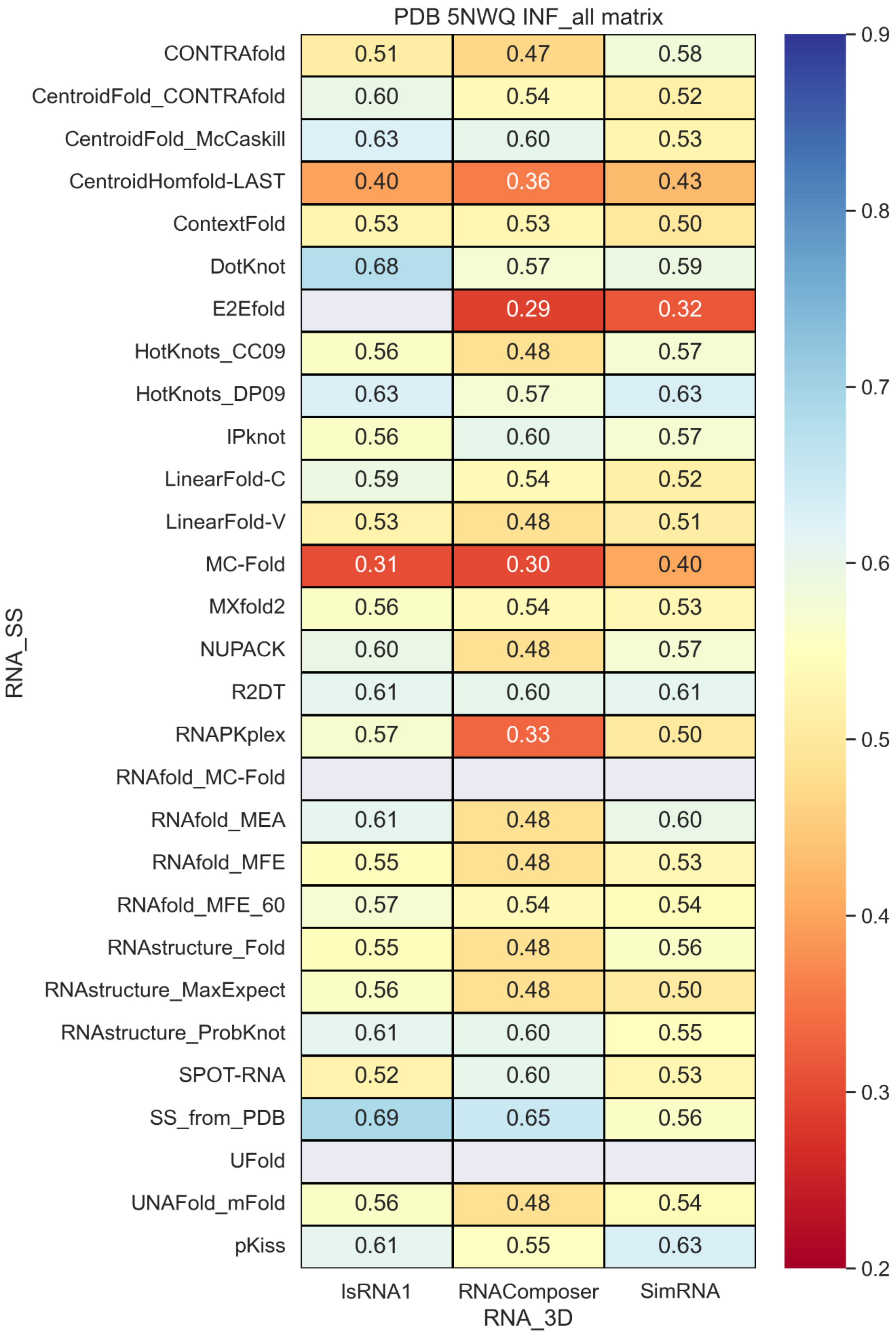


**Figure S56.** INF_all values with different RNA secondary structure methods and RNA 3D methods.

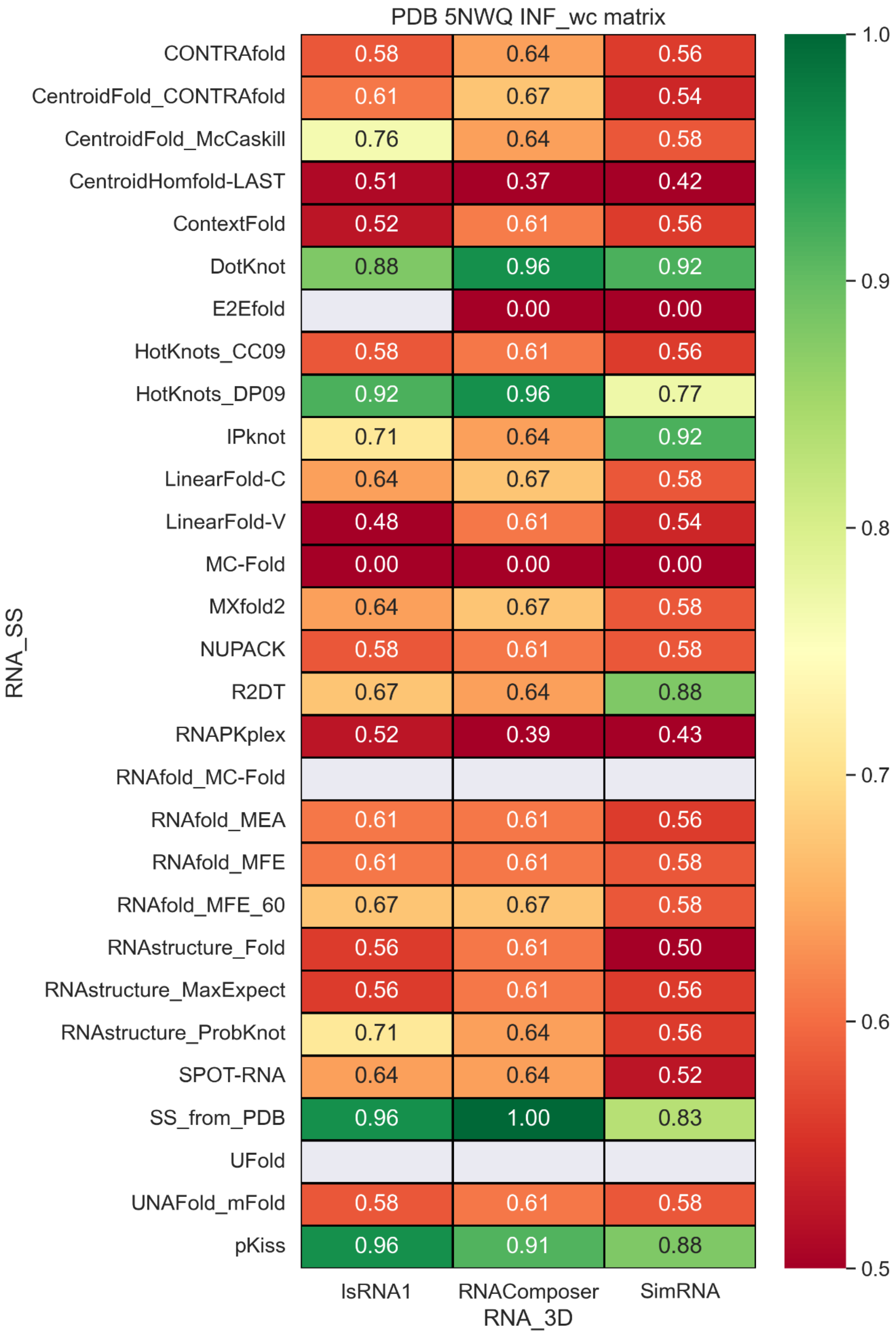


**Figure S57.** INF_wc values with different RNA secondary structure methods and RNA 3D methods.

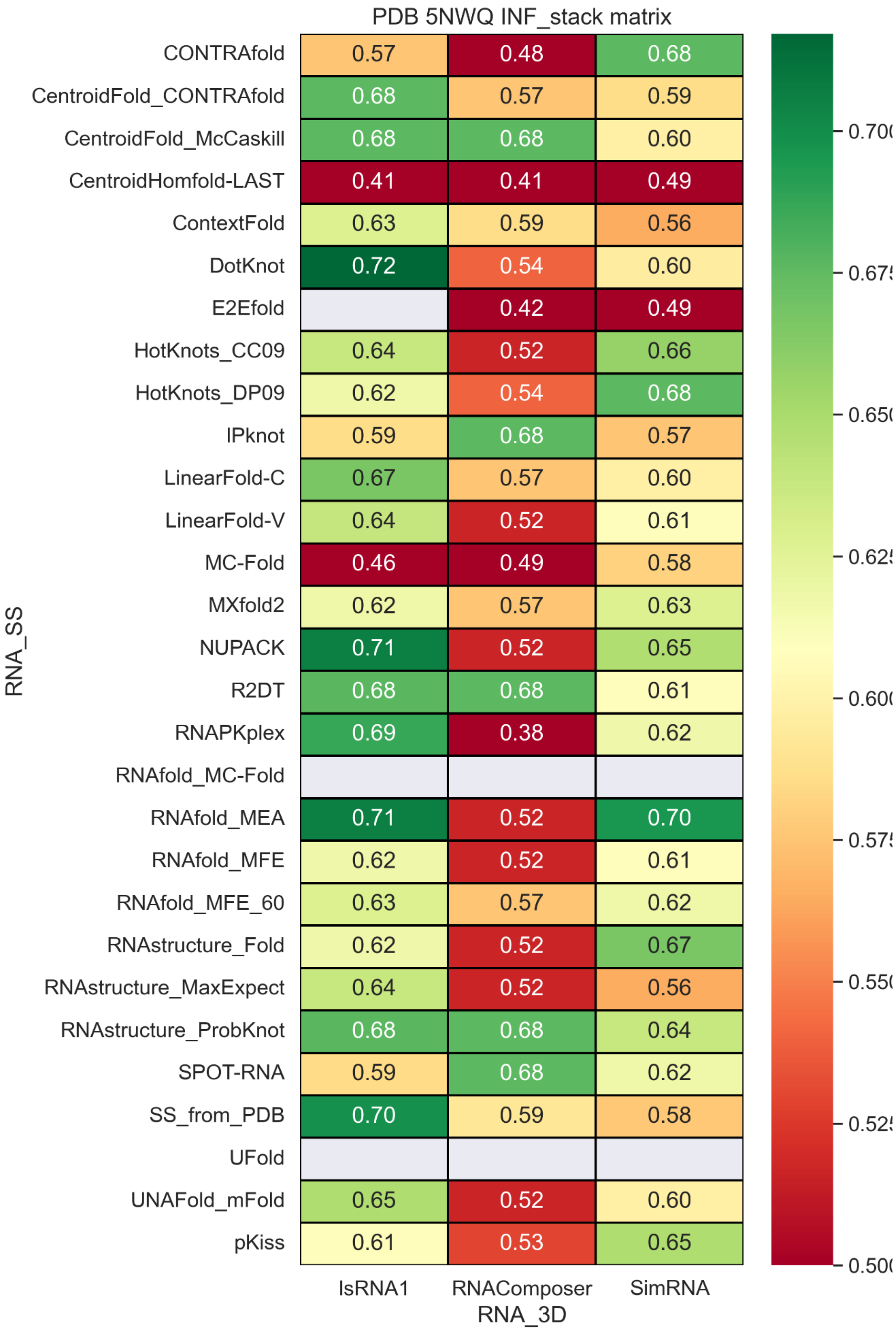


**Figure S58.** INF_stack values with different RNA secondary structure methods and RNA 3D methods.

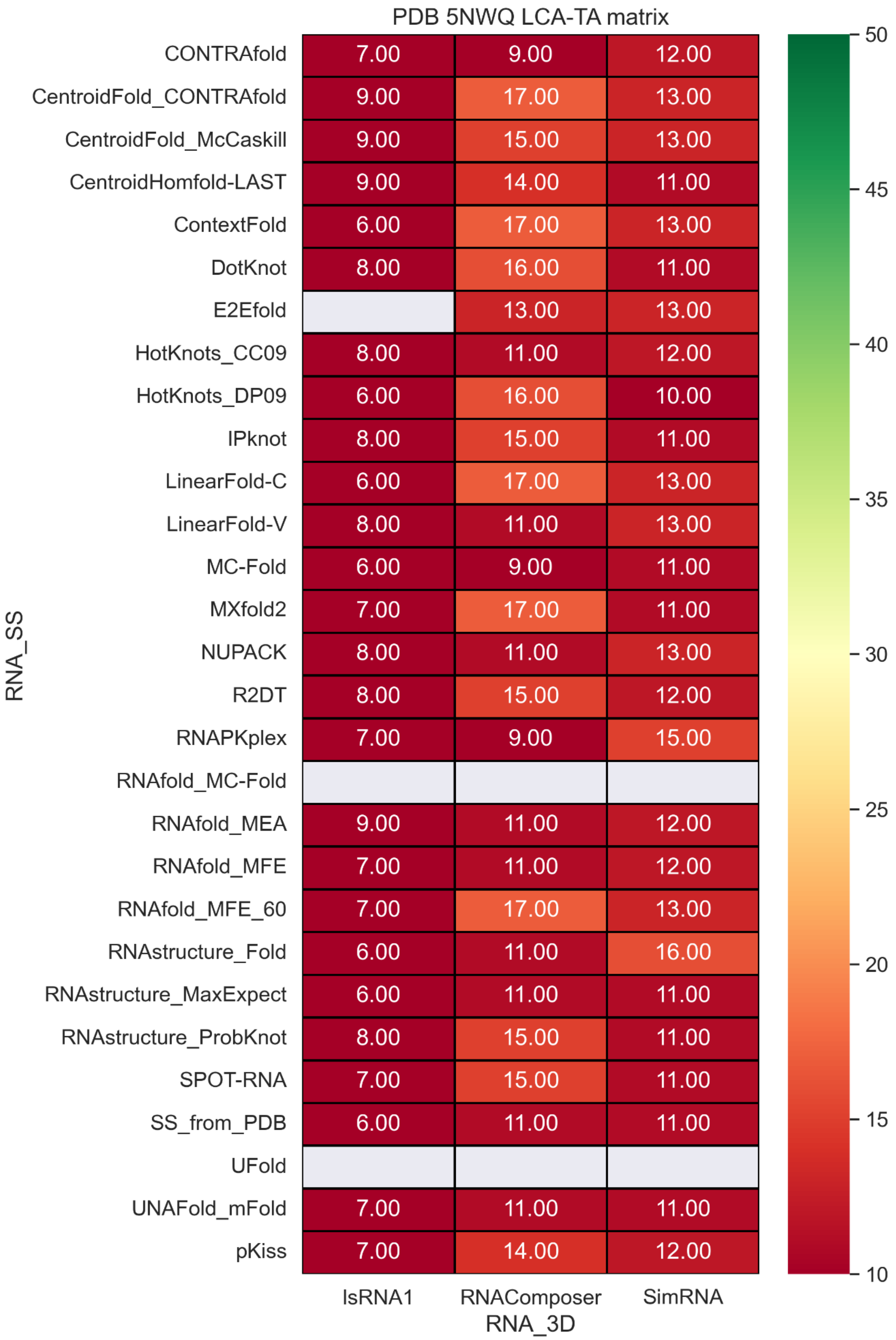


**Figure S59**. LCS-TA values with different RNA secondary structure methods and RNA 3D methods.

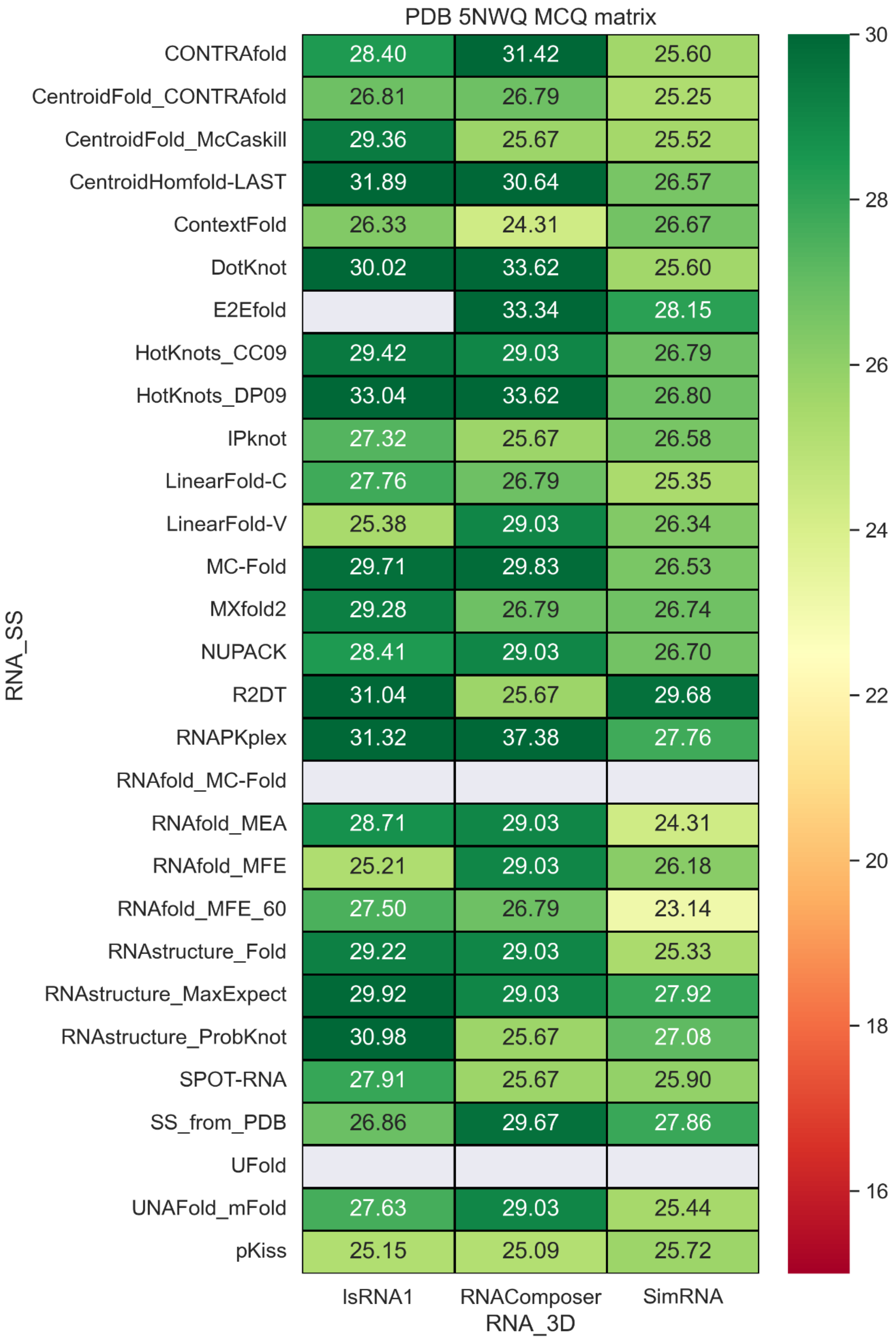


**Figure S60.** MCQ values with different RNA secondary structure methods and RNA 3D methods.

### PDB 4ENC

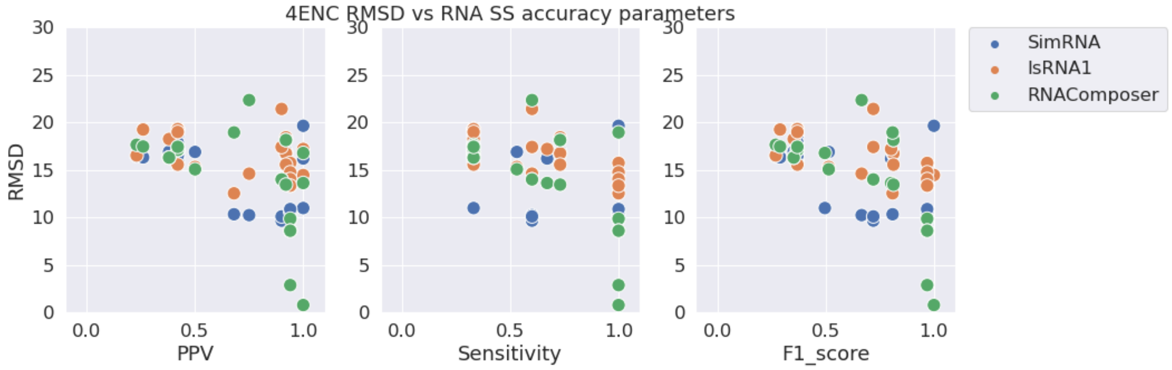


**Figure S61.** RMSD vs. PPV (left), Sensitivity (middle), and F1-score (right) for stem-loop structure PDB 4ENC.

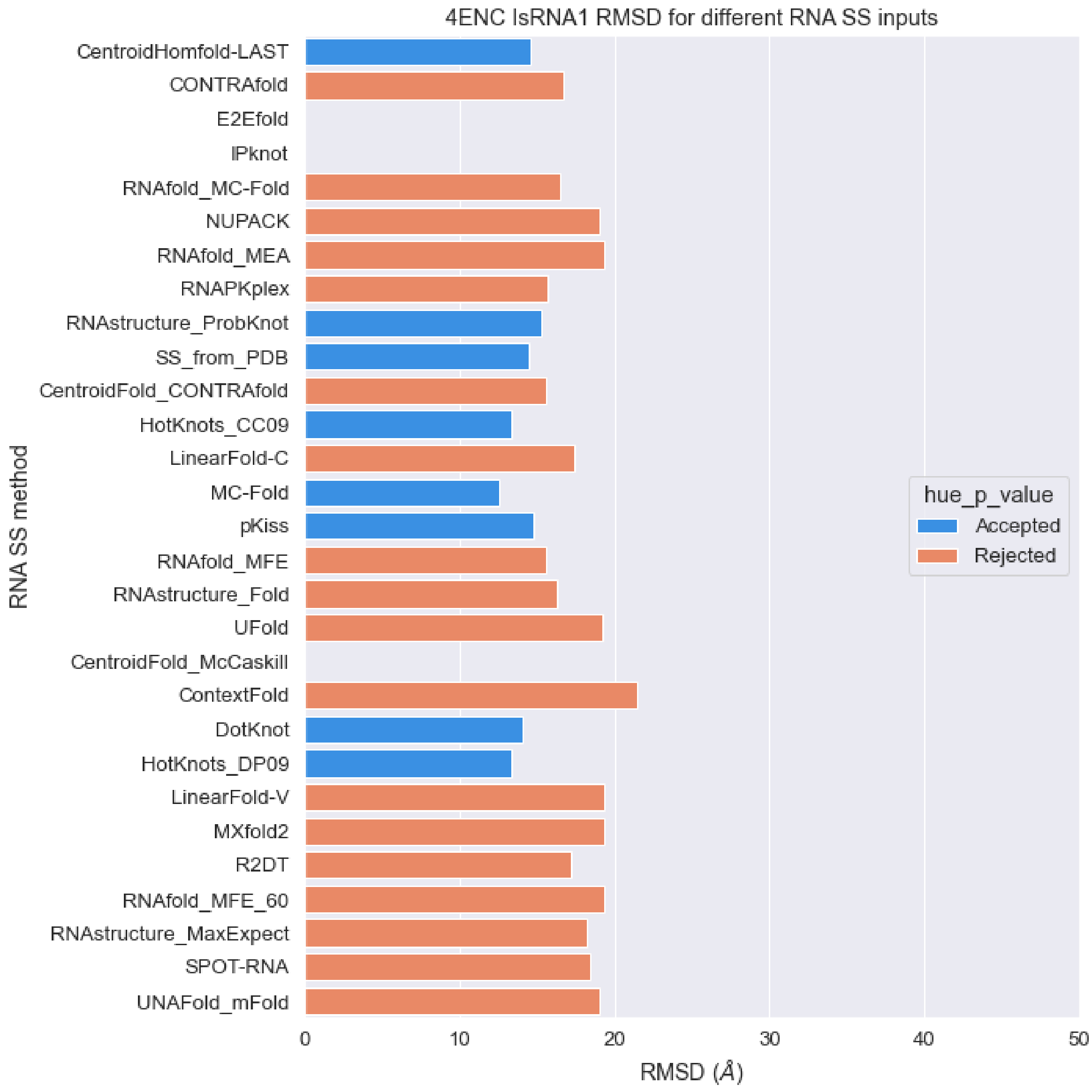


**Figure S62.** RMSD values of predicted structures for PDB 4ENC sequence with different RNA secondary structures as constraints with IsRNA1 package (orange: p-value > 0.01, blue: p-value < 0.01).

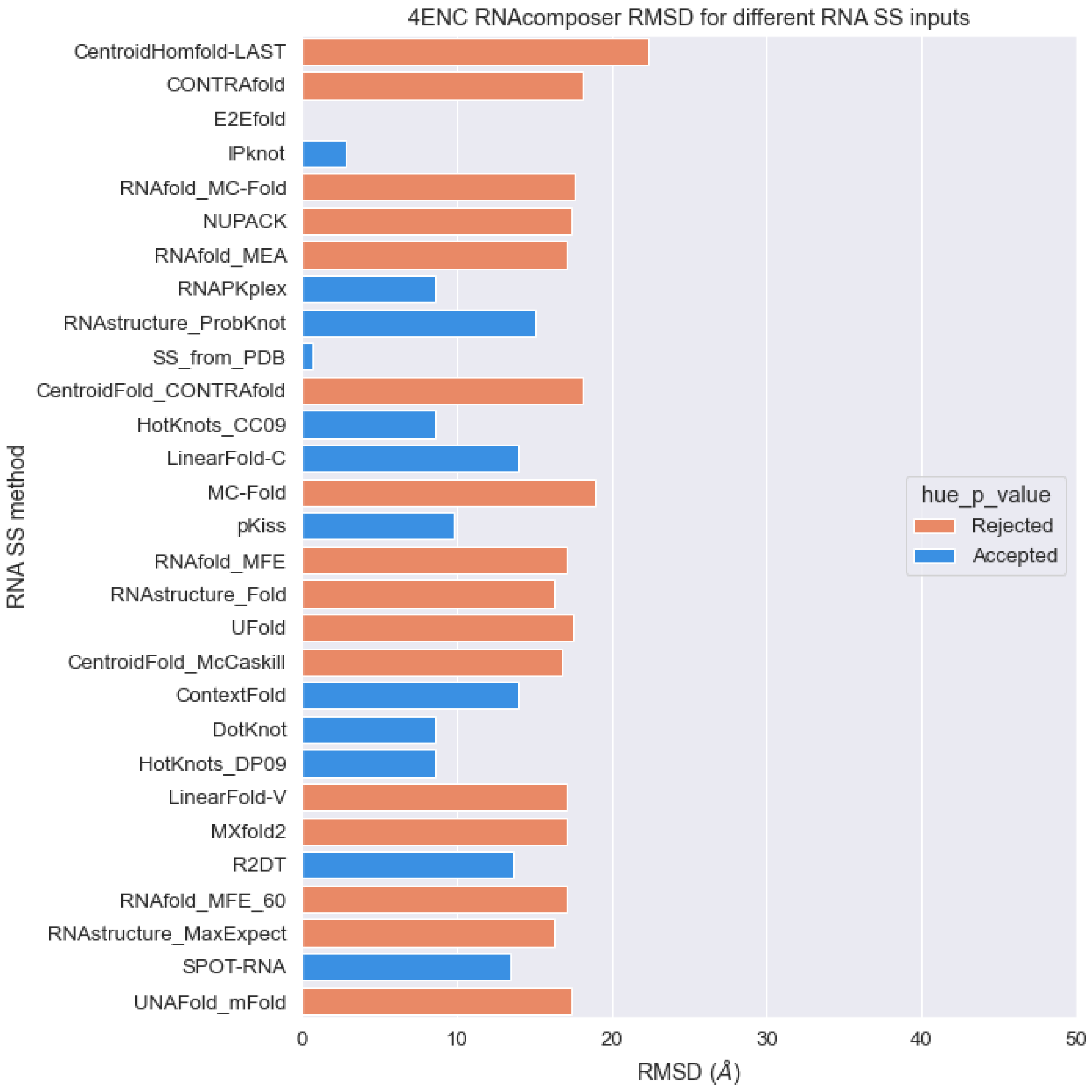


**Figure S63.** RMSD values of predicted structures for PDB 4ENC sequence with different RNA secondary structures as constraints with RNAcomposer package (orange: p-value > 0.01, blue: p-value < 0.01).

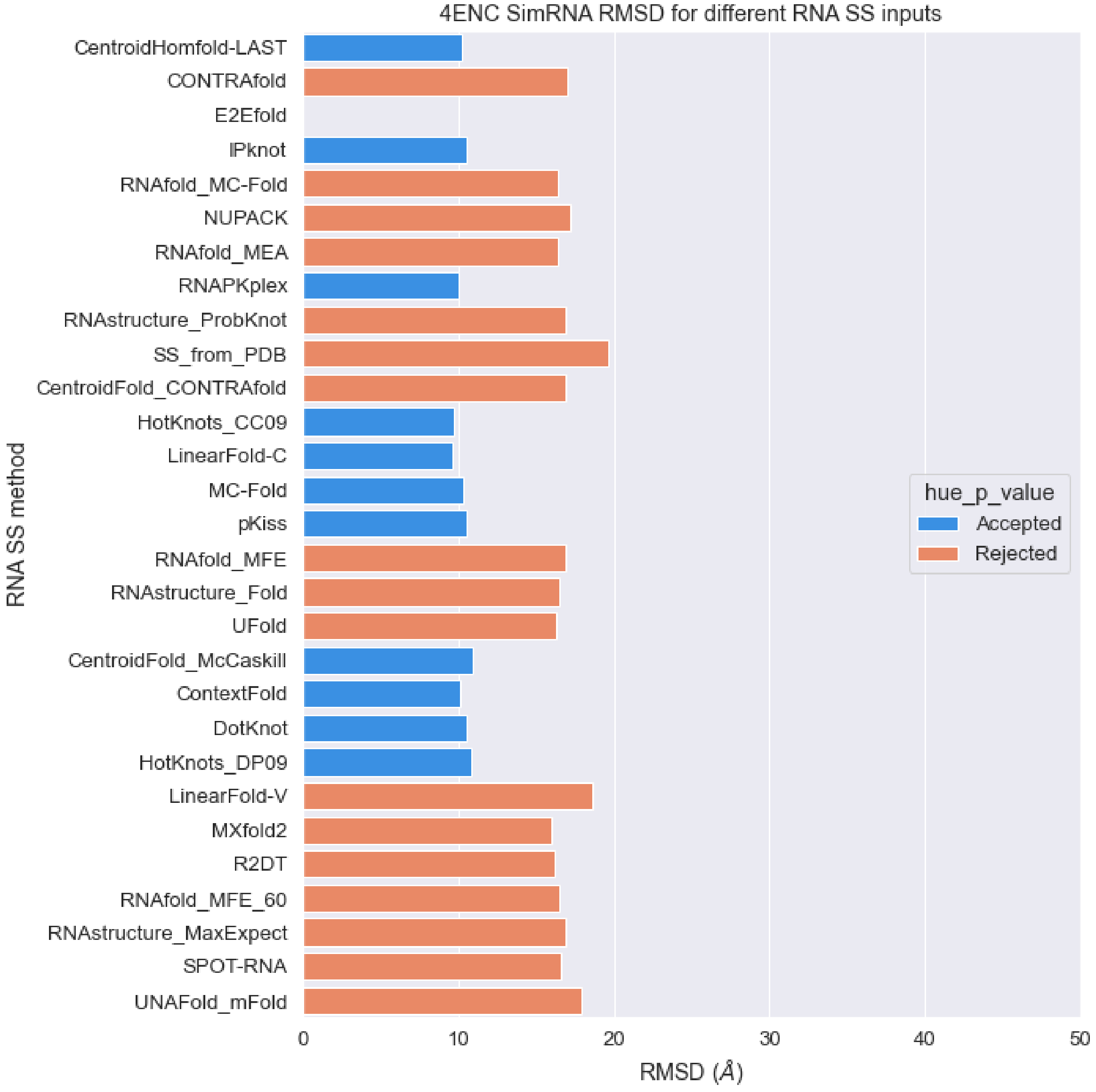


**Figure S64.** RMSD values of predicted structures for PDB 4ENC sequence with different RNA secondary structures as constraints with SimRNA package (orange: p-value > 0.01, blue: p-value < 0.01).

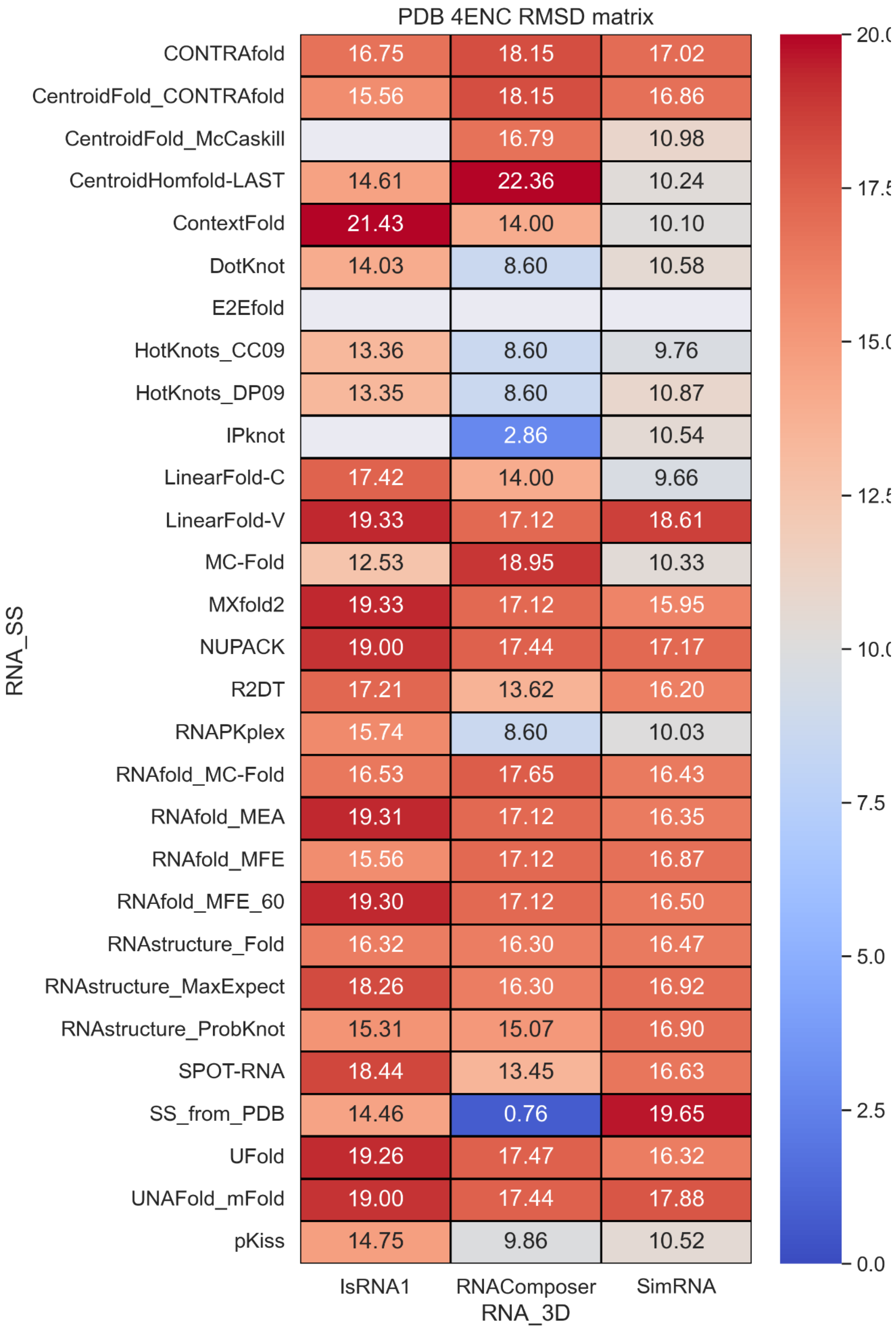


**Figure S65.** RMSD values with different RNA secondary structure methods and RNA 3D methods.

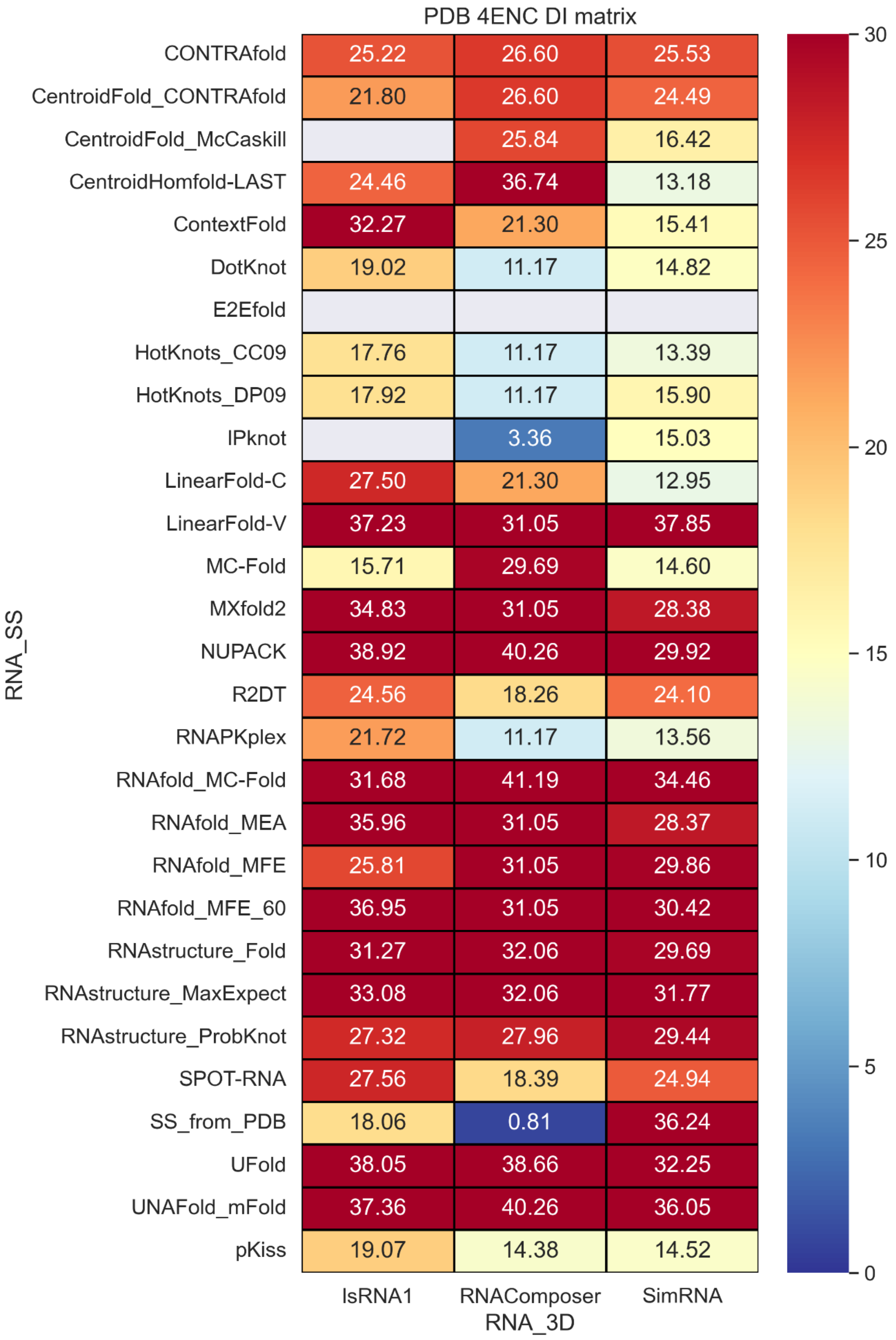


**Figure S66.** Deformation index values with different RNA secondary structure methods and RNA 3D methods.

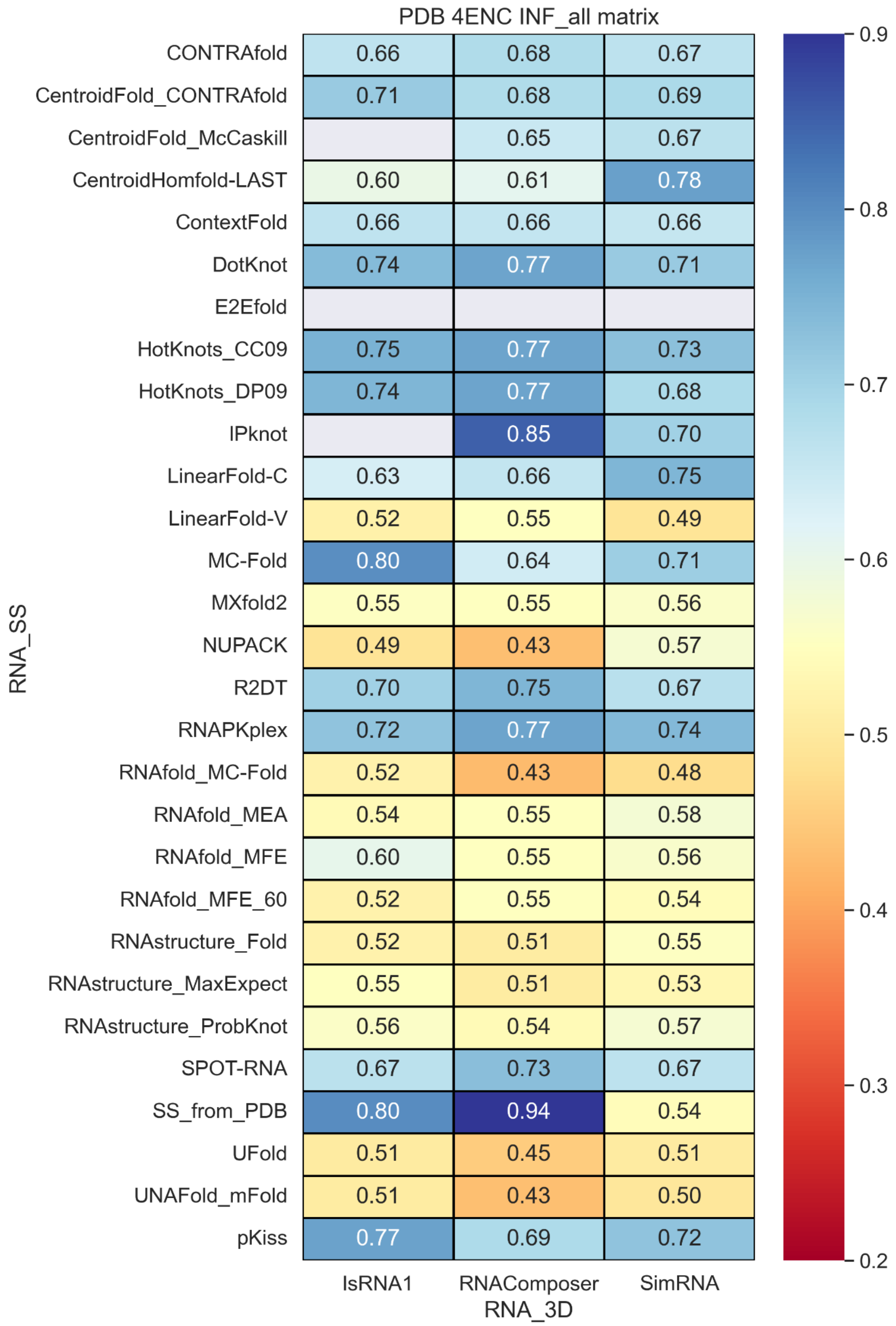


**Figure S67.** INF_all values with different RNA secondary structure methods and RNA 3D methods.

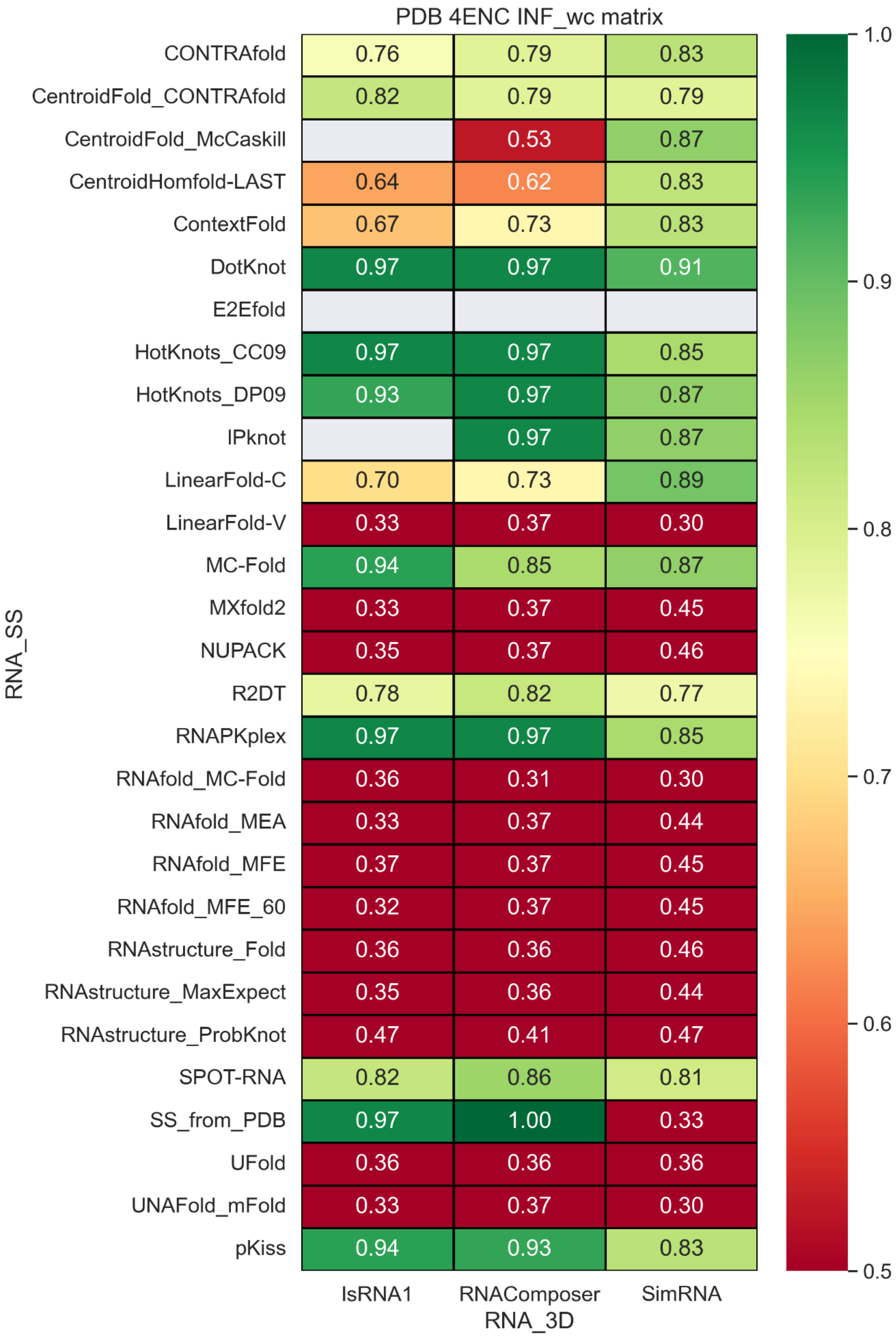


**Figure S68.** INF_wc values with different RNA secondary structure methods and RNA 3D methods.

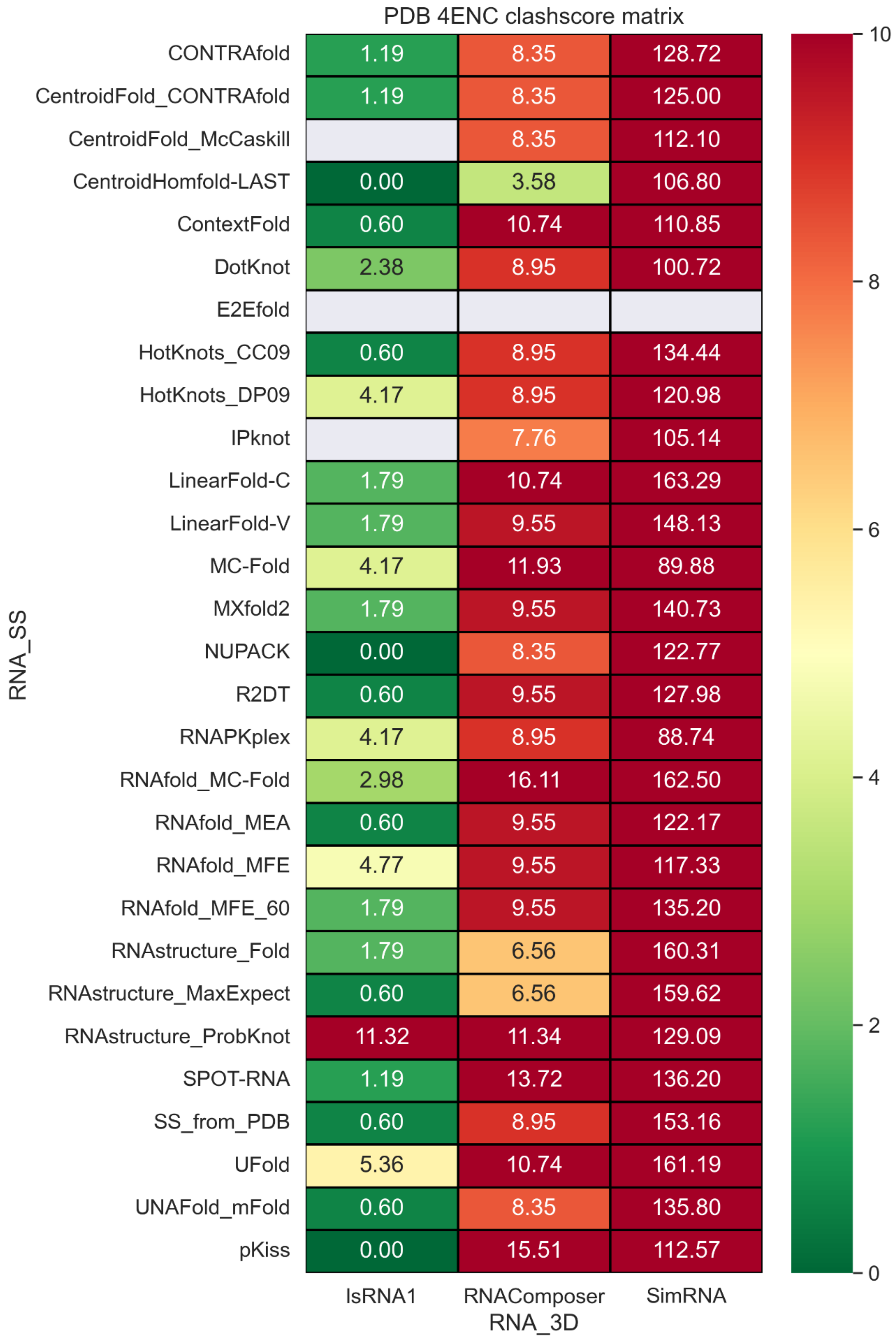


**Figure S69.** Clashscore values with different RNA secondary structure methods and RNA 3D methods.

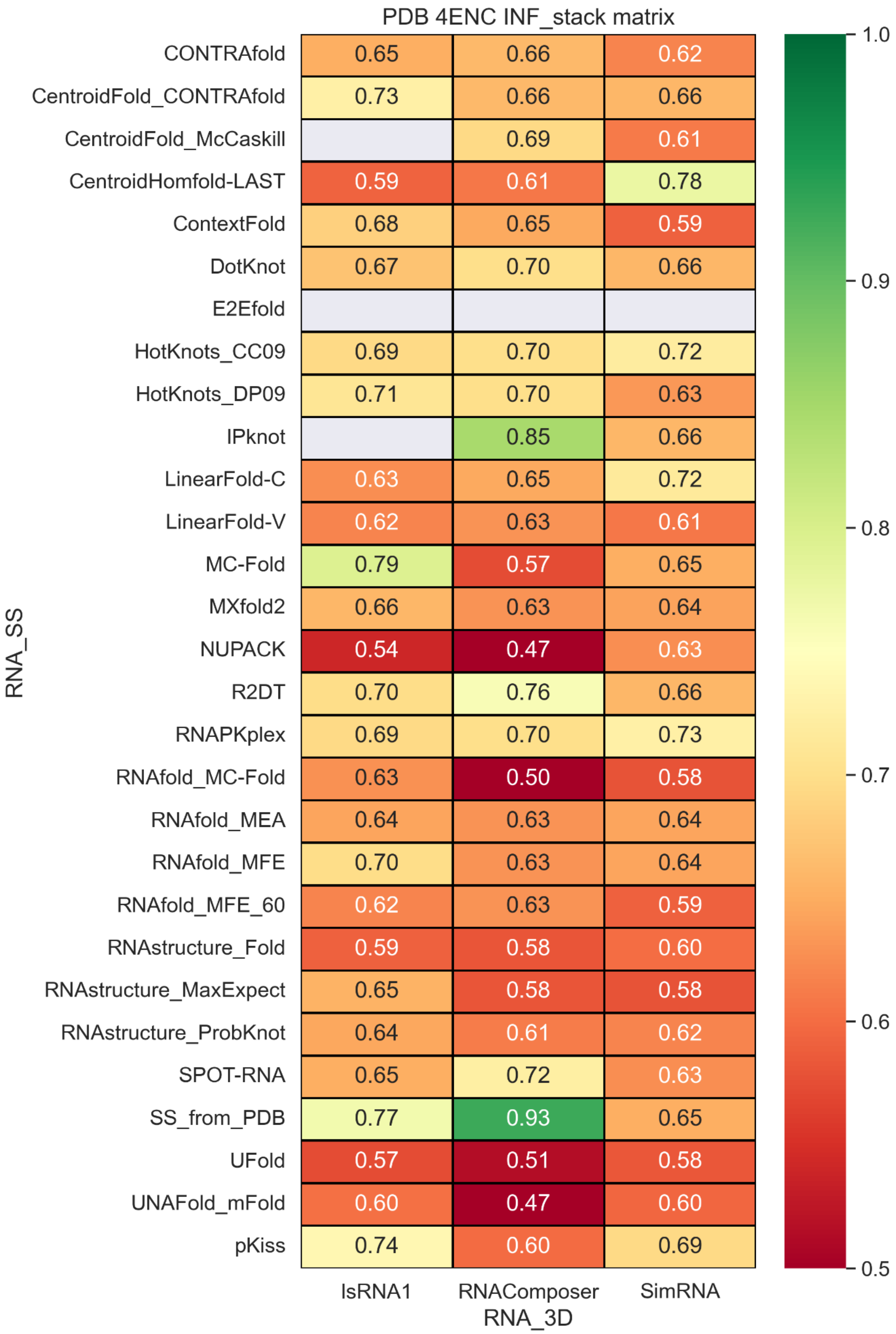


**Figure S70.** INF_stack values with different RNA secondary structure methods and RNA 3D methods.

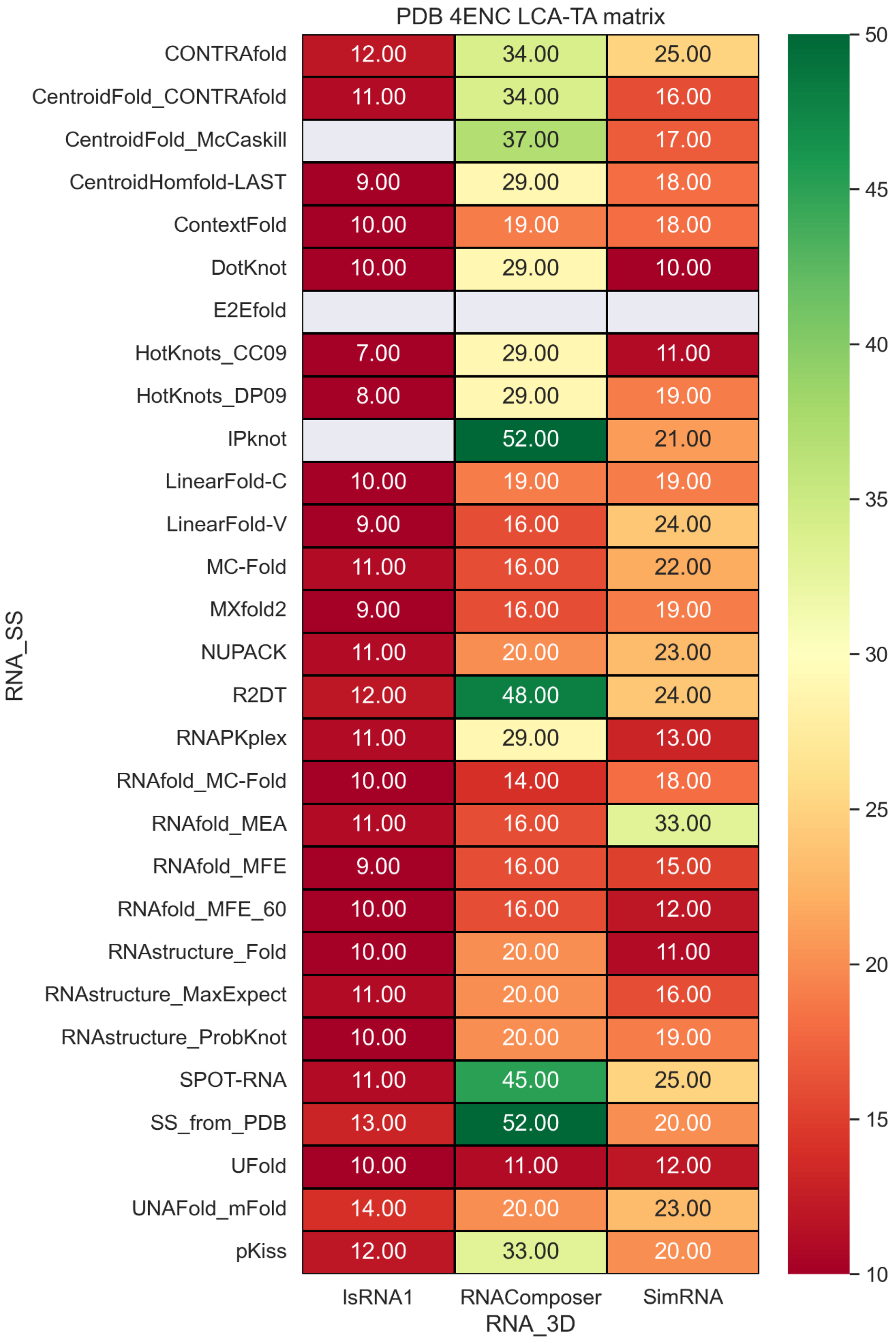


**Figure S71.** LCS-TA values with different RNA secondary structure methods and RNA 3D methods.

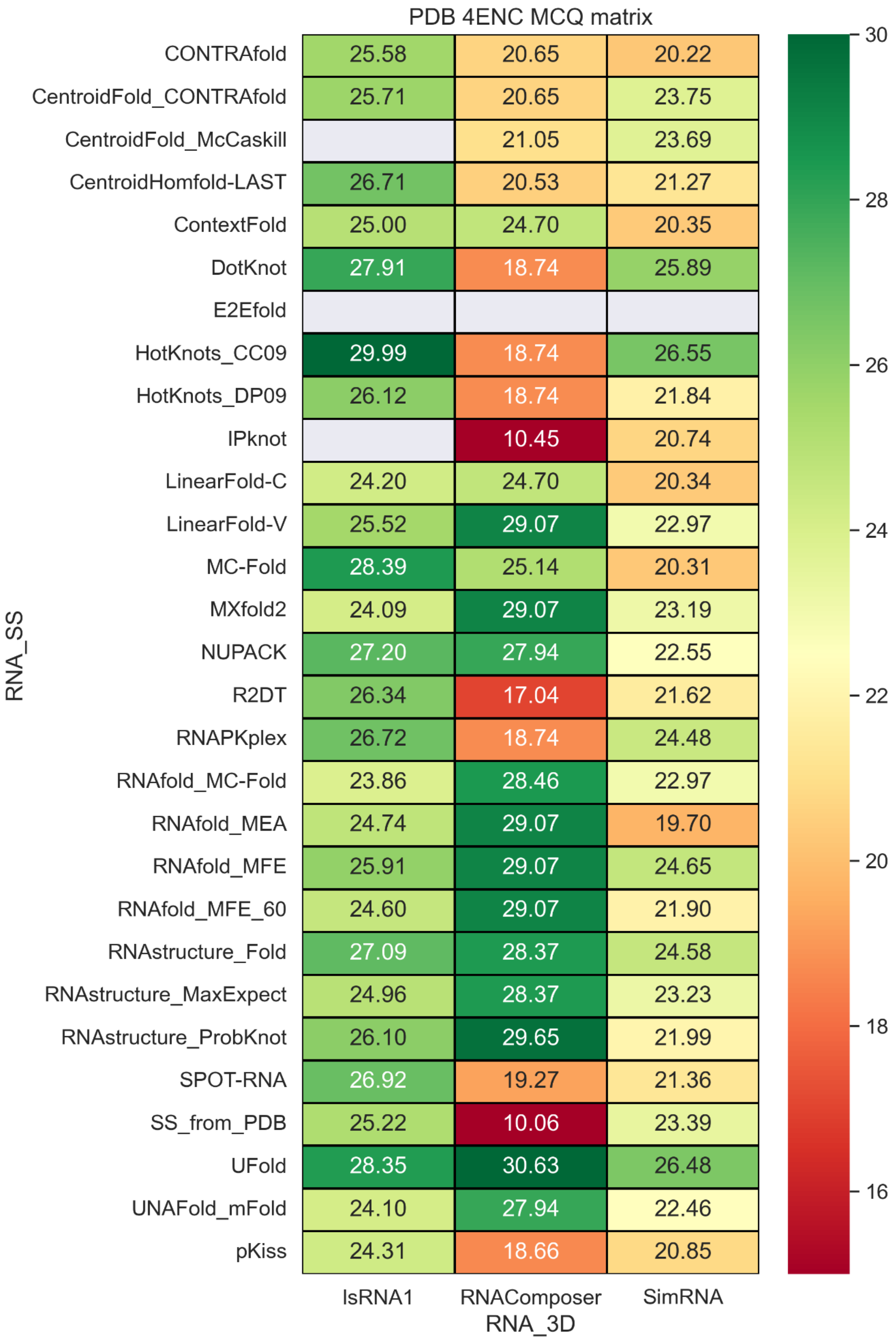


**Figure S72.** MCQ values with different RNA secondary structure methods and RNA 3D methods.

### PDB 6P2H

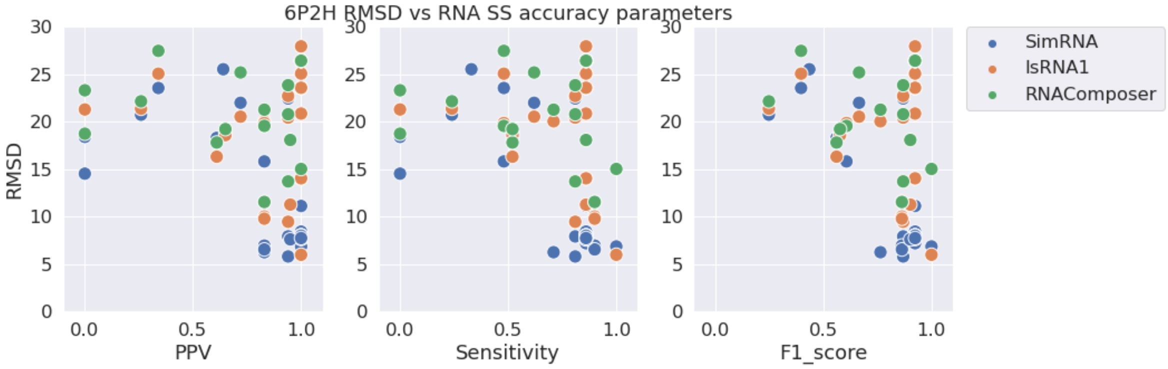


**Figure S73.** RMSD vs. PPV (left), Sensitivity (middle), and F1-score (right) for stem-loop structure PDB 6P2H.

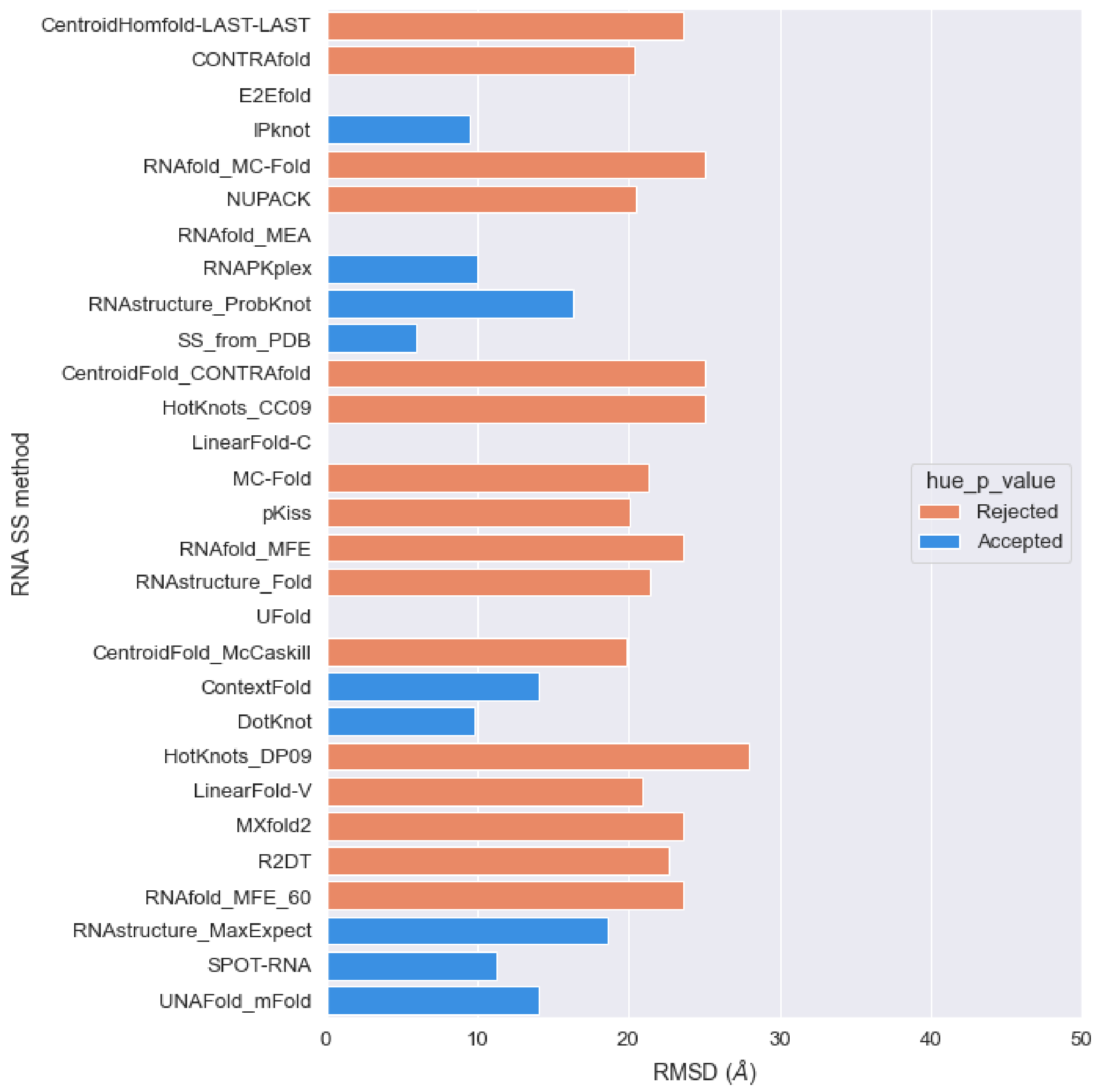


**Figure S74.** RMSD values of predicted structures for PDB 6P2H sequence with different RNA secondary structures as constraints with IsRNA1 package (orange: p-value > 0.01, blue: p-value < 0.01).

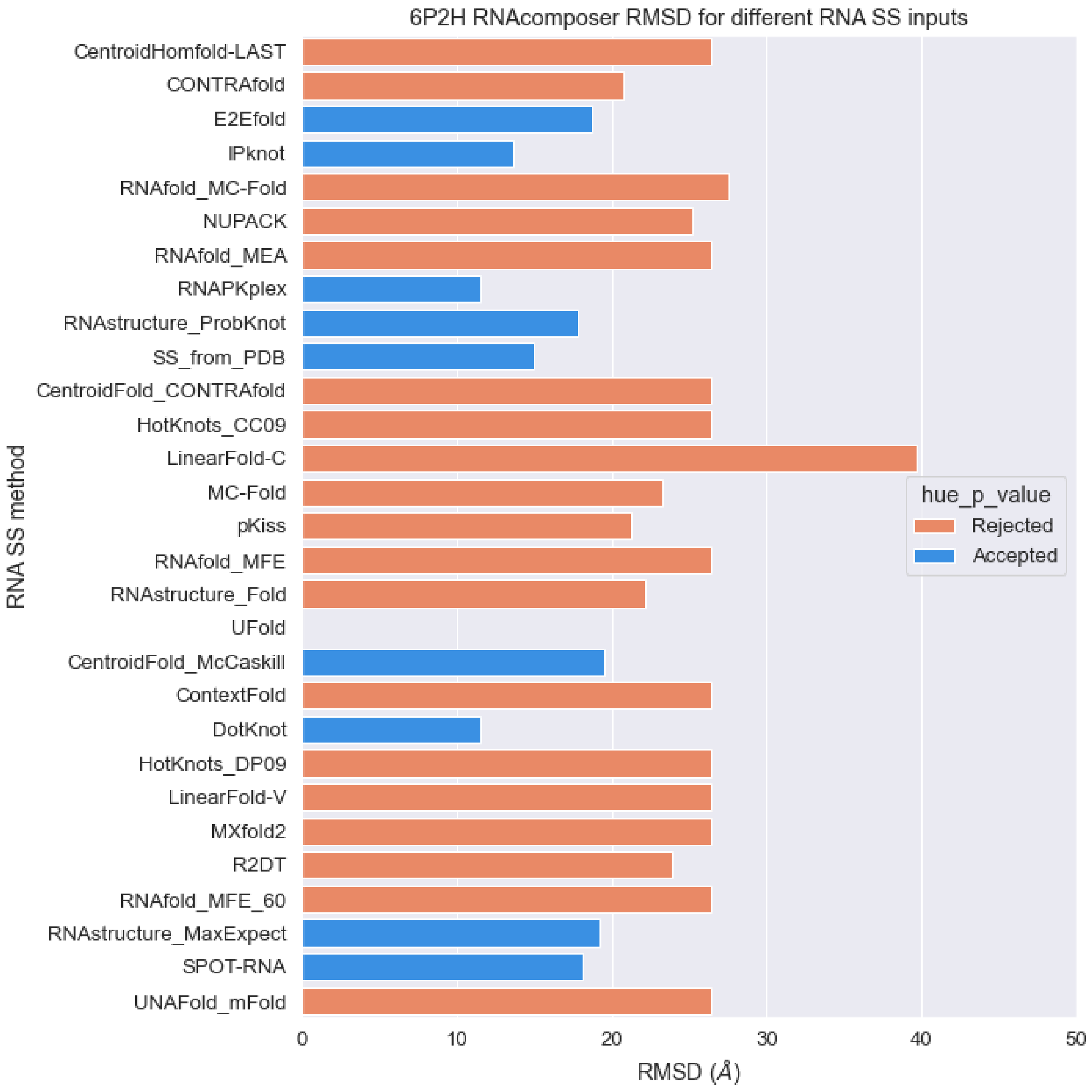


**Figure S75.** RMSD values of predicted structures for PDB 6P2H sequence with different RNA secondary structures as constraints with RNAcomoser package (orange: p-value > 0.01, blue: p-value < 0.01).

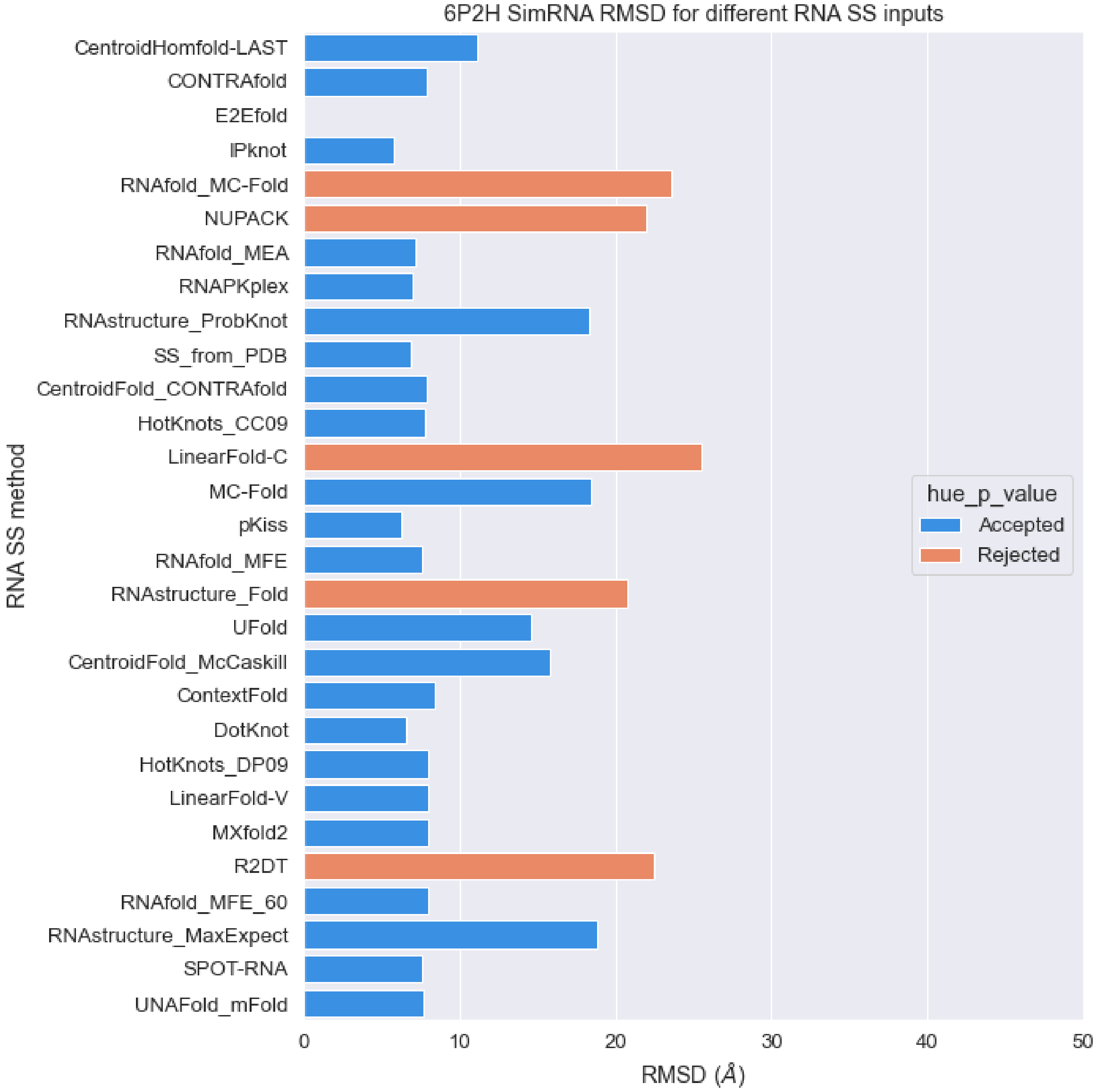


**Figure S76.** RMSD values of predicted structures for PDB 6P2H sequence with different RNA secondary structures as constraints with SimRNA package (orange: p-value > 0.01, blue: p-value < 0.01).

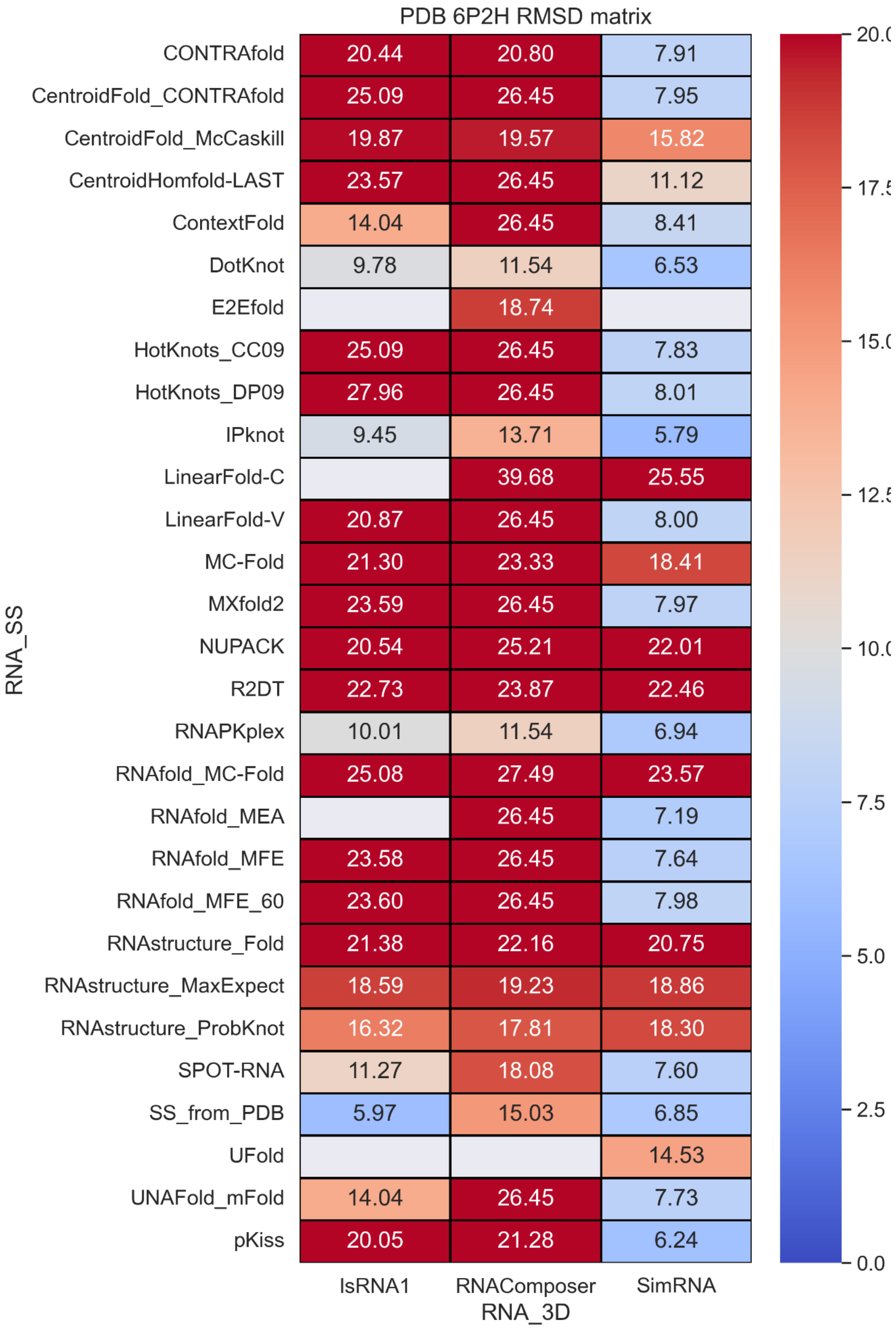


**Figure S77.** RMSD values with different RNA secondary structure methods and RNA 3D methods.

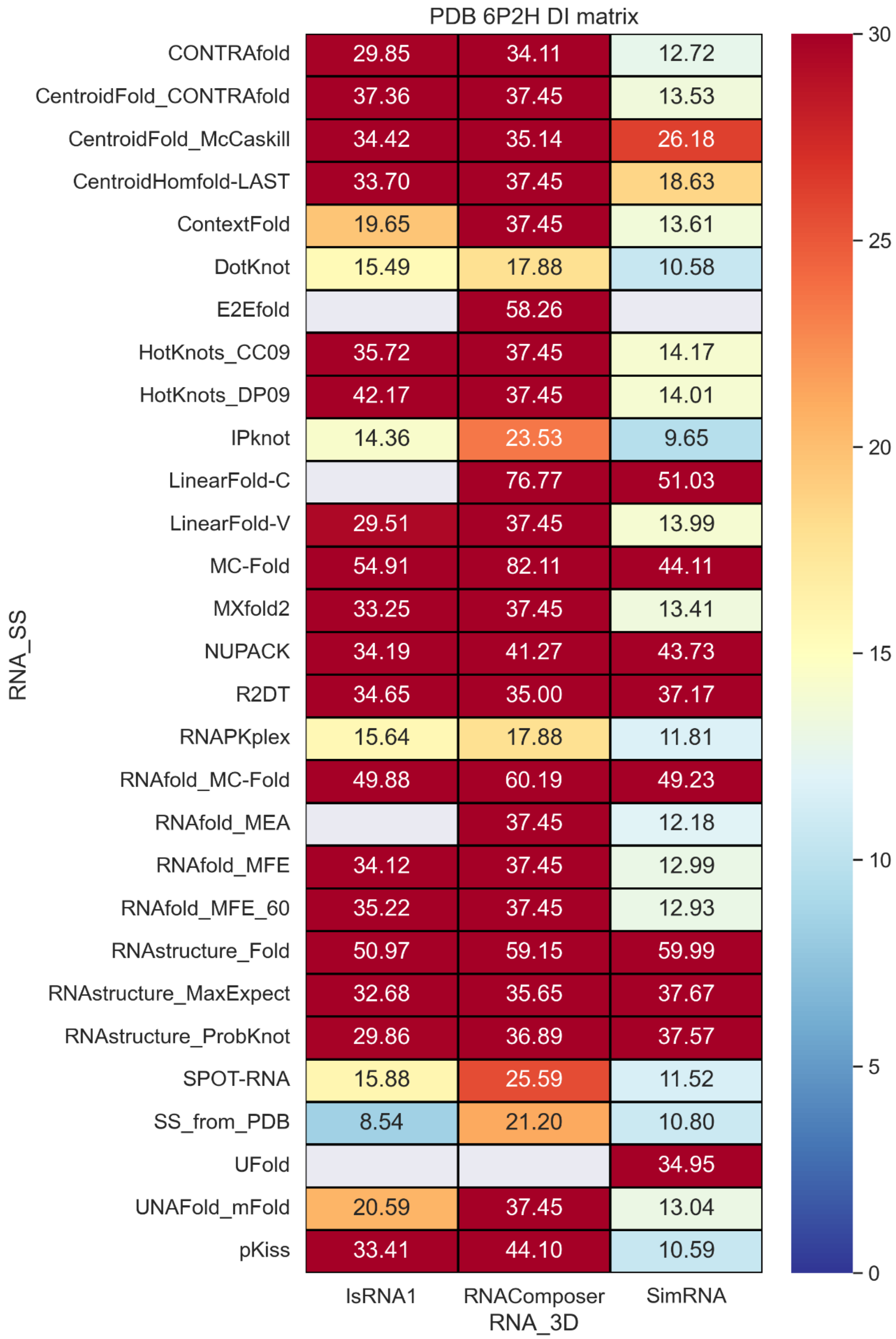


**Figure S78.** Deformation index values with different RNA secondary structure methods and RNA 3D methods.

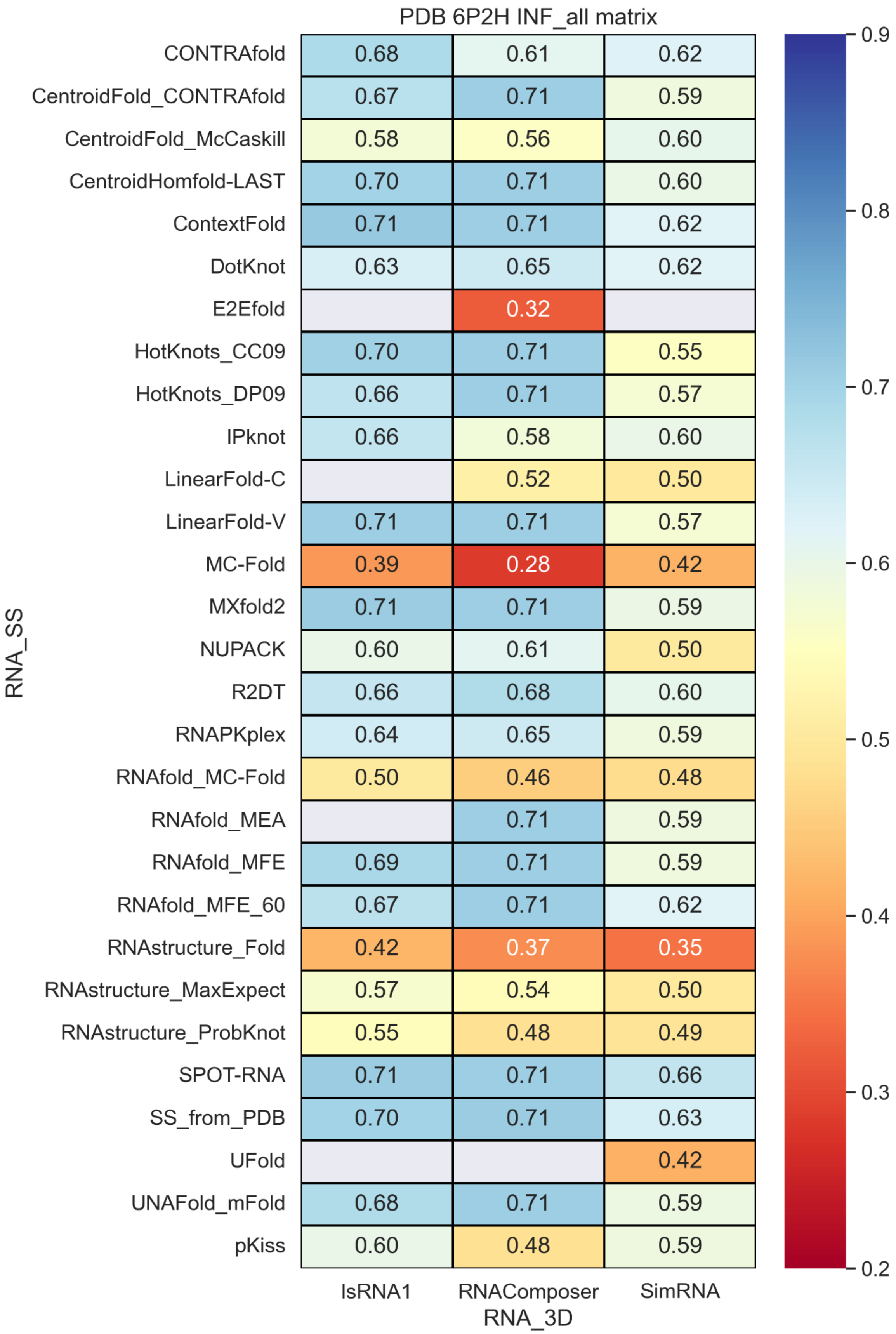


**Figure S79.** INF_all values with different RNA secondary structure methods and RNA 3D methods.

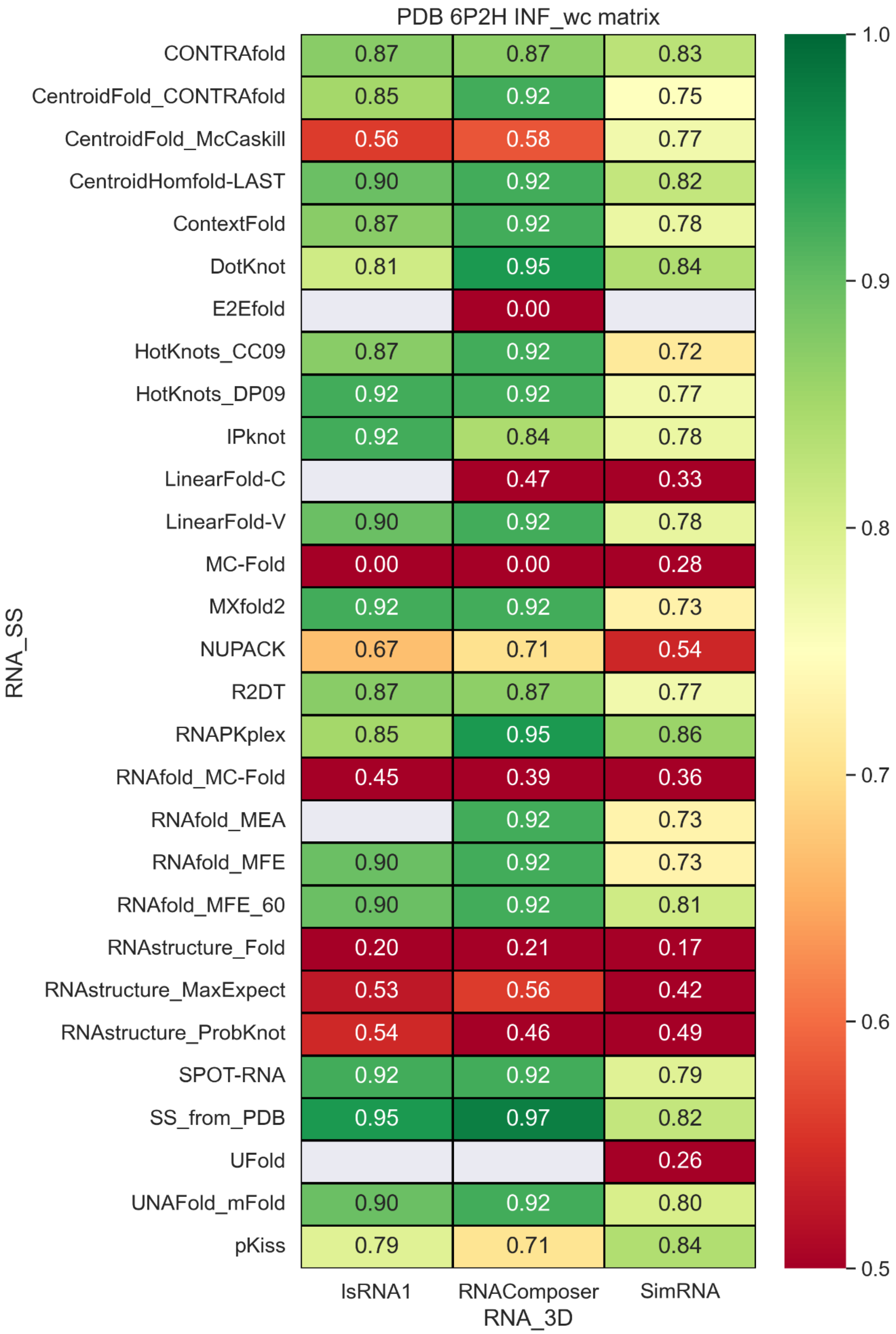


**Figure S80.** INF_wc values with different RNA secondary structure methods and RNA 3D methods.

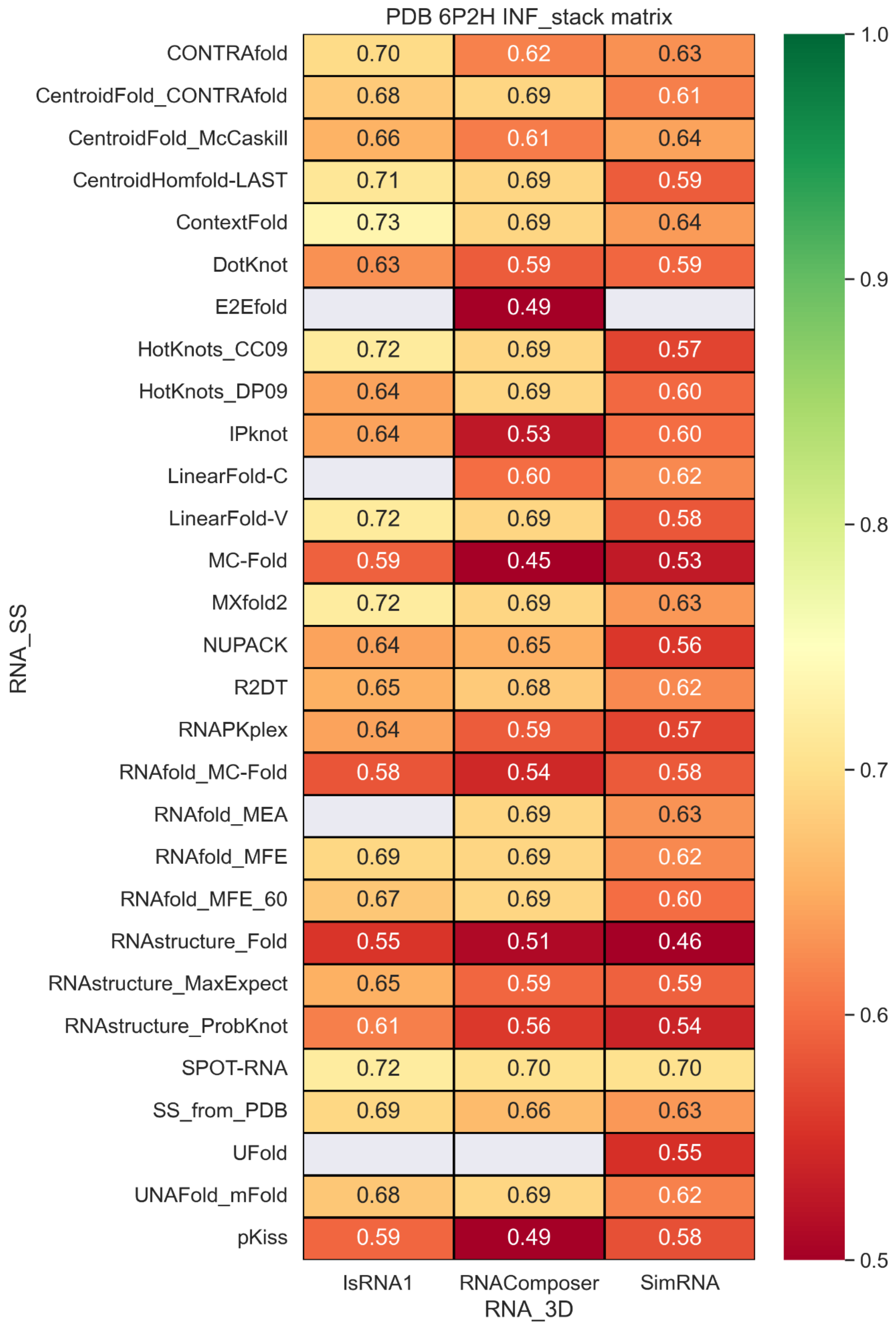


**Figure S81.** INF_stack values with different RNA secondary structure methods and RNA 3D methods.

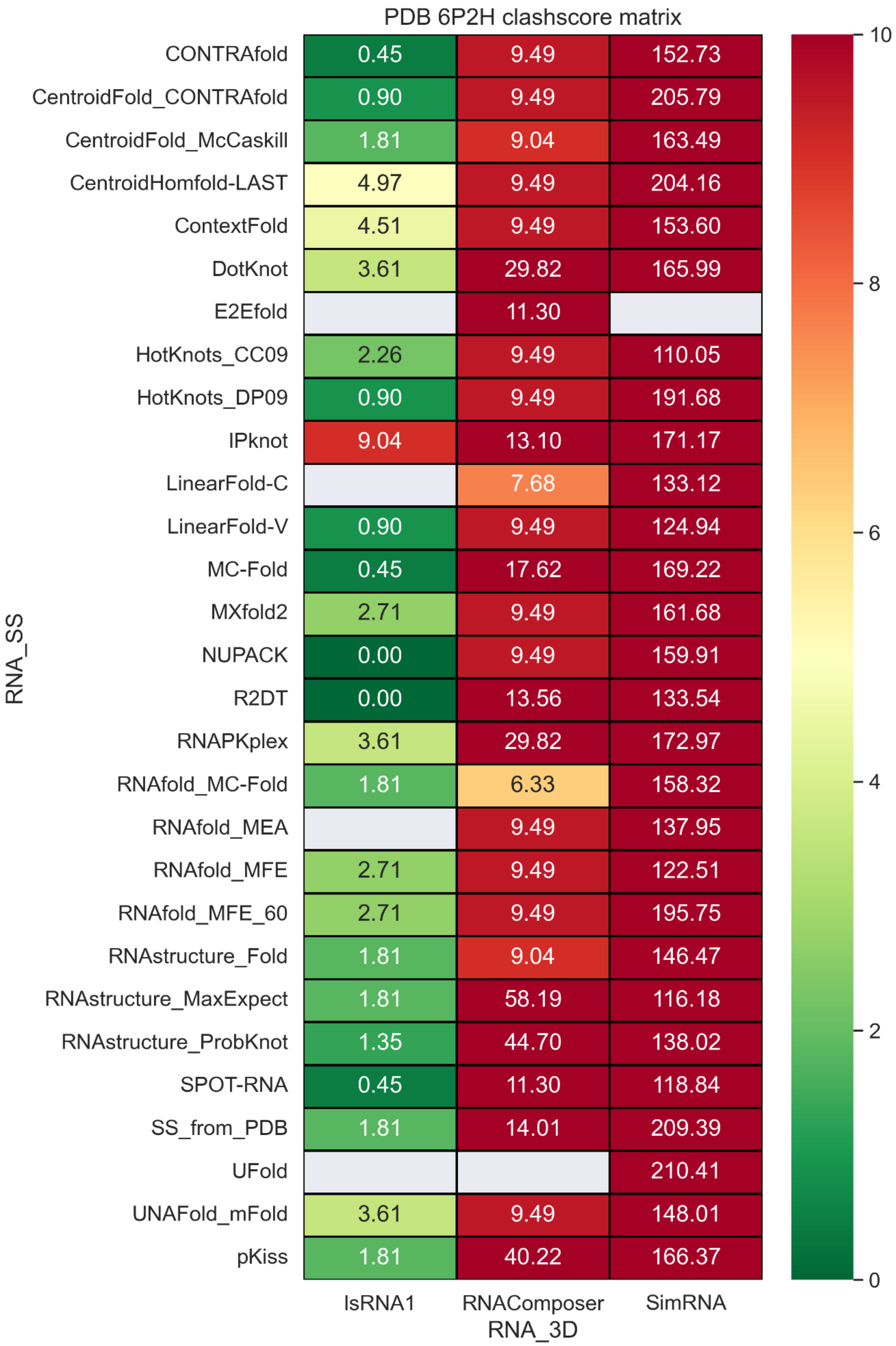


**Figure S82.** Clashscore values with different RNA secondary structure methods and RNA 3D methods.

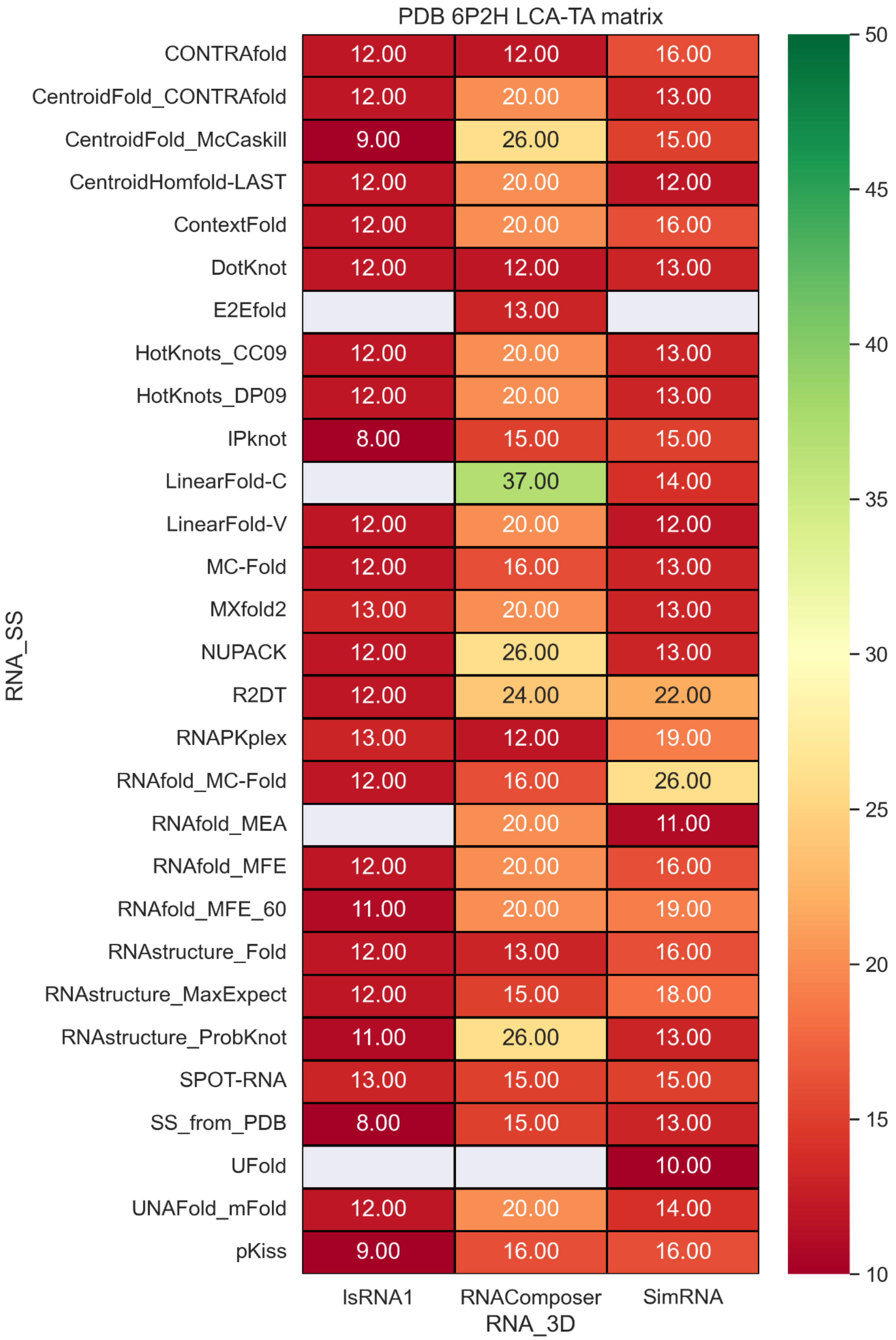


**Figure S83.** LCS-TA values with different RNA secondary structure methods and RNA 3D methods.

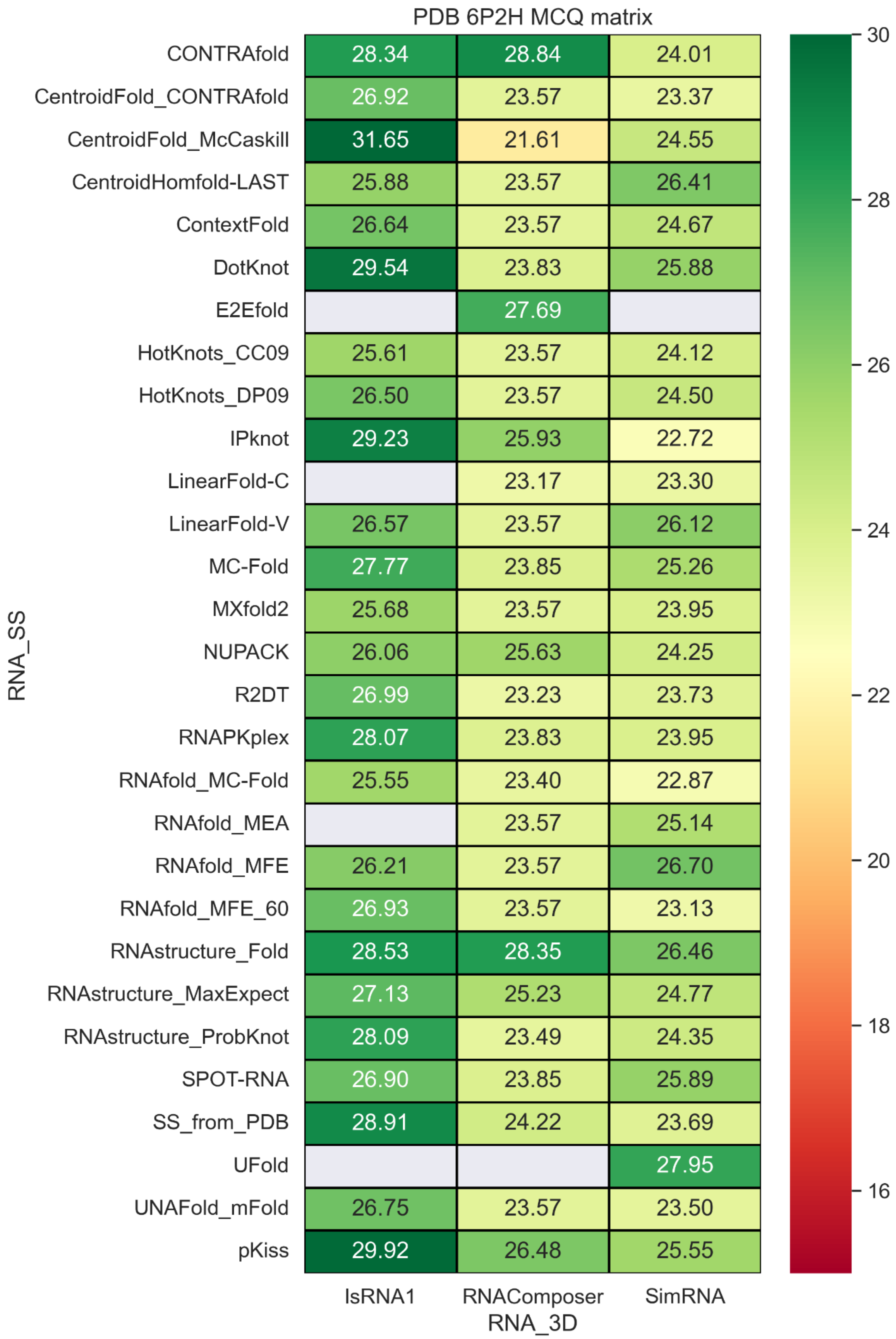


**Figure S84.** MCQ values with different RNA secondary structure methods and RNA 3D methods.

### PDB 3OWZ (Chain A)

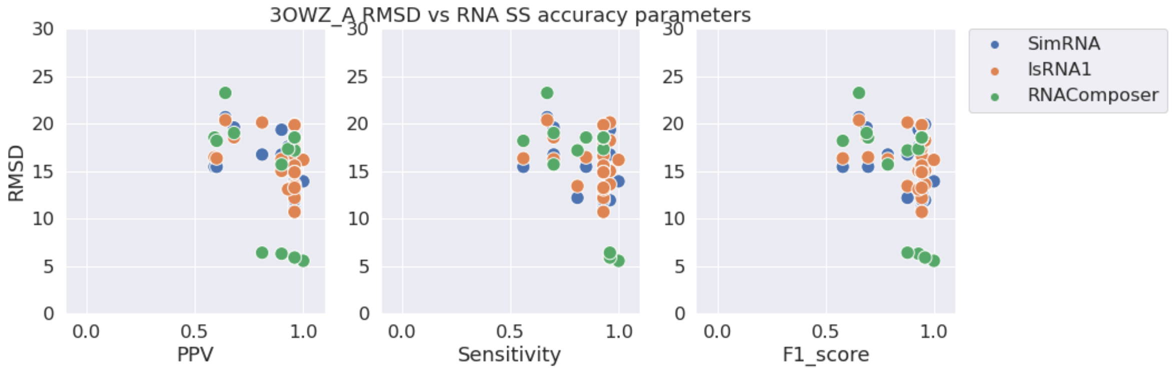


**Figure S85.** RMSD vs. PPV (left), Sensitivity (middle), and F1-score (right) for stem-loop structure PDB 3OWZ.

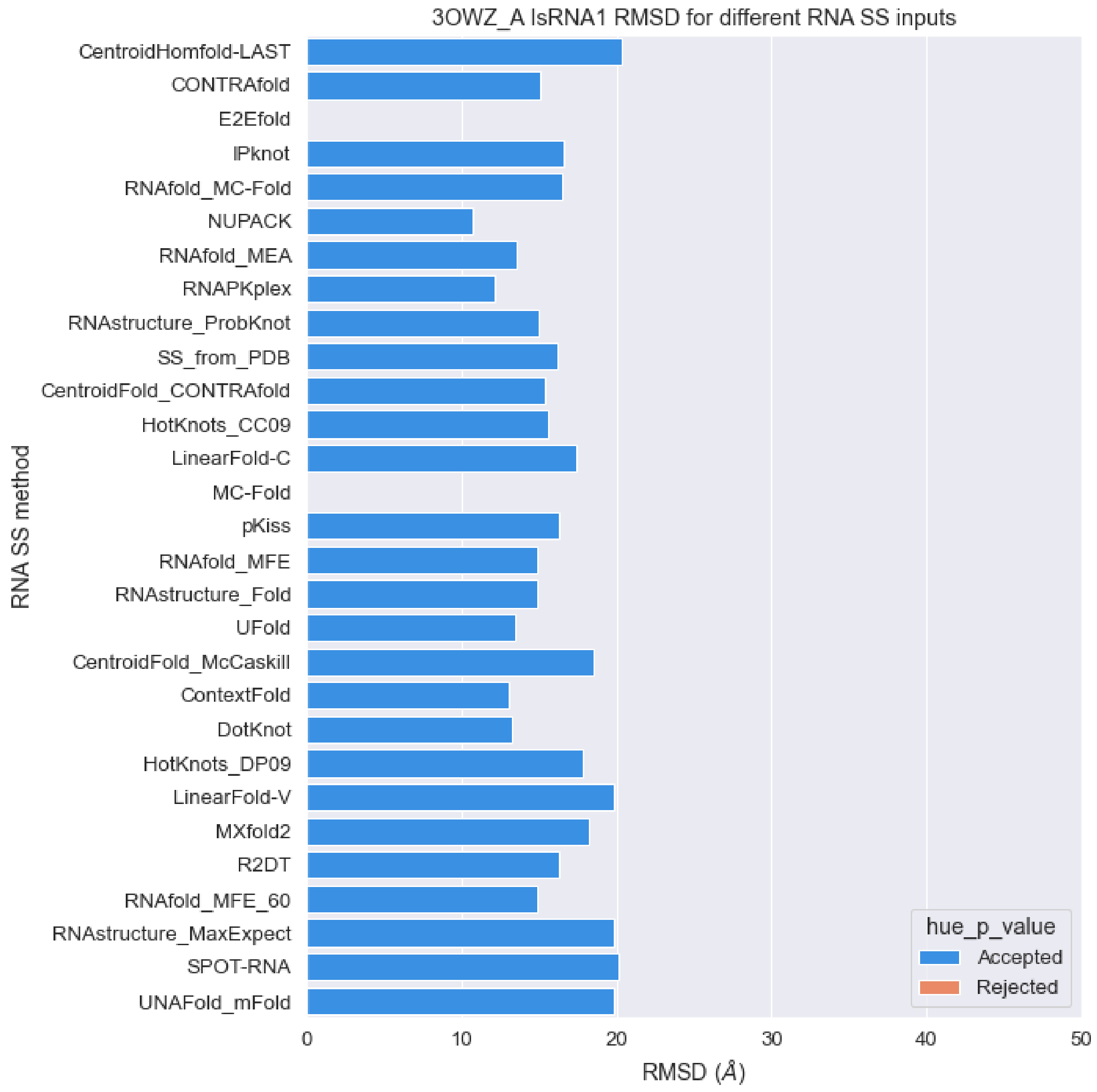


**Figure S86.** RMSD values of predicted structures for PDB 3OWZ sequence with different RNA secondary structures as constraints with IsRNA1 package (orange: p-value > 0.01, blue: p-value < 0.01).

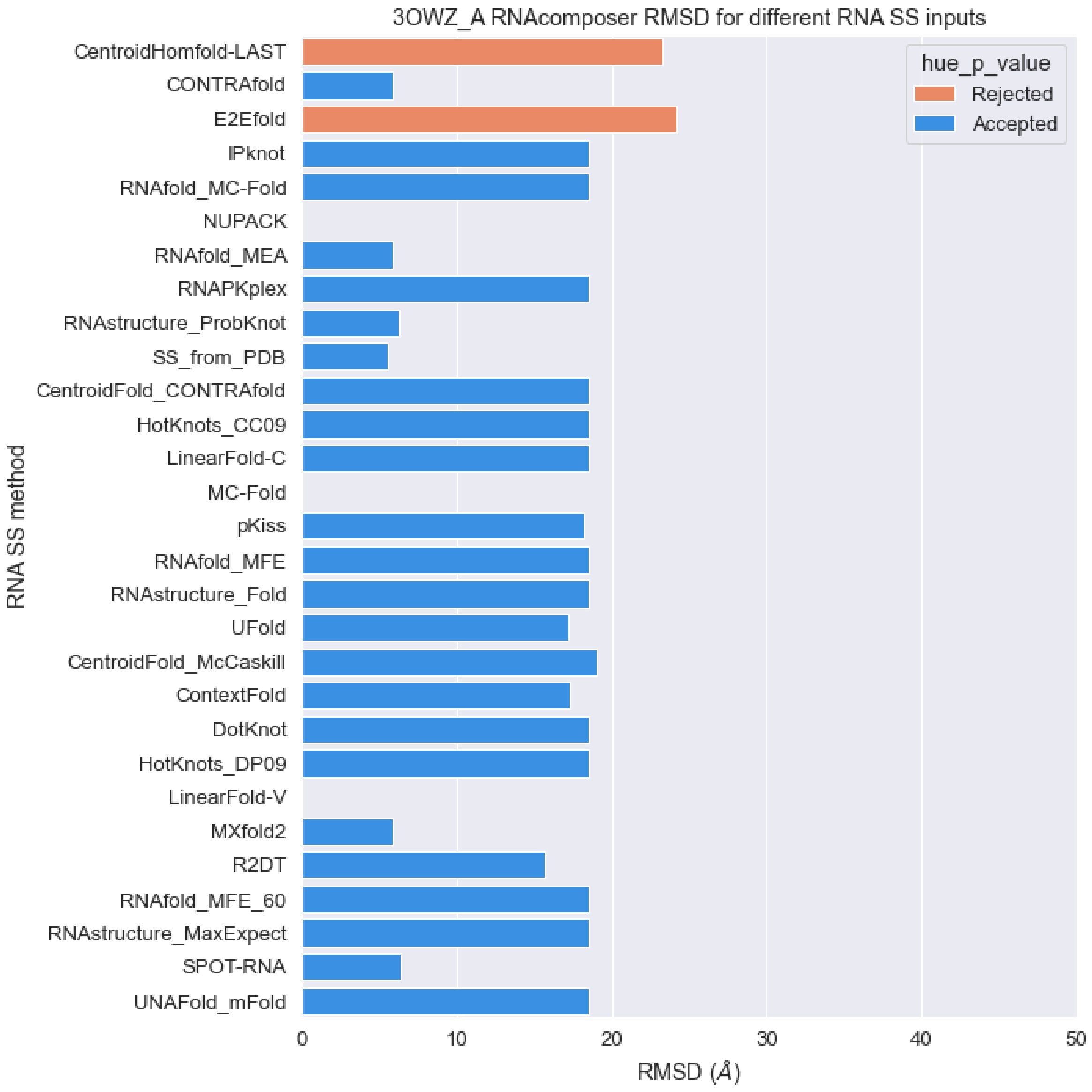


**Figure S87.** RMSD values of predicted structures for PDB 3OWZ sequence with different RNA secondary structures as constraints with RNAcomposer package (orange: p-value > 0.01, blue: p-value < 0.01).

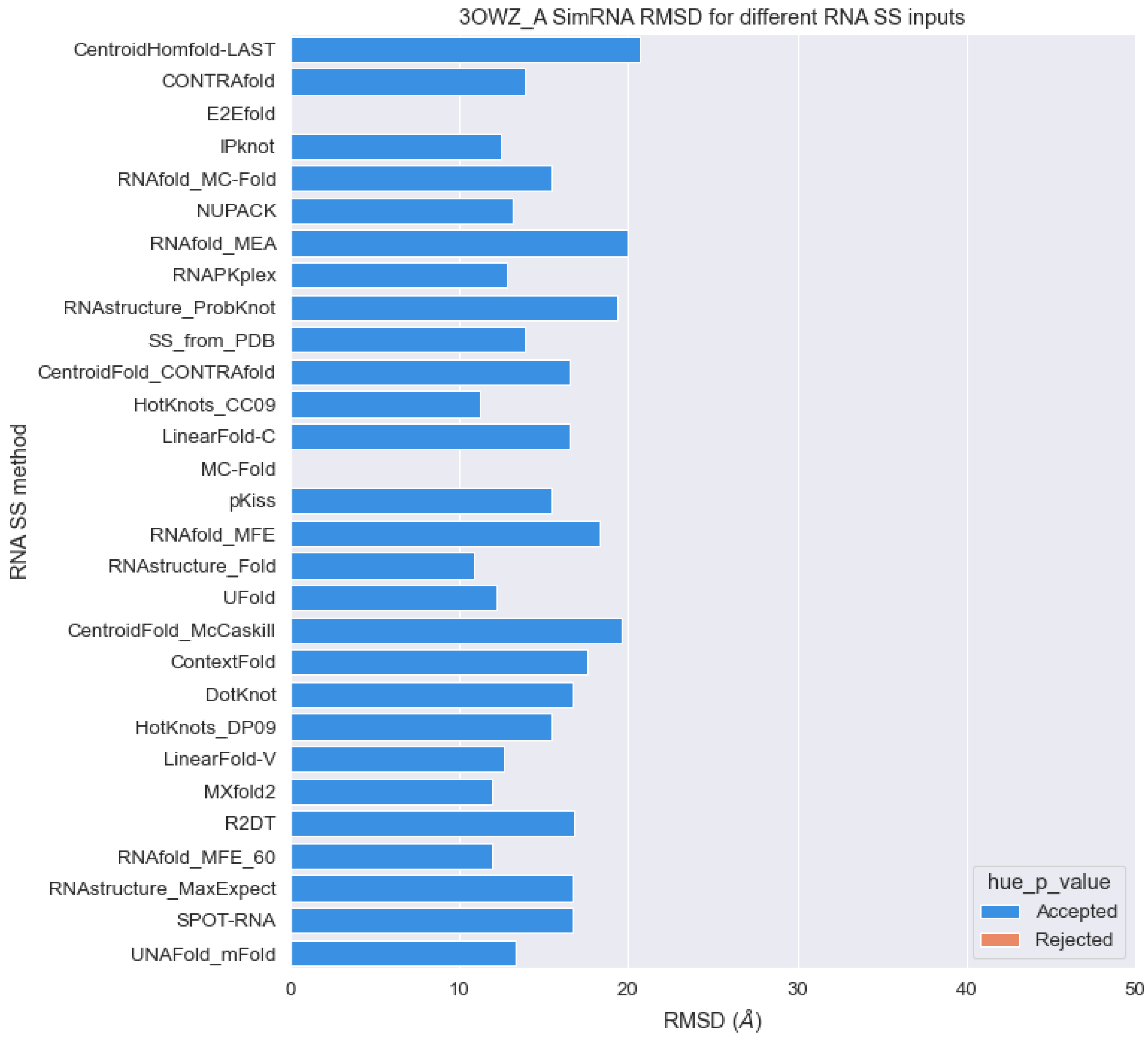


**Figure S88.** RMSD values of predicted structures for PDB 3OWZ sequence with different RNA secondary structures as constraints with SimRNA package (orange: p-value > 0.01, blue: p-value < 0.01).

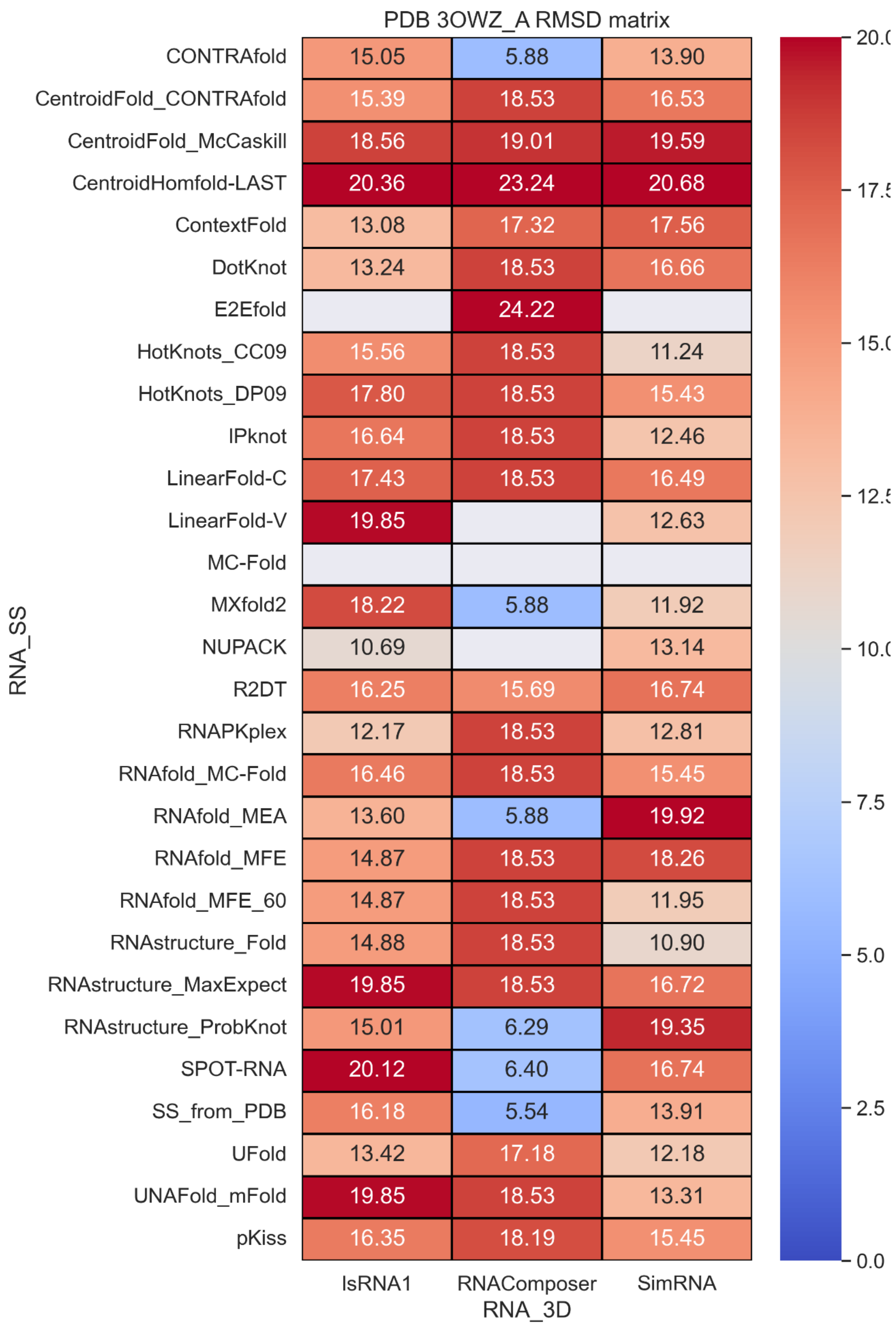


**Figure S89.** RMSD values with different RNA secondary structure methods and RNA 3D methods.

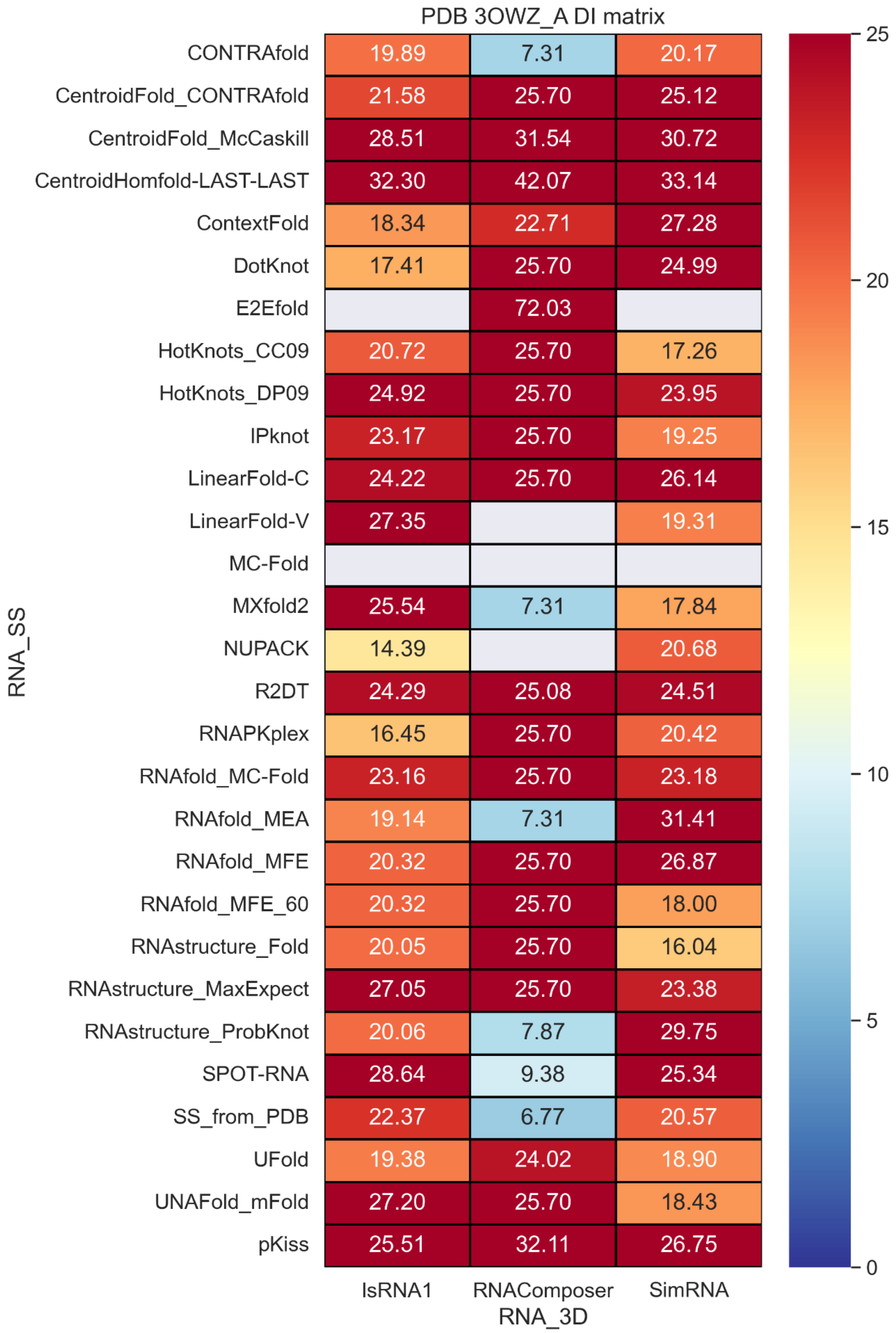


**Figure S90.** Deformation index values with different RNA secondary structure methods and RNA 3D methods.

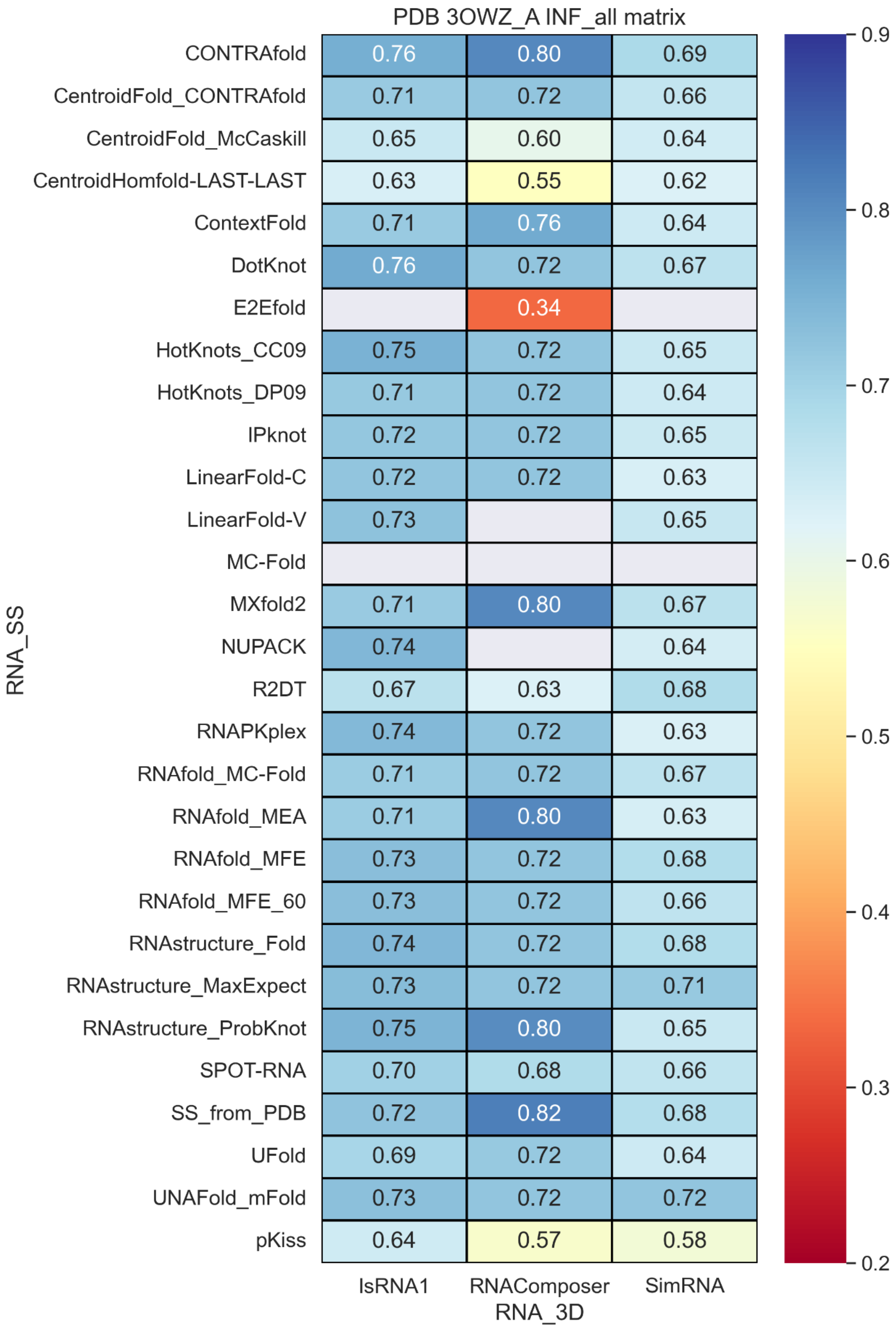


**Figure S91.** Inf_all values with different RNA secondary structure methods and RNA 3D methods.

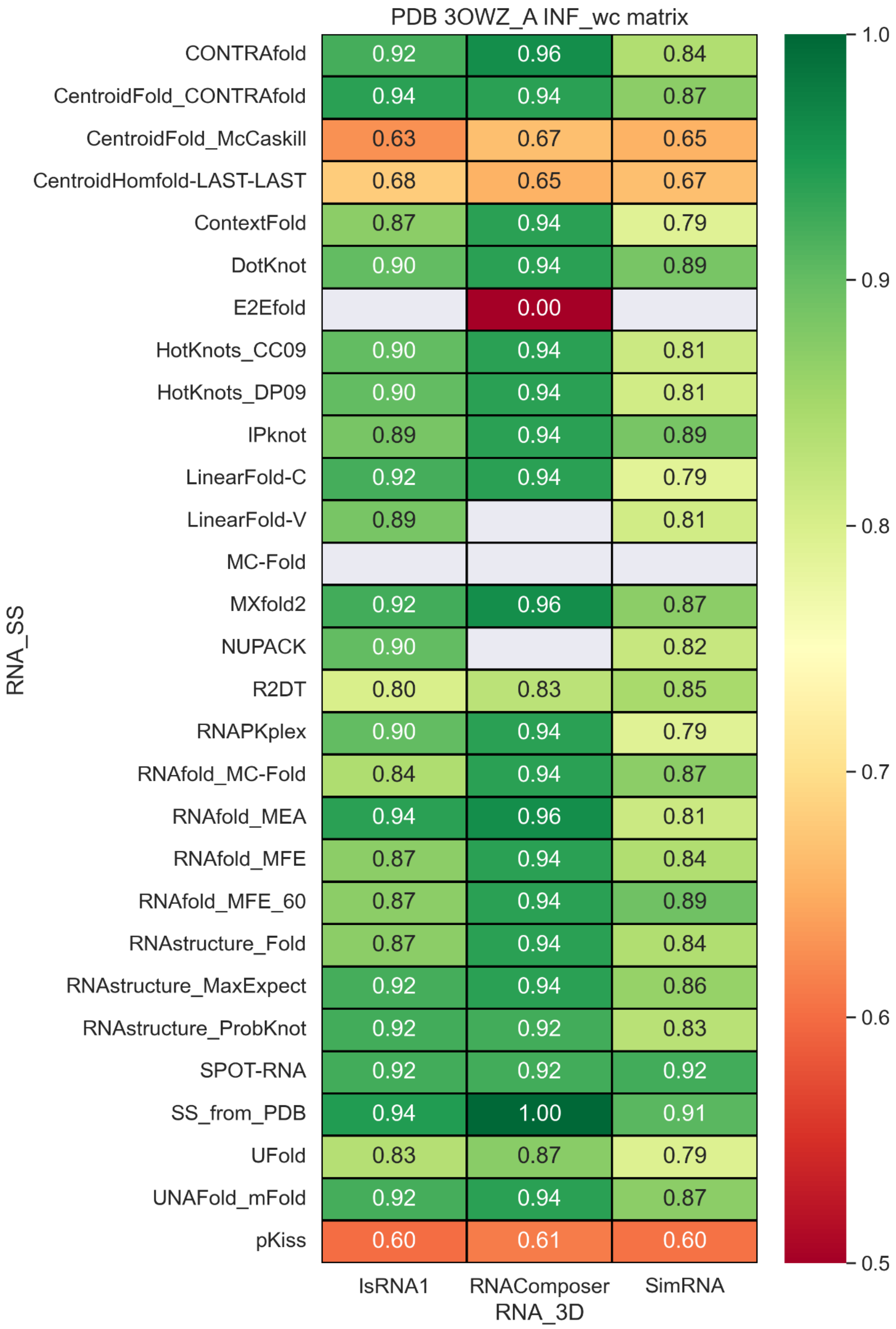


**Figure S92.** INF_wc values with different RNA secondary structure methods and RNA 3D methods.

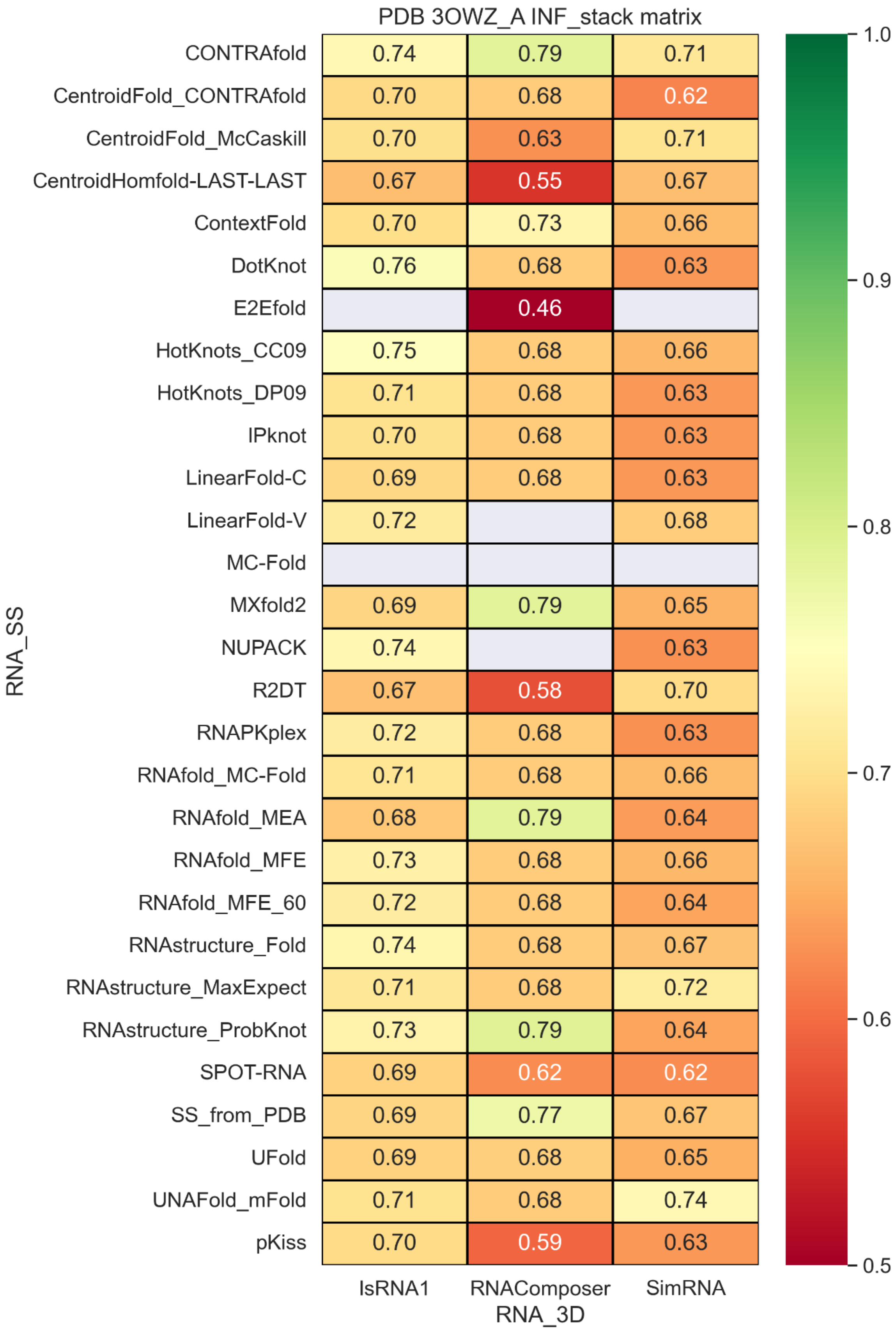


**Figure S93.** INF_stack values with different RNA secondary structure methods and RNA 3D methods.

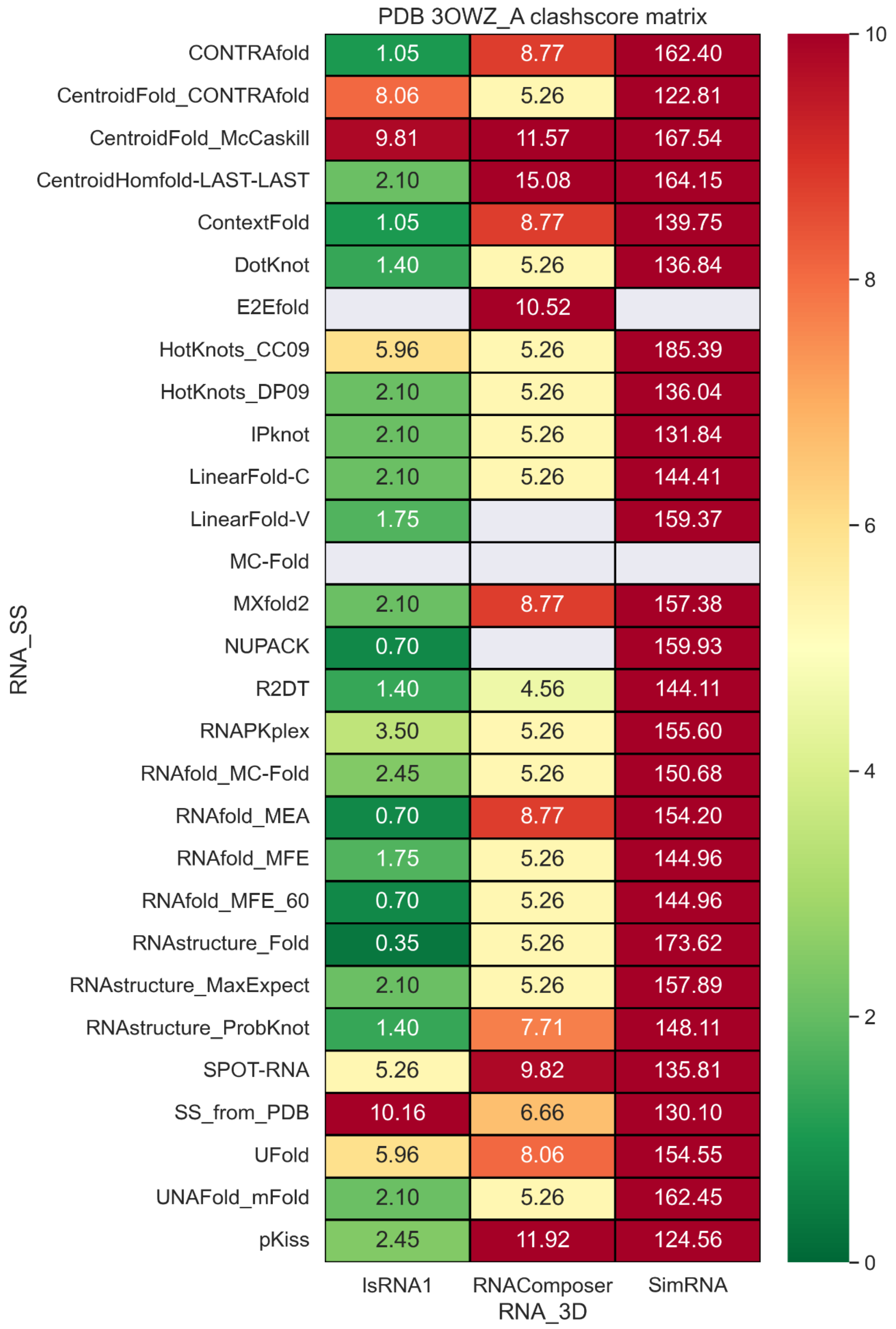


**Figure S94.** Clashscore values with different RNA secondary structure methods and RNA 3D methods.

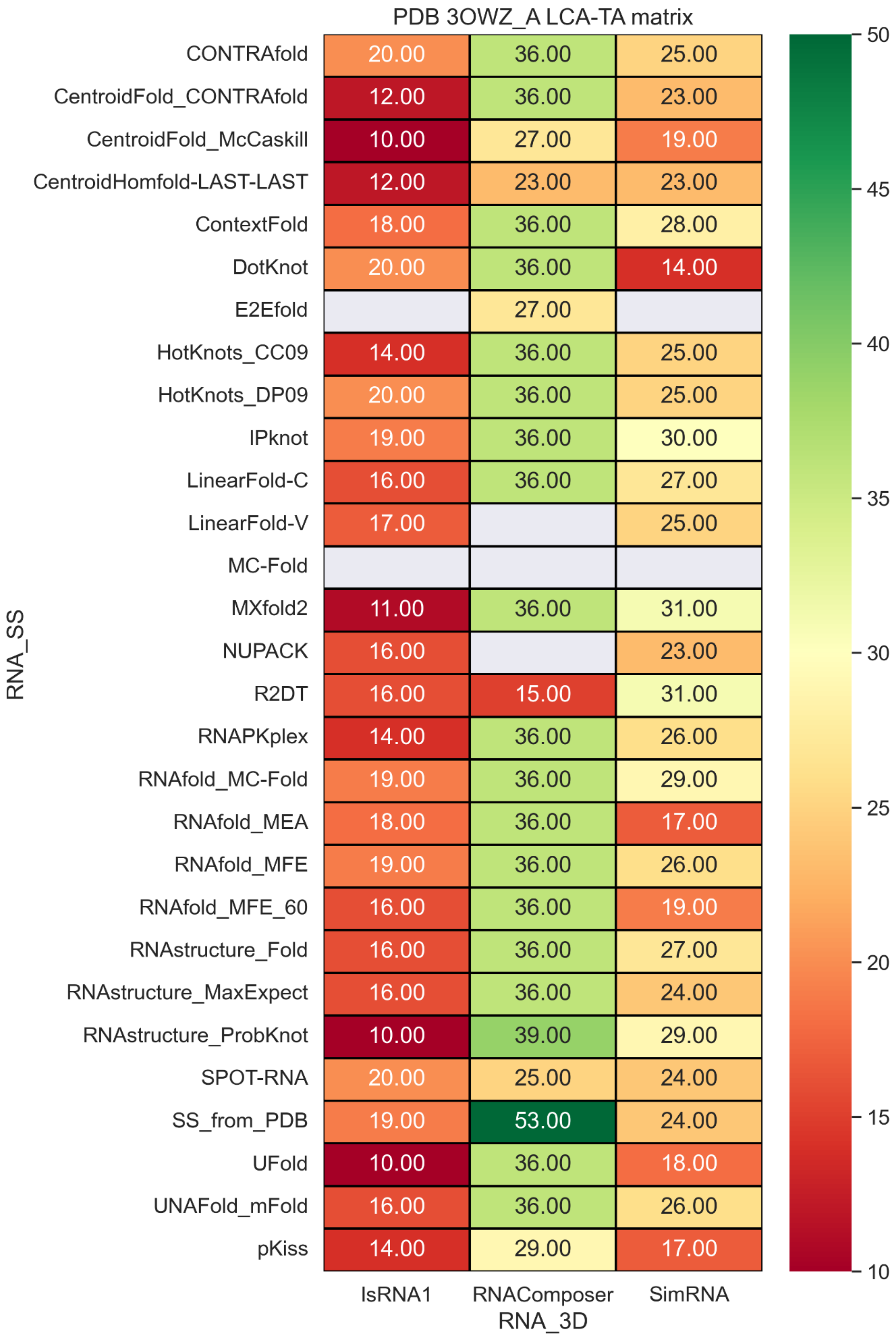


**Figure S95.** LCS-TA values with different RNA secondary structure methods and RNA 3D methods.

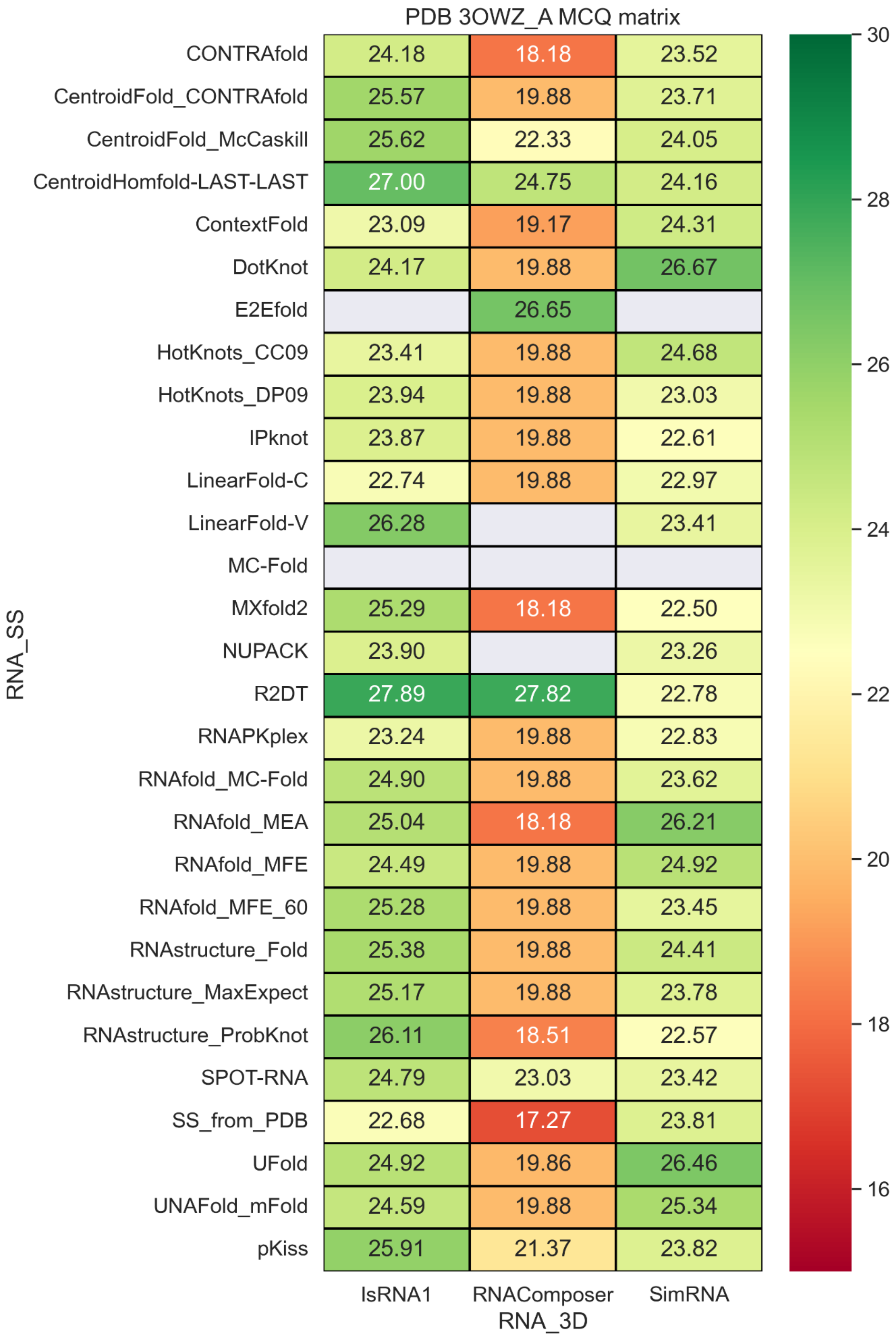


**Figure S96.** MCQ values with different RNA secondary structure methods and RNA 3D methods.
